# Explainable AI identifies recombination and chromatin environment as key predictors of subgenome evolution in maize and *Brassica*

**DOI:** 10.64898/2026.07.13.738278

**Authors:** Layla A. Schuster, Beibei Liu, Jose Cleydson F Silva, Xi Cheng, Raquel Dias, Meixia Zhao

## Abstract

Polyploidy, or whole genome duplication, reshapes genomes through biased gene loss and regulatory rewiring, yet the drivers of biased fractionation among subgenomes remain unclear. Using maize and *Brassica rapa* as model allopolyploids, we compiled 60 genomic and epigenomic features in maize and 45 in *B. rapa* and constructed supervised machine-learning models to classify genes by subgenome identity. In both species, eXplainable Artificial Intelligence (XAI) approaches identified recombination rate as the most influential and highly interconnected feature despite nonsignificant mean differences between subgenome groups. Chromosome location, transposon density, and chromatin-associated features, including the proximity of accessible chromatin regions to genes and active histone marks such as H3K9ac and H3K27ac, consistently ranked among the top contributors to subgenome classification. XAI-derived co-variation and interaction networks further revealed recombination rate as the central node connecting significant features. Together, these results highlight recombination and chromatin environment as major predictors of subgenome divergence in polyploid genomes.

**Teaser:** Recombination and chromatin environment are key predictors of unequal gene loss and subgenome dominance after polyploidy.

## Introduction

Polyploidy, the presence of more than two complete sets of chromosomes resulting from whole genome duplication (WGD), is a prevalent and recurrent phenomenon in eukaryotic evolution, particularly in flowering plants (*1–3*). By expanding the amount of genetic material, WGD generates additional gene copies and novel genomic variation that drives physiological changes, enhanced stress tolerance, phenotypic diversity, evolutionary innovation, and species diversification (*4–8*).

Polyploidy can arise through genome duplication within a species (autopolyploidy) or through hybridization between diverged species followed by genome doubling (allopolyploidy), although many polyploid genomes fall along a continuum between these two extremes (*9–13*). In allopolyploidy, interspecies hybridization often triggers genomic instability, or “genomic shock”, due to incompatibilities between the genetic and epigenetic landscapes of the parental genomes (*14*). This instability is typically resolved through rediploidization, a process involving rapid and extensive genetic and epigenetic changes marked by gene loss, chromosomal rearrangements, divergence in gene expression and function, activation and proliferation of transposable elements (TEs), and alterations in epigenetic states such as changes in small RNAs, DNA methylation, and accessible chromatin regions (ACRs) (*15–21*).

Gene loss following WGD is often driven by functional redundancy, in which one copy of duplicated genes becomes dispensable and is preferentially lost from one of the parental subgenomes, a process known as fractionation (*22, 23*). In many paleopolyploids, including *Brassica rapa* (*24*) and maize (*Zea mays*) (*16*), such fractionation is unequal, with one subgenome consistently losing more genes than the other following WGD and diploidization. For example, in maize, which experienced a WGD approximately 11.4 million years ago (MYA) (*25*), only 39% of the original duplicated gene pairs remain in the current maize genome (*16, 19, 20*). Among the singleton genes, 70% are in the dominant (under-fractionated) subgenome (maize1) and 30% reside in the recessive (over-fractionated) subgenome (maize2), demonstrating clear biased fractionation (*16, 19, 20*). Even among gene pairs in which both copies persist, copies in the recessive subgenome exhibit relaxed selection, lower levels of gene expression, higher rates of TE accumulation, and higher levels of small interfering RNAs (siRNAs) and DNA methylation near genes (*16, 19, 26, 27*). These genes are more likely to be deleted eventually because of their reduced contribution to overall function and fitness (*28*).

A similar pattern is observed in *B. rapa*, which underwent a whole genome triplication (WGT) approximately 13∼17 MYA (*29*), resulting in three subgenomes: the dominant, least fractionated (LF) subgenome and the more fractionated (MF1) and most fractionated (MF2) nondominant subgenomes (*24, 26*). The LF subgenome preserves about 70% of the genes found in *A. thaliana*, whereas gene retention is considerably reduced in the MF1 and MF2 subgenomes, which retain 46% and 36% of the corresponding genes, respectively (*24*). These pronounced differences in gene retention between LF and the MF subgenomes have led to the proposal of a two-step model for the *Brassica* WGT event, in which the MF1 and MF2 progenitor genomes merged prior to the incorporation of the LF genome (*24, 30, 31*). The recessive subgenomes (MFs) accumulate higher levels of 24-nt small RNAs and TEs in the flanking regions of retained genes, likely contributing to the lower expression of these genes relative to their homoeologs in the dominant subgenome (LF) (*17, 18*). Because 24-nt small RNAs trigger DNA methylation, this pattern corresponds to elevated methylation, particularly in the CHH (H = A, T, or C) context, in the upstream and downstream regions of genes located in the recessive genomes of both maize and *B. rapa* (*19, 32*). Interestingly, in monkeyflower, CHH methylation levels decrease within TEs and near genes in first-generation hybrids and are subsequently repatterned differently between the dominant and recessive subgenomes in the natural allopolyploid (*33*). These observations led to the TE load hypothesis, which proposes that differences in TE abundance, distribution, and epigenetic silencing between diploid progenitors are expected to be inherited in polyploids and play a decisive role in establishing subgenome dominance by driving early expression bias (*18, 33, 34*). Consistent with this, in some other allopolyploids or many autopolyploids where diploid progenitors have similar TE densities and methylation profiles, subgenome dominance and biased fractionation may not emerge (*19, 35–37*). However, the TE load theory has been challenged in synthesized *Brassica* allotetraploids, where the expected negative association between TE load and subgenome dominance is not observed, suggesting that methylated TEs near genes are not sufficient to initiate the biased gene expression that defines subgenome dominance (*38*). These findings also raise the possibility that differences in TE abundance and distribution may be a consequence, rather than a cause, of biased fractionation (*39*).

In addition to the roles of small RNAs and DNA methylation in biased gene expression, chromatin environment and *cis*-regulatory elements have emerged as important determinants of the divergence and evolution of duplicated genes (*20, 40*). In maize, the less fractionated subgenome tends to exhibit higher chromatin accessibility, indicating more open chromatin upstream of retained homoeologs compared with the more fractionated genome (*20, 32*), likely contributing to higher levels of transcription. Similarly, a greater number of ACRs has been observed in the dominant A subgenome than the recessive B/C/D subgenomes of octoploid strawberry (*21*). In cotton, the larger A-genome shows wider average nucleosome spacing in diploids and homoeologs with higher expression levels exhibit greater promoter accessibility than their homoeologous counterparts (*40*). Chromatin accessibility is also positively correlated with several active histone modifications, including H3K4me3, H3K36me3, H3K27ac, and H3K9ac, as well as the repressive mark H3K27me3 (*41*). However, no significant correlation between H3K27me3 and subgenome expression is observed in polyploid wheat (*42*).

Beyond these chromatin-based mechanisms, polyploid genomes also undergo extensive structural remodeling that further contributes to biased gene retention and subgenome divergence. After WGD, recombination plays a major role in shaping genome architecture, as increasing evidence indicates that gene loss often occurs through recombination-mediated DNA deletions rather than gradual pseudogenization (*43*). Such recombinational deletions contribute substantially to gene loss following polyploidization and may reinforce patterns of subgenome differentiation through their effects on duplicate gene retention (*43, 44*). Studies in natural and synthetic polyploids have also demonstrated that meiotic recombination patterns rapidly evolve following polyploidization to stabilize chromosome pairing (*44*). For example, autotetraploid *Arabidopsis arenosa* has evolved strengthened crossover interference to restrict recombination to homologous chromosomes, while allopolyploids such as *Arabidopsis suecica* exhibit suppressed homoeologous recombination and largely diploid-like meiotic behavior (*45*). These observations suggest that, as polyploid genomes stabilize, selection favors mechanisms that reduce inappropriate recombination and promote faithful chromosome pairing, including the suppression of homoeologous crossovers (*45, 46*).

While many studies have explored how differences in diploid progenitor genomes contribute to the establishment of subgenome dominance in allopolyploids, few have focused on identifying the key genomic and epigenomic features and their interactions that most effectively shape subgenome differentiation. In this study, we trained supervised machine learning (ML) models to classify genes as belonging to the dominant or nondominant subgenomes in maize and *B. rapa* using a comprehensive set of genomic and epigenomic features. We then applied eXplainable Artificial Intelligence (XAI) approaches to determine which features are the most associated with these classifications. We found that although recombination rates do not differ significantly different between subgenomes in either species, recombination is consistently the most important and most interconnected feature across all models. In addition, gene chromosome location, TE density, the distance between ACRs and genes, and active histone marks, such as H3K9ac and H3K27ac, are among the top contributors to subgenome classification. Together, our findings suggest that gene-associated open chromatin plays a key role in maintaining subgenome dominance.

## Results

### Machine learning models effectively classify duplicated genes in maize subgenomes

To classify genes by subgenome identity in maize, we trained predictive machine learning models on 60 features that describe various genomic and epigenomic characteristics of the subgenomes (Fig. 1A) and organized them into nine categories: recombination rates, gene location, gene expression, evolutionary distances, GC content, transposable elements (TEs), DNA methylation, histone modifications, and accessible chromatin regions (ACRs) (Fig. 2).

**Fig. 1.**
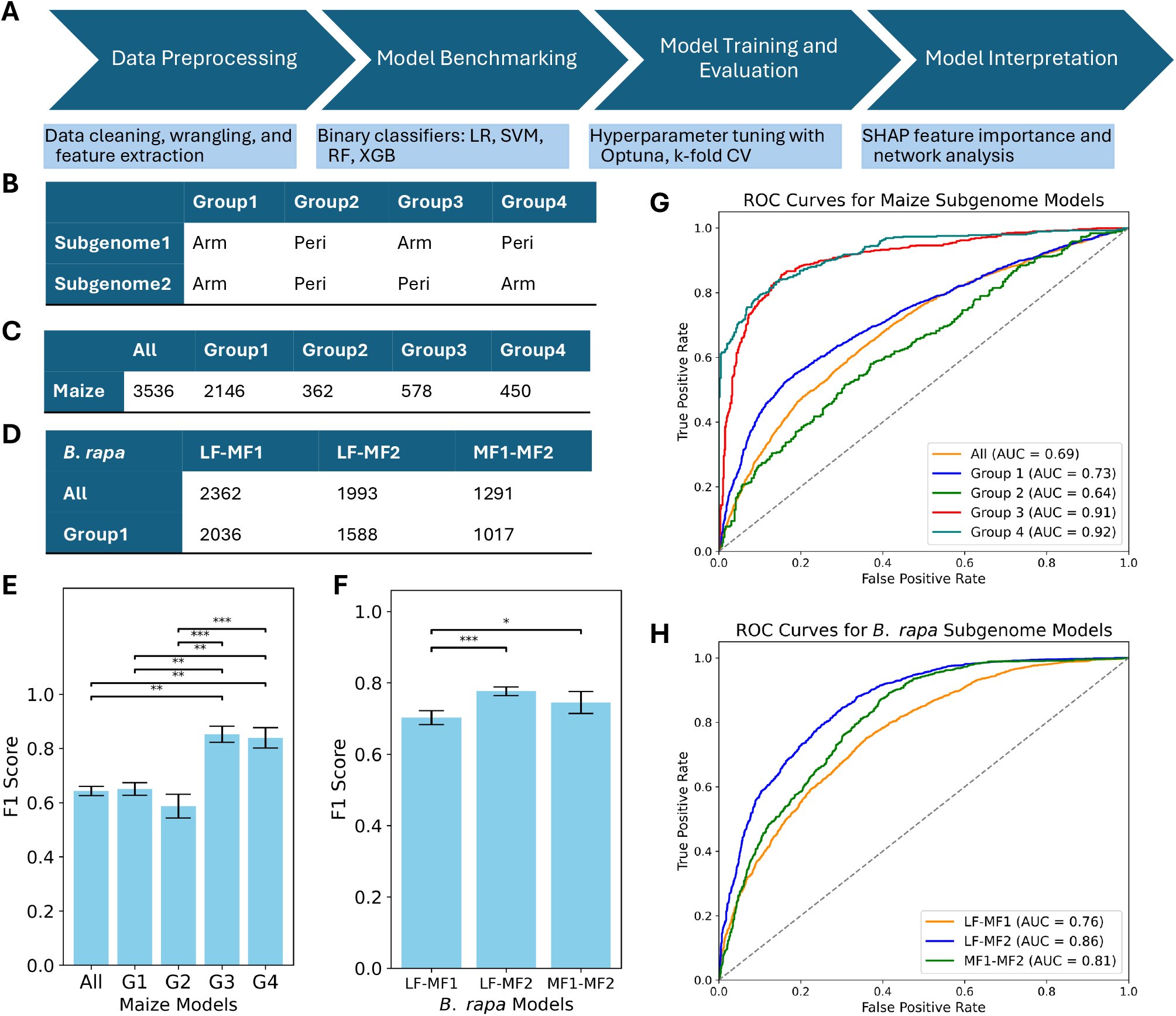
Machine learning models effectively classify duplicated genes in maize and *Brassica rapa* subgenomes. (**A**) Overview of the machine learning pipeline for maize and *B. rapa* models. Binary classifiers listed in model design and selection include logistic regression (LR), support vector machine (SVM), random forest (RF), and XGBoost (XGB). (**B**) Classification of duplicated genes into four groups based on genomic location. (**C**) Number of duplicated gene pairs in maize in the full dataset (All) and in each of the four genomic groups. (**D**) Number of duplicated gene pairs in the three all-gene *B. rapa* subgenome pairs, including the dominant LF (least fractionated) and the recessive MF1 (more fractionated) and MF2 (most fractionated) subgenomes, as well as the group1 subset containing only genes located in chromosomal arms. Mean and standard deviation of F1 scores across training folds for maize (**E**) and *B. rapa* (**F**) models (*** *P* ≤ 0.001, Mann-Whitney U test). Maize models include a binary classifier trained on the full dataset (All) and on genes in group1 (G1), group2 (G2), group3 (G3), and group4 (G4). ROC/AUC curves for five maize (**G**) and three *B. rapa* (**H**) models are shown. The gray dashed line indicates the performance of a random classifier (AUC = 0.5).

**Fig. 2.**
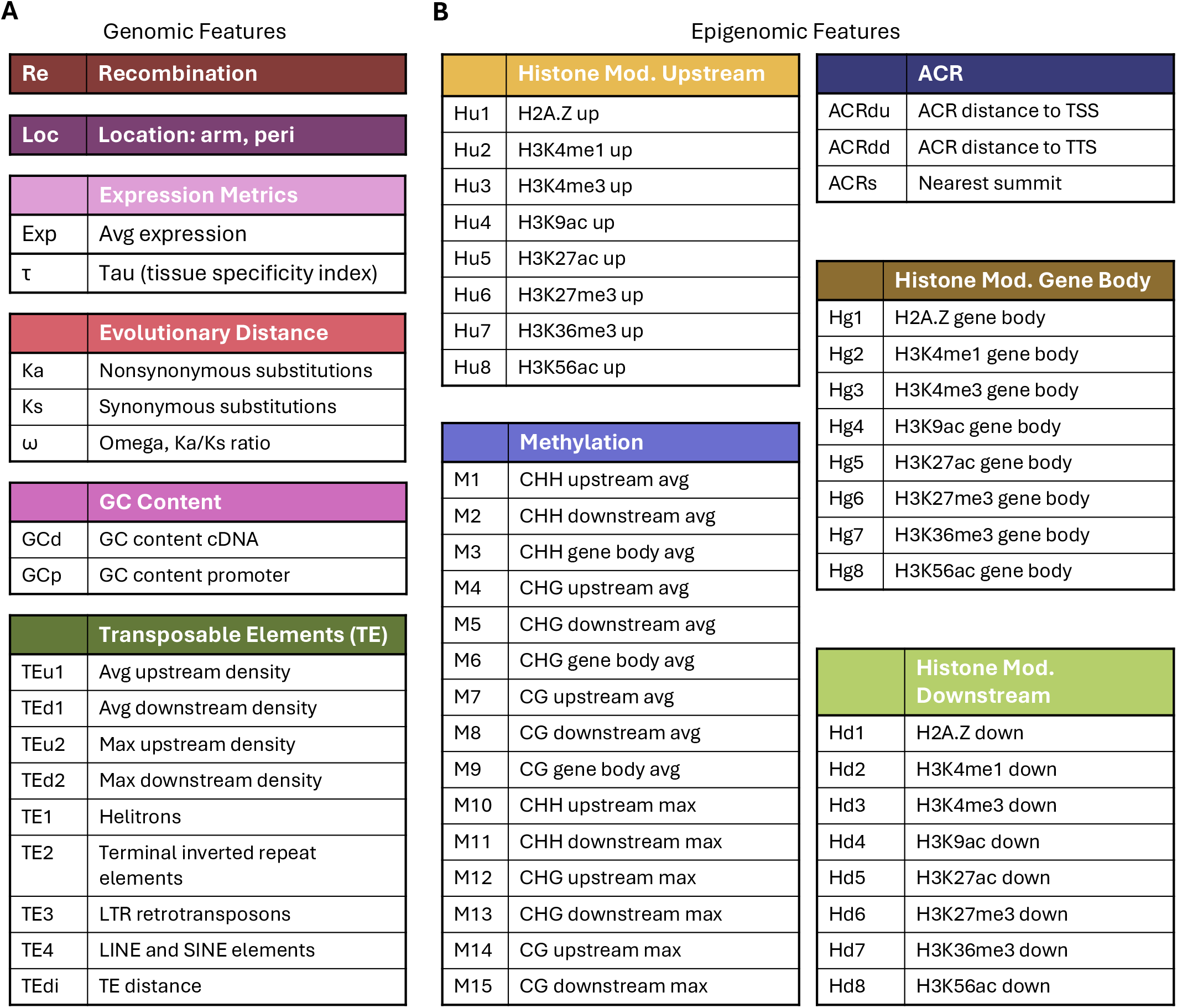
Full set of features used to train maize and *B. rapa* models. (**A**) Genomic features. (**B**) Epigenomic features. Maize models were trained using all 60 features, except for the location feature in groups 1-4, whereas *B. rapa* models were trained on all features excluding histone modifications H(u/g/d)1, 2, 4, 7, 8.

Our previous analysis performed genome-wide comparisons of genomic and epigenomic features between maize subgenomes across distinct chromatin environments, finding that the location of maize1 genes in chromosomal arms is pivotal for maintaining subgenome dominance (*20*). Following this, we separated 3,536 duplicated gene pairs in maize into four groups: group1 (M1_arm; M2_arm: 2,146 gene pairs, with both genes in chromosomal arms), group2 (M1_peri; M2_peri: 362 gene pairs, with both genes in pericentromeric regions), group3 (M1_arm; M2_peri: 578 gene pairs, with maize1 genes in chromosome arms and maize2 genes in pericentromeric regions), and group4 (M1_peri; M2_arm: 450 gene pairs, with maize1 genes in pericentromeric regions and maize2 genes in chromosome arms) (Fig. 1B and C) (*20*).

As an initial analysis, we compared distributions of feature values between subgenomes using Mann-Whitney U tests (MWU) with Benjamini-Hochberg false discovery rate (FDR) correction and quantified the magnitude of differences using the rank-biserial correlation coefficient as a measure of effect size, with features ranked by absolute effect size. In the full maize1-maize2 gene set, 34 features were significantly different between subgenomes, compared to 9 features in group1 and 0 features in group2 (Fig. S1 and Data S1). In groups 3 and 4, where maize1 and maize2 homoeologs are located within the contrasting chromosomal locations, recombination rate showed the largest effect size in both groups by a substantial margin, approximately 2.4× and 6.2× the effect size of the next ranked feature in groups 3 and 4, respectively (Fig. S2), with its direction of effect reversed between groups, consistent with the inverse locations of the duplicated gene pairs. In the full dataset, three features with the largest absolute value effect sizes were average gene expression (Exp), nonsynonymous substitutions (Ka), and H3K9ac in gene bodies (Hg4), while in group1 these were H3K9ac upstream of genes (Hu4), Ka, and Exp (Fig. S1). No features were significantly different in group2 (Fig. S1C), consistent with prior evidence that subgenome bias is largely absent when both homoeologs are in pericentromeric regions (*20*).

Next, we quantified associations among features within each dataset by computing Spearman rank correlation coefficients across all features pairs (Figs. S3-S7 and Data S2). The strongest pairwise correlations were consistently within feature categories across all datasets, particularly among DNA methylation features (e.g., r ≈ 0.91) and histone modification features (e.g., r > 0.89). As expected for groups 3 and 4 given the broad genomic differences between chromosomal arms and pericentromeric regions, recombination rate showed strong correlations with many other features, predominantly with DNA methylation features in group3 and with ACR features in group4. Interestingly, however, recombination rate showed no strong correlations with other features (|r| ≥ 0.3) in groups 1 and 2, where both homoeologs share the same chromosomal context.

For our investigation, we assembled five maize1-maize2 paired feature sets corresponding to the full maize1-maize2 gene set and four subgroups, with each used to train its own model to classify genes as belonging to the maize1 or maize2 subgenome (Fig. 1B). Following data preparation, class balance was preserved across all groups for model training. For the model trained on the full maize1 and maize2 gene set, all 60 features were included. For the subgroup models (groups 1-4), the gene location feature was excluded because it is the feature used to separate the duplicated genes into four groups (*20*). Model benchmarking was performed to evaluate classification performance across four different model architectures: logistic regression (LR), support vector machine (SVM), random forest (RF), and XGBoost (XGB). In pairwise comparisons of per-fold F1 scores (the harmonic mean of precision and recall, summarizing model performance by balancing false positives and false negatives) using two-sided Wilcoxon signed-rank tests for the all, group1 and group2 benchmark models, XGB significantly outperformed all other architectures in the all and group2 datasets, and significantly outperformed LR and SVM but not RF in group1 (Figs. S8-S11 and Table S1). Hence, XGBoost (XGB) was selected as the final model architecture. Following hyperparameter optimization with Optuna (Tables S2 and S3), final XGB models were evaluated using stratified 10-fold cross-validation (CV). Mean F1 scores across folds, ranked from lowest to highest, were 0.59 (group2), 0.64 (all), 0.65 (group1), 0.84 (group4), and 0.85 (group3) (Figs. 1E and S12), which is mostly consistent with the ranking of corresponding AUC (Figs. 1G and S13).

### Recombination drives predicted genomic dominance in maize

To better understand the contribution of each feature to the model’s predictions and to compare feature importance profiles across models, we applied SHAP (SHapley Additive exPlanations) analysis to the five maize models. SHAP is an XAI method based on collaborative game theory that assigns each feature a value representing its contribution to a given prediction (*47*). These values are calculated for each feature and each individual gene, providing both a global view of feature importance (via aggregated SHAP values) and a gene-specific view of how each feature influences individual predictions. A positive SHAP value indicates that a feature increases the model’s confidence in assigning a gene to a particular class, while a negative SHAP value indicates that the feature decreases the model’s confidence in that assignment. Because each SHAP value reflects a feature’s contribution toward predicting one specific class, the direction of that contribution depends on which class is designated as positive. In the maize models, maize1 serves as the negative class and maize2 as the positive class; thus, negative SHAP values push model predictions towards maize1 while positive SHAP values push them towards maize2.

To quantify global feature contributions, we separately summed positive and negative SHAP values, representing the aggregated contributions of each feature toward maize2 and maize1, respectively. Intriguingly, across all maize models, recombination rate was consistently the most influential feature. In the full maize1-maize2 gene set model, in addition to recombination rate, other top features included gene chromosomal location, ACR distance to transcription start site (TSS) (ACRdu), Ka, and average gene expression (Exp) (Figs. 3A and S14). In the group1 and group2 models, ACRdu and several open chromatin features, including upstream H3K9ac (Hu4) and H3K36me3 (Hu7) and gene body H2A.Z (Hg1), ranked among the top five features, though the specific features and their rankings differed between groups (Figs. 3, S15, and S16). In groups 3 and 4, recombination was the dominant feature, as expected given the broad genomic differences between chromosomal arms and pericentromeric regions (Figs. S17 and S18).

**Fig. 3.**
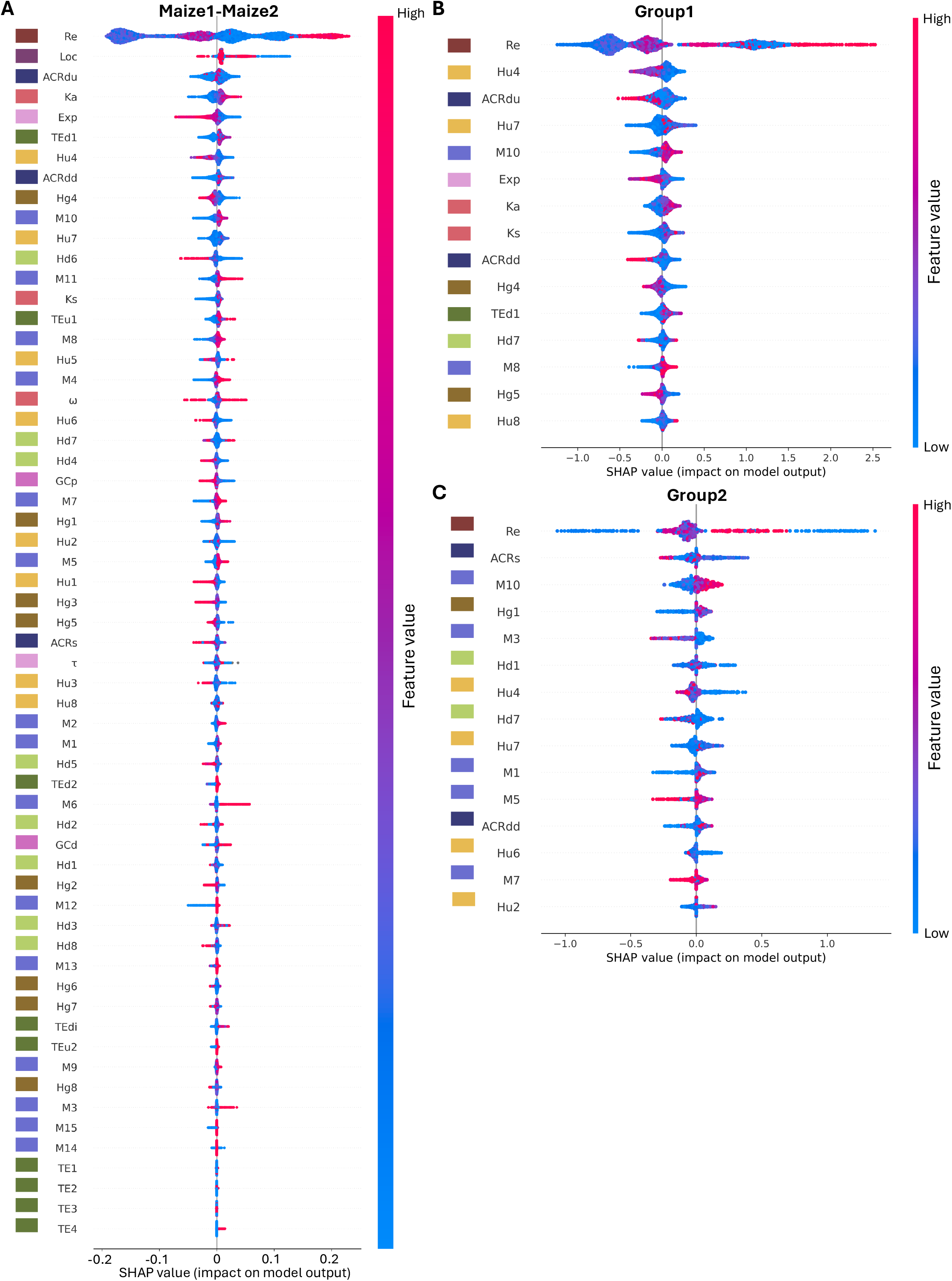
Recombination is the most influential feature in shaping subgenome dominance in maize. SHAP summary beeswarm plots for maize subgenome pair comparisons where SHAP values are shown for maize1-maize2 (all features, A), group1 (top 15 features, B), and group2 (top 15 features, C), computed across all test-fold samples (genes) from the final XGBoost models. Features are ranked by mean absolute SHAP value (highest at top). Each point represents one gene; horizontal position indicates the SHAP value (impact on model output) and point color encodes the raw feature value (red = high, blue = low). Positive SHAP values favor maize2; negative values favor maize1.

Given that recombination rate was identified as the most influential feature across all maize models, we examined its correlation with each of the other features individually (Figs. S19-S23 and Data S2). In the full gene set, recombination rate was strongly positively correlated with chromosome location (r = 0.63), consistent with the well-established association between chromosomal arm position and elevated recombination rates. In contrast, recombination rate showed no strong correlations with any other feature (|r| ≥ 0.3) in groups 1 and 2, where both homoeologs share the same chromosomal context.

To further explore the influence of important features on model predictions, we visualized feature importance using SHAP beeswarm plots (Figs. 3 and S24-S32). While the aggregated SHAP profiles reveal the overall directional influence of each feature and the magnitude of its total contribution across the dataset, the beeswarm plots complement this by showing how individual feature values relate to SHAP magnitude at the level of individual genes, making it possible to identify whether a feature’s effect on classification is monotonic or nonlinear. For features with a monotonic effect on the model’s output, high or low feature values are associated with either consistently positive or consistently negative SHAP values. Conversely, high or low feature values with both positive and negative SHAP values suggest a nonlinear relationship. Because gene chromosomal location was ranked as the second most influential feature (Figs. 3 and S14), the contrasting chromosomal locations of groups 3 and 4 could disproportionately influence model predictions and biased the ranking of other features. To mitigate this confounding effect, we focused our analysis on groups 1 and 2, where genes are located within the same chromosomal location. In the full, group1, and group2 profiles, recombination rate showed a nonlinear relationship, where both high and low recombination rates are associated with predictions toward either subgenome (Figs. 3, S24, and S25). Notably, the negative SHAP values for recombination in group1 show multiple clusters of varying SHAP magnitude, suggesting that recombination’s influence on subgenome classification depends on interactions with other features rather than acting independently (Figs. 3B and S24). To characterize these clusters, we classified genes into two groups based on their recombination SHAP value (cluster 1, left cluster: -0.80 to -0.40, n = 1,560; cluster 2, right cluster: -0.40 to 0, n = 1,172) and compared their raw feature values between groups using MWU tests with Benjamini-Hochberg correction (Fig. 3B and Data S3). Cluster 1 was significantly enriched for maize1 genes relative to cluster 2 (68.8% vs. 53.9%, Fisher’s exact test, OR = 1.88, *P* = 2.44×10⁻¹⁵). Restricting the analysis to maize1 genes, we identified 13 features that differed significantly between clusters. Compared with cluster 2, cluster 1 genes had downstream ACRs located farther from genes, lower levels of downstream histone modifications, higher DNA methylation, and lower recombination rates, indicating that they reside in a relatively less active chromatin environment despite both clusters consisting of maize1 genes located in chromosome arms. These findings suggest that the clustered SHAP values reflect variation in the broader chromatin and genomic context rather than differences in recombination rate alone, consistent with the context-dependent nature of SHAP values, in which each feature’s contribution is evaluated in the presence of all other features.

Per-label SHAP analysis, in which feature importance is computed separately for maize1 and maize2 individual genes, revealed that the top five features were consistent across labels within each model (Figs. S28-32). In group2, the top five features were identical between labels, suggesting that the model draws on the same genomic signals regardless of the subgenome being classified, despite its overall lower classification performance in this group.

### ACR distance to genes, histone modifications, and DNA methylation contribute to subgenome classification across maize models

Following WGD, genes retained in the maize1 subgenome consistently exhibit higher expression levels than their homoeologs in maize2 (*16, 19*), an expression bias partially explained by differences in chromatin accessibility between subgenomes, with maize1 ACRs found to be more accessible than maize2 in chromosomal arms (*20*). Consistent with this, beyond recombination rate, several feature categories related to chromatin accessibility and modification contributed consistently to subgenome classification in the full maize1-maize2 model. ACR distance to TSS (ACRdu) and ACR distance to transcription termination site (TTS) (ACRdd) features ranked at three and eight (Fig. 3A), respectively, reflecting the well-documented enrichment of accessible chromatin regions near maize1 genes in chromosomal arms (*20*). Active histone modifications, including upstream H3K9ac (Hu4) and upstream H3K36me3 (Hu7), and upstream maximum CHH methylation (M10) also ranked within the top 20% of features. Despite the well-established expression bias between maize1 and maize2, average gene expression ranked 5th in the full gene set and 6th in group1 but fell outside the top 15 in group2, suggesting that other features capture more unique predictive information in these models (Fig. 3).

In group1, Hu4 (2nd), ACRdu (3rd), Hu7 (4th), M10 (5th), and ACRdd (9th) were all within the top 20% of features, consistent with the pattern observed in the full gene set (Fig. 3B). In group2, Hu4, Hu7, ACRdd, and M10 remained within the top 20% of features. In contrast, ACRdu did not rank in the top 15 features; instead, ACR nearest summit values (ACRs) and H2A.Z in gene bodies (Hg1) and downstream regions of genes (Hd1) ranked among the top features, a pattern not observed in group1 or the full gene set (Fig. 3C). DNA methylation features, particularly CHH methylation in upstream regions of genes (M10), were among the top five features in both groups 1 and 2 (Fig. 3B and C). Overall, these findings suggest that features associated with open chromatin environments are important in shaping maize subgenome differentiation.

### Recombination rate is the most highly connected feature in maize co-variation networks

To investigate how features jointly influence model predictions in maize, we computed pairwise SHAP co-variation scores and used them to construct feature co-variation networks (Figs. 4 and S33 and Data S4). The SHAP co-variation score between a pair of features reflects how consistently their individual contributions to model predictions move together: a positive score indicates that when one feature contributes more strongly to a given prediction, the other tends to do so as well, whereas a negative score indicates that the two features tend to vary in opposition in their contributions. Networks were filtered to retain only feature pairs with significant co-variation using a permutation-based empirical null distribution (n = 1,000, α = 0.05). Across all five maize models, recombination rate (Re) was the most highly connected feature, ranking first by node degree in every network (Figs. 4D and S33).

**Fig. 4.**
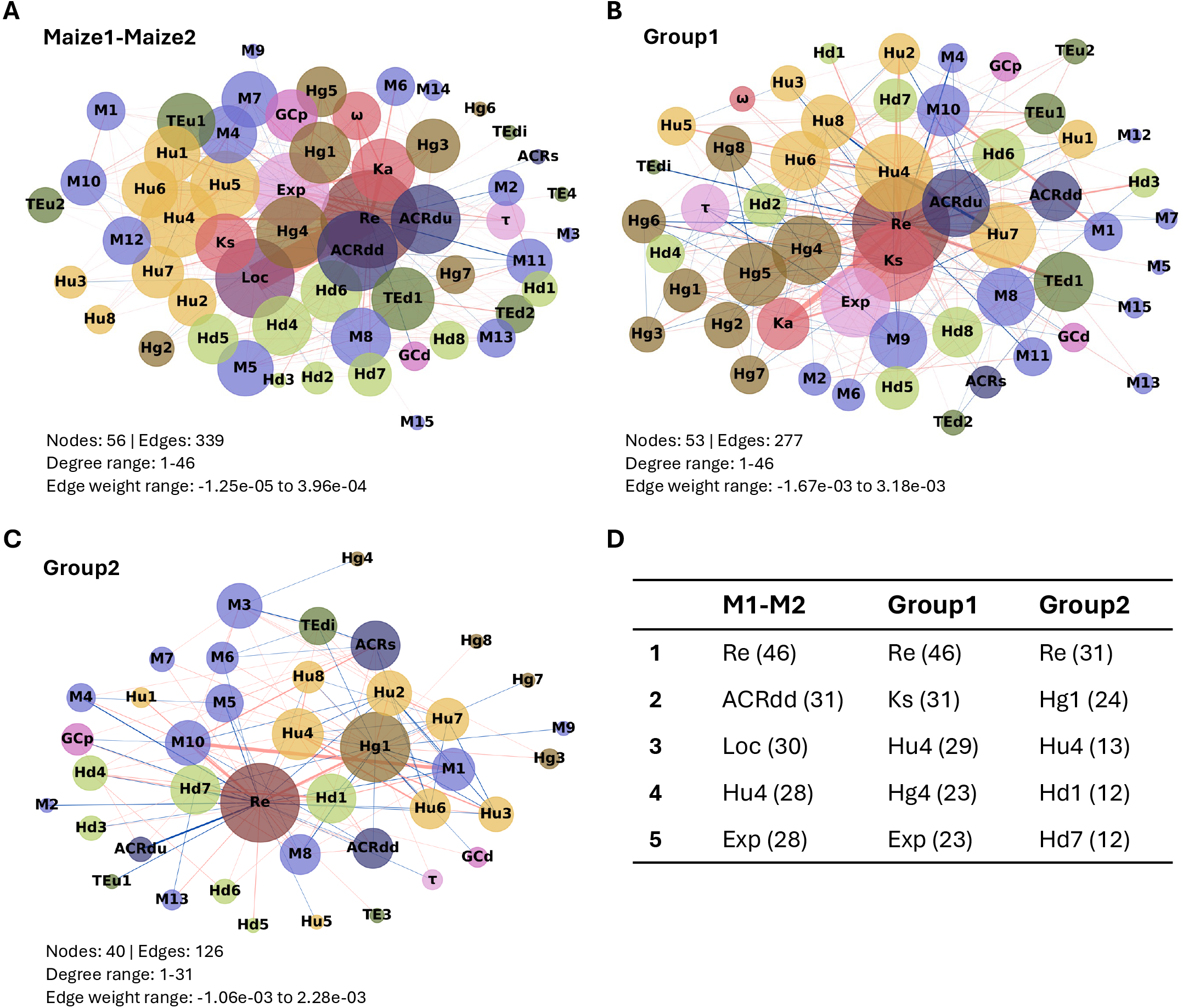
Recombination rate is the most highly connected feature in maize co-variation networks. Networks for maize1-maize2 all genes (**A**), group1 (**B**), and group2 (**C**). Nodes represent features, colored by feature group, and sized by degree (number of significant co-varying partners). Edges connect feature pairs with significant pairwise SHAP co-variation (permutation test, n = 1,000, α = 0.05). Edge color indicates direction: positive co-variation (features vary together in model importance; red) or negative co-variation (features vary in opposition; blue), with edge width scaled to reflect relative co-variation magnitude within each panel. Each panel reports the total number of nodes and edges, the range of node degrees (minimum to maximum), and the range of edge weights (minimum to maximum co-variation score). The top five features by node degree and their corresponding degrees are listed for each network (**D**).

In the full maize1-maize2 gene set network (Fig. 4A), 56 features formed 339 significant co-varying pairs. The top 15 features by node degree spanned eight feature categories, including recombination rate, chromosomal location, ACR features, active histone marks, average expression, downstream TE density, evolutionary rate, and DNA methylation. Recombination rate had the highest degree with 46, followed by ACR distance to the TTS (ACRdd, degree 31), chromosomal location (Loc, degree 30), upstream H3K9ac (Hu4, degree 28), and average expression (Exp, degree 28) (Fig. 4D). The strongest single co-varying pair was Loc and Re (co-variation score = 3.96 × 10⁻⁴), which was approximately seven times stronger than the next strongest pair (Ka-Re, 5.7 × 10⁻⁵), suggesting that chromosomal location and recombination rate are tightly coupled in their joint contribution to subgenome classification across the full gene set. Among the top 15 most connected nodes, co-variation was predominantly positive, forming a broadly coordinated cluster of open chromatin and accessibility features, including ACRdd, ACRdu, Hu4, Hg4, and Hu5, all positively linked to Re. Negative co-variation edges within the top-15 nodes connected Re to several CHH methylation marks (M11 and M2), and linked Loc to Hu4, indicating that chromosomal location and upstream H3K9ac tend to vary in opposition in their contributions to prediction.

In the group1 network (Fig. 4B), 53 features formed 277 significant co-varying pairs. Recombination rate again had the highest degree of 46, followed by synonymous substitution rate (Ks, degree 31), upstream H3K9ac (Hu4, degree 29), gene body H3K9ac (Hg4, degree 23), and average expression (Exp, degree 23) (Fig. 4D). The highest-degree features in group1 were more epigenomically concentrated, with seven histone marks and three methylation features among the top 15, compared to a more heterogeneous feature composition in the all-genes network. The strongest edge in the group1 network was between Ka and Re (3.18 × 10⁻³), approximately 1.6× stronger than the Re-TEd1 edge (average downstream TE density, 2.03 × 10⁻³) and the Re-Hu4 edge (2.02 × 10⁻³). Ks was positively linked to Re and Exp, but negatively linked to several chromatin marks, including Hu4, Hg4, and Hg5, suggesting that genes with higher synonymous divergence between subgenomes tend to show lower SHAP contributions from these histone marks to subgenome classification. The strongest negative edge in the group1 network was between Hu4 and Hu7 (−1.67 × 10⁻³), and Re was negatively linked to several repressive chromatin marks, including Hg6 and M4, as well as TEdi, consistent with the association between high-recombination genomic environments and open, transcriptionally active chromatin.

The group2 network showed a markedly different co-variation structure (Fig. 4C). Group2 retained 40 features forming 126 significant co-varying pairs, fewer than either the all-genes or group1 networks, and the proportion of negative edges was notably higher (41% of all significant edges) compared to group1 (30%) and the all-genes network (18%), indicating a greater degree of opposing co-variation structure in the subgenome prediction of genes in pericentromeric regions. Recombination rate remained the most connected feature (degree 31), followed by four histone marks, including Hg1 (degree 24), Hu4 (degree 13), Hd1 (degree 12), and Hd7 (degree 12) (Fig. 4D). In contrast to group1 and the all-genes network, the top 15 features by degree in group2 were largely restricted to histone modification (7 of 15) and DNA methylation features (5 of 15). The strongest co-variation in the group2 network was not a Re-involving pair, but a positive connection between M1 and M10 (2.28 × 10⁻³), reflecting a tightly co-varying cluster of CHH methylation features. The negative Re-ACRdu edge indicates that for pericentromeric genes, when recombination rate contributes more strongly to subgenome classification, open chromatin proximity tends to contribute in the opposing direction, a pattern that contrasts with the positive Re-ACR co-variation seen in the all-genes and group1 networks. Interestingly, gene body H2A.Z (Hg1) and downstream H2A.Z (Hd1) were negatively connected, suggesting that positional context of H2A.Z, rather than its overall abundance, may help distinguish dominant from recessive genes in pericentromeric regions.

### Feature interactions reveal redundancy among recombination and chromatin features

To complement the co-variation analysis, we constructed SHAP feature interaction networks for each maize model (Figs. 5 and S34 and Data S5). SHAP interaction values measure the degree to which pairs of features jointly influence model predictions beyond what would be expected from their individual contributions alone. A negative interaction score indicates that two features are partially redundant in their combined contribution to classification, whereas a positive score indicates that their combined contribution exceeds the sum of their individual effects. Across models, interaction networks were substantially sparser than their corresponding co-variation networks and negative edges predominated in all three networks (Figs. 4 and 5).

**Fig. 5.**
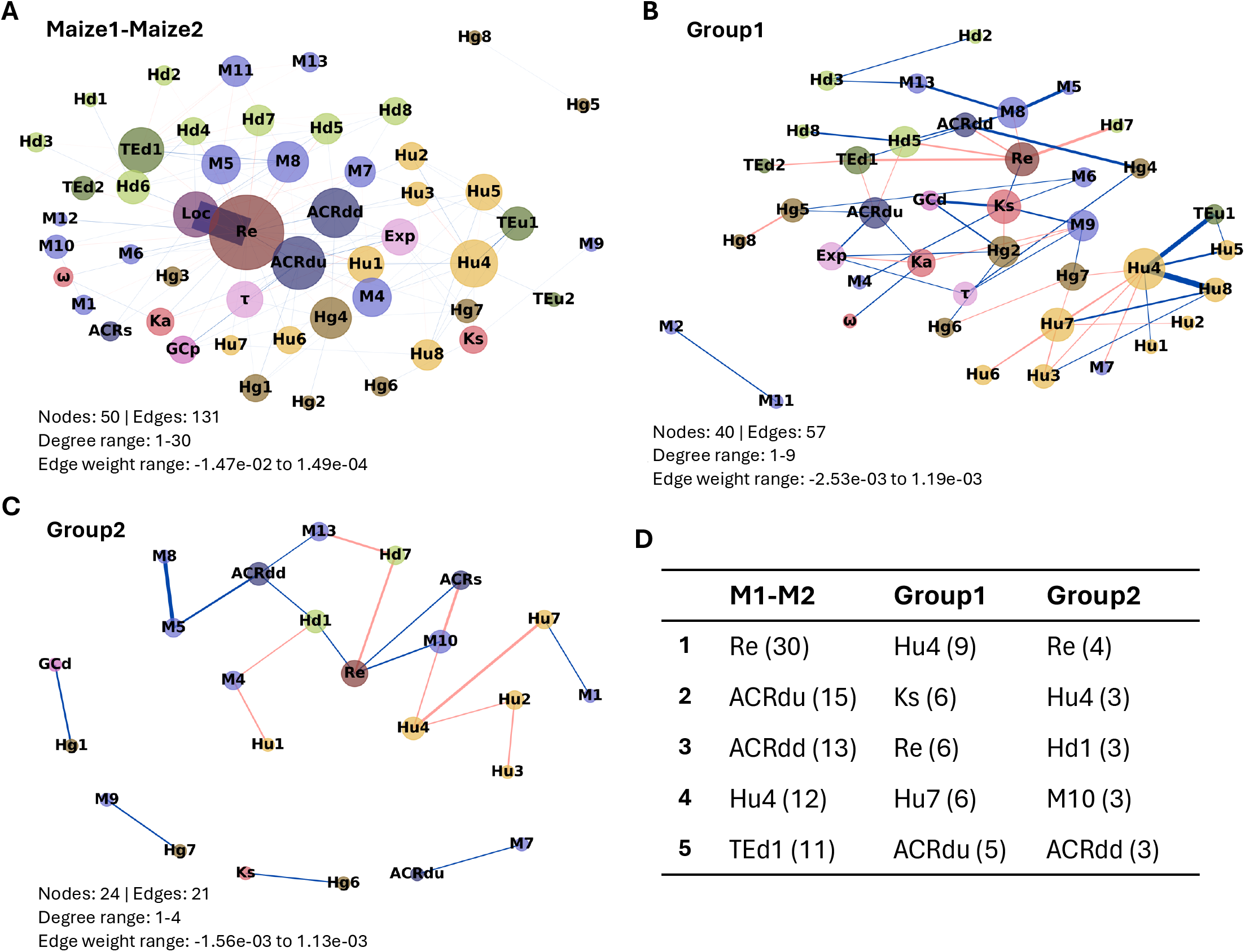
Feature interactions reveal redundancy among recombination and chromatin features. Interaction networks shown for the maize all-genes dataset (**A**), group1 (**B**), and group2 (**C**). SHAP interaction values quantify the extent to which pairs of features jointly influence model predictions beyond the sum of their individual effects. For each feature pair, mean Shapley interaction values were computed across all test-set genes. Edges connect feature pairs whose absolute mean interaction value exceeded a robust effect-size threshold (median + 12 × median absolute deviation, computed from non-zero pairs within each group). Edge color indicates interaction direction: positive (pink) edges denote synergistic interactions, where the combined effect of two features exceeds the sum of their individual effects, whereas negative (blue) edges denote antagonistic interactions, where the combined effect is less than the sum. Node size is proportional to degree (number of retained edges), and node color indicates feature category, as in Fig. 4. The top five features by node degree and their corresponding degrees are listed for each network (**D**).

In the all-genes interaction network (Fig. 5A), 50 features formed 131 significant interaction pairs, of which 88 (67%) were negative. Recombination rate was the most connected feature (degree 30), followed by ACRdu (degree 15), ACRdd (degree 13), and Hu4 (degree 12) (Fig. 5D). The strongest interaction edge was between Loc and Re (−1.47 × 10⁻²), a negative interaction approximately 60× larger in absolute magnitude than the next strongest edge. This finding contrasts sharply with the co-variation network, where Loc-Re represented the strongest positive edge. These opposing signs reflect distinct relationships captured by the two metrics. Loc and Re co-vary positively in their individual contributions to predictions, indicating that when one contributes more to a prediction, the other tends to do so as well. However, their interaction value is negative, indicating that their combined contribution is less than the sum of their individual contributions. This pattern is consistent with partial redundancy, as genes located in chromosomal arms generally have elevated recombination rates. Consequently, once one feature captures this signal, the other provides limited additional predictive information.

In the group1 interaction network (Fig. 5B), 40 features formed 57 significant interaction pairs, of which 34 (60%) were negative. Unlike both the all-genes interaction network and co-variation network, recombination rate was not the most connected feature in group1. Instead, Hu4 had the highest degree (9), whereas Re, Ks, and Hu7 were tied at degree 6 (Fig. 5D). The strongest interactions in group1 were negative, including Hu4-Hu8 (−2.53 × 10⁻³) and TEu1-Hu4 (−1.90 × 10⁻³), suggesting substantial partial redundancy between active histone modification and TE density features in the classification of arm-located gene pairs. The group2 interaction network was the sparsest among the primary models, with only 21 significant edges among 24 features and a maximum node degree of 4 (Fig. 5C). The substantially reduced connectivity in the group2 interaction and co-variation networks suggests that feature interactions play a limited role in the classification of duplicated genes in pericentromeric regions. In the all-genes network, the per-label structure was broadly symmetrical, with Re remaining the most connected feature and Loc-Re representing the strongest negative interaction in both. In contrast, the group1 per-label networks showed greater asymmetry: Re was tied with Hu4 for the highest degree in the dominant maize1 network but was clearly the most connected feature in the recessive maize2 network (degree 14 versus 10) (Fig. S35 and Data S6).

For groups 3 and 4, in the co-variation networks, recombination rate was again the most highly connected feature in both groups (degree 48 in group3 and degree 43 in group4), consistent with the primary group networks, and total edge counts were comparable (149 significant edges in group3 and 117 in group4) (Fig. S33 and Data S4). In contrast, the interaction networks revealed a pronounced asymmetry between the two control groups (Fig. S34 and Data S5). Group3, in which maize1 homoeologs are in chromosomal arms, had 146 significant interaction edges among 51 features. Recombination rate was the most connected feature (degree 39) and ACR related features were also highly connected. Notably, the three strongest interaction edges were all negative and involved Re or ACR features. Despite these strong antagonistic interactions, positive edges slightly outnumbered negative edges overall (52% of significant interactions), distinguishing group3 from the primary interaction networks, where negative interactions predominated. In contrast, group4, in which maize1 homoeologs are in pericentromeric regions, retained only 4 significant interaction edges among six features. This asymmetry in interaction network complexity between the two control groups suggests that positioning maize1 genes within chromosomal arms corresponds to a far richer interaction structure among recombination and chromatin accessibility. Such a pattern is consistent with the role for chromosomal arm environment in maintaining subgenome dominance (*20*).

### Epigenomic and genomic features encode subgenome identity across *B. rapa* subgenomes

To further validate our findings in maize, we applied the same machine learning pipeline in a second polyploid system, *B. rapa* (Fig. 1A), a species that underwent a whole genome triplication (WGT), resulting in a least-fractioned dominant subgenome (LF) and two more-fractionated recessive subgenomes (MF1 and MF2) (*24, 26, 48*). Given this genome structure, we trained three binary classifiers to distinguish subgenomic identity within each subgenome pair (LF-MF1, LF-MF2, and MF1-MF2), using the gene pairs shown in Fig. 1D. Features were organized into the same nine categories used in maize (Fig. 2). However, histone modification data available for *B. rapa* was limited to three histone marks, H3K4me3, H3K27ac, and H3K27me3, compared to eight in maize (*49*), resulting in a final feature set of 45 features per model. To parallel the chromosomal context analysis in maize, we defined a group1 subset for each subgenome pair retaining only gene pairs in which both homeologs are located on chromosomal arms (Fig. 1D). Subsets analogous to maize groups 2-4 were not analyzed due to insufficient gene counts in *B. rapa*.

To compare feature distributions between subgenomes, we performed two-sided MWU tests with Benjamini-Hochberg FDR correction and reported effect sizes as rank-biserial correlation coefficients (Figs. S36 and S37 and Data S1). In the all-genes datasets, 16 and 20 features were significantly different in LF-MF1 and LF-MF2, respectively, whereas no features reached significance in MF1-MF2 after FDR correction (Fig. S36). Across both LF-MF1 and LF-MF2 comparisons, H3K27ac marks (Hg5, Hu5, and Hd5) and H3K4me3 (Hg3, Hu3, and Hd3) marks in gene bodies and their upstream and downstream regions were enriched in LF, while gene body and upstream and downstream H3K27me3 marks (Hg6, Hu6, and Hd6) were enriched in the recessive subgenomes MF1 and MF2. Average expression was also higher in LF, and tissue specificity (τ) was higher in the recessive subgenomes in both comparisons (Data S1). Recombination rate differed significantly only in LF-MF2 (r_rb = +0.095, higher in LF), the subgenome pair exhibiting the greatest degree of fractionation divergence. In the group1 datasets, 13 and 17 features remained significant in LF-MF1 and LF-MF2 group1, respectively, with the same histone modification patterns retained (Fig. S37 and Data S1). In contrast to the all-genes MF1-MF2 comparison, recombination rate was the only significant feature in the MF1-MF2 group1 dataset (r_rb = +0.151, higher in MF1).

Pairwise Spearman rank correlations across all features and between recombination rate and all other features revealed the same patterns as observed in maize (Figs. S38-S49 and Data S2). Recombination rate showed no strong correlations with any other feature (|r| ≥ 0.3) in any of the three group1 datasets. In the all-genes datasets, recombination rate exhibited only a weak positive correlation with chromosomal location (r = 0.26-0.32), with no other features approaching the threshold.

As in maize, we benchmarked four model architectures, LR, SVM, RF, and XGB, and selected XGB as the final model architecture given its superior or equivalent performance across the majority of datasets (Figs. S50-S56 and Table S4). Across both all-genes and group1 models, LF-MF2 achieved the highest classification performance, consistent with the greater degree of fractionation difference between the LF and MF2 subgenomes relative to the other subgenome pairs (*24, 26*)) (Figs. S57-58 and Table S5).

### Recombination and chromatin environment are important in *B. rapa* subgenome dominance

For SHAP analysis of the *B. rapa* models, LF was defined as the negative class and MF1 or MF2 as the positive class for the LF-MF1 and LF-MF2 models, such that negative SHAP values push predictions towards LF classification and positive SHAP values towards MF1 or MF2 classification. In the MF1-MF2 model, MF1 was the negative class and MF2 the positive class. Aggregated directional SHAP profiles confirm recombination rate as the most influential feature across all *B. rapa* models (Figs. S59-S64) and beeswarm plots further reveal a non-monotonic relationship between recombination rate values and their SHAP contributions (Figs. 6 and S65-S70). Beyond recombination rate, chromosome location (Loc) ranked among the most influential features, placing second in the LF-MF2 and MF1-MF2 models and twelfth in the LF-MF1 model (Figs. 6 and S65-S70). Histone modification features, particularly H3K27me3 and H3K27ac, also ranked among the top contributors across all three models. Per-label SHAP analysis, in which feature importance was summarized separately for dominant and recessive subgenome genes, revealed a high degree of concordance between labels across all six models (Figs. S71-S76). For example, in the LF-MF2 model, the top five features for LF genes were Re, Loc, Exp, Hg6, and ACRdd. The same five features were also in the top five for MF2 genes, although their ordering differed slightly (Fig. S72).

**Fig. 6.**
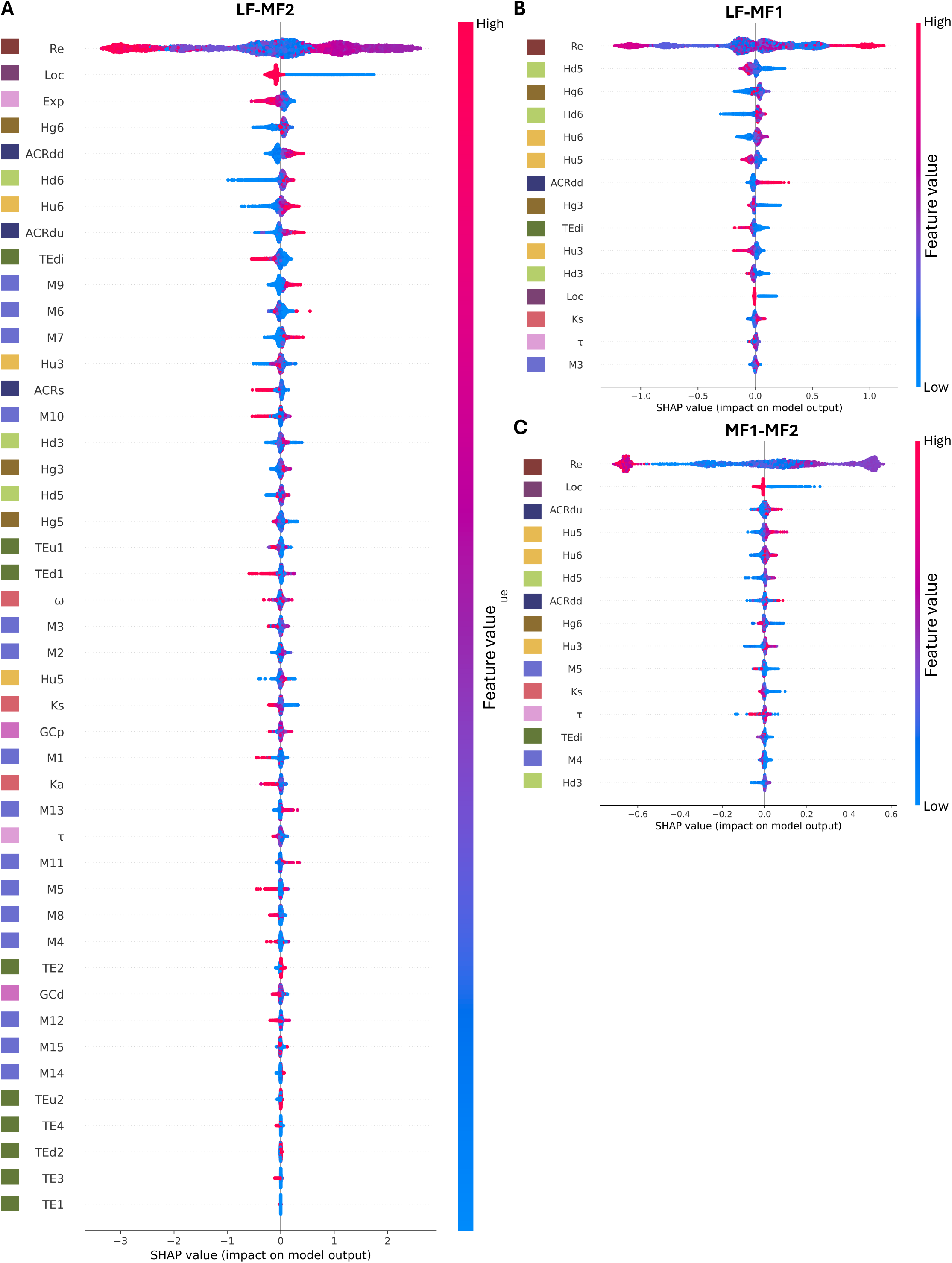
Recombination is the most influential feature in shaping subgenome dominance in *B. rapa*. SHAP summary beeswarm plots for *B. rapa* subgenome pair comparisons where SHAP values are shown for LF-MF2 (all features, **A**), LF-MF1 (top 15 features, **B**), and MF1-MF2 (top 15 features, **C**), computed across all test-fold genes from the final XGBoost models. Features are ranked by mean absolute SHAP value (highest at top). Each point represents one gene; horizontal position indicates the SHAP value (impact on model output) and point color encodes the raw feature value (red = high, blue = low). Positive SHAP values favor the more non-dominant subgenome (MF2 in LF-MF2 and MF-MF2, MF1 in LF-MF1) and negative values favor the more dominant subgenome (LF in LF-MF1 and LF-MF2, MF1 in MF-MF2).

In the LF-MF2 model, recombination rate (Re) was approximately 5.9× more influential than the second-ranked feature, chromosomal location (Loc) based on mean absolute SHAP values, followed by average gene expression (Exp), H3K27me3 in gene bodies (Hg6), and ACR distance to TTS (ACRdd) (Figs. 6 and S66). This pattern is analogous to the maize all-genes model, where Re and Loc also ranked first and second. In the LF-MF1 model, Re was approximately 7.8× more influential than the second-ranked feature, downstream H3K27ac (Hd5). H3K27me3 marks occupied three of the top five positions, gene body (Hg6), downstream (Hd6), and upstream (Hu6). This finding is consistent with both MWU results and SHAP beeswarm directionality, which showed greater H3K27me3 enrichment in, and predictive of MF1, and greater H3K27ac enrichment in and predictive of LF (Figs. 6 and S65). In the group1 models, the top-ranked features for LF-MF1 remained dominated by both H3K27ac and H3K27me3 histone marks, while the LF-MF2 group1 was dominated by H3K27me3 features. This pattern is broadly analogous to maize group1, where active histone modifications also ranked prominently beyond recombination rate, although the specific marks differed.

The MF1-MF2 comparison exhibited a distinct pattern. In the all-genes model, Re was approximately 17× more influential than the second-ranked feature, Loc, followed by ACR distance to TSS (ACRdu), upstream H3K27ac (Hu5), and upstream H3K27me3 (Hu6) (Fig S67). In the MF1-MF2 group1 model, Re was approximately 35× more influential than the second-ranked feature, synonymous substitution rate (Ks), followed by average downstream TE density (TEd1), downstream CG methylation (M8), and upstream H3K27me3 (Hu6) (Fig. S70). In both MF1-MF2 SHAP profiles, the substantial margin between Re and all other features is notable. This pattern is particularly striking given that no features showed statistically significant distributional differences between MF1 and MF2 (Figs. S36 and S37 and Data S1), suggesting that the model captures positional and recombination-associated signals that are not detectable through pairwise distributional comparisons alone.

### Recombination and histone modification features are the most connected nodes in *B. rapa* subgenome co-variation and interaction networks

We constructed co-variation networks for all six *B. rapa* subgenome pair models using the same permutation-based approach applied in maize (Figs. 7 and S77). Recombination rate was the most connected node in the three all-genes networks, with degrees of 31, 40, and 36 in LF-MF1, LF-MF2, and MF1-MF2, respectively, consistent with the maize all-genes result (Figs. 7D and 4D). In both LF-MF1 and LF-MF2 models, Re and Loc formed the strongest co-varying pair. This relationship was considerably more dominant in the LF-MF2 network, where the Re-Loc edge was 3.4× stronger than the next strongest edge, compared with only 1.3× margin in LF-MF1 (Fig. 7D and Data S4). In LF-MF2, the four next strongest edges after Re-Loc also involved Re, further emphasizing recombination rate as the central organizing feature of this network. Beyond Re and Loc, histone modification features, including H3K27me3 in LF-MF2 and both H3K27me3 and H3K27ac marks in LF-MF1, ranked among the most highly connected nodes, consistent with their prominence in the SHAP feature importance analysis (Fig. 6). The MF1-MF2 all-genes network exhibited a distinct structure, with fewer significant edges overall (91, compared to 128 and 136 in LF-MF1 and LF-MF2, respectively), Re-ACRdu replacing Re-Loc as the strongest co-varying pair, and a higher proportion of negative co-variation edges (∼38%, compared to ∼18-28% in the LF-MF1 and LF-MF2 models) (Fig. 7C). Together, these patterns indicate a higher degree of opposing co-variation structure between MF1 and MF2 than between either recessive subgenome and LF.

**Fig. 7.**
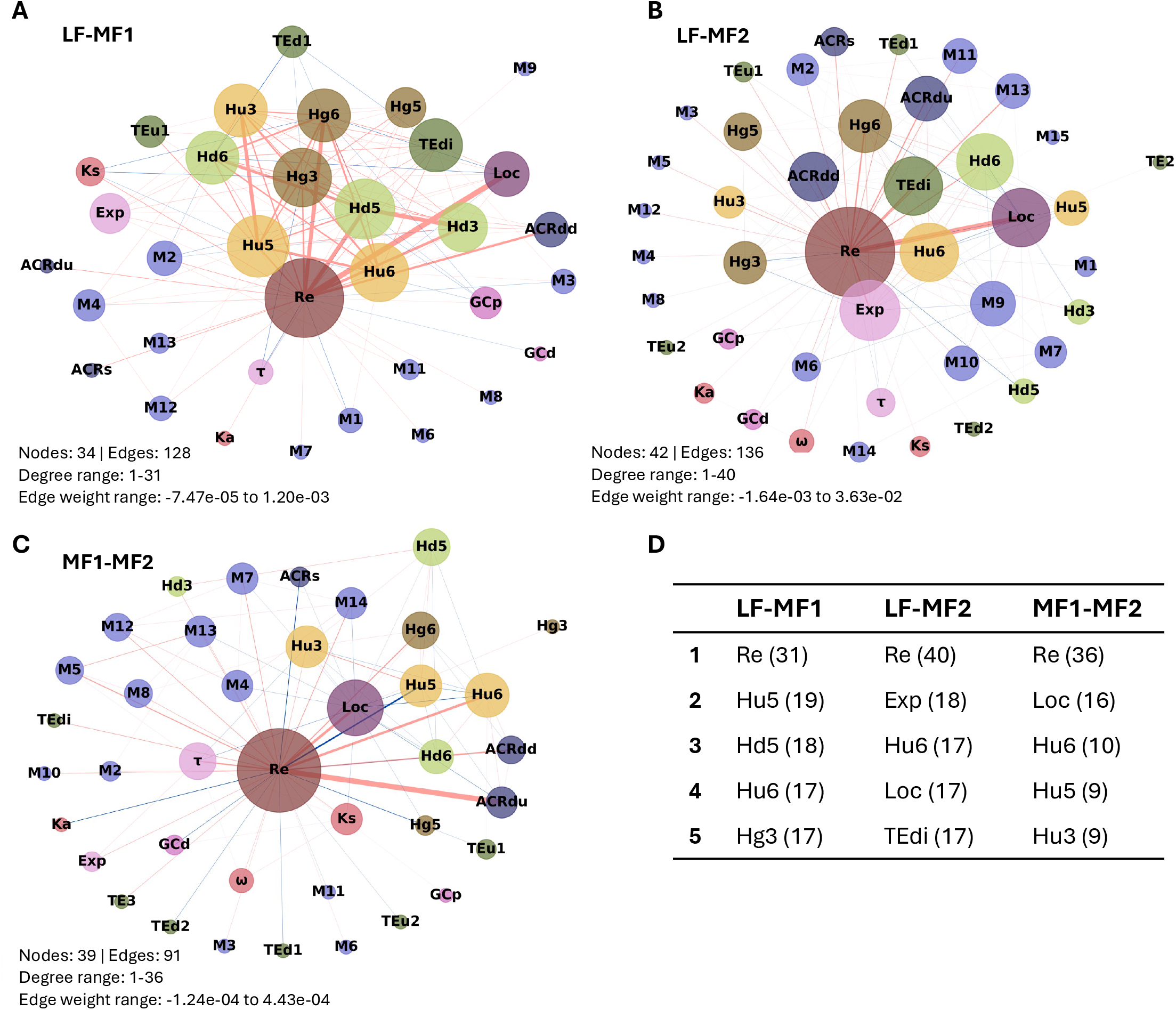
Recombination and histone modification features are the most connected nodes in *B. rapa* subgenome co-variation networks. Networks for LF-MF1 (**A**), LF-MF2 (**B**), and MF1-MF2 (**C**), each using all genes. Nodes represent features, colored by feature group and sized by degree (number of significant co-varying partners). Edges connect feature pairs with significant pairwise SHAP co-variation (permutation test, n = 1,000; α = 0.05). Edge color indicates direction: positive co-variation (features vary together in model importance; red) or negative co-variation (features vary in opposition; blue), with edge width scaled to reflect relative co-variation magnitude within each panel. Each panel reports the total number of nodes and edges and the range of node degrees (minimum to maximum) in the network. The top five features by node degree and their corresponding degrees are listed for each network (**D**).

In the group1 networks, chromosomal location was excluded as the group-defining variable, as in the maize subgroup analysis. Recombination rate nevertheless remained the most connected node in all three networks, with degrees of 32, 37, and 37 in LF-MF1, LF-MF2, and MF1-MF2 group1, respectively (Fig. S77 and Data S4). Without location as a co-varying partner, the strongest edges shifted to epigenomic features. In LF-MF1 group1, Re-Hd5 (downstream H3K27ac) was the strongest co-varying pair and Hd5 was also among the most structurally prominent features, appearing in three of the six strongest edges in the network. In LF-MF2 group1, Re remained the most connected node, and five of the six strongest co-varying pairs involved Re. TEdi and ACRdd were its two strongest partners, followed by all three H3K27me3 marks (Hg6, Hu6, and Hd6). In the MF1-MF2 group1 network, Re co-varied positively with Ka and downstream CG methylation (M8) while H3K27ac marks showed negative co-variation with Re. This network also exhibited a higher proportion of negative edges (42%) than both LF-MF1 and LF-MF2 group1 networks (∼12-14%), indicating a greater prevalence of opposing relationships among predictive features.

SHAP interaction networks also revealed that recombination rate was the most connected node in all six interaction networks, with degrees ranging from 18 to 27, consistent with the co-variation network results (Figs. 78 and 79 and Data S5). In contrast to the predominantly positive co-variation structure observed in the LF all-genes and group1 networks, negative interactions, indicative of partial redundancy in feature contributions, accounted for the majority of above-threshold edges across all six interaction networks (54-67%). In LF-MF1, the strongest interaction in both all-genes and group1 datasets was the positive Hd5-Re pair, indicating a synergistic relationship between downstream H3K27ac and recombination rate in classifying duplicated genes. In LF-MF2, Re was again the most connected node in both datasets but did not form the strongest interaction. Instead, the highest-scoring edges involved negative interactions between upstream H3K4me3 (Hu3) and gene expression in the all-genes network, and between two H3K27me3 marks (Hd6-Hg6) in the group1 network, suggesting partial redundancy among these features in distinguishing LF and MF2 genes. In the MF1-MF2 models, Re-Loc was the strongest interaction in the all-genes network, while Re-Ks was the strongest in group1. The MF1-MF2 group1 interaction network was the sparsest of the six, retaining only 43 above-threshold edges across 29 nodes. This sparsity is consistent with the overwhelming dominance of Re in this model, leaving relatively few additional features with sufficient SHAP contributions to generate strong pairwise interactions.

Per-label interaction networks for all six *B. rapa* datasets revealed that Re remained the most connected node across all twelve networks (Figs. S80 and S81 and Data S7). A consistent asymmetry between labels was observed in the LF-MF2 and MF1-MF2 all-genes models. In both cases, Re-Loc was the strongest interaction edge in both the dominant and recessive label networks, but the interaction signs were reversed. The interaction was negative in the dominant label network, indicating partial redundancy between recombination rate and chromosomal location when classifying dominant subgenome genes, but positive in the recessive label network, indicating synergy between these two features when classifying recessive subgenome genes.

### Recombination rate is largely sufficient for subgenome classification in *B. rapa* but not maize

To quantify the contribution of recombination rate to subgenome classification, we performed feature ablation analyses under three conditions: all features included as in the models described previously, recombination rate excluded, and recombination rate as the sole predictor (Figs. 8 and S82-S93 and Data S8).

**Fig. 8.**
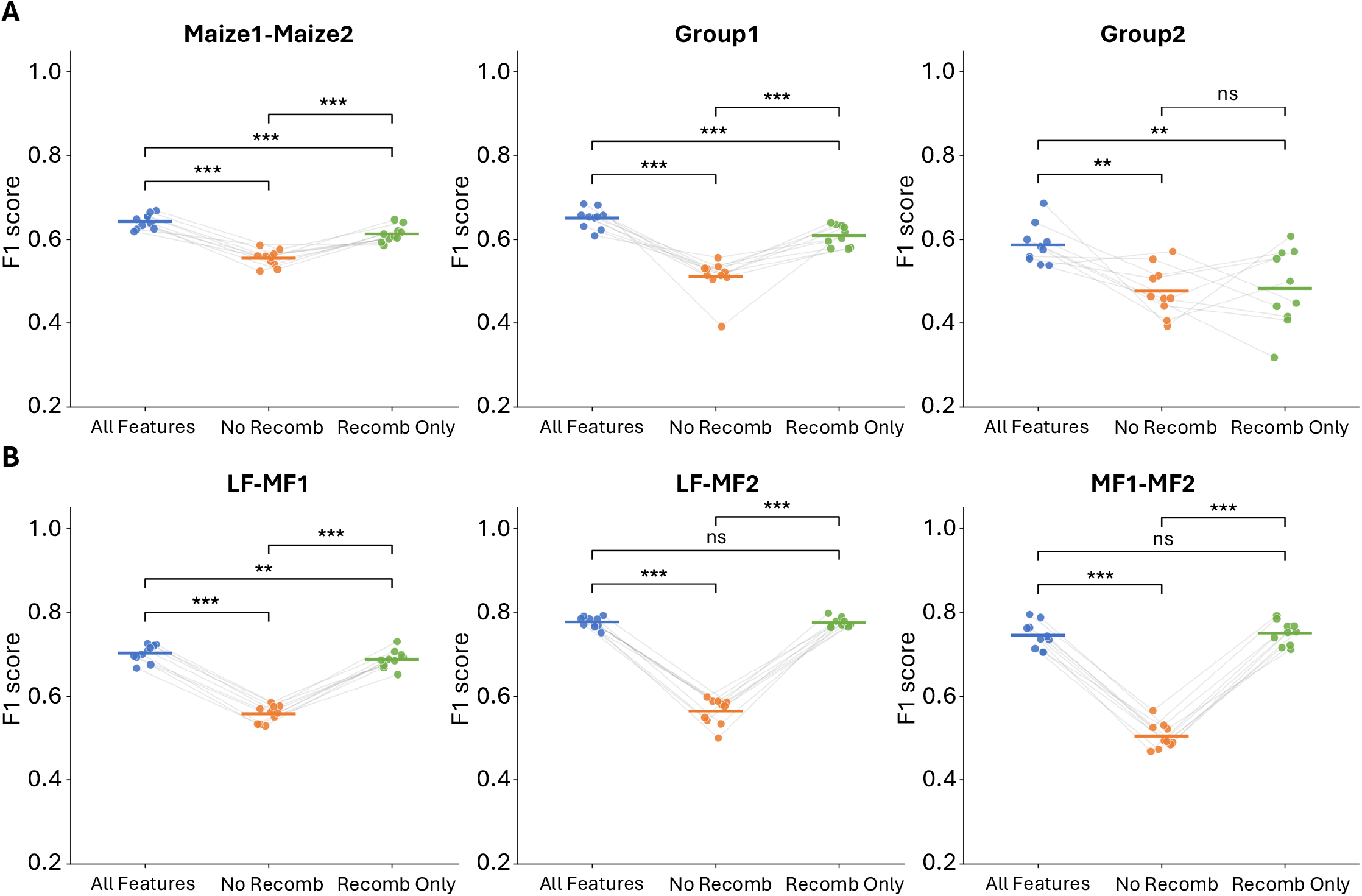
Recombination rate is largely sufficient for subgenome classification in *B. rapa* but not maize. F1 scores from each of the 10 stratified CV folds are shown under three feature conditions: all features (blue), recombination rate excluded (orange), and recombination rate as the sole feature (green) for maize (**A**) and *B. rapa* (**B**). Each point represents one fold, and gray lines connect matching folds across conditions to make the paired structure explicit. The horizontal bar within each condition indicates the mean F1 across folds. The same Optuna-tuned XGBoost hyperparameters and identical fold splits were used across all three conditions; models were not retrained under reduced feature conditions. Pairwise differences in F1 between conditions were assessed using two-sided Student’s paired *t*-tests on the 10 per-fold scores. Benjamini-Hochberg FDR correction was applied within each group independently (three tests: All Features vs. No Recombination, All Features vs. Recombination Only, No Recombination vs. Recombination Only; * *P* < 0.05, ** *P* < 0.01, *** *P* < 0.001, ns, not significant).

In maize all-genes, group1, and group2 datasets, removing recombination rate significantly reduced classification performance. In the all-genes model, mean F1 declined from 0.643 to 0.555 upon recombination removal (Student’s paired *t*-test, BH-corrected *P* < 0.001, Figs. 8A and S82). Using recombination rate as the sole predictor also resulted in significantly lower performance than the full model (mean F1 = 0.613, *P* < 0.001), indicating that the remaining epigenomic and genomic features contribute predictive information beyond recombination rate alone (Data S8). A consistent pattern was observed in group1, where F1 declined from 0.650 to 0.511 without recombination rate (*P* < 0.001), and recombination rate alone (F1 = 0.609) remained significantly less predictive than the all-features model (*P* < 0.001, Fig. S83). Notably, however, AUC showed no significant difference between the full model and recombination-only conditions for group1 (Fig. S92), in contrast to the F1 results. Group2 exhibited a distinct pattern. Although removing recombination rate significantly reduced F1 relative to the all-features model (F1: 0.587→0.477, *P* < 0.01, Fig. S84), recombination rate alone performed no better than the no-recombination rate condition by F1 (F1 = 0.483 vs. 0.477, ns). AUC comparisons yielded a different result: the all-features and recombination-only models were not significantly different, while recombination-only had significantly higher AUC than the no-recombination condition (*P* < 0.001, Fig. S92), indicating metric-dependent effects in pericentromeric genes.

In contrast to maize, recombination rate alone largely captured the full predictive signal across *B. rapa* subgenome comparisons. Removing recombination rate significantly reduced F1 across all three all-genes models, with mean declines of 0.145, 0.212, and 0.240 for LF-MF1, LF-MF2, and MF1-MF2 respectively (all *P* < 0.001, Fig. 8B and S85-S87, and Data S8). However, using recombination rate as the sole predictor recovered near-full performance in LF-MF2 and MF1-MF2. In both cases, F1 did not differ significantly from the all-features model (mean differences of 0.001 and −0.006, both ns). AUC comparisons were concordant for MF1-MF2, showing no significant difference between the all-features model and only-recombination condition. In LF-MF2, however, AUC revealed a small but significant reduction of approximately 0.02 when only recombination rate was used (*P* < 0.001, Fig. S93), indicating a modest discrepancy between F1 and AUC. LF-MF1 was the only all-genes model in which recombination rate alone remained significantly less predictive than the all-features model by F1 (mean F1 = 0.688 vs. 0.703, *P* < 0.01, Figs. 8B and S85). Group1 results largely mirrored the all-genes analyses. Recombination alone matched all-features performance in both LF-MF2 and MF1-MF2 group1 datasets (both ns by F1, Figs. S89-S91). Moreover, the LF-MF1 exception observed in the all-genes analysis did not persist in group1, where the difference between the all-features model and only-recombination model conditions was not significant (*P* = 0.081, ns, Figs. S88 and S91).

Taken together, these results revealed a marked contrast between maize and *B. rapa*. In maize, the all-features set consistently outperformed recombination alone across all-genes, group1, and group2 datasets by F1, demonstrating that additional genomic and epigenomic features contribute substantial predictive information. In *B. rapa*, by contrast, recombination rate alone was sufficient to recover nearly all predictive performance in most comparisons, particularly for LF-MF2 and MF1-MF2. Although several AUC comparisons revealed small but significant differences between the all-features model and recombination-alone conditions, the overall pattern indicates that recombination rate captures a substantially larger fraction of the subgenome-classification signal in *B. rapa* than in maize.

### TE density and TE distance to genes are among top features differentiating subgenomes in maize and *B. rapa*

Differences in TE abundance, distribution, and epigenetic silencing between diploid progenitors have long been hypothesized as contributors to the differential expression of duplicated genes, as silenced TEs often exert deleterious effects on the expression of nearby genes (*18, 19, 33, 34, 50*). We examined TE type, TE density, and TE distance to genes across a total of nine features (Fig. 2). TE types were generally weak predictors across both species. Interspersed nucleotide elements (LINEs) and short interspersed nuclear elements (SINEs) were consistently ranked at the bottom across all models. Other TE types, including helitrons (TE1), terminal inverted repeat elements (TE2), and LTR retrotransposons (TE3), also tended to show low rankings across feature importance profiles. Because miniature inverted-repeat transposable elements (MITEs) have been found to associate with recombination hotspots in maize and other species (*51, 52*), we evaluated MITEs as an independent feature in our model. However, MITEs were not a significant predictor of subgenome evolution (Fig. S94).

In contrast, TE density features ranked substantially higher. In maize, average downstream TE density (TEd1) was the most prominent TE-related feature, ranking 6th in the all-genes model and 11th in group1 (Fig. 3A and B). Beyond its individual SHAP contribution, TEd1 also ranked fifth by node degree in the all-genes interaction network (Fig. 5A), indicating that downstream TE density joins with other features to influence model predictions beyond its direct effect alone.

In *B. rapa*, TE distance to genes (TEdi) was the most consistently prominent TE-related feature across models, ranking ninth in both the LF-MF1 and LF-MF2 all-genes models and 13th in MF1-MF2 (Fig. 6). Its importance increased further in the group1 subsets, rising to seventh in LF-MF1 group1 and sixth in LF-MF2 group1 (Figs. S62-S63). This prominence is consistent with its position as the fifth most connected node in the LF-MF2 co-variation network (Fig. 7). Together, these results indicate that TE density represents the strongest TE-associated signal in maize subgenome differentiation. In contrast, TE distance to genes is the most consistently important TE-related feature across *B. rapa* models.

## Discussion

Our XAI approach identified recombination as the most influential feature for subgenome classification in both maize and *B. rapa*, together with closely associated features including gene chromosome location, and open chromatin features such as the distance of ACRs to genes and active histone marks (Figs. 3 and 6). These results are consistent with previous work showing that chromatin environment is critical for the maintenance of gene dominance in maize (*20*), cotton (*40*), and strawberry (*21*). In maize, we previously showed that biased fractionation is largely attenuated when homoeologous genes are positioned in pericentromeric regions (*20*). Consistently, our XAI classification models achieved the highest performance when maize1 and maize2 genes were located in contrasting chromosome contexts (groups 3 and 4, Fig. 1B, E, and G), reflecting strong differences in recombination rate, TE associated features, and chromatin state between chromosomal arms and pericentromeric regions. Together, these results indicate that subgenome dominance classification is strongly shaped by chromosomal context and recombination landscape, while the contribution of additional features depends on species and genomic context. More broadly, these patterns align with the emerging view of polyploidy as a continuum, in which genomes range from more autopolyploid-like to more allopolyploid-like states depending on the degree of subgenome divergence and interaction. In this framework, chromosomal context and recombination landscape can be viewed as key axes along which subgenomes differentiate (*53, 54*).

Recombination rate emerged as the most influential predictor of genome dominance in both maize and *B. rapa*, even though recombination rates were not significantly different among subgenomes in several comparisons (Figs. 3 and 6, and Figs. S1, S2, S19-S23, and S36, S37, S44-S49, and Data S1). In maize groups 3 and 4, where duplicated genes are in contrasting chromatin locations, recombination remained a dominant feature, but its SHAP pattern was context dependent rather than uniformly monotonic, reflecting the strong coupling between recombination, chromosomal location, and other chromatin-associated features (Figs. S26 and S27).

From a machine learning perspective, recombination can be interpreted as a highly informative meta-feature that captures part of the joint signal associated with genomic architecture, including chromatin accessibility, epigenetic state, and TE density (*55, 56*). Its predictive value likely reflects both its direct association with subgenome classification and indirect correlation with multiple covarying genomic features, although the relative contribution of these effects differs between maize and *B. rapa*. This interpretation aligns with recent multi-omics and machine learning studies in *B. napus*, where repeat-rich, highly methylated regions were predictive of reduced recombination frequencies, whereas gene-dense, euchromatic regions were enriched for recombination events (*57*). Correspondingly, feature interaction networks place recombination near the center in both species, but the surrounding network structure varies by species and chromosomal context (Figs. 5, S78, and S79). In maize, recombination appears to act as a more context-dependent feature whose predictive value is distributed across a broader network of genomic and epigenomic signals. This is reflected in the stronger dependence of maize models on multiple correlated features, the richer interaction structure, and the fact that recombination alone does not recover full classification performance (Figs. 8A, S92, and Data S8). In contrast, *B. rapa* models are more heavily dominated by recombination, with simpler network structure and ablation results showing that recombination alone recovers much of the predictive signal (Figs. 8B, S91 and S93). These results suggest that subgenome classification in maize is more complex and multi-layered, whereas in *B. rapa* it is more strongly concentrated around recombination.

Our results further suggest that recombination does not act in isolation but integrates multiple aspects of genome organization, including TE density, chromatin accessibility, and histone modifications, particularly in maize, where these features co-vary with chromosome location and fractionation status. SHAP analysis indicates that the models use recombination in combination with other covarying factors rather than as a fully independent variable, but the strength of this dependence differs by species: maize shows a more diffuse, context-dependent signal, whereas *B. rapa* is more strongly dominated by recombination itself. These patterns are consistent with TE load theory and support the idea that TE accumulation and its epigenetic consequences shape recombination landscapes and genome dominance (*18, 19, 33, 34, 58*). However, while SHAP values reflect the contribution of features to model predictions, they do not establish causality or directionality and therefore require further validation. Our binary classification framework and selected features capture key elements of subgenome dominance, but additional uncollected biological factors may also influence crossover dynamics.

From a biological and evolutionary perspective, recombination hotspots are generally suppressed in condensed pericentromeric regions and enriched in gene-rich euchromatic regions, particularly near TSSs and TTSs, which are characterized by reduced nucleosome occupancy, enhanced chromatin accessibility, DNA hypomethylation, low H3K9me2, and increased levels of H3K4me4 and H2A.Z (*59–62*). Regions with elevated recombination typically experience more efficient purifying selection and reduced effects of linked selection (*63, 64*), facilitating faster removal of deleterious mutations and thereby promoting gene retention and maintaining functional constraints (*65, 66*). In contrast, in TE-rich, low-recombination regions, natural selection is less efficient in purging deleterious mutations. Consequently, likely due to dosage effects (*67*), when both gene copies are retained, genes in low-recombination regions show reduced expression because of accumulated deleterious mutations and eventually lose function over sufficiently long evolutionary timescales (*20, 65*). Our findings suggest that recombination integrates multiple aspects of genome organization and evolutionary history, particularly in maize, where its predictive signal emerges from a broader network of correlated epigenomic and structural features. However, this layered interpretation is less pronounced in *B. rapa*, where recombination more directly captures the dominant signal underlying subgenome classification.

Our findings are consistent with cytological studies showing that chromosome pairing behavior, homoeologous recombination, and chromosome restructuring play important roles in polyploid genome evolution (*44–46, 68, 69*). Although our recombination data are derived from modern mapping populations (*20, 70–72*), they likely retain signatures of historical chromosomal organization and selection. Following polyploidization, interactions between homoeologous chromosomes can generate homoeologous exchanges and structural rearrangements, leading to unequal genome restructuring and differential retention of genetic material among subgenomes (*44, 73*). Homoeologous recombination often occurs during the first generation following polyploidization and persists over subsequent generations but gradually declines as allopolyploids become established and meiosis stabilizes, likely reflecting selection for mechanisms that stabilize chromosome pairing and promote diploid-like meiotic behavior (*69*). For example, adaptive changes affecting crossover control have been documented in autotetraploid *Arabidopsis arenosa* (*45*), while reduced homoeologous crossovers associated with MSH4 copy number evolution have been reported in allopolyploid *B. napus* (*46*). The strong predictive importance of recombination identified in our models is therefore consistent with the broader view that chromosome pairing and recombination dynamics contribute to long-term subgenome differentiation. Although our analyses do not directly measure meiotic processes or historical homoeologous exchanges, the genomic and epigenomic signatures captured by our models may reflect the cumulative effects of these chromosome-level evolutionary processes.

By identifying genomic and epigenomic features that most strongly modulate genomic dominance, our XAI approach provides evolutionary insights into how genome architecture shapes subgenome differentiation while also informing strategies for crop improvement, where hybridization is often the first step. In species in which chromatin accessibility, histone modifications, or DNA methylation make major contributions, subgenome dominance may not be entirely fixed and could, in principle, be altered through changes in these regulatory states. Identifying the features that most strongly influence subgenome dominance, and understanding how they interact with recombination and chromosomal context, provides a framework for developing strategies to modulate dominance patterns rather than relying solely on selection of existing genetic variation. More broadly, this work demonstrates how integrative, explainable machine learning can reveal the genomic and epigenomic factors underlying polyploid genome evolution while providing a foundation for the rational improvement of polyploid crops.

## Materials and Methods

### Gene classification by subgenomes

The list of duplicated gene pairs in maize, based on the version 4 reference genome, was obtained from https://doi.org/10.6084/m9.figshare.7926674.v1 (*16, 74*). Of the 4,578 duplicated gene pairs in maize with syntelogs in sorghum, 3,536 were retained after excluding pairs with missing feature data. Following a previous study (*20*), retained genes were classified into four groups according to their genomic location: group1, both maize1 and maize2 genes located in chromosomal arms; group2, both genes located in pericentromeric regions; group3, maize1 genes located in chromosomal arms and maize2 genes in pericentromeric regions; and group4, maize1 genes located in pericentromeric regions and maize2 genes in chromosomal arms (Fig. 1B and C). Duplicated genes in *B. rapa* with syntelogs in Arabidopsis were obtained from previous research (*18, 26*). Three pairwise subgenome comparisons were analyzed: LF-MF1, LF-MF2, and MF1-MF2, comprising 4,294, 3,666, and 2,504 gene pairs in the source dataset, respectively. After excluding pairs with missing feature data, 2,362, 1,993, and 1,291 pairs were retained for analysis, with group1 subsets of 2,036, 1,588, and 1,017 pairs, respectively (Fig. 1D).

### Feature data collection and statistical characterization in maize and *B. rapa*

Genomic and epigenomic features for maize and *B. rapa* were compiled from publicly available resources (see detailed feature collection in the *Supplementary Materials and Methods*). For each species, these included recombination rate and gene chromosomal location (one feature), gene expression metrics (two features), evolutionary distances (three features), GC content (two features), transposable elements (nine features), DNA methylation (15 features), and ACRs (three features). Histone modification data comprised 24 features in maize and nine in *B. rapa* (Fig. 2). In total, 60 features were analyzed in maize and 45 in *B. rapa*. Detailed procedures for feature processing and analysis are provided in the *Supplementary Materials and Methods*.

Prior to model development, feature distributions were compared between dominant and non-dominant subgenome genes using two-sided Mann-Whitney U tests with Benjamini-Hochberg FDR correction. Effect sizes were quantified using rank-biserial correlation (Data S1, Figs. S1, S2, S36, and S37). In both the all-genes maize1-maize2 dataset and the all-genes *B. rapa* datasets, chromosomal location was excluded as a binary categorical variable. Pairwise relationships among features were assessed using Spearman rank correlations with Bonferroni correction across all feature pairs, with missing values excluded on a per-pair basis. Correlations between recombination rate and all other features were examined specifically given its potential confounding relationship with subgenome dominance patterns (Data S2, Figs. S3-S12 and S38-S49).

### Predictive model development

Classifier models included five maize models, including the full maize1 and maize2 whole genome duplicated genes and four groups. For *B. rapa*, six models were implemented, including whole genome triplicated genes for the three subgenomes in three pairwise comparisons (LF vs. MF1, LF vs. MF2, and MF1 vs. MF2) and group1 subsets of these three pairwise comparisons.

### Benchmarking

For both maize and *B. rapa*, four model architectures were benchmarked to identify the best-performing classifier for our datasets: logistic regression (LR), support vector machine (SVM), random forest (RF), and XGBoost (XGB). Because LR, SVM, and RF cannot handle missing values natively, samples (here indicate individual genes) containing any missing values were excluded prior to benchmarking. XGBoost was evaluated on the same reduced dataset to ensure a like-for-like comparison across all four architectures. Feature scaling was performed using StandardScaler, fitted only on the training fold to prevent data leakage. Scaling was applied to LR and SVM because these models are sensitive to feature magnitude; RF and XGBoost are split-based and therefore scale-invariant, so no scaling was applied. All classifiers were evaluated using stratified 10-fold cross-validation with default hyperparameter settings, except that probability estimation was enabled for the SVM and log loss was used as the evaluation metric for XGBoost. Models were implemented using scikit-learn (v1.3.0) (*75*) and xgboost (v1.7.6) (*76*). Per-fold F1 scores from each baseline architecture (LR, SVM, and RF) were compared against XGBoost using two-sided Wilcoxon signed-rank tests. Benjamini-Hochberg FDR correction was applied across the three comparisons within each dataset independently. Results are summarized in Tables S1 and S4.

### Hyperparameter optimization

XGBoost models were optimized using Optuna (v3.2.0) (*77*) with a Tree-structured Parzen Estimator (TPE) sampler, aiming to maximize mean binary F1 score computed on the held-out test folds within stratified 10-fold cross-validation. Within each fold, the training data were further partitioned into a 90% fitting set and a 10% inner validation set used exclusively for early stopping. Training halted after 30 consecutive rounds without improvement in validation log loss, ensuring that the held-out test fold did not influence training or stopping decisions. Features with biologically meaningful missing values, specifically tissue specificity index (τ, undefined for genes with zero expression across all sampled tissues) and ACR distance features (undefined for genes at chromosome boundaries), were retained in the feature matrix, as XGBoost natively routes missing values at each split without imputation. Hyperparameter search spaces and final tuned values for maize and *B. rapa* are provided in Tables S2, S3, and S5.

### Final model training and evaluation

Final XGBoost models were trained using stratified 10-fold cross-validation with the same inner validation split and early stopping procedure described above. Model performance was evaluated using mean F1 score and ROC/AUC across folds. Potential overfitting was assessed by examining training and validation log loss curves across boosting rounds for each fold. Performance metrics are shown in Figs. 1E, 1G, S17 and S18 for maize and in Figs. 1F, 1H, S57, S58 for *B. rapa*.

### SHAP feature importance and network analysis

#### SHAP feature importance

Feature importance was assessed using SHAP values to interpret the contribution of individual features to model classifications (shap v0.45.0) (*47*). Within each cross-validation fold, a SHAP TreeExplainer was fitted to the inner training set (the 90% fitting partition) and SHAP values were computed for the held-out test fold, ensuring that SHAP attribution reflected model generalization rather than training fit. Because each individual gene appeared in exactly one test fold during cross-validation, SHAP values were concatenated across folds to produce a complete, non-overlapping dataset covering all samples (genes). To examine asymmetry in feature importance between subgenomes, SHAP values were also computed separately for dominant and recessive subgenome genes by partitioning the aggregated array by true label after concatenation. For Figs. S14-S18 and S59-64, positive and negative SHAP values were summed separately for each feature and visualized using back-to-back horizontal bar plots. For Figs. 3 and 6, S24-32 and S65-76, SHAP beeswarm plots were generated using shap.summary_plot. Per-label shap beeswarm plots are shown in Figs. S28-S32 and S71-S76.

#### Co-variation networks

For each model, pairwise SHAP co-variation scores were computed from the aggregated SHAP value matrix. The co-variation score for a feature pair was defined as the mean of the products of their SHAP values across all genes, where a positive score indicates that the two features tend to contribute in the same direction across predictions and a negative score indicates opposing contributions. Scores were computed for all upper-triangle feature pairs. Statistical significance was assessed using an empirical null distribution generated from 1,000 permutations in which the rows of the SHAP array were shuffled across rows, thereby disrupting between-sample co-variation while preserving within-sample feature structure. All upper-triangle scores across all permutations were pooled into a single null distribution. Two-sided empirical p-values were computed as the proportion of null scores whose absolute value equaled or exceeded the observed score, and feature pairs with p ≤ 0.05 were retained as network edges. Networks were constructed and visualized using NetworkX (v3.2.1) (*78*) with a Kamada-Kawai layout. Edge width was scaled to reflect co-variation magnitude and edge color indicated direction. Node size was proportional to degree, defined as the number of retained edges for that node, and node color denoted feature category. Co-variation networks are shown in Figs. 4 and S33 for maize and Figs. 7 and S77 for *B. rapa* and network edge weights and node degrees are available in Data S4.

#### Interaction networks

Shapley interaction values computed during cross-validation were used to construct feature interaction networks for each model. For each upper-triangle feature pair, the mean Shapley interaction value was computed across all samples (genes). To distinguish meaningful interactions from the large number of near-zero pairs, a robust effect size threshold was applied. For each model, the threshold was defined as the median plus 12 times the median absolute deviation (MAD) of the absolute mean interaction values, computed from non-zero pairs only. A uniform multiplier (k = 12) was applied across all models, selected based on exploratory analyses of the interaction value distributions, providing a consistent standard of evidence across groups with different absolute interaction scales. Feature pairs whose absolute mean interaction value exceeded the group-specific threshold were retained as network edges. Positive edges indicate synergistic interactions, in which the combined contribution of two features to a prediction exceeds the sum of their individual effects, whereas negative edges indicate antagonistic interactions, in which the combined contribution is less than the sum of their individual effects. Networks were visualized using the same approach as the co-variation networks. Interaction networks are shown in Figs. 5, S34, and S35 for maize and Figs. S78-81 for *B. rapa.* Network edge weights and node degrees are available in Data S5. Edge weights and node degree data from per-label interaction networks are available in Data S6 for maize and in Data S7 for *B. rapa*.

### Recombination rate ablation analysis

To quantify the contribution of recombination rate to model performance, three feature configurations were evaluated for each model: (i) the full feature set including recombination rate, (ii) all features excluding recombination rate, and (iii) recombination rate as the sole feature. Hyperparameters were tuned on the full feature set and held fixed across all three configurations to isolate the marginal contribution of recombination rate without confounding effects from re-optimization. Model performance under each configuration was assessed using the same stratified 10-fold cross-validation procedure described above. Pairwise differences in per-fold F1 scores between conditions were evaluated using paired t-tests, and pairwise differences in AUC were evaluated using the DeLong test, which accounts for correlations between AUC estimates derived from the same set of genes. Benjamini-Hochberg FDR correction was applied independently within each model group and separately for each performance metric, resulting in three statistical comparisons per group per metric. Results of the ablation analyses are shown in Figs. 8, S82-S84 (maize), S85-S90 (*B. rapa*), and S91-S93 (metric comparisons). Associated data are provided in Data S8.

## Acknowledgements

We thank the University of Florida HiPerGator Supercomputer for providing computational resources for this analysis.

## Author contributions

Conceptualization: L.A.S., R.D., and M.Z. Methodology: L.A.S., J.C.F.S., R.D., and M.Z. Investigation: L.A.S., X.C., B.L., and M.Z. Visualization: L.A.S. Supervision: R.D. and M.Z. Writing—original draft: L.A.S. and J.C.F.S. Writing—review & editing: R.D. and M.Z.

## Data and materials availability

All data and code needed to evaluate and reproduce the results in the paper are present in the paper and/or the Supplementary Materials. All raw data used in this study are publicly available.

