## Supplementary Information for "Explainable AI identifies recombination and chromatin environment as key predictors of subgenome evolution in maize and *Brassica*"

Layla A. Schuster *et al.*

**This PDF file includes:**

Supplementary Text
Figs. S1 to S94
Tables S1 to S5
Legend for Data S1-S8
References (1 to 28)

**Other Supplementary Materials for this manuscript include the following:**

Data S1-S8

**Supporting Text**

### Supplementary Materials and Methods

#### Recombination Rates

Recombination rate (Re) contributes one feature to each model, representing the estimated local crossover frequency at each gene's chromosomal position in units of centimorgans per megabase (cM/Mb). In both species, per-gene recombination rates are smoothed estimates interpolated from genetic maps rather than direct per-gene measurements, but the interpolation method and spatial resolution of the estimates differ between the two species as described below.

Maize: recombination rates (cM/Mb) were estimated using 6,257 of the 10,188 genetic markers from the integrated map previously described (*1*). These markers were analyzed using MareyMap (*2*), and gene-level recombination rates were assigned based on the midpoint position of each gene along the genetic map.

*B. rapa*: recombination rates were estimated using 430 genetic markers following the same approach as in maize (*3, 4*). The recombination rate file covers only genes assigned to the main chromosomes. Gene pairs located on scaffold chromosomes lack genetic map positions and therefore have no recombination rate estimate available and were excluded from the final feature set.

#### Gene Location

Gene chromosomal location (Loc) contributes one feature to each model, indicating whether a gene resides in a chromosome arm or a pericentromeric region. For maize, location was used to stratify gene pairs into four structural groups: group1 (maize 1 arm-maize 2 arm), group2 (maize1 pericentromeric-maize2 pericentromeric), group3 (maize1 arm-maize2 pericentromeric), and group4 (maize1 pericentromeric-maize2 arm). For *B. rapa*, the same grouping scheme was applied, but only group1 was used because the number of gene pairs in the other categories was insufficient for robust analysis (*5, 6*).

Maize: The boundaries between chromosomal arms and pericentromeric regions for all ten maize chromosomes were defined following our previous research (*5, 6*). Briefly, gene density (genes/Mb) and repeat density (Mb/Mb) were calculated in 1-Mb windows with 500-kb shifts along each chromosome, and recombination rates were estimated for each window using the integrated genetic map described above. Based on the characteristic pattern of suppressed recombination, reduced gene density, and elevated TE density in pericentromeric regions, each chromosome was manually partitioned into two arms and one pericentromeric region. Each gene was assigned to chromosomal arms or pericentromeric regions accordingly.

*B. rapa*: Each gene was classified as residing in either a chromosome arm region or a pericentromeric region following the same boundary criteria described above and applied to the *B. rapa* v1.5 genome annotation. Because *B. rapa* chromosomes are substantially shorter than those of maize, gene density, TE density, and recombination rates were calculated in 500-kb windows with 100-kb shifts. The final location annotation included 39,608 genes across the ten main chromosomes. Genes located on scaffolds were excluded from the analysis.

**GC Content**

GC content contributes two features per gene in both species: GC content of the coding sequence (GCd) and GC content of a 170-bp core promoter window (GCp; −165 to +5 bp relative to the transcription start site, TSS), following previous research (*7*). The core promoter window definition is identical across species. TSS coordinate sources differ as described below.

Maize: Coding sequence GC content was calculated from the primary transcript CDS sequences in the Phytozome Maize B73 v4 annotation. Promoter GC content was calculated using a 170-bp core promoter window spanning −165 to +5 bp relative to the TSS. Experimental TSS coordinates from previous research were used whenever available (*7*); otherwise, the TSS was defined as the mRNA start coordinate from the Maize GDB B73 v4 GFF3 annotation. Of the 9,156 target genes, 7,889 were assigned experimental TSS coordinates (*7*) and the remaining 1,267 used annotation-derived TSS positions. Because all target genes were located on fully assembled chromosomes, no promoter windows crossed chromosome boundaries, and no missing values were introduced for this feature.

*B. rapa*: Coding sequence GC content was calculated by extracting annotated CDS exon intervals from the *B. rapa* v1.5 GFF annotation, retrieving the corresponding sequences from the v1.5 genome assembly, concatenating exons in genomic order, and calculating the GC fraction of the resulting coding sequence. Because the *B. rapa* v1.5 annotation contains a gene model per locus, no isoform selection was required. Promoter GC content was calculated using the same 170-bp core promoter window as in maize. The TSS was defined as the mRNA feature start coordinate for plus-strand genes and the mRNA end coordinate for minus-strand genes, with minus-strand sequences reverse-complemented prior to GC calculation. No experimental TSS dataset was available for *B. rapa*. Therefore, all TSS coordinates were obtained from the genome annotation. Genes whose promoter window extended beyond a chromosome or scaffold boundary received NaN for this feature. All such cases were confined to scaffold-resident genes.

**Gene Expression**

Two expression features were derived for each gene in both species: average expression level (Exp) and the tissue specificity index tau (τ). Average expression was calculated as the mean of $\log_2(\mathrm{FPKM} + 1)$ values across all tissues. Tissue specificity was quantified using τ, which measures the extent to which expression is restricted to a subset of tissues. Genes with FPKM = 0 across all tissues have mathematically undefined τ. Because these cases are biologically meaningful, τ was retained as NaN rather than treated as missing data. The number of tissue conditions used for τ calculation differs between species. Therefore, τ values are not numerically comparable across species.

Maize: Expression data were obtained from RNA-seq FPKM estimates across 24 developmental stages and tissue types, including germinating seeds, primary root, shoot apical meristem, multiple leaf stages, whole seed at multiple timepoints, endosperm, embryo, ear, and tassel (*6, 8*). Expression magnitude was quantified as the arithmetic mean of $\log_2(\mathrm{FPKM} + 1)$ values across all 24 tissues. Tissue specificity was measured using the τ index, calculated from the $\log_2$-transformed values according to $\tau = \Sigma(1 - x\_i/x\_max) / (N - 1)$, with N = 24. Genes with FPKM = 0 in all 24 tissues received NaN for τ. This applied to 66 maize1 genes and 82 maize2 genes, and their NaN values were retained throughout the analysis. No NaN values were introduced for average expression level.

*B. rapa*: Gene expression data were obtained from GEO accession GSE43245 (*9*), which provides Cufflinks FPKM estimates for eight tissue samples from *B. rapa* Chiifu mapped to the v1.5 genome reference. The sampled tissues included callus, flower, two leaf samples (Leaf1 and Leaf2), two root samples (Root1 and Root2), silique, and stem. Expression magnitude was quantified as the arithmetic mean of $\log_2$(FPKM + 1) values across all eight tissue columns. Expression breadth was quantified using the τ tissue specificity index, calculated from six biologically distinct tissue conditions. Prior to the τ calculation, the two leaf tissues were averaged into a single leaf value, and the two root samples were averaged to a single root value. The τ index was calculated using the same formula as for maize, but with N = 6. Genes with FPKM = 0 across all eight tissues (n = 1,491 genome-wide) have mathematically undefined τ values and were assigned NaN for this feature. Average expression was calculated from all eight tissues averaging to maximize the information used in estimating expression magnitude. A total of 340 master doublet genes were absent from the GSE43245 expression dataset. These genes were assigned NaN for both expression features and their associated gene pairs were excluded during the final data-filtering step.

**Evolutionary Distance**

Evolutionary distance on genes was quantified using three metrics: the nonsynonymous substitution rate (Ka), the synonymous substitution rate (Ks), and their ratio (Ka/Ks), referred to as omega (ω). For maize, *Sorghum bicolor* (reference genome version 3) was used as the outgroup, and for *B. rapa*, Arabidopsis was used as the outgroup. A list of maize (v4) genes and their sorghum syntelogs was obtained from a previous study (*10*) and Arabidopsis syntelogs with *B. rapa* were obtained from previous studies (*11, 12*).

For each gene pair, coding sequences were aligned using MUSCLE (*13*) or ClustalW (*14*) or PRANK (*15*). Sequence alignments were evaluated sequentially in the order MUSCLE, ClustalW, and PRANK because gaps introduced during alignment occasionally prevented downstream evolutionary-rate estimation. When an alignment generated by one method could not be used, the next method in the sequence was applied. Ka, Ks, and ω values were then estimated from the resulting codon alignments using the yn00 program implemented in PAML version 4.7 (*16*). Evolutionary distance metrics for *B. rapa* genes were calculated using the same procedure, except that Arabidopsis syntelogs were used as the corresponding outgroup sequences (*13-16*).

**Transposable Elements**

Transposable element features comprise nine features per gene: average upstream TE density (TEu1), maximum upstream TE density (TEu2), average downstream TE density (TEd1), maximum downstream TE density (TEd2), four one-hot encoded TE type indicators (TE1-TE4), and the genomic distance to the nearest TE (TEdi). Regional TE density was calculated as the proportion of base-pair coverage in 100 bp windows with 10 bp increments across the 2 kb upstream and downstream flanking regions of each gene, following our previous method (*6, 8*). Mean and maximum density were subsequently calculated for the upstream and downstream regions separately. TE elements were classified into four functional groups: TE1, Helitron DNA transposons; TE2, terminal inverted repeat (TIR) DNA transposons; TE3, long terminal repeat (LTR) retrotransposons; TE4, long interspersed nuclear elements (LINEs) and short interspersed nuclear elements (SINEs). Both species were processed using the same density window parameters.

Maize: TE annotations were derived from a curated library of 1,526 consensus and exemplar TEs mapped to the maize B73 v4 reference genome using RepeatMasker with a sequence divergence cutoff of <20%, capturing both intact elements and structurally fragmented remnants (*6*). TE distance (TEdi) was defined as the absolute genomic distance between the proximal edge of the nearest TE and the nearest transcriptional boundary of the gene (TSS for upstream elements and TTS for downstream elements). Genes directly overlapping a TE were assigned a distance of zero. TE type was assigned based on the nearest TE. When nested insertions resulted in multiple TE classes occurring at the same minimum-distance boundary, a single TE type was retained using a strict categorical hierarchy priority order: TE1 > TE2 > TE3 > TE4.

*B. rapa*: TE distance and TE type were derived from the *B. rapa* v1.5 TE annotation. TE distance was calculated using the complete TE annotation, including unclassified elements. When multiple equidistant TEs were returned, the minimum distance was retained. TE type was determined from a separate analysis restricted to TEs with known family designations, including the four groups of TEs mentioned above. Consequently, TE distance and TE type do not necessarily correspond to the same TE for a given gene: TE distance captures the absolute nearest annotated TE regardless of classification, while TE type reflects the nearest element with a known family designation. TE density was calculated using the same procedure applied to maize. Specifically, TE density was computed from 100 bp windows with 10 bp increments across the 2 kb upstream and downstream regions of genes. Mean and maximum density were then calculated separately for the upstream and downstream regions.

**Accessible Chromatin Regions (ACRs)**

ACR features contribute three variables per gene: the fold enrichment score of the most relevant ACR summit (ACRs), and the distances to the nearest ACR summit in the upstream (ACRdu) and downstream (ACRdd) directions. In both species, ACR peaks were identified from ATAC-seq data using MACS2 (*17*). Distance calculations were based on ACR summit coordinates rather than peak boundaries, as summits represent the positions of maximum Tn5 transposase insertion frequency. For genes lacking an ACR in a given direction, the corresponding distance feature was assigned NaN. These values are biologically meaningful and were retained rather than exclusion of the gene from the dataset.

Maize: ATAC-seq peak calls were obtained from previous research (*18*), which identified 32,111 ACRs in young B73 leaves. For genes with one or more ACRs overlapping the gene body, the maximum summit fold enrichment among all overlapping ACRs was assigned as the ACRs feature. For genes without an overlapping ACR, the fold enrichment of the nearest flanking ACR summit was used. Upstream distance (ACRdu) was defined as the distance from the TSS to the nearest ACR summit in the upstream direction, whereas downstream distance (ACRdd) was defined from the TTS to the nearest downstream ACR summit. When an ACR overlapped the gene body, its summit was assigned to either the upstream or downstream category according to the nearest gene boundary. No maximum distance threshold was imposed. A small number of genes located at chromosome boundaries lacked an ACR in one direction and consequently received NaN for the corresponding distance feature (acr_Lup_M1: 1 gene; acr_Ldown_M1: 2 genes; acr_Lup_M2: 1 gene). These genes were retained and exempted from pair exclusion.

*B. rapa*: ACR data were obtained from three previous studies, combining peaks called across eight libraries into a single unified peak set (*17, 19-21*). Identification of ACR peaks followed the same workflow as maize (*18*). In brief, raw reads from six samples were trimmed using Trimmomatic v0.36 to remove adapters and low-quality reads (*22*). The trimmed reads were then aligned to the *B. rapa* v1.5 genome using Bowtie2 v2.5.4 under the following parameters -X 1000 -N 1 (*23*). PCR duplicates were removed with Picardtools. Peaks were called using MACS2 v2.2.7.1 (*17*) and merged across samples. Annotation and distribution of peaks were analyzed using Bedtools v2.30.0 (*24*). The same three features were calculated per gene using the same strand-aware procedure as described for maize. Genes located on scaffolds were assigned NaN for all three ACR features and were removed during the final data-filtering step. Genes located on main chromosomes that lacked an ACR in either the upstream or downstream direction received NaN for the corresponding distance feature and were retained in the dataset, consistent with the treatment applied in maize.

**DNA Methylation**

DNA methylation features comprise 15 variables per gene: nine mean methylation features and six maximum methylation features. Mean methylation was calculated for each combination of cytosine context (CG, CHG, and CHH) and genomic region (2 kb upstream region, gene body, and 2 kb downstream region), yielding nine features. Maximum methylation was calculated for each cytosine context in the upstream and downstream flanking regions only, yielding six additional features. Maximum gene body methylation was excluded from both species to avoid confounding effects associated with variation in gene length. All methylation values were bounded between 0 and 1 represented the proportion of methylated cytosines within a genomic window. Cytosines supported by fewer than three mapped reads were excluded from all analyses.

Maize: Whole-genome bisulfite sequencing (WGBS) data were obtained from our previous research from two biological replicates of immature ears of the reference line B73 (*25*). Data processing followed previously established pipelines (*6, 8, 25, 26*). Briefly, reads were aligned against the maize v4 reference genomes using Bismark with the parameters -n 2 -I 50 -N 1 (*27*), followed by PCR duplicate removal using the Bismark deduplication package. Cytosine methylation calls were extracted using Bismark methylation extractor, bismark2bedGraph, and coverage2cytosine. For feature calculation, the 2 kb upstream and downstream regions were divided into 40 fixed 50 bp windows. Gene bodies were divided into 40 proportionally sized bins to accommodate variation in gene length. Mean methylation was calculated per window across each region, and the two biological replicates were averaged at the gene level before integration into the feature matrix. Genes absent from the methylation dataset were assigned NaN for all methylation features, and their associated pairs were excluded during the final data-filtering step.

*B. rapa*: Whole genome bisulfite methylation levels were obtained from wild type leaves of three biological replicates and mapped to the *B. rapa* v1.5 genome (*28*). All feature definitions, cytosine contexts (CG, CHG, CHH), genomic regions (2 kb upstream, gene body, 2 kb downstream), windowing strategy (40 windows per region), filtering threshold (<3 mapped reads), and exclusion of maximum gene-body methylation were identical to those used in maize. Mean methylation was calculated as the average methylation value across windows within each region, and maximum methylation was defined as the highest single-window value within each flanking region. Genes absent from the methylation dataset were assigned NaN for all methylation features, and their associated gene pairs were removed during the final data-filtering step.

**Histone Modifications**

Histone modification data were quantified across three genomic regions per gene: the 2 kb upstream (Hu), the gene body (Hg), and the 2 kb downstream (Hd). Signal enrichment was computed in 40 windows per region as $\log_2$((normalized treatment reads + 1) / (normalized input reads + 1)). Upstream and downstream flanking regions used fixed ~50 bp windows, while gene bodies were divided into proportionally sized windows to accommodate variation in gene lengths, following the same approach as for DNA methylation. Window values were then averaged to yield a single signal score per gene per region.

Maize: Histone modification data for maize were obtained from previous research (*18*), which profiled ChIP-seq in young B73 seedling leaves. Eight chromatin marks were included: seven histone modifications (H3K4me1, H3K4me3, H3K9ac, H3K27ac, H3K27me3, H3K36me3, and H3K56ac) and one histone variant (H2A.Z). The BED file containing reads mapped to the B73 v4 reference genome (*18*) was used to calculate histone enrichment for each mark within each genomic window. All 9,156 genes in the analysis were present across all histone feature files prior to merging, and no missing data were introduced at this step.

*B. rapa*: Histone modification data for three marks, H3K27ac, H3K27me3, and H3K4me3, were obtained from previous research (*20*). Raw reads were trimmed using Trimmomatic v0.36 (*22*), aligned to the *B. rapa* v1.5 reference genome using Bowtie2 v2.5.4 (-X 1000 -N 1) (*23*), and PCR duplicates were removed using Picard. Signal enrichment was quantified and averaged across windows using the same procedure described for maize, yielding nine feature values per gene. All 31,806 gene pairs were present across all histone feature files with no missing data introduced at the merge step.

**Data S1.** Mann-Whitney U test statistics and effect sizes for pairwise comparisons of feature distribution between subgenomes in maize and *B. rapa*.

**Data S2.** Spearman correlation coefficients between features, including pairwise feature correlations and correlations with recombination rate in maize and *B. rapa*.

**Data S3.** Mann-Whitney U test results comparing raw feature values between recombination rate SHAP value clusters (Fig. 3B) in maize group1.

**Data S4.** SHAP covariate network edge weights and node degrees for maize and *B. rapa*.

**Data S5.** SHAP interaction network edge weights and node degrees for maize and *B. rapa*.

**Data S6.** Maize SHAP interaction network edge weights and node degrees stratified by subgenome label.

**Data S7.** *B. rapa* SHAP interaction network edge weights and node degrees stratified by subgenome label.

**Data S8.** Ablation study performance statistics for maize and *B. rapa*.

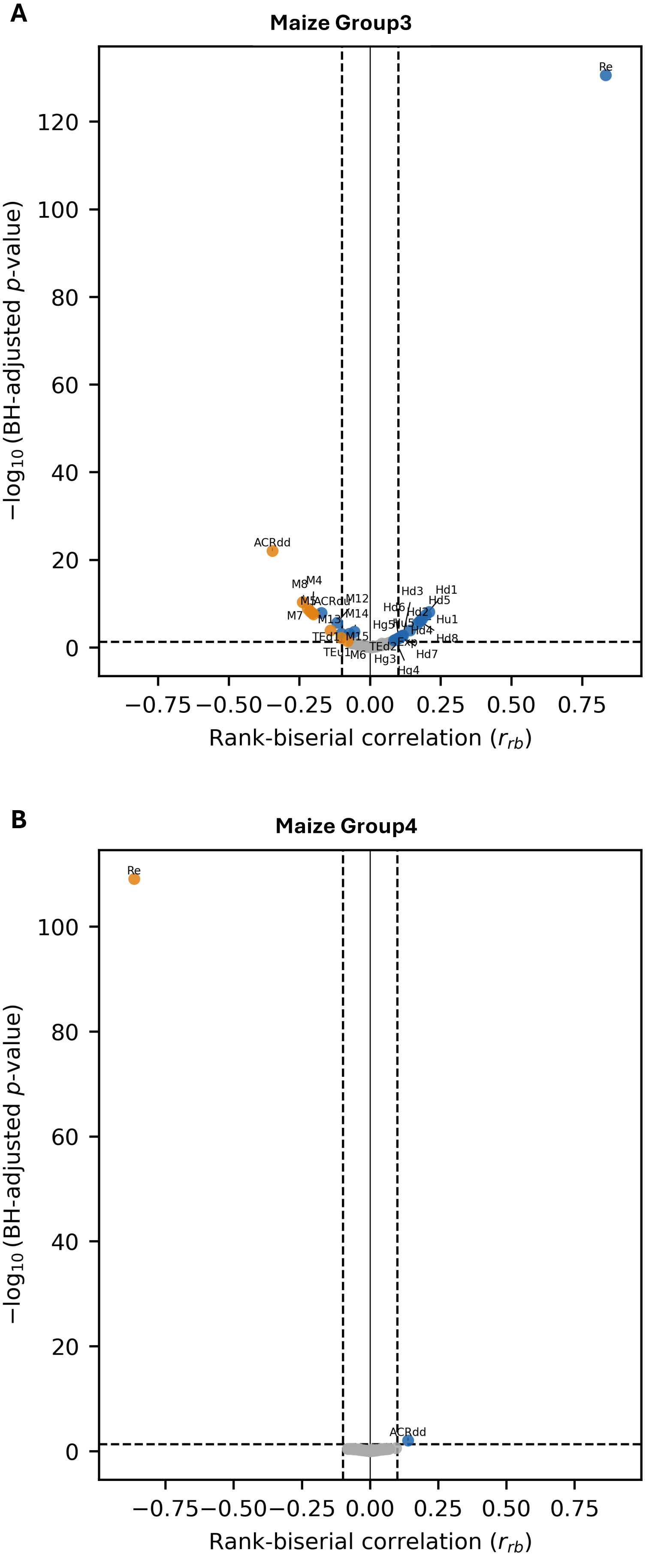


**Fig. S2. Volcano plots of maize feature distributions.** Volcano plots showing the results of two-sided Mann-Whitney U tests comparing genomic feature distributions between maize1 (dominant, WGD=0) and maize2 (recessive, WGD=1) subgenome homeologs for group3 (**A**) and group4 (**B**). The x-axis shows the rank-biserial correlation (*r*~rb~), a measure of effect size ranging from −1 to +1. The y-axis shows the −log~10~ Benjamini-Hochberg (BH) FDR-adjusted *P*-value. Blue points indicate features with significantly higher values in maize1; orange points indicate features with significantly higher values in maize2. Grey points are not significant. The horizontal dashed line marks the FDR significance threshold (BH-adjusted *P* = 0.05). Vertical dashed lines mark an effect size threshold of |*r*~rb~| = 0.1. No features reached significance in the group2 comparison after FDR correction.

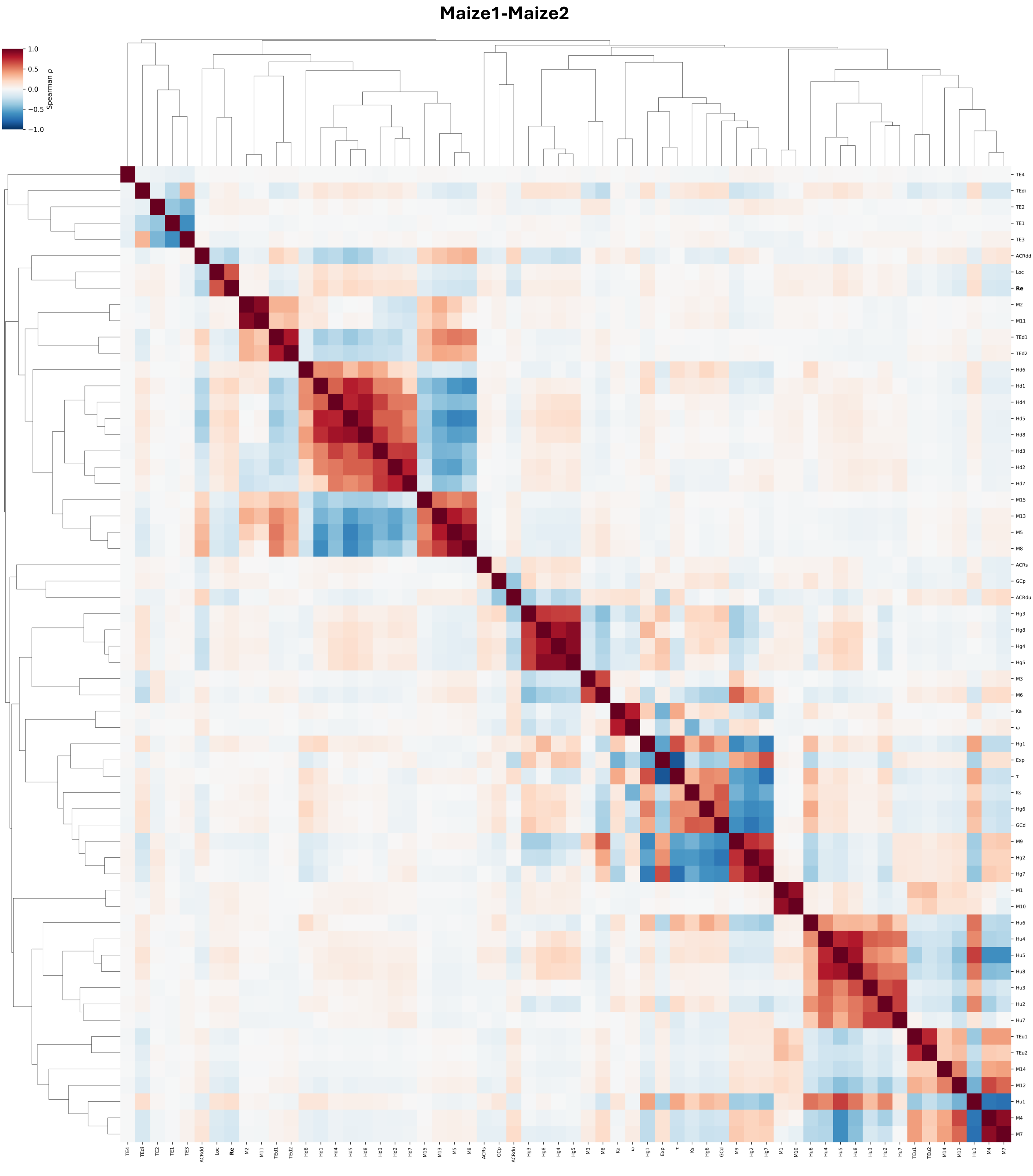


**Fig. S3. Spearman correlation heatmap of maize1-maize2 genomic and epigenomic features.** Pairwise Spearman rank correlation coefficients were computed across all features using complete pairwise deletion for missing values. Features are clustered on both axes using average linkage on a distance matrix defined as 1 − |ρ|, where ρ is the Spearman correlation coefficient; this metric groups co-varying feature blocks symmetrically regardless of the direction of correlation. The color scale represents the Spearman ρ value (red = positive correlation, blue = negative correlation). Recombination rate (Re) is shown in bold.

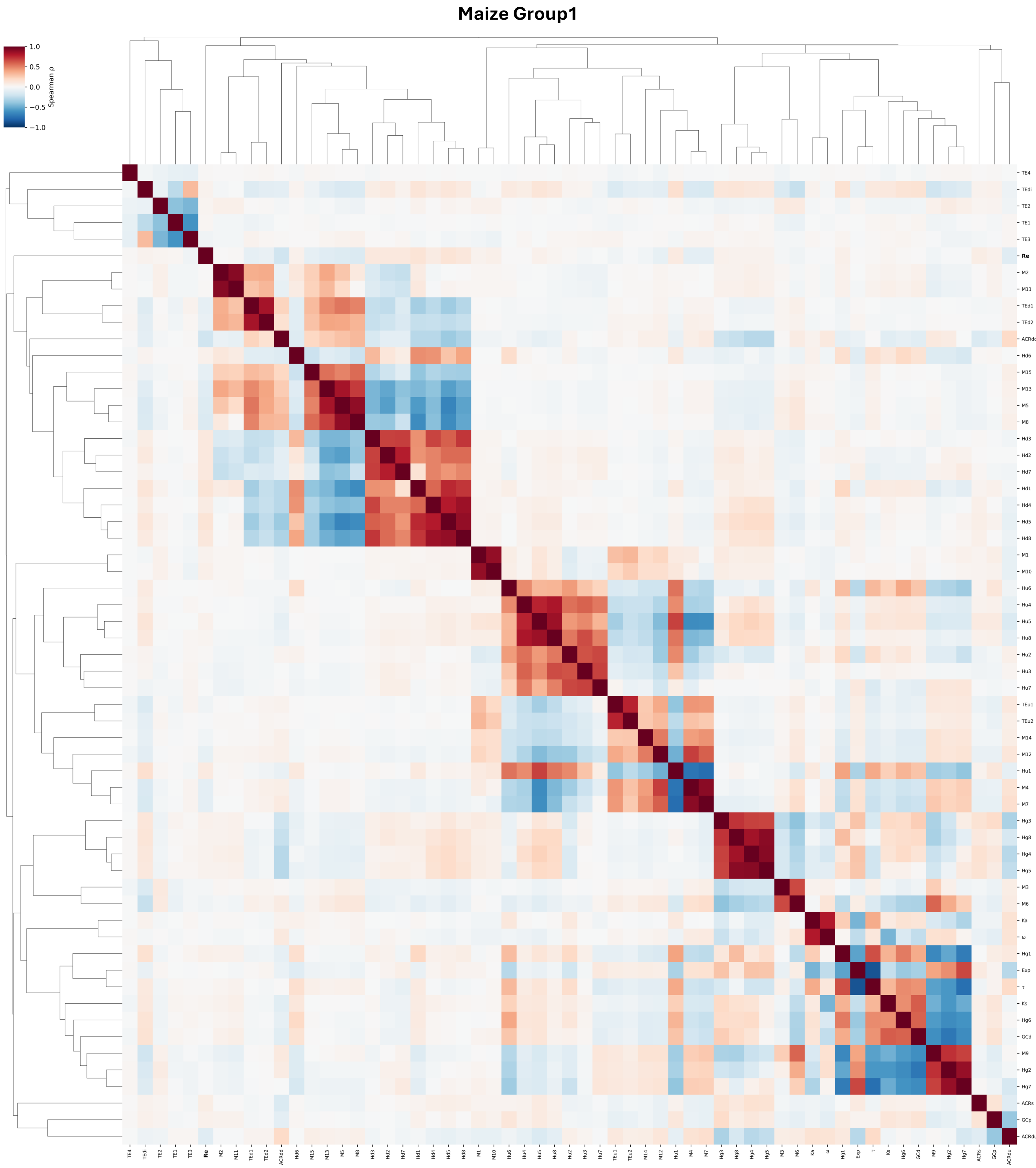


**Fig. S4. Spearman correlation heatmap of maize group1 genomic and epigenomic features.** Pairwise Spearman rank correlation coefficients were computed across all features using complete pairwise deletion for missing values. Features are clustered on both axes using average linkage on a distance matrix defined as 1 − |ρ|, where ρ is the Spearman correlation coefficient; this metric groups co-varying feature blocks symmetrically regardless of the direction of correlation. The color scale represents the Spearman ρ value (red = positive correlation, blue = negative correlation). Recombination rate (Re) is shown in bold.

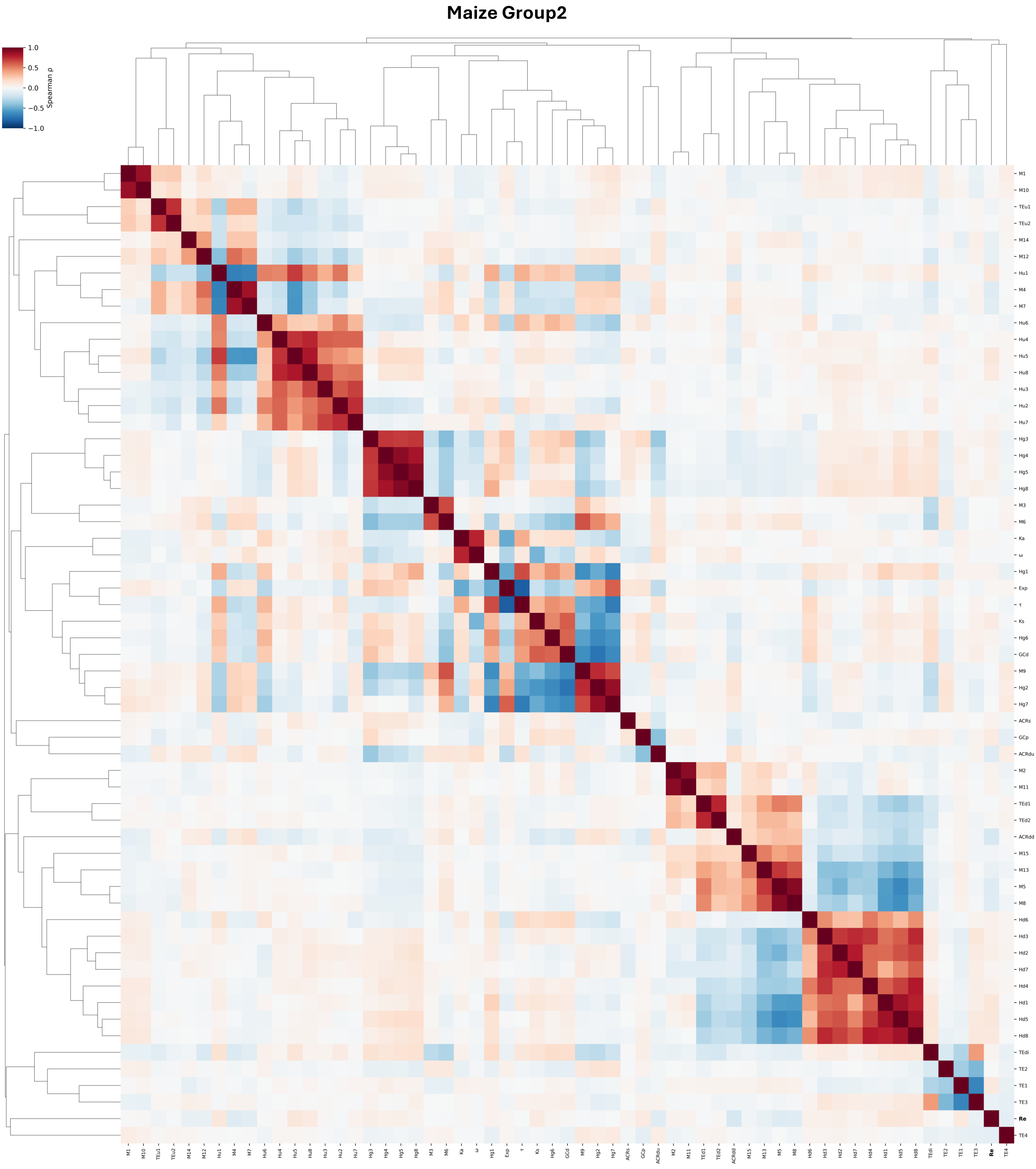


**Fig. S5. Spearman correlation heatmap of maize group2 genomic and epigenomic features.** Pairwise Spearman rank correlation coefficients were computed across all features using complete pairwise deletion for missing values. Features are clustered on both axes using average linkage on a distance matrix defined as 1 − |ρ|, where ρ is the Spearman correlation coefficient; this metric groups co-varying feature blocks symmetrically regardless of the direction of correlation. The color scale represents the Spearman ρ value (red = positive correlation, blue = negative correlation). Recombination rate (Re) is shown in bold.

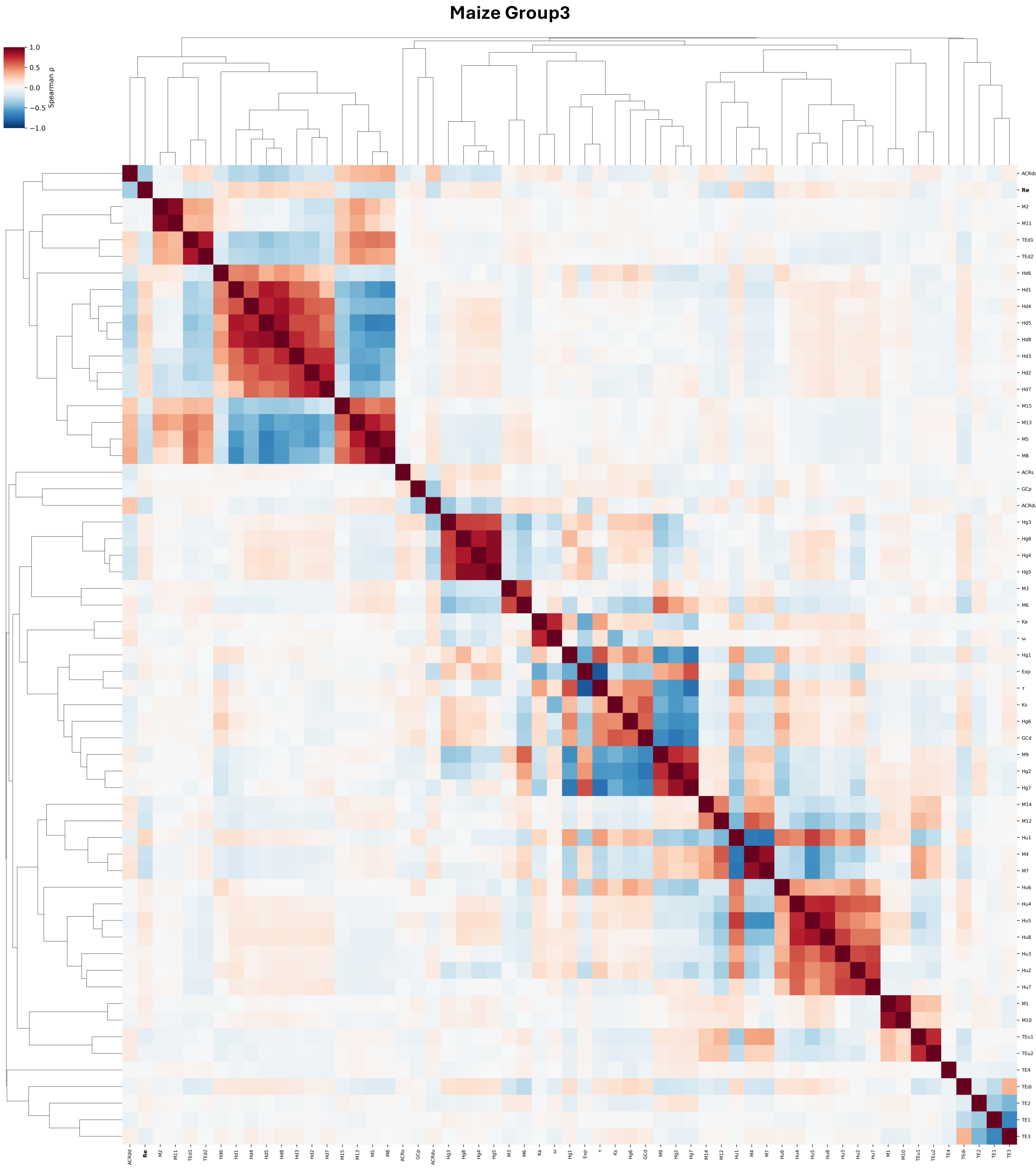


**Fig. S6. Spearman correlation heatmap of maize group3 genomic and epigenomic features.** Pairwise Spearman rank correlation coefficients were computed across all features using complete pairwise deletion for missing values. Features are clustered on both axes using average linkage on a distance matrix defined as 1 − |ρ|, where ρ is the Spearman correlation coefficient; this metric groups co-varying feature blocks symmetrically regardless of the direction of correlation. The color scale represents the Spearman ρ value (red = positive correlation, blue = negative correlation). Recombination rate (Re) is shown in bold.

**Maize Group4**

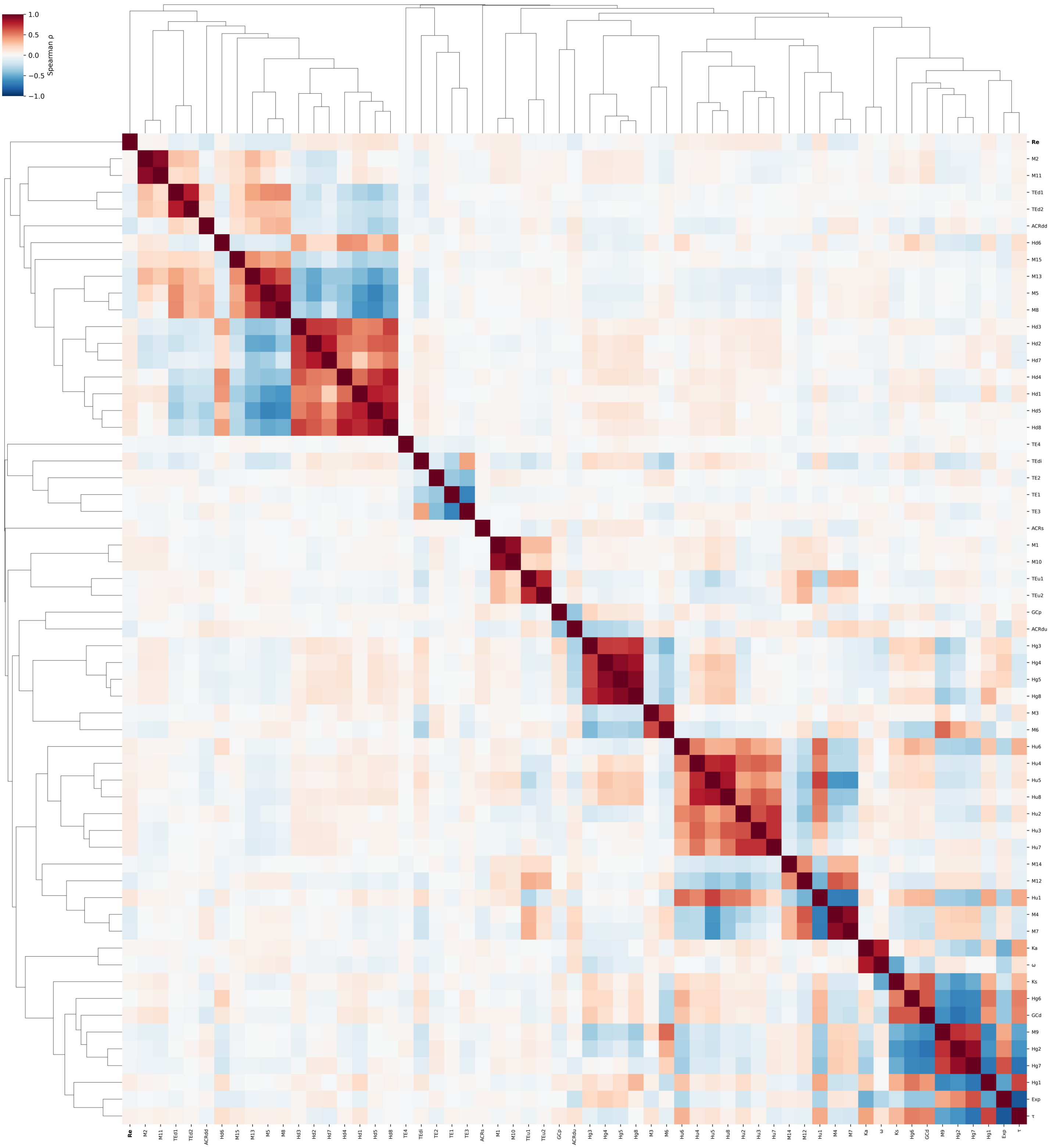


**Fig. S7. Spearman correlation heatmap of maize group4 genomic and epigenomic features.** Pairwise Spearman rank correlation coefficients were computed across all features using complete pairwise deletion for missing values. Features are clustered on both axes using average linkage on a distance matrix defined as 1 − |ρ|, where ρ is the Spearman correlation coefficient; this metric groups co-varying feature blocks symmetrically regardless of the direction of correlation. The color scale represents the Spearman ρ value (red = positive correlation, blue = negative correlation). Recombination rate (Re) is shown in bold.

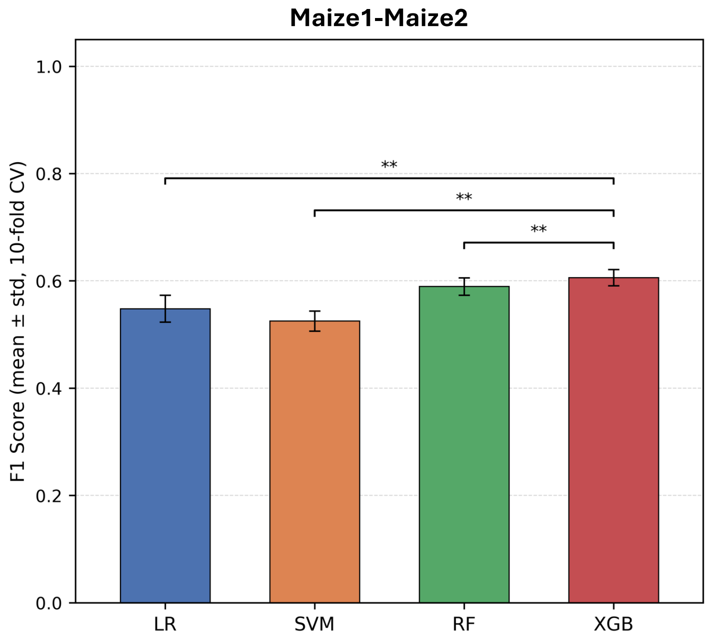


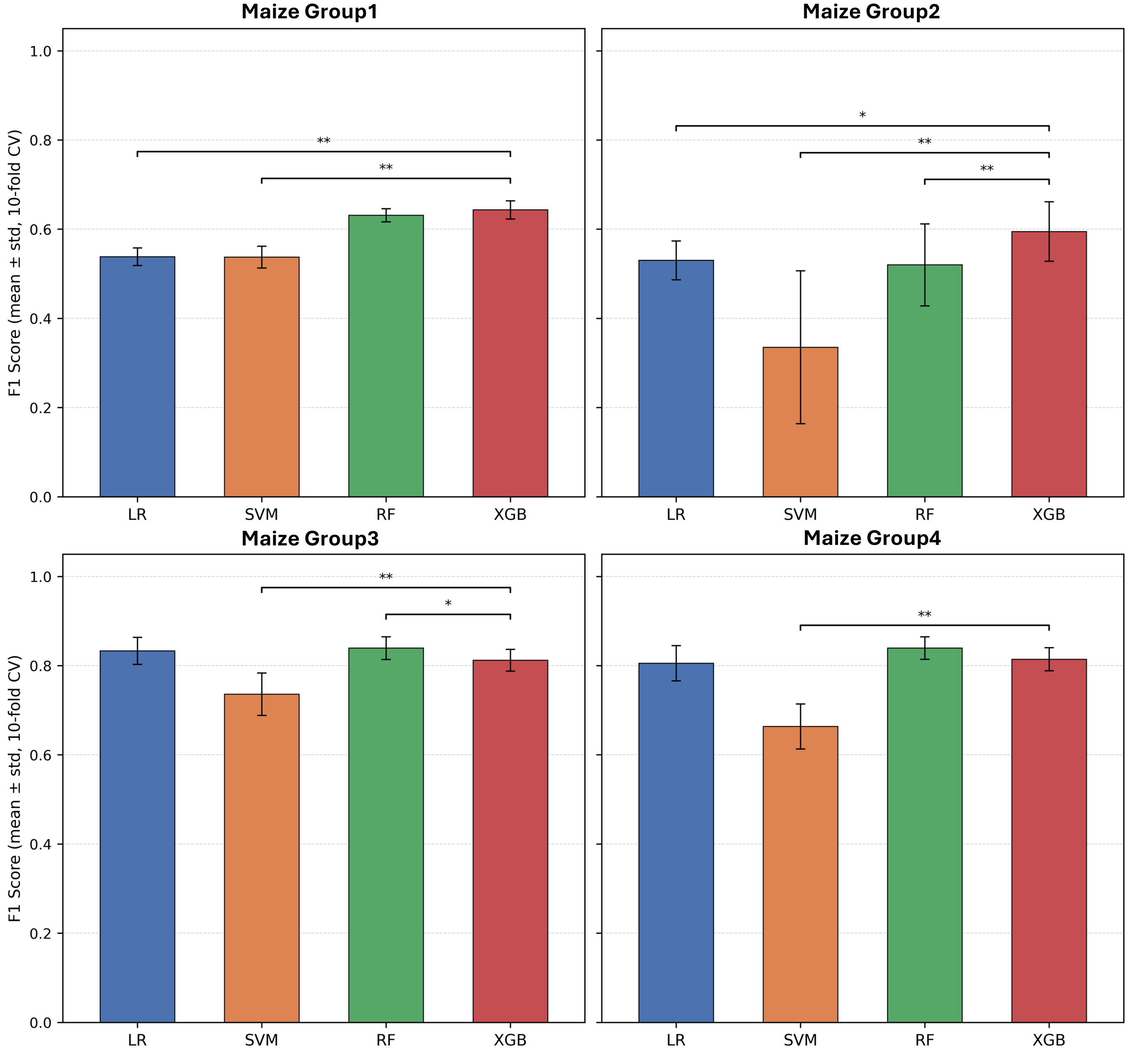


**Fig. S8. Comparison of classification performance across model architectures for each maize gene group.** Mean F1 score (± standard deviation across 10 stratified cross-validation folds) is shown for each model architecture: Logistic Regression (LR), Support Vector Machine (SVM), Random Forest (RF), and XGBoost (XGB). Pairwise significance was assessed between each baseline architecture and XGB using two-sided Wilcoxon signed-rank tests on paired fold-level F1 scores, with Benjamini-Hochberg false discovery rate correction applied within each group independently. Only comparisons reaching adjusted $P < 0.05$ are annotated (* $P < 0.05$, ** $P < 0.01$, *** $P < 0.001$). Comparisons across groups are not made. All models used default hyperparameters; feature scaling was applied per fold for LR and SVM only.

### Maize1-Maize2

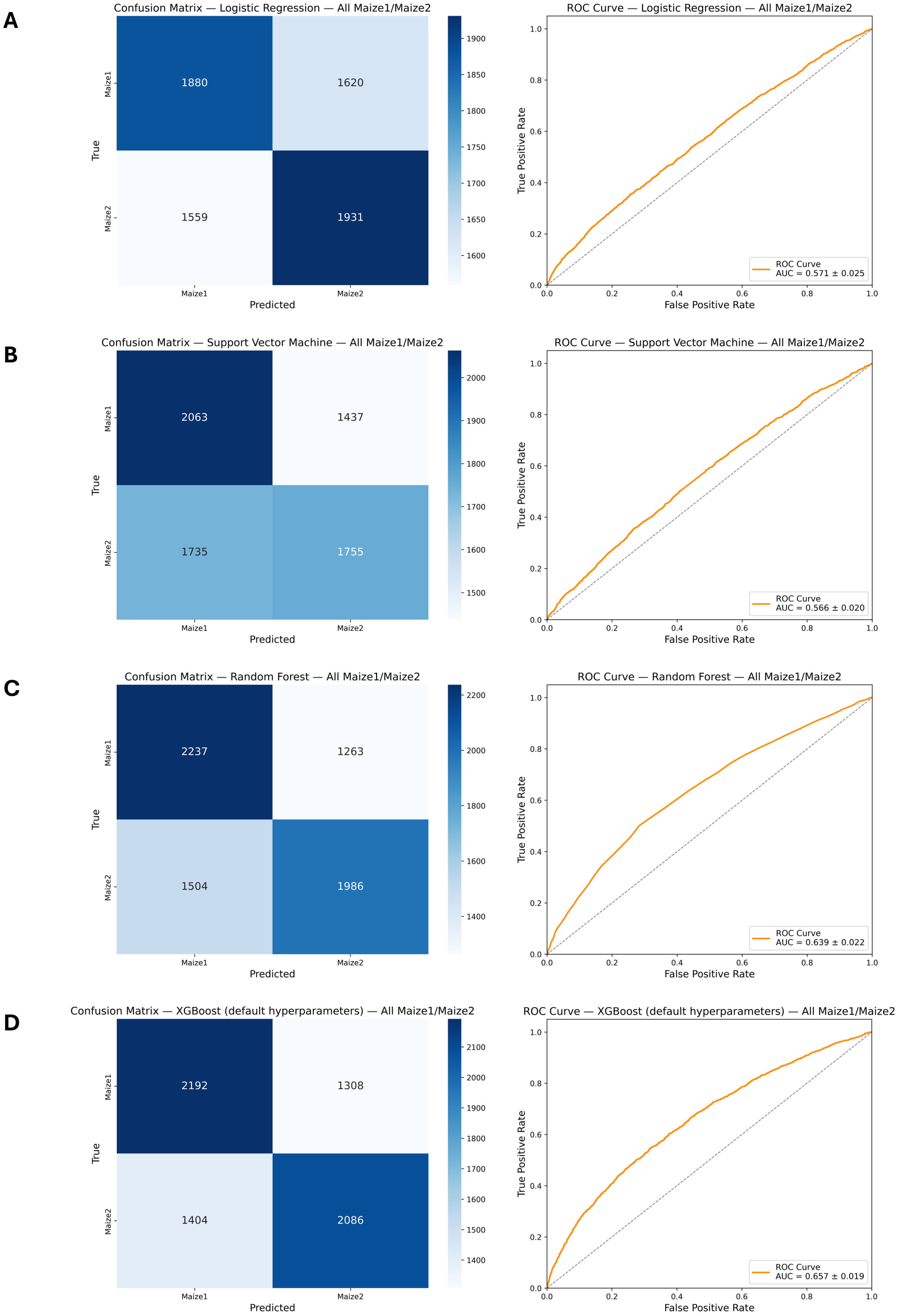


**Fig. S9. Benchmarking of unoptimized model performance on maize1-maize2 dataset.** Confusion matrices and ROC/AUC from logistic regression (**A**), support vector machine (**B**), random forest (**C**), and XGBoost (**D**) model architectures.

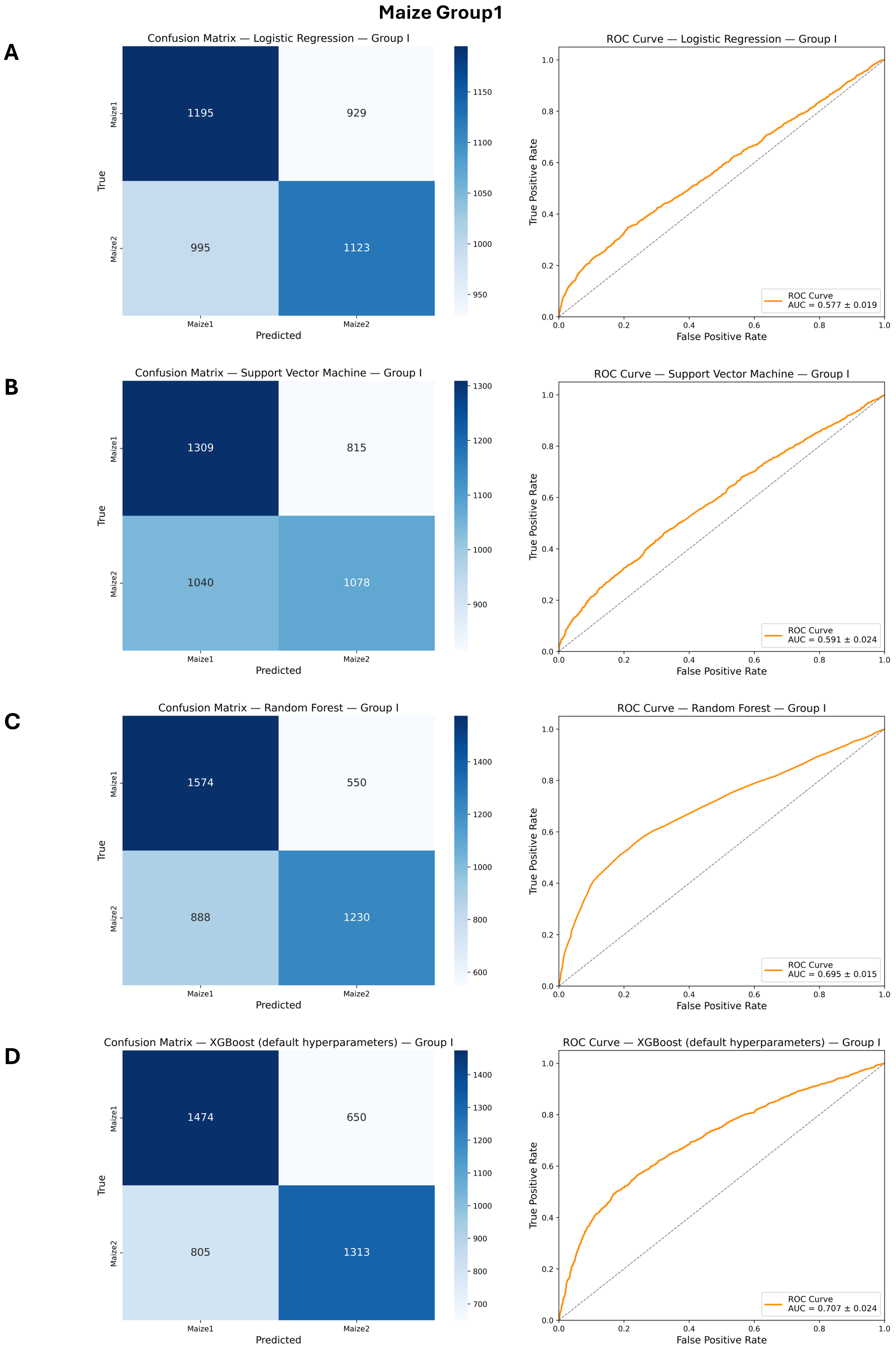


**Fig. S10. Benchmarking of unoptimized model performance on maize group1 dataset.** Confusion matrices and ROC/AUC from logistic regression (**A**), support vector machine (**B**), random forest (**C**), and XGBoost (**D**) model architectures.

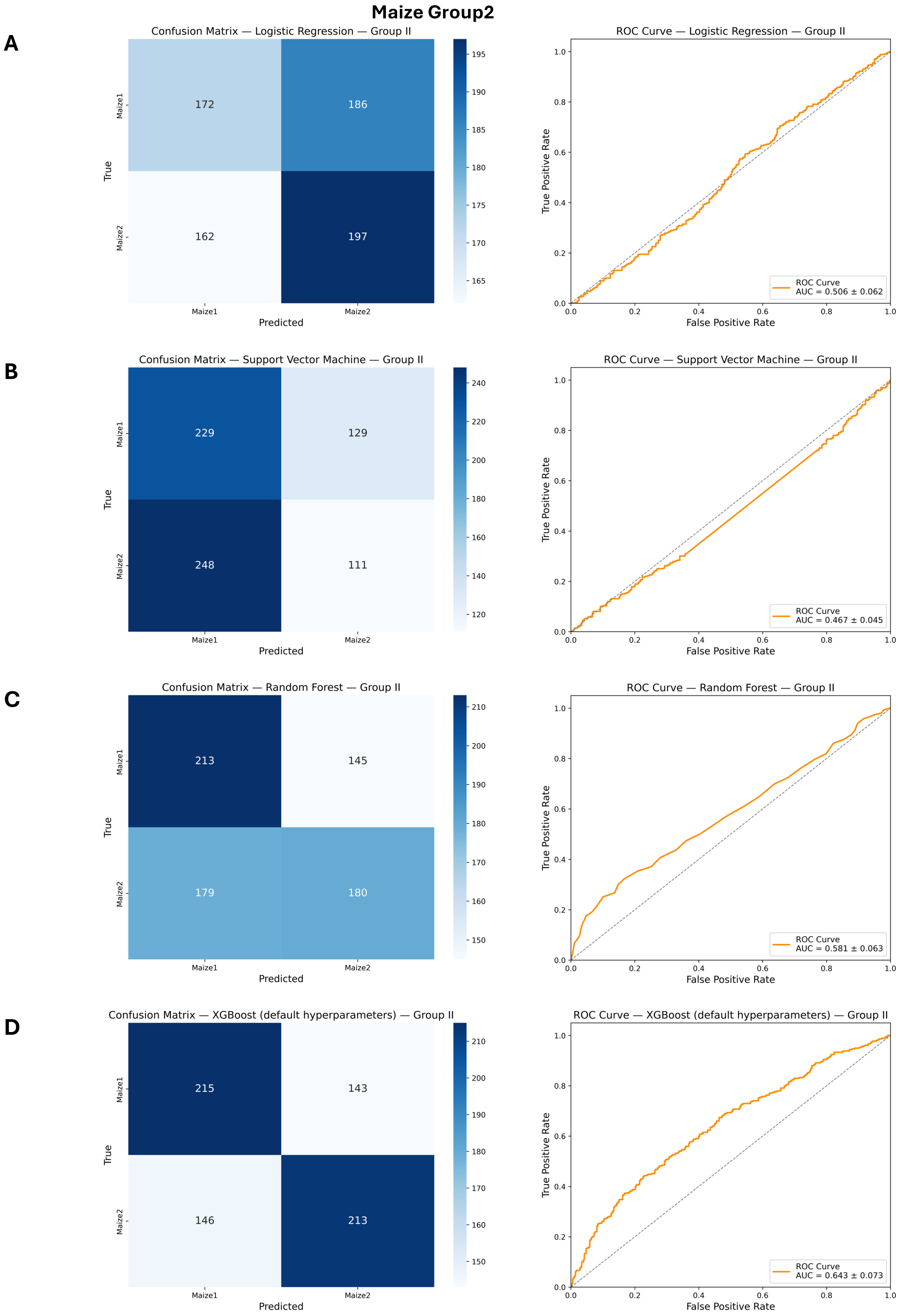


**Fig. S11. Benchmarking of unoptimized model performance on maize group2 dataset.** Confusion matrices and ROC/AUC from logistic regression (**A**), support vector machine (**B**), random forest (**C**), and XGBoost (**D**) model architectures.

**A**

| | Mean F1 ± std |
|---|---|
| **Maize1-Maize2** | 0.643±0.017 |
| **Group1** | 0.650±0.023 |
| **Group2** | 0.587±0.044 |
| **Group3** | 0.852±0.029 |
| **Group4** | 0.839±0.037 |

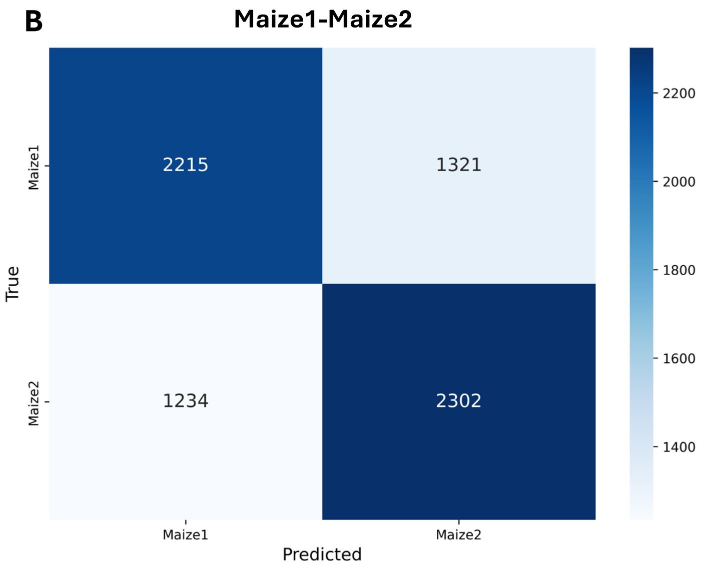


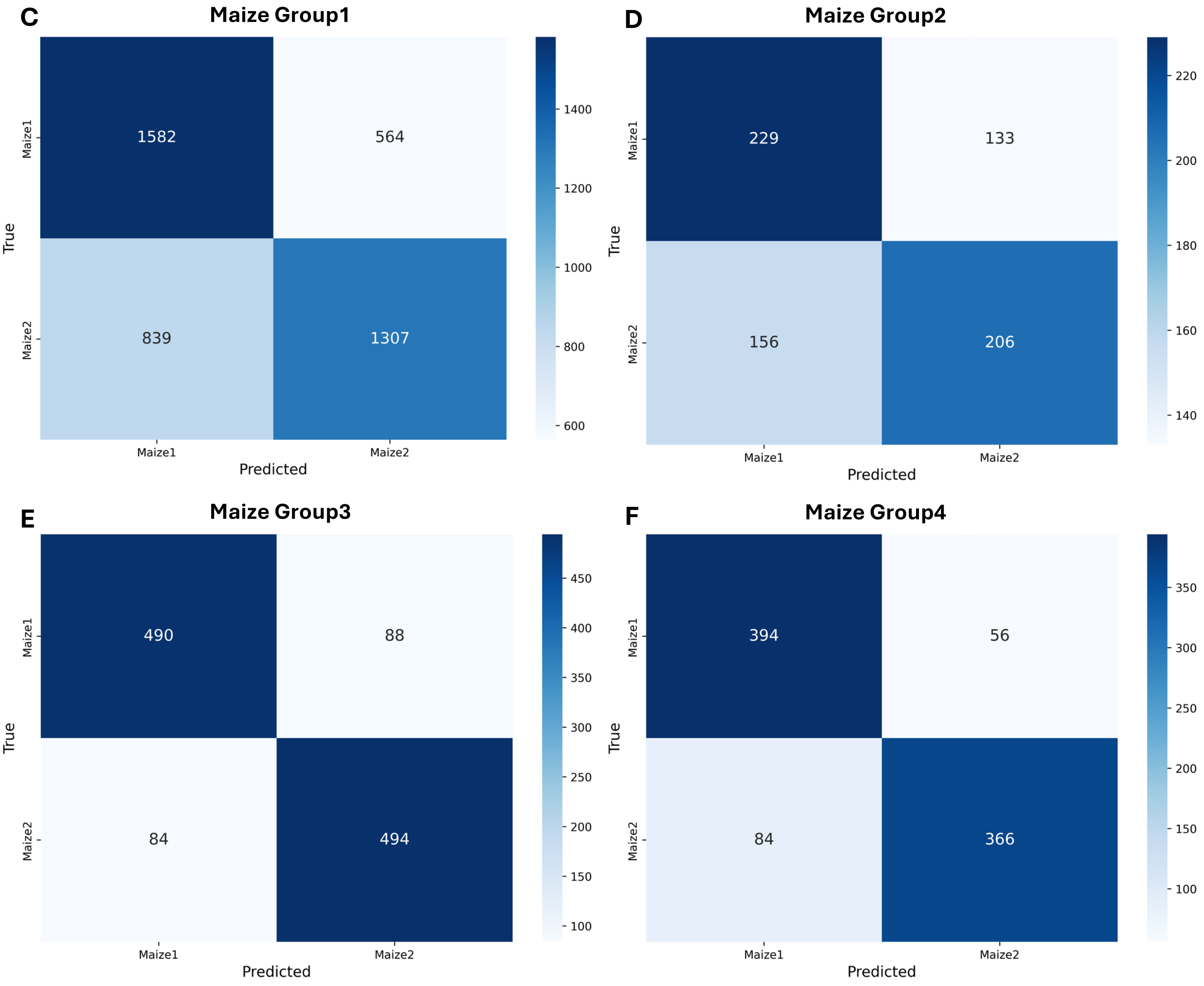


**Fig. S12. Performance metrics for five final XGBoost maize models.** F1 scores for five final maize models, mean and standard deviation across 10 folds (**A**) and confusion matrices for maize1-maize2 all-genes (**B**), group1 (**C**), group2 (**D**), group3 (**E**), group4 (**F**) models.

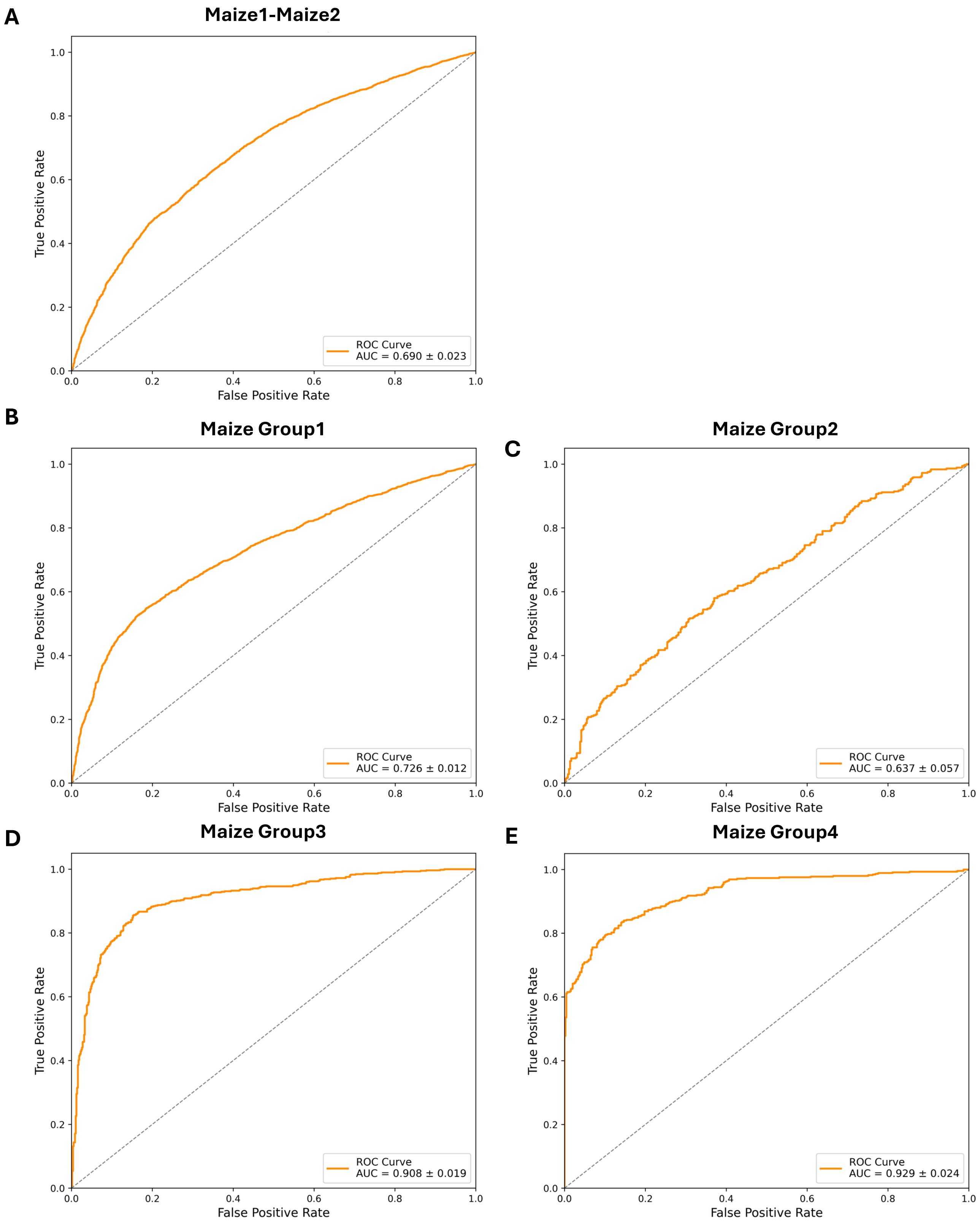


**Fig. S13. ROC/AUC for five final maize models.** ROC curves for maize1-maize2 all-genes model (**A**), group1 (**B**), group2 (**C**), group3 (**D**), and group4 (**E**) showing mean AUC score and standard deviation across 10 folds.

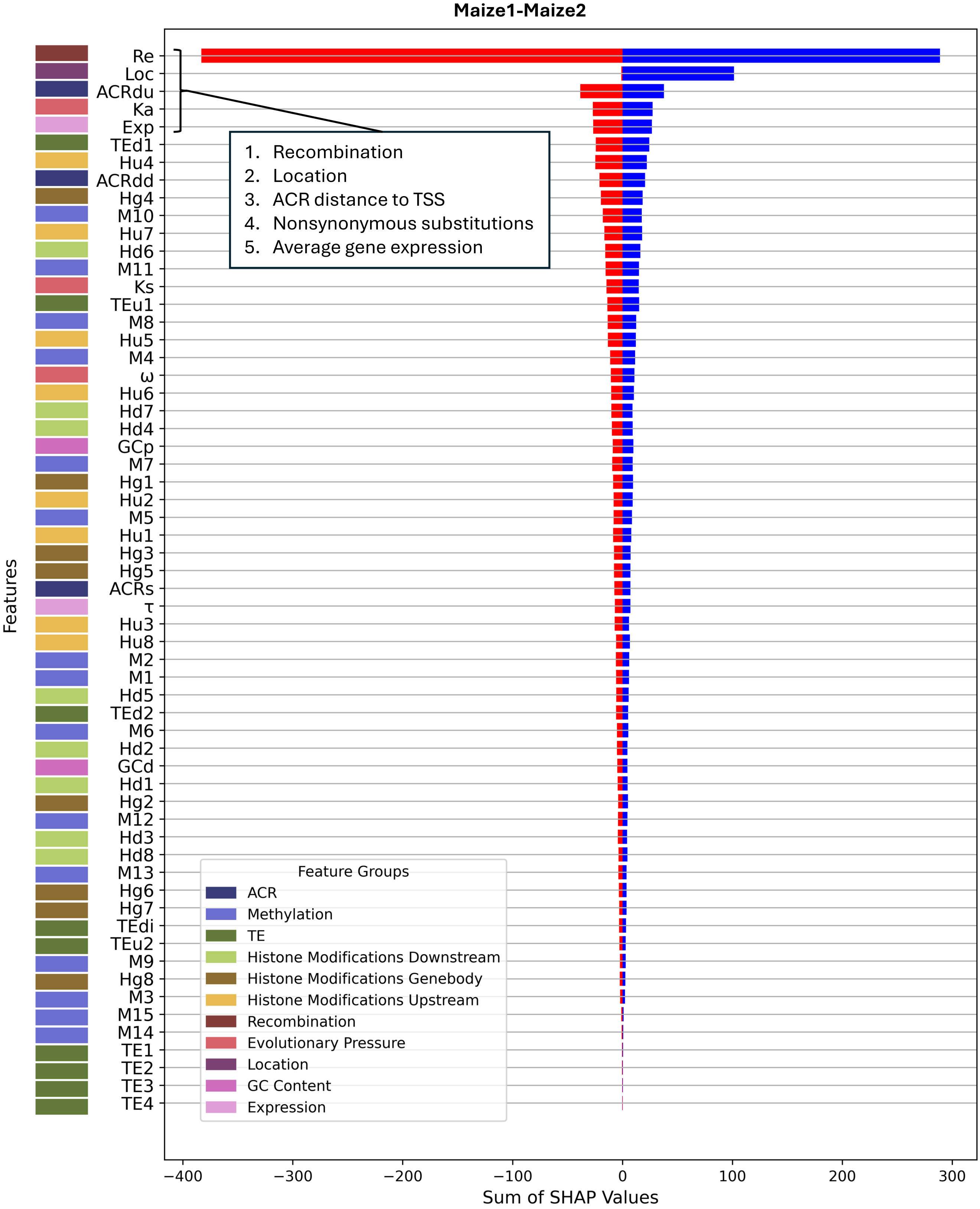


**Fig. S14. Directional SHAP feature importance profile for maize1-maize2.** Features are ranked by the sum of their absolute SHAP values across all test-fold samples (ascending order, highest-importance features at top). For each feature, SHAP values are summed separately across positive contributions (blue; favoring maize2 classification) and negative contributions (red; favoring maize1 classification), reflecting total directional influence across the test set rather than per-sample magnitude. Colored tiles in the left margin indicate feature group membership. The five features contributing most to classification, as determined by summed absolute SHAP value, are listed in the inset.

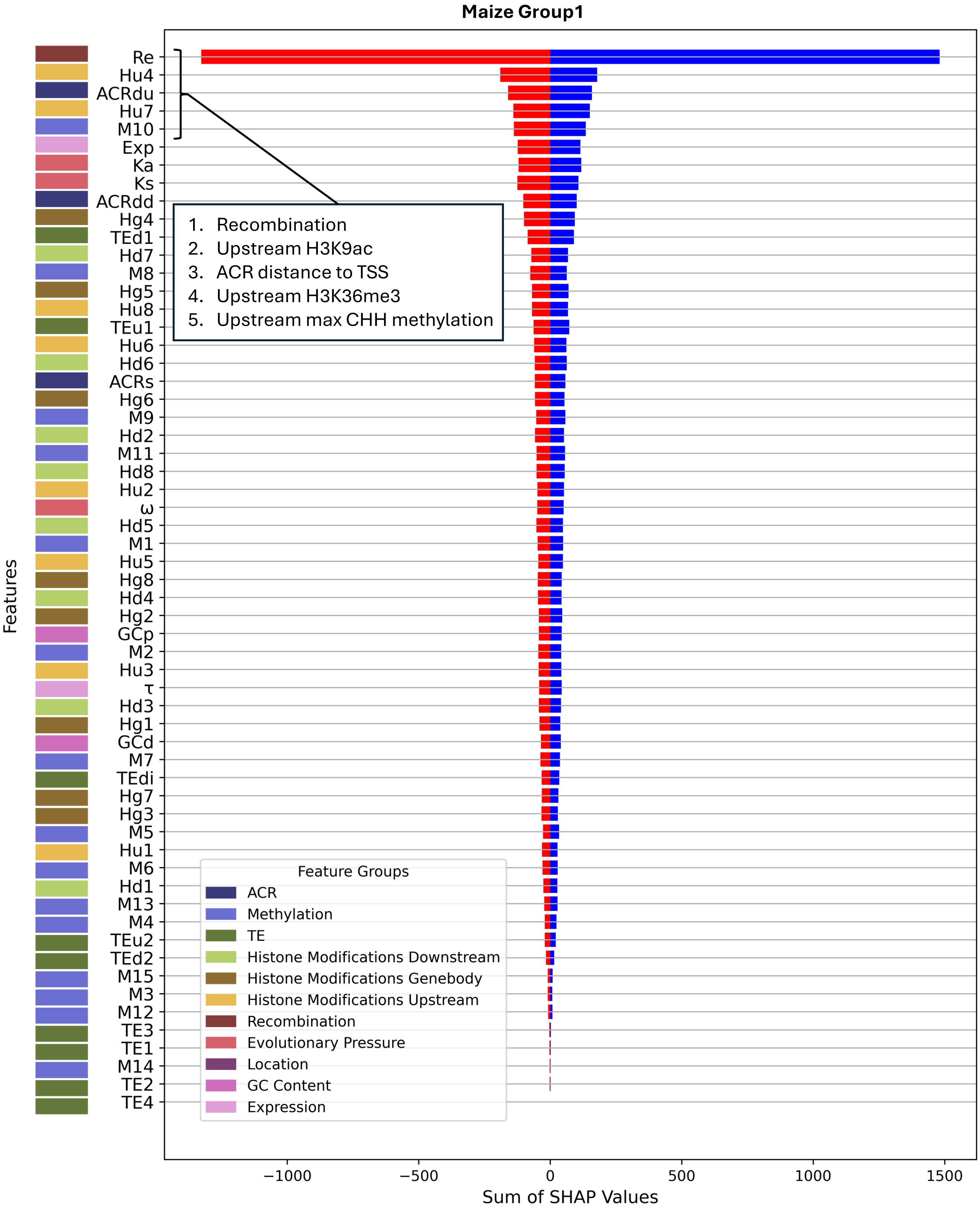


**Fig. S15. Directional SHAP feature importance profile for maize group1.** Features are ranked by the sum of their absolute SHAP values across all test-fold samples (ascending order, highest-importance features at top). For each feature, SHAP values are summed separately across positive contributions (blue; favoring maize2 classification) and negative contributions (red; favoring maize1 classification), reflecting total directional influence across the test set rather than per-sample magnitude. Colored tiles in the left margin indicate feature group membership. The five features contributing most to classification, as determined by summed absolute SHAP value, are listed in the inset.

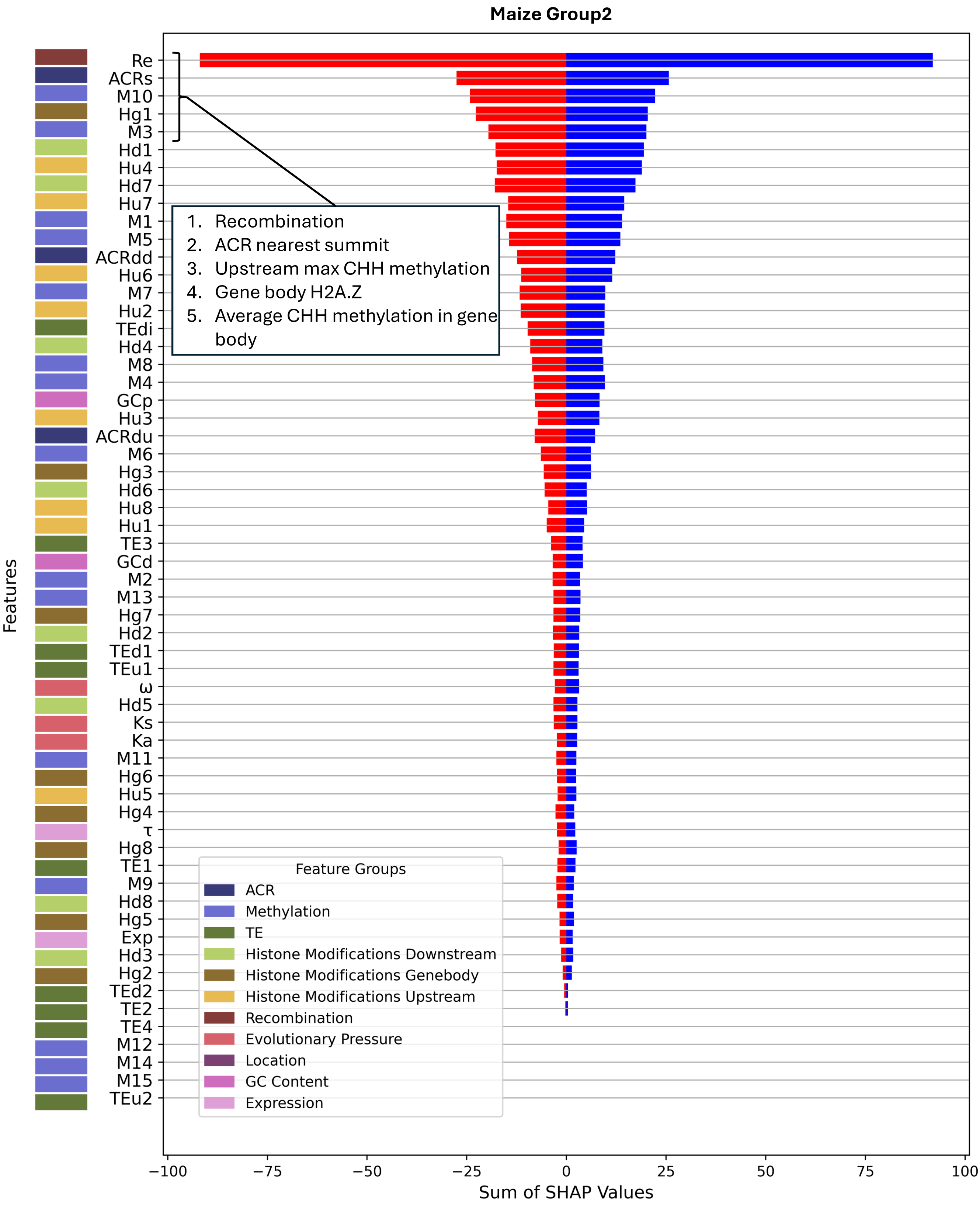


**Fig. S16. Directional SHAP feature importance profile for maize group2.** Features are ranked by the sum of their absolute SHAP values across all test-fold samples (ascending order, highest-importance features at top). For each feature, SHAP values are summed separately across positive contributions (blue; favoring maize2 classification) and negative contributions (red; favoring maize1 classification), reflecting total directional influence across the test set rather than per-sample magnitude. Colored tiles in the left margin indicate feature group membership. The five features contributing most to classification, as determined by summed absolute SHAP value, are listed in the inset.

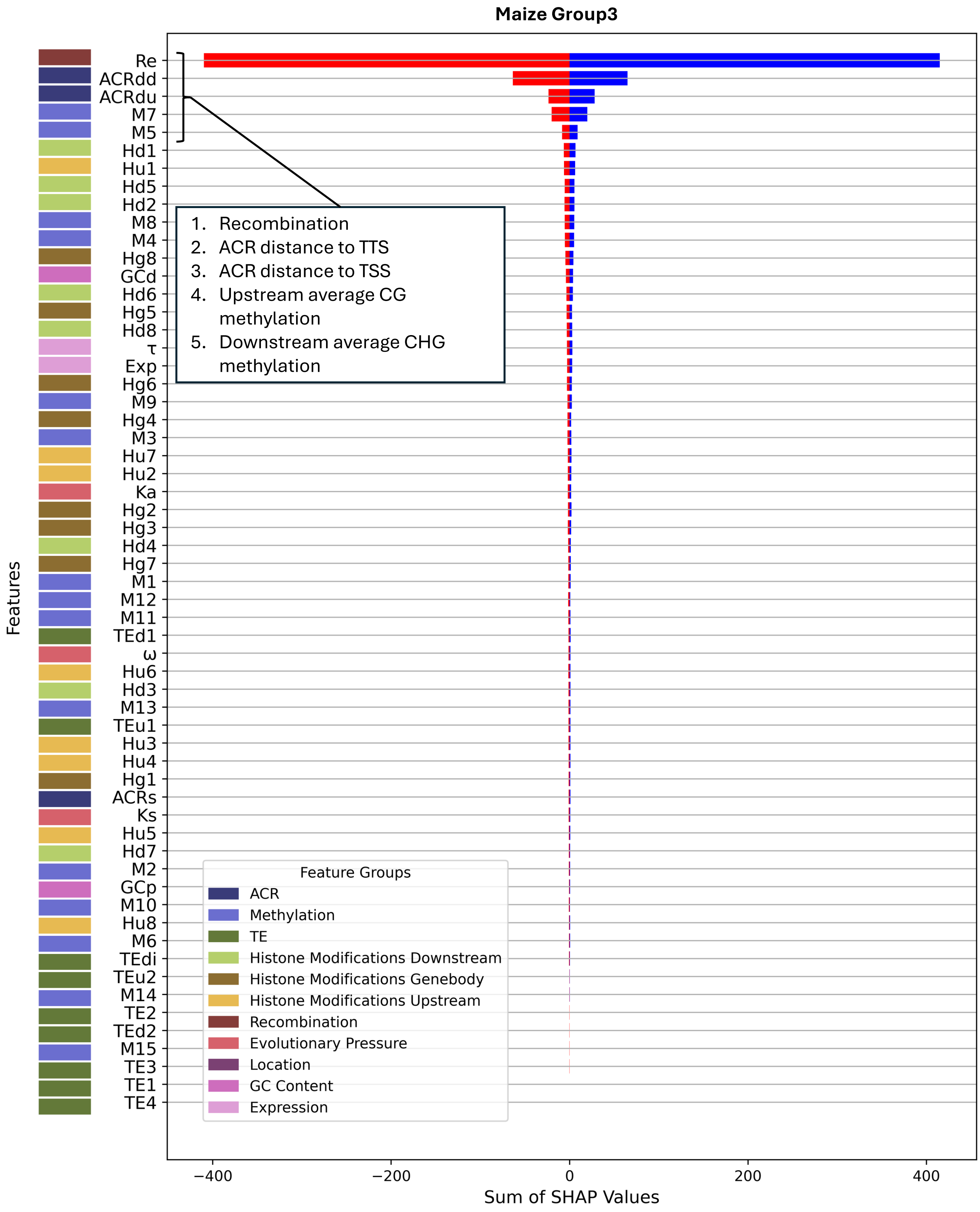


**Fig. S17. Directional SHAP feature importance profile for maize group3.** Features are ranked by the sum of their absolute SHAP values across all test-fold samples (ascending order, highest-importance features at top). For each feature, SHAP values are summed separately across positive contributions (blue; favoring maize2 classification) and negative contributions (red; favoring maize1 classification), reflecting total directional influence across the test set rather than per-sample magnitude. Colored tiles in the left margin indicate feature group membership. The five features contributing most to classification, as determined by summed absolute SHAP value, are listed in the inset.

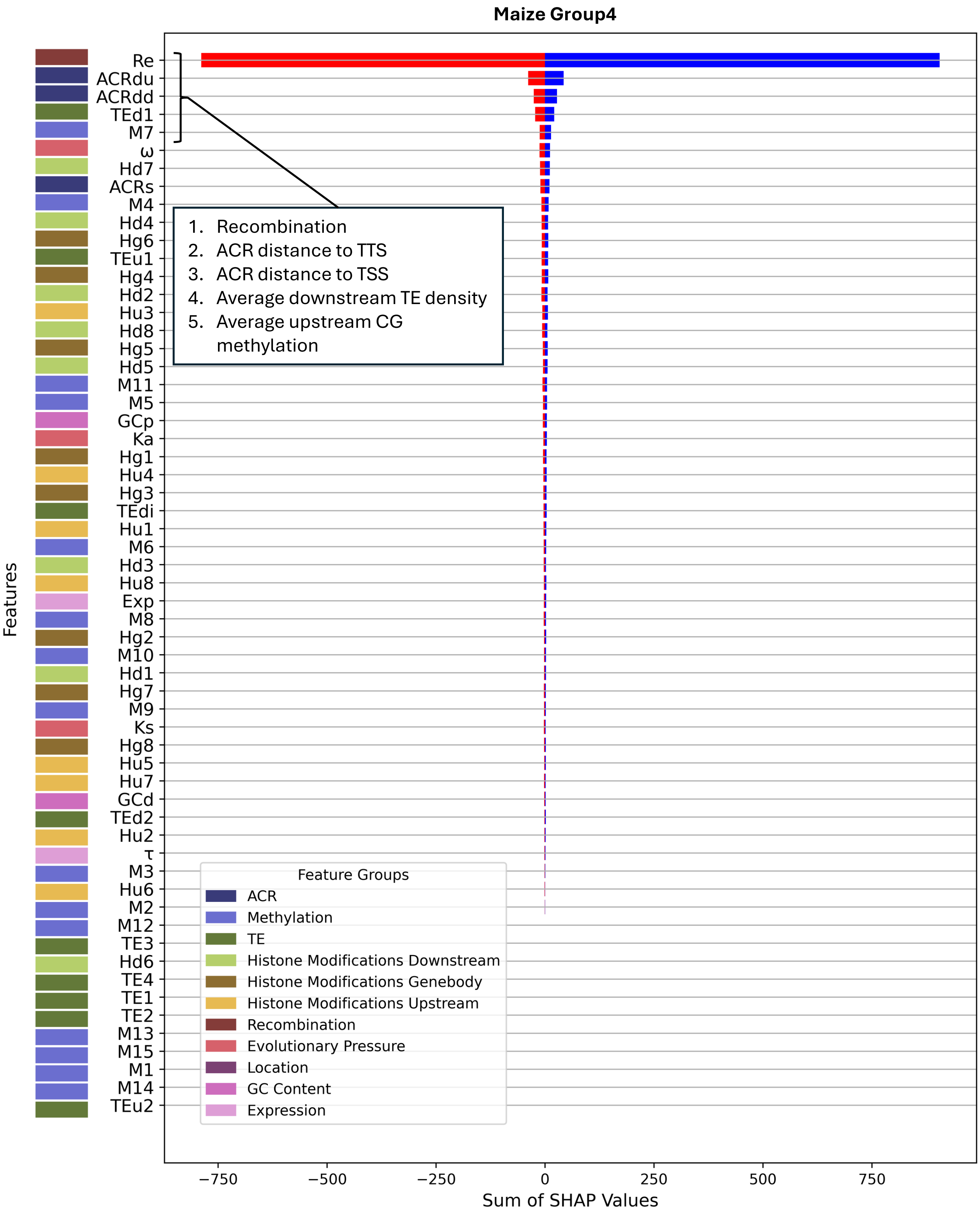


**Fig. S18. Directional SHAP feature importance profile for maize group4.** Features are ranked by the sum of their absolute SHAP values across all test-fold samples (ascending order, highest-importance features at top). For each feature, SHAP values are summed separately across positive contributions (blue; favoring maize2 classification) and negative contributions (red; favoring maize1 classification), reflecting total directional influence across the test set rather than per-sample magnitude. Colored tiles in the left margin indicate feature group membership. The five features contributing most to classification, as determined by summed absolute SHAP value, are listed in the inset.

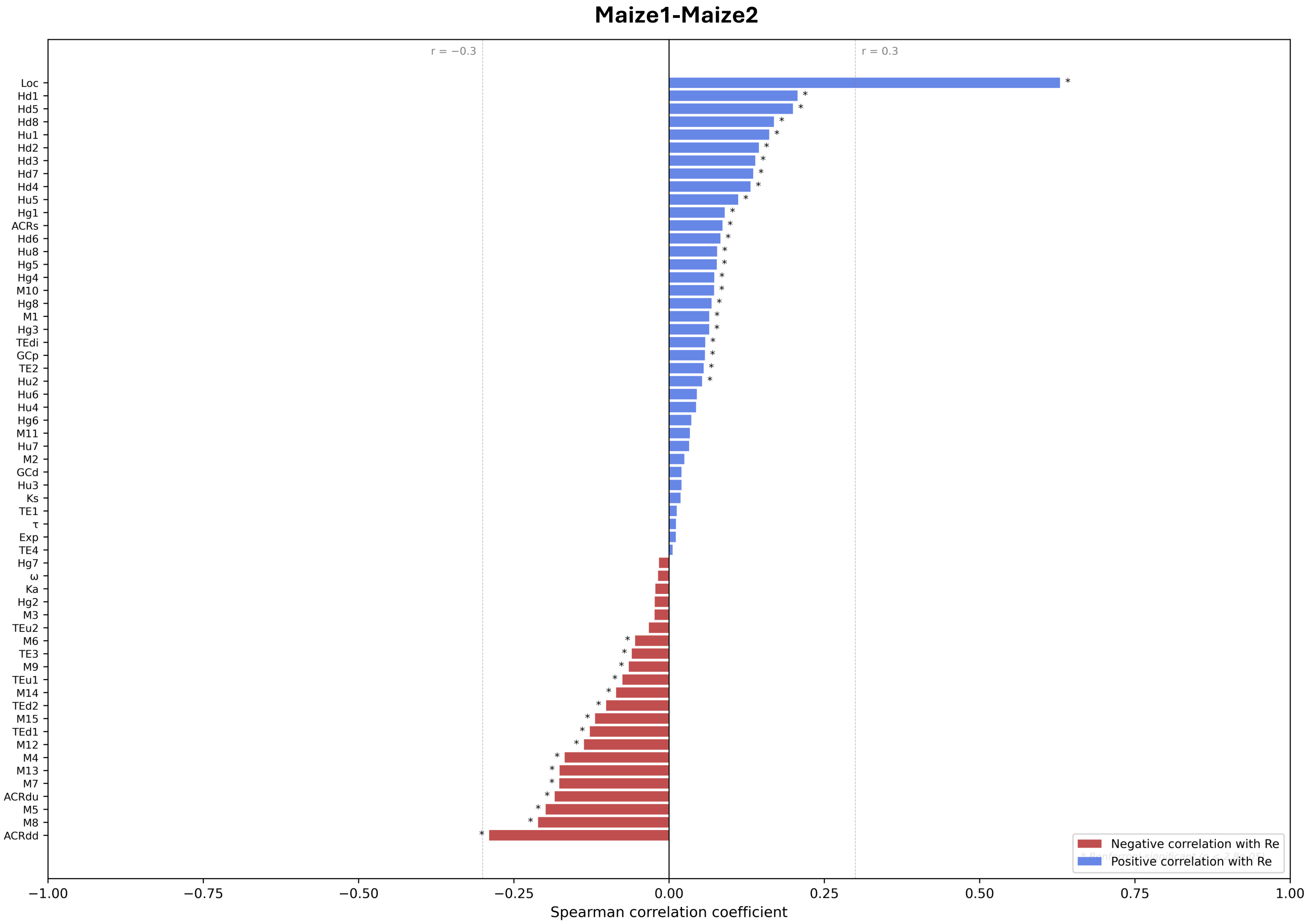


**Fig. S19. Spearman correlation of recombination rate (Re) with all maize1-maize2 features.** Features are ranked by Spearman ρ with Re (most negative at bottom). Bar color indicates direction of correlation: negative (red) or positive (blue). Asterisks indicate feature pairs surviving Bonferroni correction across all n(n−1)/2 pairwise tests within the group (α = 0.05). Dashed reference lines are drawn at ρ = ±0.3.

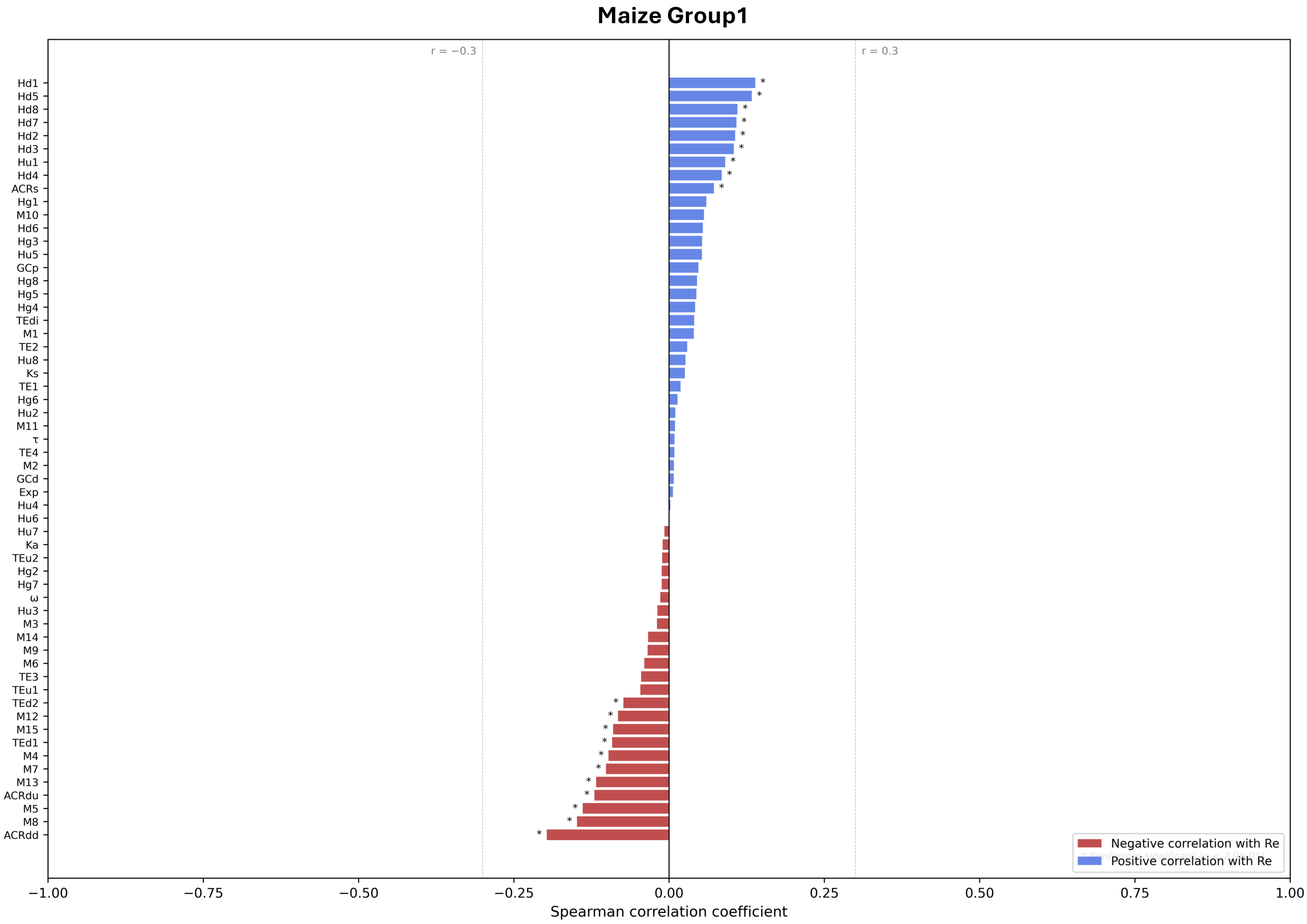


**Fig. S20. Spearman correlation of recombination rate (Re) with all maize group1 features.** Features are ranked by Spearman ρ with Re (most negative at bottom). Bar color indicates direction of correlation: negative (red) or positive (blue). Asterisks indicate feature pairs surviving Bonferroni correction across all $n(n-1)/2$ pairwise tests within the group ($\alpha = 0.05$). Dashed reference lines are drawn at $\rho = \pm 0.3$.

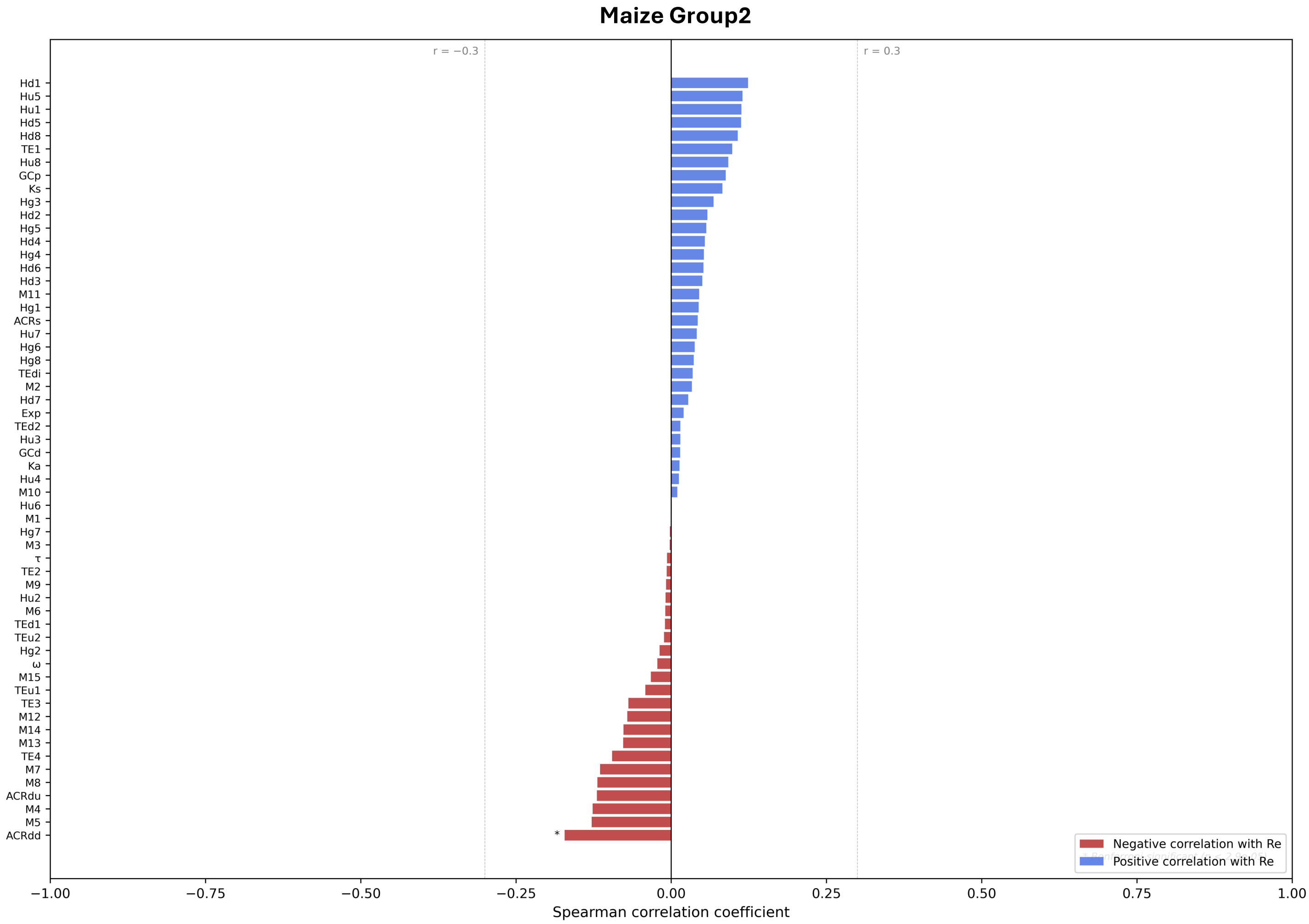


**Fig. S21. Spearman correlation of recombination rate (Re) with all maize group2 features.** Features are ranked by Spearman ρ with Re (most negative at bottom). Bar color indicates direction of correlation: negative (red) or positive (blue). Asterisks indicate feature pairs surviving Bonferroni correction across all $n(n-1)/2$ pairwise tests within the group ($\alpha = 0.05$). Dashed reference lines are drawn at $\rho = \pm 0.3$.

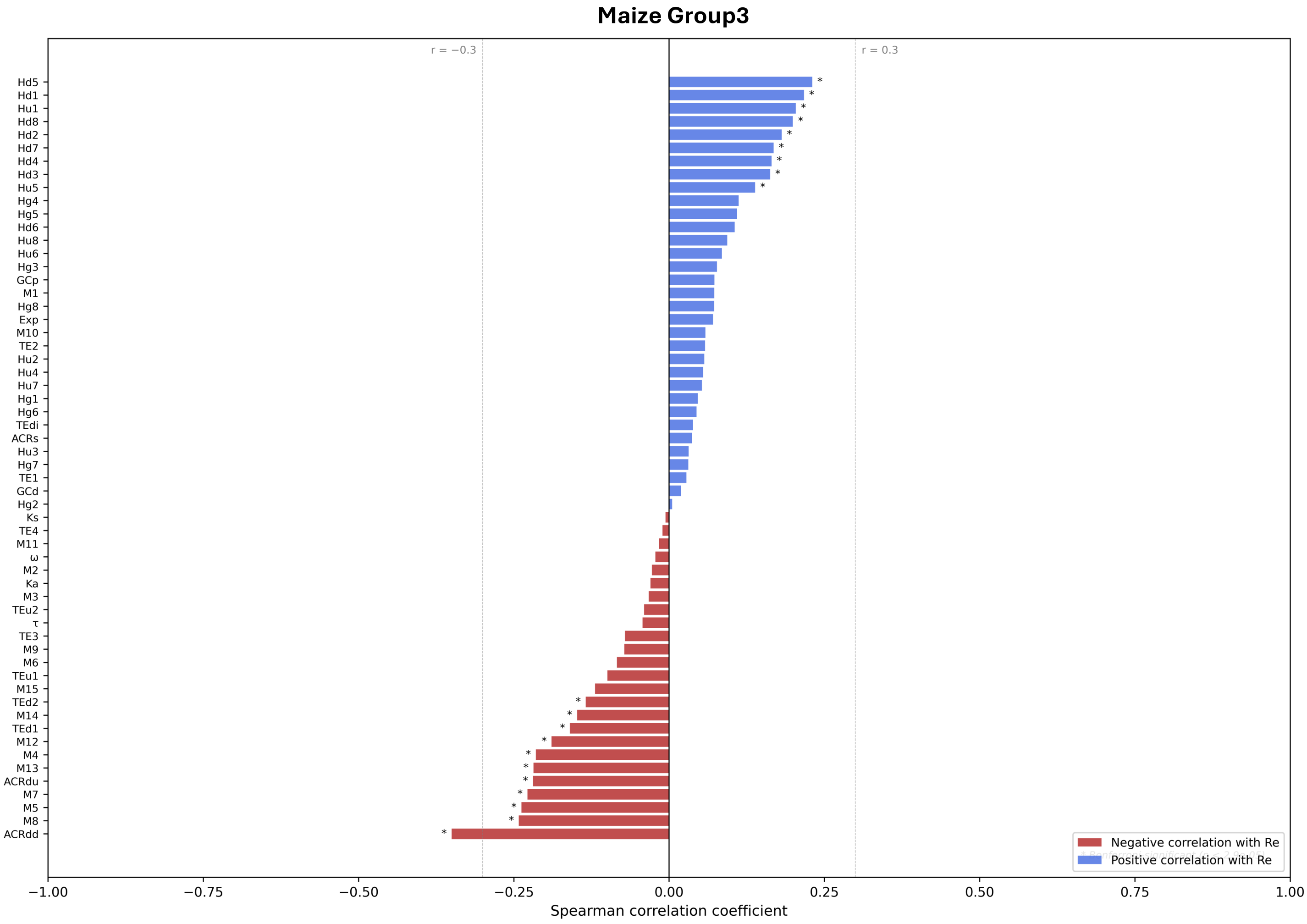


**Fig. S22. Spearman correlation of recombination rate (Re) with all maize group3 features.** Features are ranked by Spearman ρ with Re (most negative at bottom). Bar color indicates direction of correlation: negative (red) or positive (blue). Asterisks indicate feature pairs surviving Bonferroni correction across all n(n−1)/2 pairwise tests within the group (α = 0.05). Dashed reference lines are drawn at ρ = ±0.3.

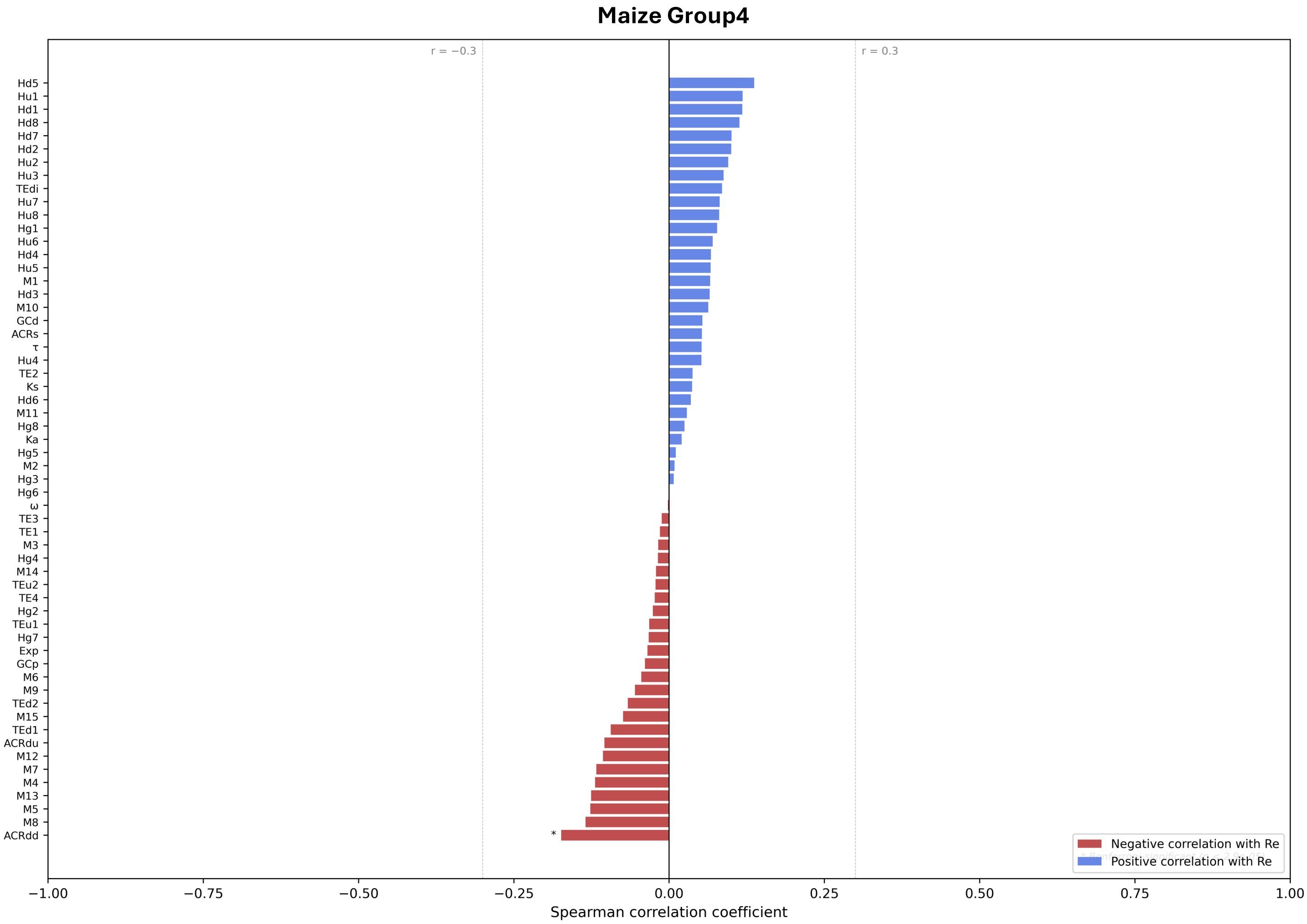


**Fig. S23. Spearman correlation of recombination rate (Re) with all maize group4 features.** Features are ranked by Spearman ρ with Re (most negative at bottom). Bar color indicates direction of correlation: negative (red) or positive (blue). Asterisks indicate feature pairs surviving Bonferroni correction across all n(n−1)/2 pairwise tests within the group (α = 0.05). Dashed reference lines are drawn at ρ = ±0.3.

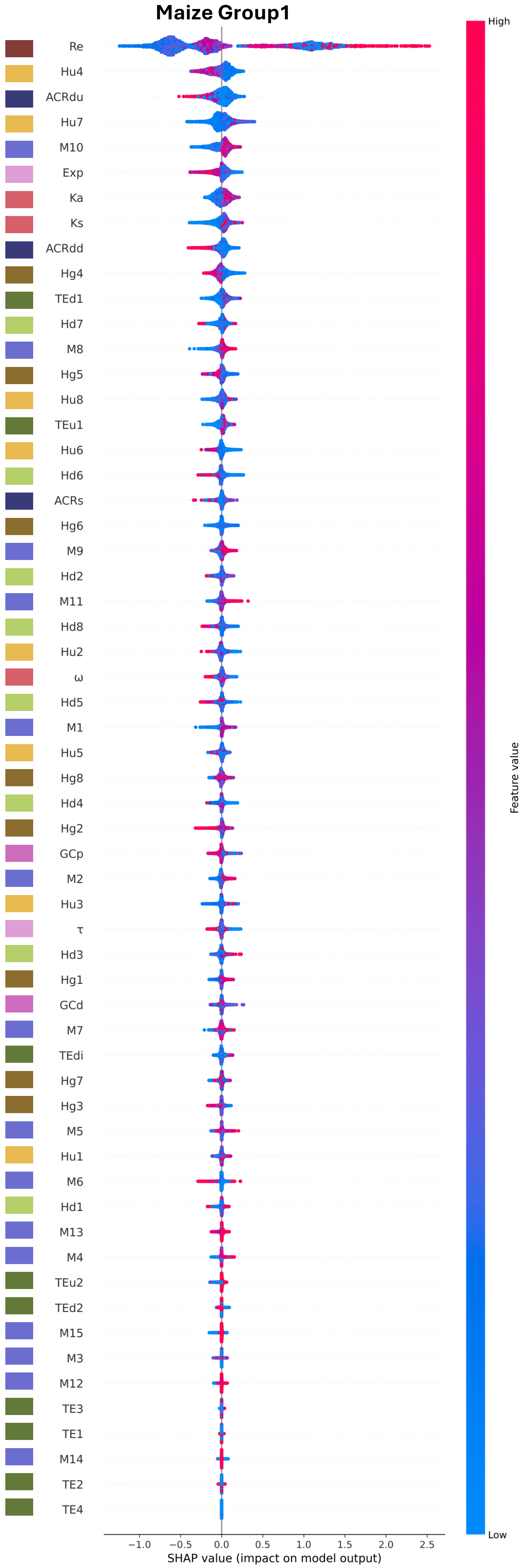


| Re | Recombination |
|---|---|

| | Expression Metrics |
|---|---|
| Exp | Avg expression |
| τ | Tau (tissue specificity index) |

| | GC Content |
|---|---|
| GCd | GC content cDNA |
| GCp | GC content promoter |

| | Transposable Elements (TEs) |
|---|---|
| TEu1 | Avg upstream density |
| TEd1 | Avg downstream density |
| TEu2 | Max upstream density |
| TEd2 | Max downstream density |
| TE1 | Helitrons |
| TE2 | Terminal inverted repeat elements |
| TE3 | LTR retrotransposons |
| TE4 | LINE and SINE elements |
| TEdi | TE distance |

| | Methylation |
|---|---|
| M1 | CHH upstream avg |
| M2 | CHH downstream avg |
| M3 | CHH gene body avg |
| M4 | CHG upstream avg |
| M5 | CHG downstream avg |
| M6 | CHG gene body avg |
| M7 | CG upstream avg |
| M8 | CG downstream avg |
| M9 | CG gene body avg |
| M10 | CHH upstream max |
| M11 | CHH downstream max |
| M12 | CHG upstream max |
| M13 | CHG downstream max |
| M14 | CG upstream max |
| M15 | CG downstream max |

| | Histone Mod. Upstream |
|---|---|
| Hu1 | H2A.Z up |
| Hu2 | H3K4me1 up |
| Hu3 | H3K4me3 up |
| Hu4 | H3K9ac up |
| Hu5 | H3K27ac up |
| Hu6 | H3K27me3 up |
| Hu7 | H3K36me3 up |
| Hu8 | H3K56ac up |

| | Histone Mod. Gene Body |
|---|---|
| Hg1 | H2A.Z gene body |
| Hg2 | H3K4me1 gene body |
| Hg3 | H3K4me3 gene body |
| Hg4 | H3K9ac gene body |
| Hg5 | H3K27ac gene body |
| Hg6 | H3K27me3 gene body |
| Hg7 | H3K36me3 gene body |
| Hg8 | H3K56ac gene body |

| | Histone Mod. Downstream |
|---|---|
| Hd1 | H2A.Z down |
| Hd2 | H3K4me1 down |
| Hd3 | H3K4me3 down |
| Hd4 | H3K9ac down |
| Hd5 | H3K27ac down |
| Hd6 | H3K27me3 down |
| Hd7 | H3K36me3 down |
| Hd8 | H3K56ac down |

| | ACRs |
|---|---|
| ACRdu | ACR distance to TSS |
| ACRdd | ACR distance to TTS |
| ACRs | Nearest summit |

| | Evolutionary Pressure |
|---|---|
| Ka | Nonsynonymous substitutions |
| Ks | Synonymous substitutions |
| ω | Omega, Ka/Ks ratio |

**Fig. S24. SHAP summary beeswarm plot for maize group1.** Features are ranked by mean absolute SHAP value (highest at top). Each point represents one test-fold sample, with horizontal position indicating the SHAP value (impact on model output); points are jittered vertically for visibility where they overlap. Point color encodes the raw feature value for that sample, ranging from low (blue) to high (red). Positive SHAP values indicate a contribution toward maize2 classification; negative SHAP values indicate a contribution toward maize1 classification. SHAP values were computed on held-out test-fold samples across 10 stratified cross-validation folds.

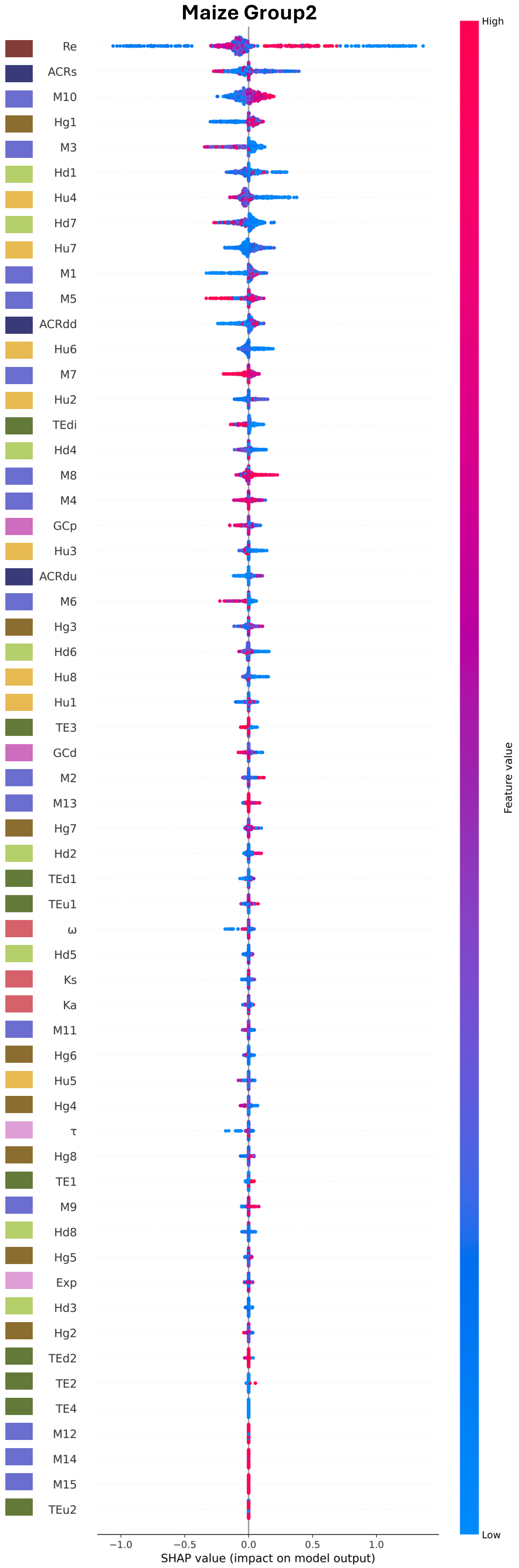


| Re | Recombination |
|---|---|

| | Expression Metrics |
|---|---|
| Exp | Avg expression |
| τ | Tau (tissue specificity index) |

| | GC Content |
|---|---|
| GCd | GC content cDNA |
| GCp | GC content promoter |

| | Transposable Elements (TEs) |
|---|---|
| TEu1 | Avg upstream density |
| TEd1 | Avg downstream density |
| TEu2 | Max upstream density |
| TEd2 | Max downstream density |
| TE1 | Helitrons |
| TE2 | Terminal inverted repeat elements |
| TE3 | LTR retrotransposons |
| TE4 | LINE and SINE elements |
| TEdi | TE distance |

| | Methylation |
|---|---|
| M1 | CHH upstream avg |
| M2 | CHH downstream avg |
| M3 | CHH gene body avg |
| M4 | CHG upstream avg |
| M5 | CHG downstream avg |
| M6 | CHG gene body avg |
| M7 | CG upstream avg |
| M8 | CG downstream avg |
| M9 | CG gene body avg |
| M10 | CHH upstream max |
| M11 | CHH downstream max |
| M12 | CHG upstream max |
| M13 | CHG downstream max |
| M14 | CG upstream max |
| M15 | CG downstream max |

| | Histone Mod. Upstream |
|---|---|
| Hu1 | H2A.Z up |
| Hu2 | H3K4me1 up |
| Hu3 | H3K4me3 up |
| Hu4 | H3K9ac up |
| Hu5 | H3K27ac up |
| Hu6 | H3K27me3 up |
| Hu7 | H3K36me3 up |
| Hu8 | H3K56ac up |

| | Histone Mod. Gene Body |
|---|---|
| Hg1 | H2A.Z gene body |
| Hg2 | H3K4me1 gene body |
| Hg3 | H3K4me3 gene body |
| Hg4 | H3K9ac gene body |
| Hg5 | H3K27ac gene body |
| Hg6 | H3K27me3 gene body |
| Hg7 | H3K36me3 gene body |
| Hg8 | H3K56ac gene body |

| | Histone Mod. Downstream |
|---|---|
| Hd1 | H2A.Z down |
| Hd2 | H3K4me1 down |
| Hd3 | H3K4me3 down |
| Hd4 | H3K9ac down |
| Hd5 | H3K27ac down |
| Hd6 | H3K27me3 down |
| Hd7 | H3K36me3 down |
| Hd8 | H3K56ac down |

| | ACRs |
|---|---|
| ACRdu | ACR distance to TSS |
| ACRdd | ACR distance to TTS |
| ACRs | Nearest summit |

| | Evolutionary Pressure |
|---|---|
| Ka | Nonsynonymous substitutions |
| Ks | Synonymous substitutions |
| ω | Omega, Ka/Ks ratio |

**Fig. S25. SHAP summary beeswarm plot for maize group2.** Features are ranked by mean absolute SHAP value (highest at top). Each point represents one test-fold sample, with horizontal position indicating the SHAP value (impact on model output); points are jittered vertically for visibility where they overlap. Point color encodes the raw feature value for that sample, ranging from low (blue) to high (red). Positive SHAP values indicate a contribution toward maize2 classification; negative SHAP values indicate a contribution toward maize1 classification. SHAP values were computed on held-out test-fold samples across 10 stratified cross-validation folds.

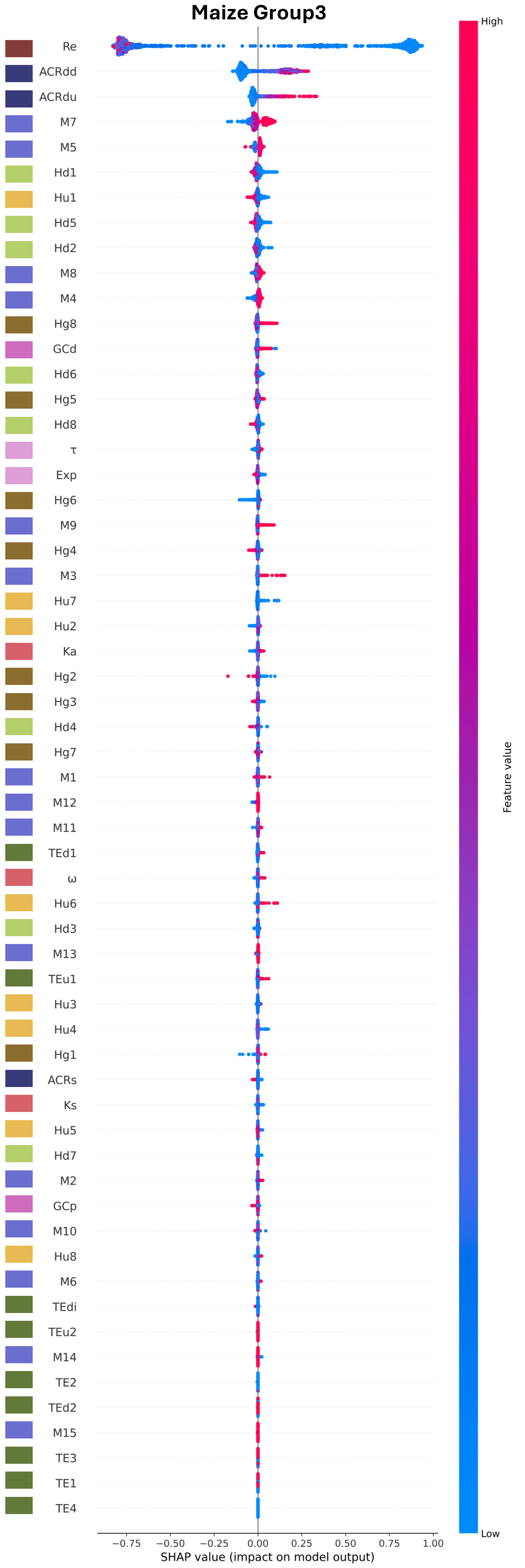


| Re | Recombination |
|---|---|

| | Expression Metrics |
|---|---|
| Exp | Avg expression |
| τ | Tau (tissue specificity index) |

| | GC Content |
|---|---|
| GCd | GC content cDNA |
| GCp | GC content promoter |

| | Transposable Elements (TEs) |
|---|---|
| TEu1 | Avg upstream density |
| TEd1 | Avg downstream density |
| TEu2 | Max upstream density |
| TEd2 | Max downstream density |
| TE1 | Helitrons |
| TE2 | Terminal inverted repeat elements |
| TE3 | LTR retrotransposons |
| TE4 | LINE and SINE elements |
| TEdi | TE distance |

| | Methylation |
|---|---|
| M1 | CHH upstream avg |
| M2 | CHH downstream avg |
| M3 | CHH gene body avg |
| M4 | CHG upstream avg |
| M5 | CHG downstream avg |
| M6 | CHG gene body avg |
| M7 | CG upstream avg |
| M8 | CG downstream avg |
| M9 | CG gene body avg |
| M10 | CHH upstream max |
| M11 | CHH downstream max |
| M12 | CHG upstream max |
| M13 | CHG downstream max |
| M14 | CG upstream max |
| M15 | CG downstream max |

| | Histone Mod. Upstream |
|---|---|
| Hu1 | H2A.Z up |
| Hu2 | H3K4me1 up |
| Hu3 | H3K4me3 up |
| Hu4 | H3K9ac up |
| Hu5 | H3K27ac up |
| Hu6 | H3K27me3 up |
| Hu7 | H3K36me3 up |
| Hu8 | H3K56ac up |

| | Histone Mod. Gene Body |
|---|---|
| Hg1 | H2A.Z gene body |
| Hg2 | H3K4me1 gene body |
| Hg3 | H3K4me3 gene body |
| Hg4 | H3K9ac gene body |
| Hg5 | H3K27ac gene body |
| Hg6 | H3K27me3 gene body |
| Hg7 | H3K36me3 gene body |
| Hg8 | H3K56ac gene body |

| | Histone Mod. Downstream |
|---|---|
| Hd1 | H2A.Z down |
| Hd2 | H3K4me1 down |
| Hd3 | H3K4me3 down |
| Hd4 | H3K9ac down |
| Hd5 | H3K27ac down |
| Hd6 | H3K27me3 down |
| Hd7 | H3K36me3 down |
| Hd8 | H3K56ac down |

| | ACRs |
|---|---|
| ACRdu | ACR distance to TSS |
| ACRdd | ACR distance to TTS |
| ACRs | Nearest summit |

| | Evolutionary Pressure |
|---|---|
| Ka | Nonsynonymous substitutions |
| Ks | Synonymous substitutions |
| ω | Omega, Ka/Ks ratio |

**Fig. S26. SHAP summary beeswarm plot for maize group3.** Features are ranked by mean absolute SHAP value (highest at top). Each point represents one test-fold sample, with horizontal position indicating the SHAP value (impact on model output); points are jittered vertically for visibility where they overlap. Point color encodes the raw feature value for that sample, ranging from low (blue) to high (red). Positive SHAP values indicate a contribution toward maize2 classification; negative SHAP values indicate a contribution toward maize1 classification. SHAP values were computed on held-out test-fold samples across 10 stratified cross-validation folds.

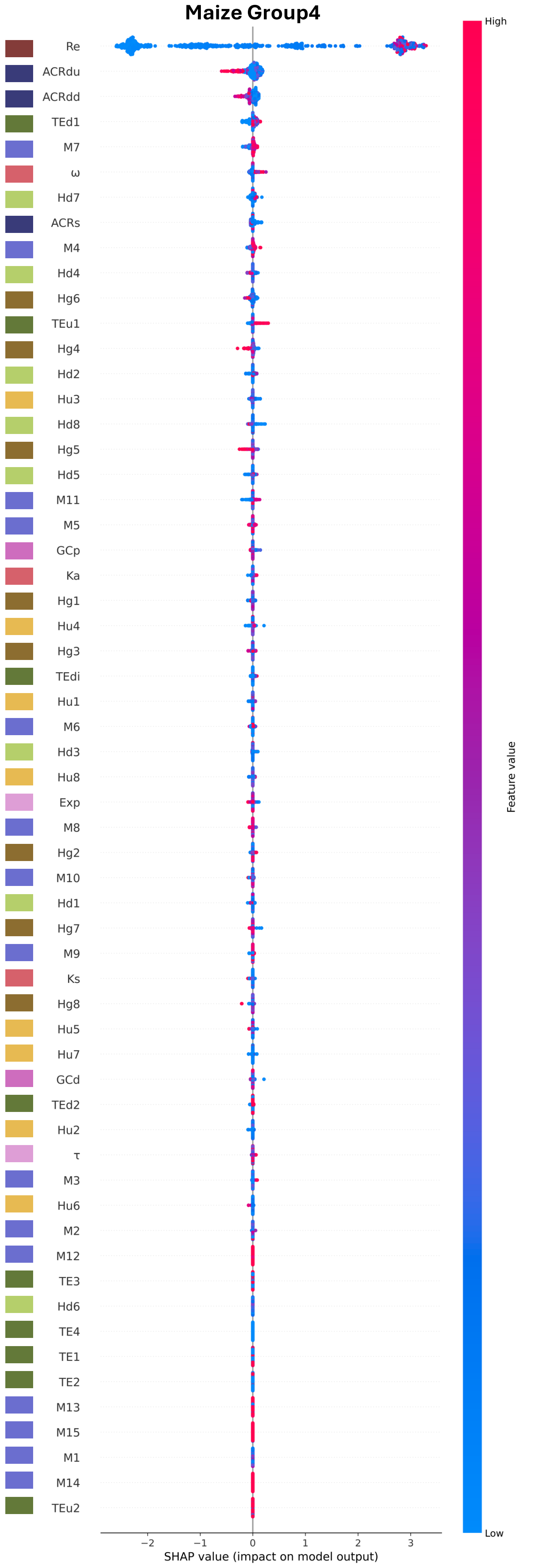


| Re | Recombination |
|---|---|

| | Expression Metrics |
|---|---|
| Exp | Avg expression |
| τ | Tau (tissue specificity index) |

| | GC Content |
|---|---|
| GCd | GC content cDNA |
| GCp | GC content promoter |

| | Transposable Elements (TEs) |
|---|---|
| TEu1 | Avg upstream density |
| TEd1 | Avg downstream density |
| TEu2 | Max upstream density |
| TEd2 | Max downstream density |
| TE1 | Helitrons |
| TE2 | Terminal inverted repeat elements |
| TE3 | LTR retrotransposons |
| TE4 | LINE and SINE elements |
| TEdi | TE distance |

| | Methylation |
|---|---|
| M1 | CHH upstream avg |
| M2 | CHH downstream avg |
| M3 | CHH gene body avg |
| M4 | CHG upstream avg |
| M5 | CHG downstream avg |
| M6 | CHG gene body avg |
| M7 | CG upstream avg |
| M8 | CG downstream avg |
| M9 | CG gene body avg |
| M10 | CHH upstream max |
| M11 | CHH downstream max |
| M12 | CHG upstream max |
| M13 | CHG downstream max |
| M14 | CG upstream max |
| M15 | CG downstream max |

| | Histone Mod. Upstream |
|---|---|
| Hu1 | H2A.Z up |
| Hu2 | H3K4me1 up |
| Hu3 | H3K4me3 up |
| Hu4 | H3K9ac up |
| Hu5 | H3K27ac up |
| Hu6 | H3K27me3 up |
| Hu7 | H3K36me3 up |
| Hu8 | H3K56ac up |

| | Histone Mod. Gene Body |
|---|---|
| Hg1 | H2A.Z gene body |
| Hg2 | H3K4me1 gene body |
| Hg3 | H3K4me3 gene body |
| Hg4 | H3K9ac gene body |
| Hg5 | H3K27ac gene body |
| Hg6 | H3K27me3 gene body |
| Hg7 | H3K36me3 gene body |
| Hg8 | H3K56ac gene body |

| | Histone Mod. Downstream |
|---|---|
| Hd1 | H2A.Z down |
| Hd2 | H3K4me1 down |
| Hd3 | H3K4me3 down |
| Hd4 | H3K9ac down |
| Hd5 | H3K27ac down |
| Hd6 | H3K27me3 down |
| Hd7 | H3K36me3 down |
| Hd8 | H3K56ac down |

| | ACRs |
|---|---|
| ACRdu | ACR distance to TSS |
| ACRdd | ACR distance to TTS |
| ACRs | Nearest summit |

| | Evolutionary Pressure |
|---|---|
| Ka | Nonsynonymous substitutions |
| Ks | Synonymous substitutions |
| ω | Omega, Ka/Ks ratio |

**Fig. S27. SHAP summary beeswarm plot for maize group4.** Features are ranked by mean absolute SHAP value (highest at top). Each point represents one test-fold sample, with horizontal position indicating the SHAP value (impact on model output); points are jittered vertically for visibility where they overlap. Point color encodes the raw feature value for that sample, ranging from low (blue) to high (red). Positive SHAP values indicate a contribution toward maize2 classification; negative SHAP values indicate a contribution toward maize1 classification. SHAP values were computed on held-out test-fold samples across 10 stratified cross-validation folds.

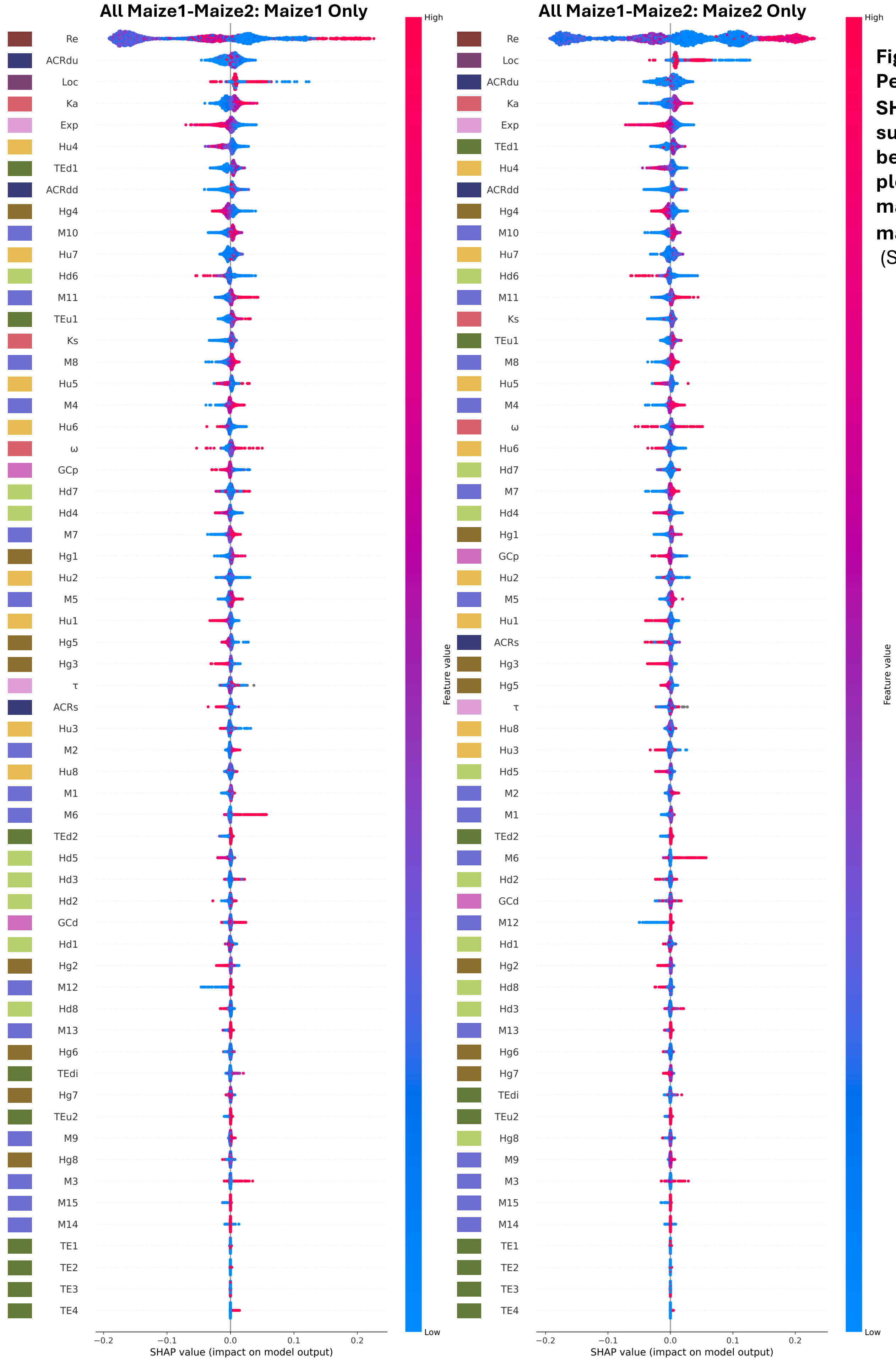


**Fig. S28. Per-label SHAP summary beeswarm plots for maize1-maize2.** (See pg 33)

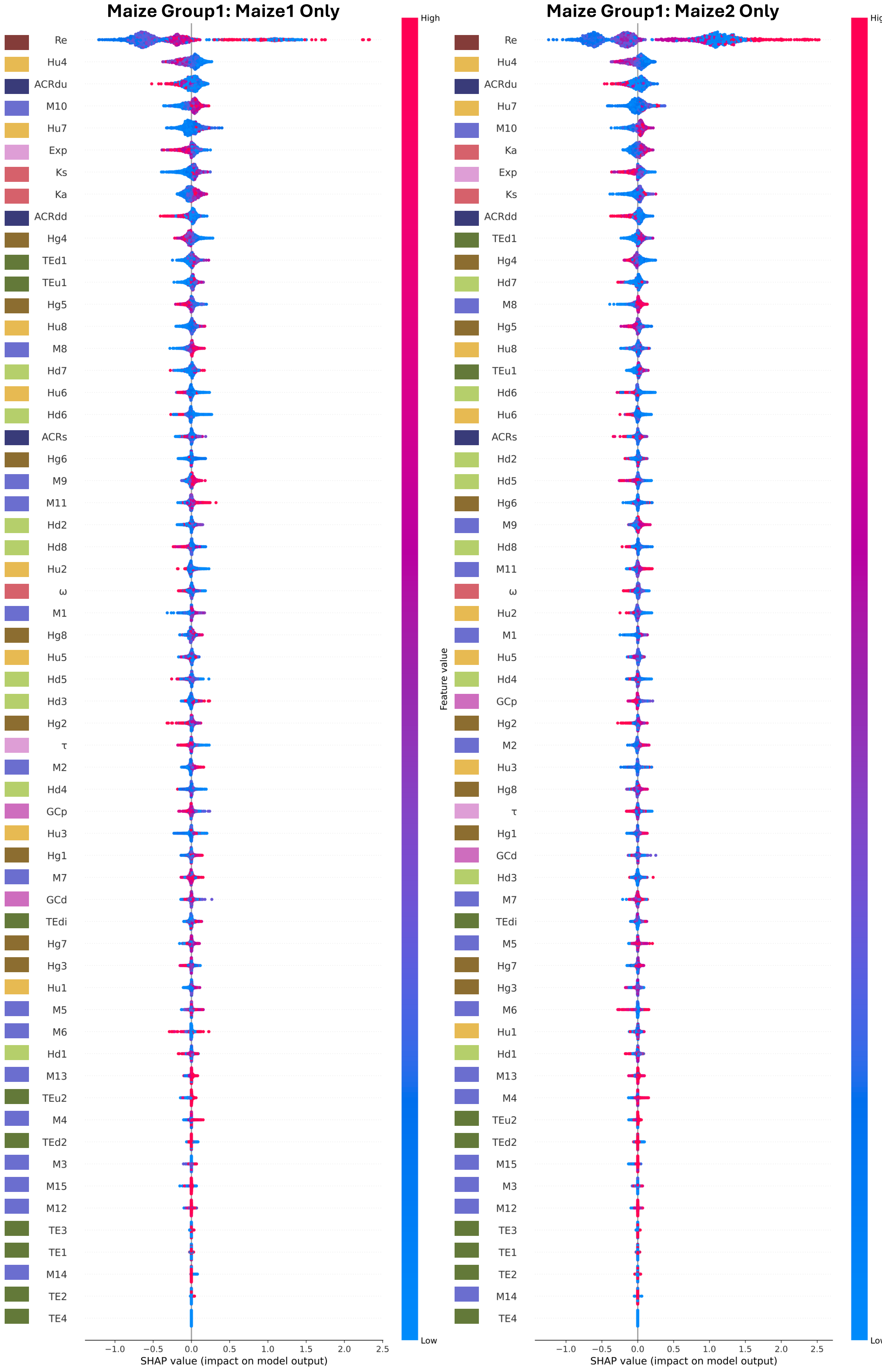


**Fig. S29. Per-label SHAP summary beeswarm plots for maize group1.** (See pg 33)

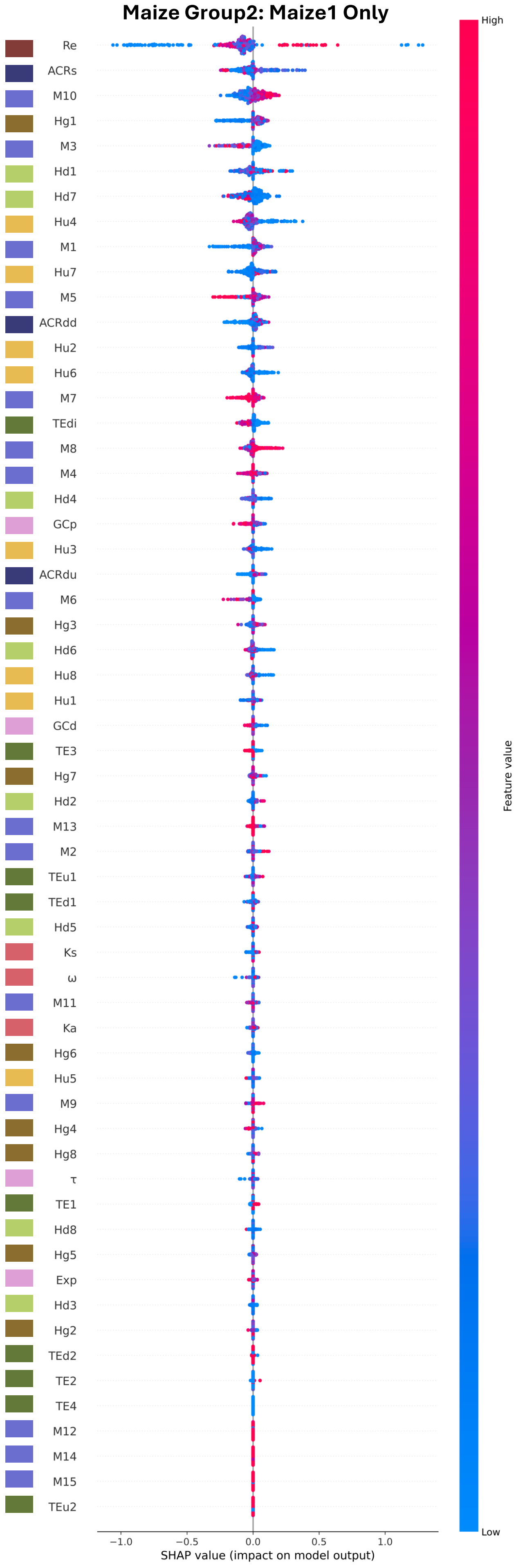


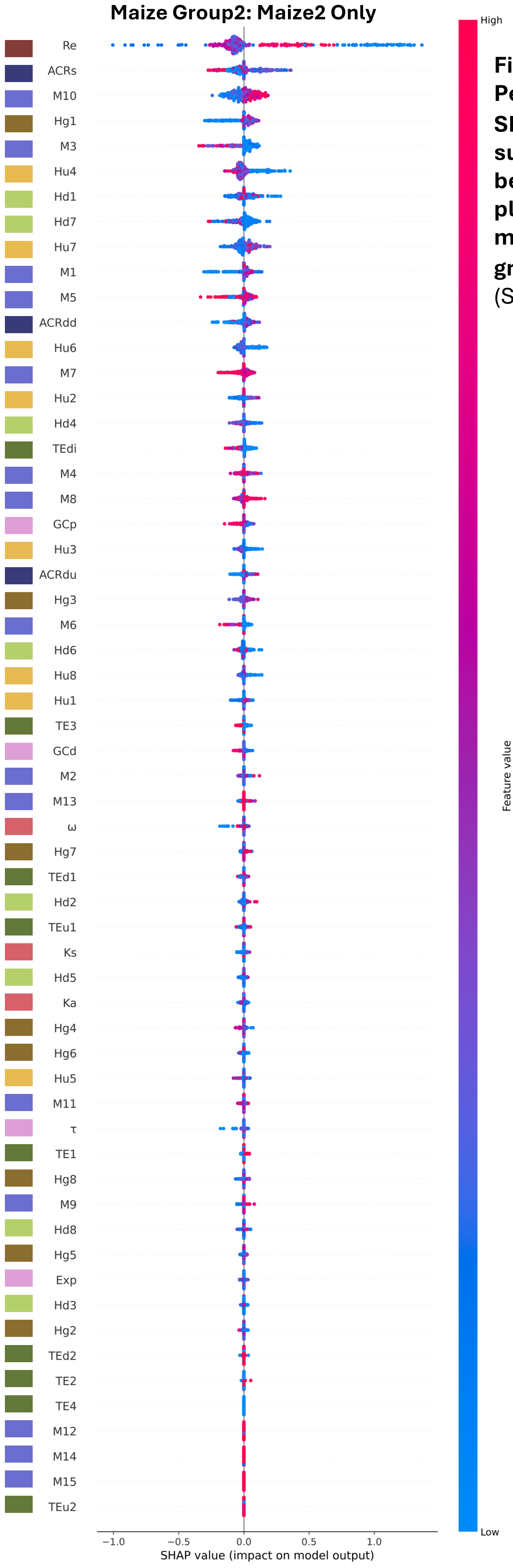


**Fig. S30. Per-label SHAP summary beeswarm plots for maize group2.** (See pg 33)

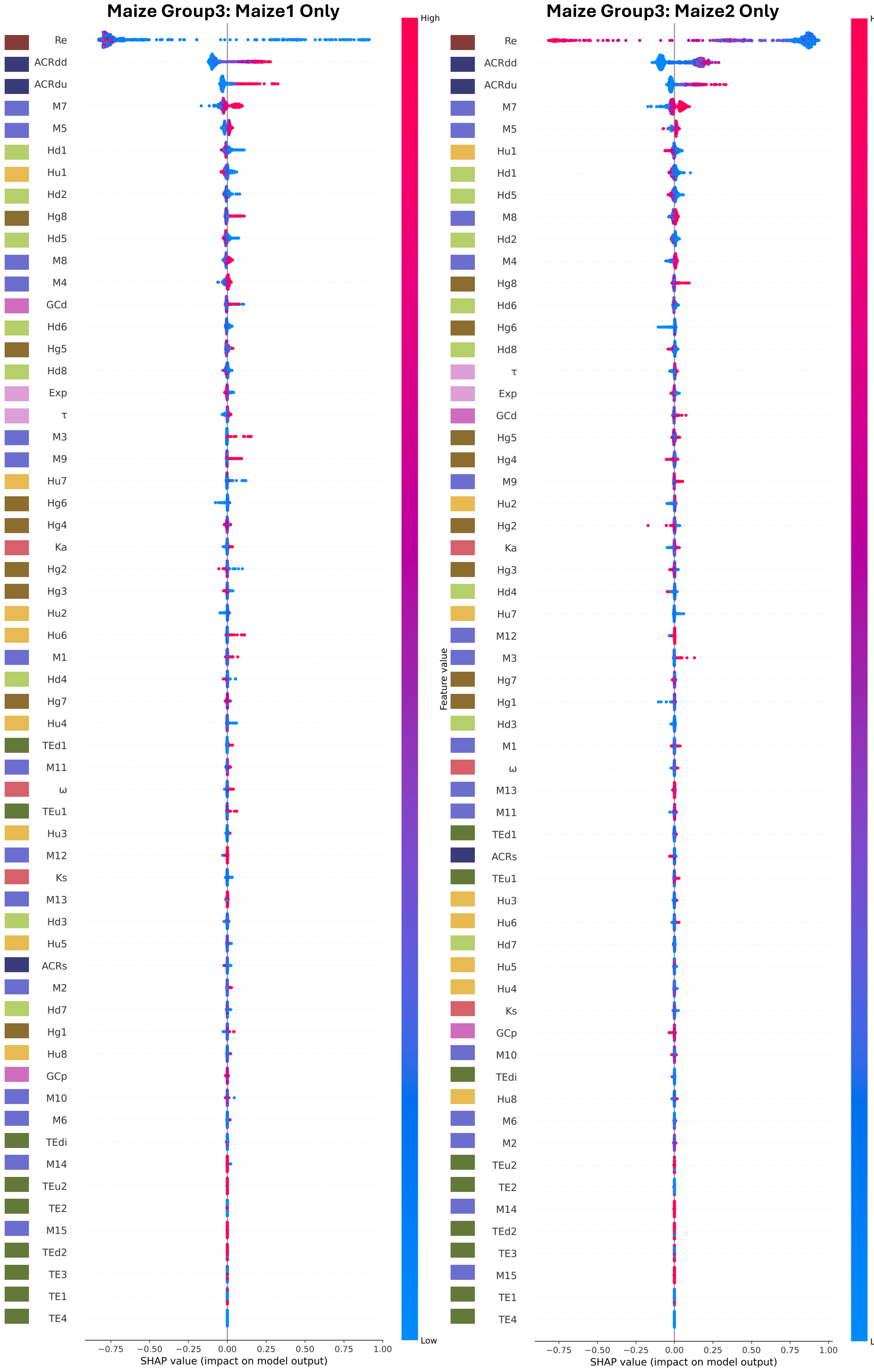


**Fig. S31. Per-label SHAP summary beeswarm plots for maize group3.** (See pg 33)

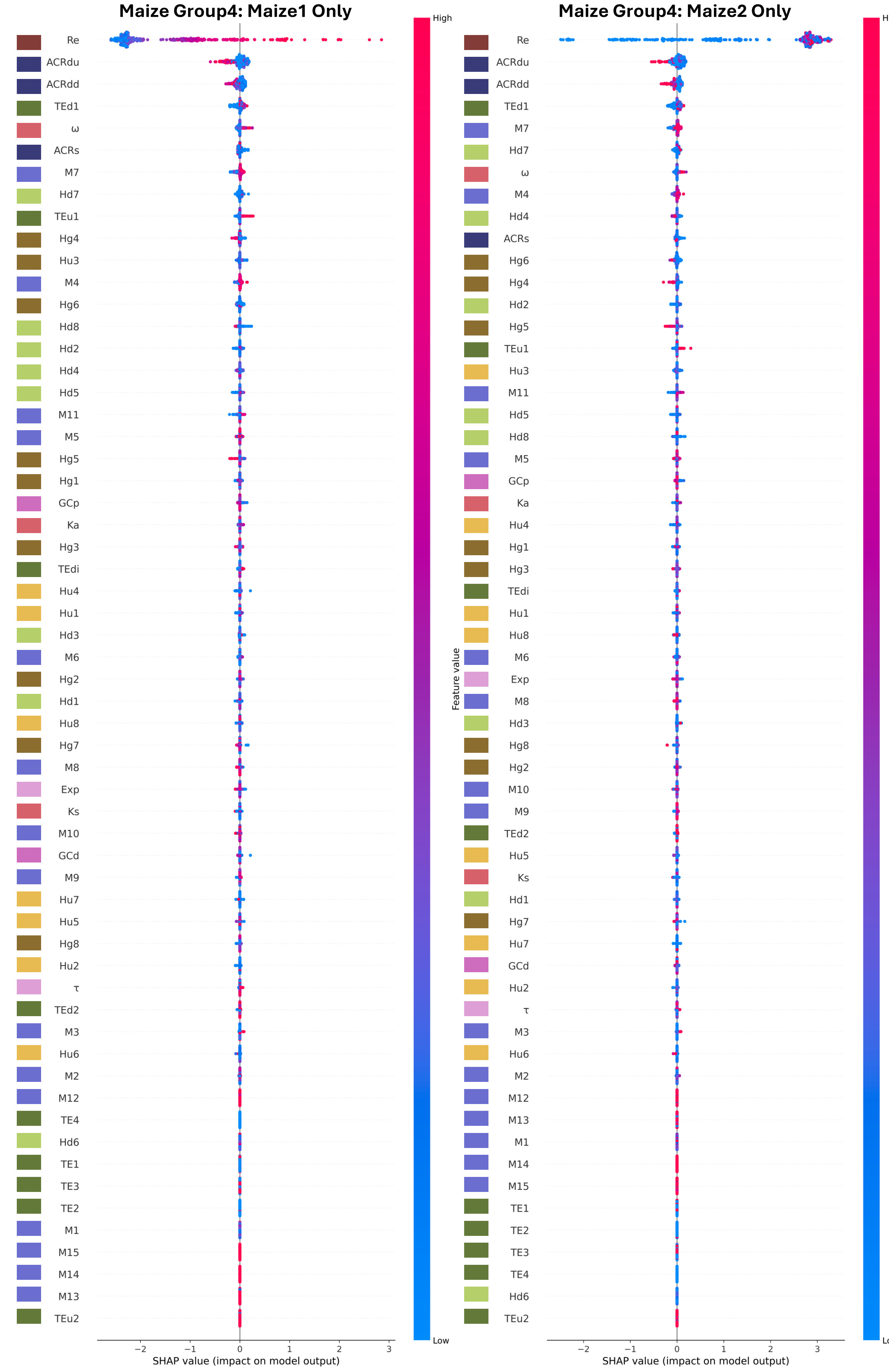


**Fig. S32. Per-label SHAP summary beeswarm plots for maize group4.** (See pg 33)

**Per-label SHAP summary beeswarm plots for maize.** SHAP values are shown separately for maize1 (dominant subgenome; left) and maize2 (recessive subgenome; right) samples. Within each panel, features are ranked independently by mean absolute SHAP value for that label's samples (highest at top); rank order therefore differs between panels and reflects asymmetry in the features driving dominant versus submissive subgenome classification. Each point represents one test-fold sample, with horizontal position indicating the SHAP value (impact on model output) and vertical jitter applied where points overlap. Point color encodes the raw feature value for that sample, ranging from low (blue) to high (red); gray points indicate missing (NaN) feature values. Positive SHAP values indicate a contribution toward maize2 classification; negative SHAP values indicate a contribution toward maize1 classification, and this directionality is consistent across both panels. SHAP values were computed on held-out test-fold samples across 10 stratified cross-validation folds. For SHAP values computed across all samples regardless of label, see Figure S24-S27. For summed directional SHAP contributions per feature, see Figure S14-S18.

**Fig. S28. Per-label SHAP summary beeswarm plots for maize1-maize2.**
**Fig. S29. Per-label SHAP summary beeswarm plots for maize group1.**
**Fig. S30. Per-label SHAP summary beeswarm plots for maize group2.**
**Fig. S31. Per-label SHAP summary beeswarm plots for maize group3.**
**Fig. S32. Per-label SHAP summary beeswarm plots for maize group4.**

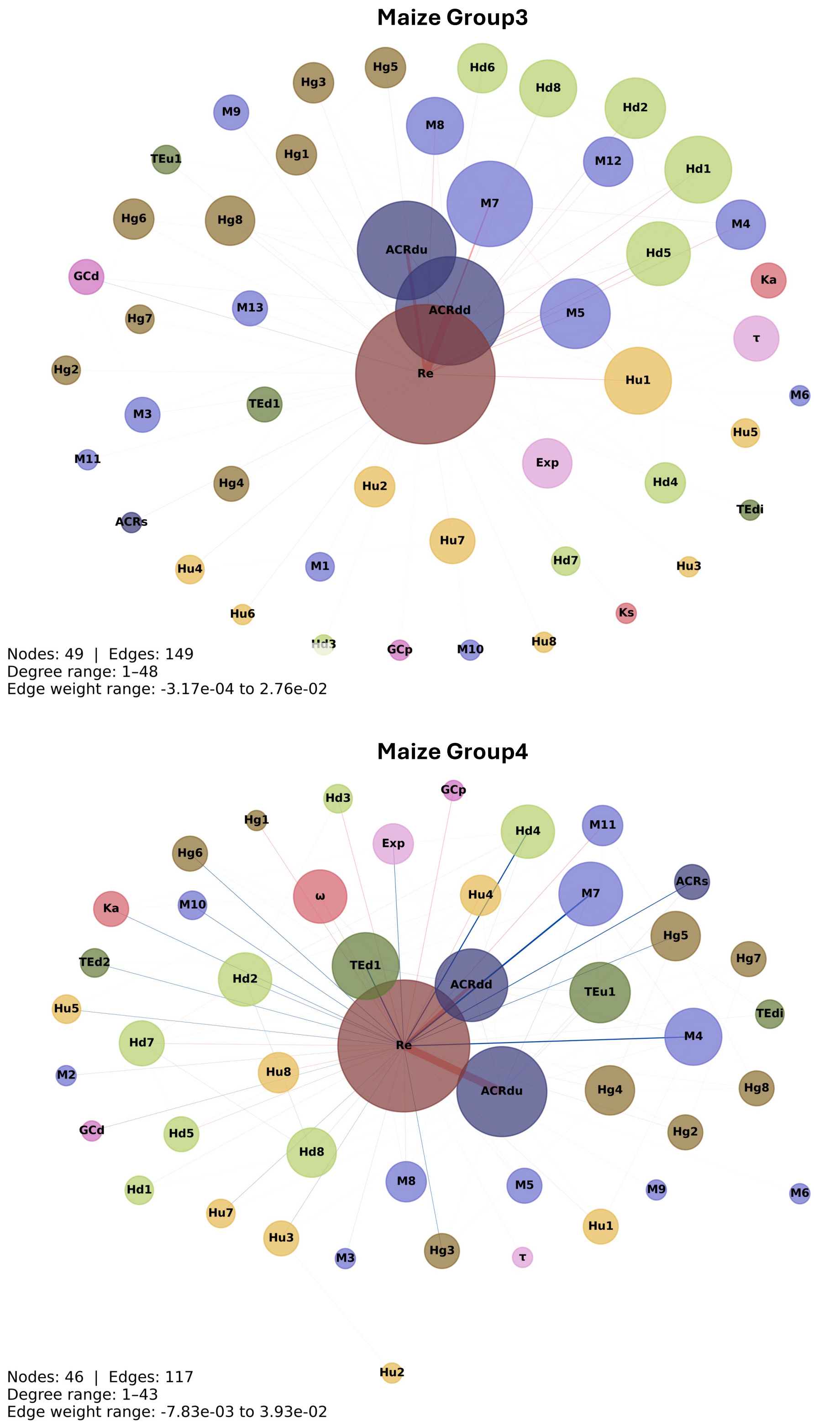


**Fig. S33. SHAP co-variation networks for maize subgenome classification.** Networks for maize group3 (top) and group4 (bottom). Nodes represent features, colored by feature group, and sized by degree (number of significant co-varying partners). Edges connect feature pairs with significant pairwise SHAP co-variation (permutation test, n = 1,000, α = 0.05), with color indicating direction: positive co-variation (features vary together in model importance; red) or negative co-variation (features vary in opposition; blue), and width scaled to reflect relative co-variation magnitude within each panel independently. Each panel reports the total node and edge count, degree range (minimum to maximum node degree in that network), and edge weight range (minimum to maximum co-variation score in that network).

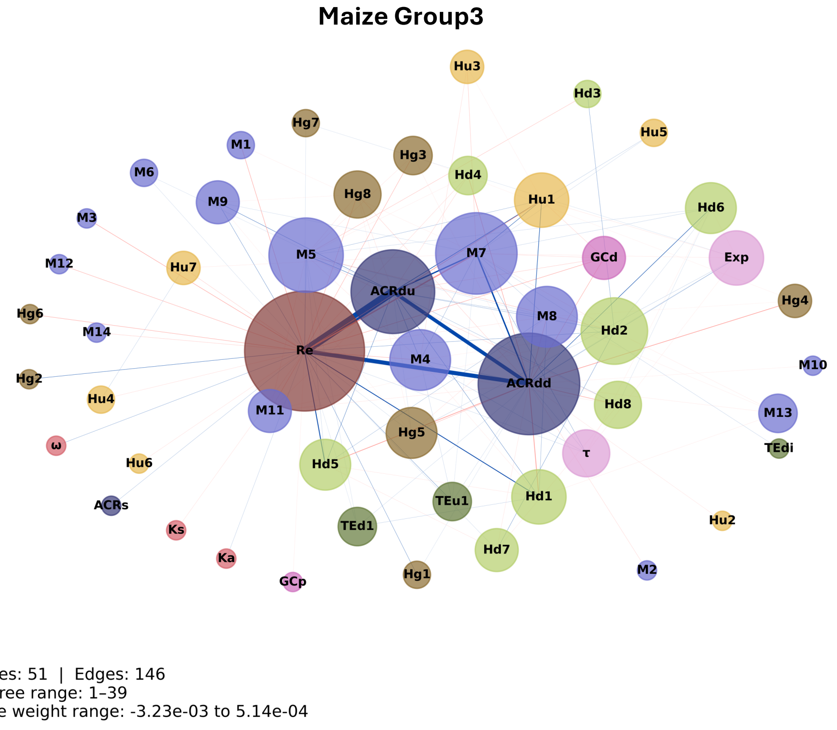


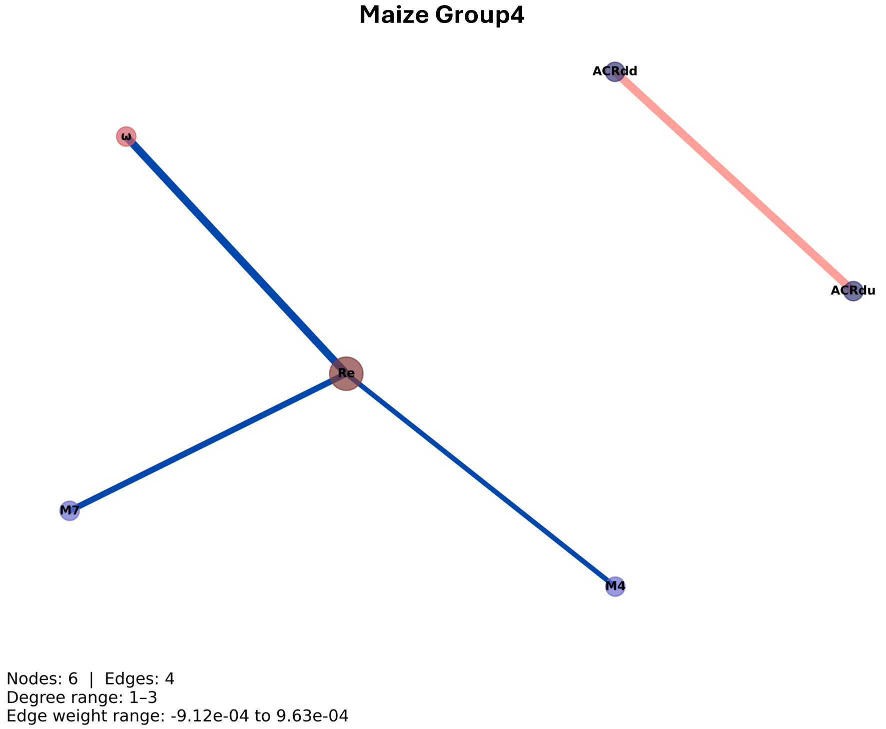


**Fig. S34. SHAP feature interaction networks for maize.** Interaction networks shown for maize group3 (**A**) and group4 (**B**). SHAP interaction values quantify the degree to which pairs of features jointly influence model predictions beyond what is expected from their individual contributions alone. For each feature pair, mean Shapley interaction values were computed across all test-set samples. Edges connect pairs whose absolute mean interaction value exceeded a robust effect size threshold (median + 12 × median absolute deviation, computed from non-zero pairs within each group) and are colored by interaction direction: positive (pink) edges indicate synergistic interactions, in which the combined contribution of two features exceeds the sum of their individual effects, and negative (blue) edges indicate antagonistic interactions, in which the combined contribution is less than the sum. Node size is proportional to degree (number of retained edges) and node color indicates feature category, as in Fig. 4.

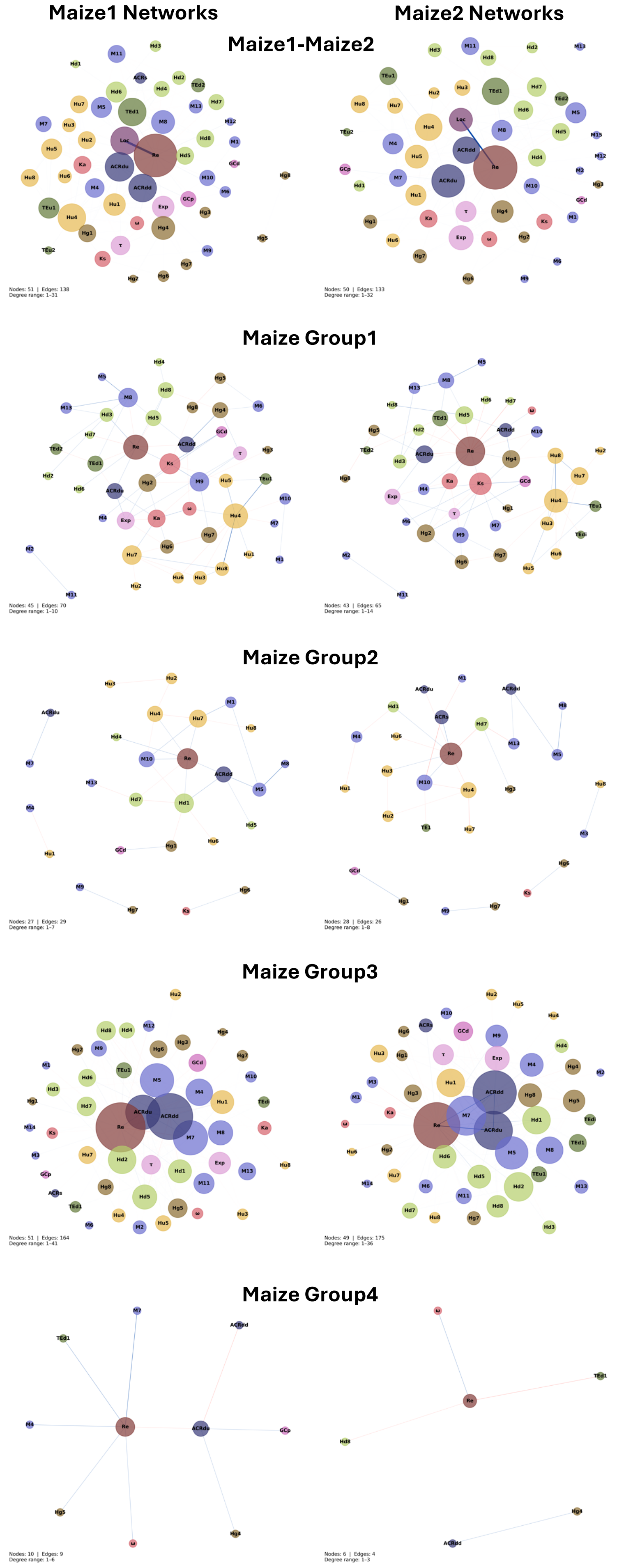


**Fig. S35. SHAP feature interaction networks for maize per label.** Networks show pairwise Shapley interaction values computed separately for genes of the dominant subgenome (maize1, left column) and the non-dominant subgenome (maize2, right column). Edges connect feature pairs whose absolute mean Shapley interaction value exceeded the group-specific robust effect size threshold (median + 12 × median absolute deviation of non-zero pairs), computed from all-samples interactions and applied uniformly to both per-label subsets within each group to ensure directly comparable filtering across columns. Positive (pink) edges indicate synergistic interactions and negative (blue) edges indicate antagonistic interactions. Node size is proportional to degree and node color indicates feature category, as in Fig. 4.

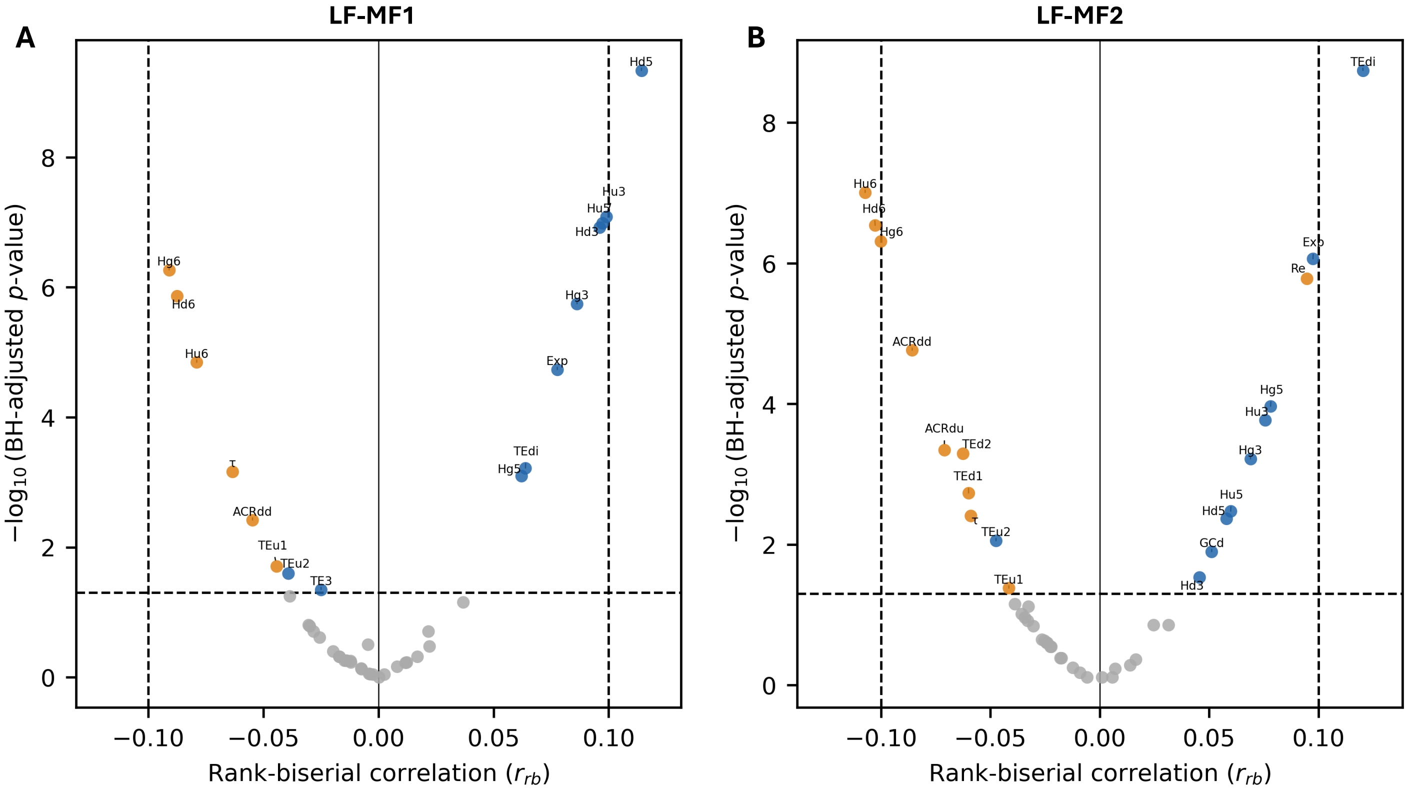

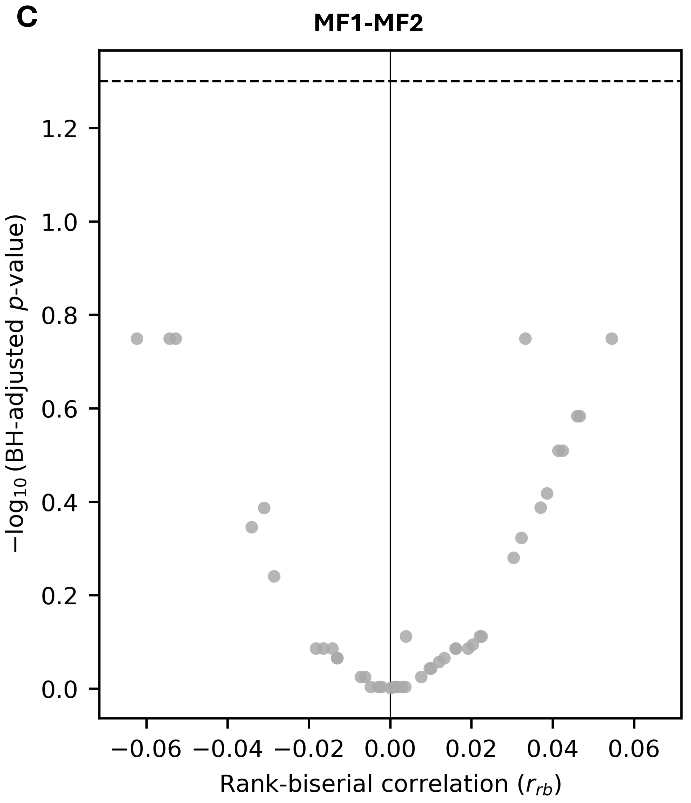


**Fig. S36. Volcano plots of *B. rapa* feature distributions.** Volcano plots showing the results of two-sided Mann-Whitney U tests comparing genomic feature distributions between subgenome pairs in *Brassica rapa* for LF-MF1 (**A**), LF-MF2 (**B**), and MF1-MF2 (**C**). The x-axis shows the rank-biserial correlation (*r*~rb~), a measure of effect size ranging from −1 to +1. The y-axis shows the −log~10~ Benjamini-Hochberg (BH) FDR-adjusted *p*-value. In each comparison, blue points indicate features with significantly higher values in the more dominant subgenome (LF in LF-MF1 and LF-MF2 comparisons; MF1 in the MF1-MF2 comparison); orange points indicate features with significantly higher values in the less dominant subgenome. Grey points are not significant. The horizontal dashed line marks the FDR significance threshold (BH-adjusted *P* = 0.05). Vertical dashed lines mark an effect size threshold of |*r*~rb~| = 0.1. No features reached significance in the MF1-MF2 comparison after FDR correction.

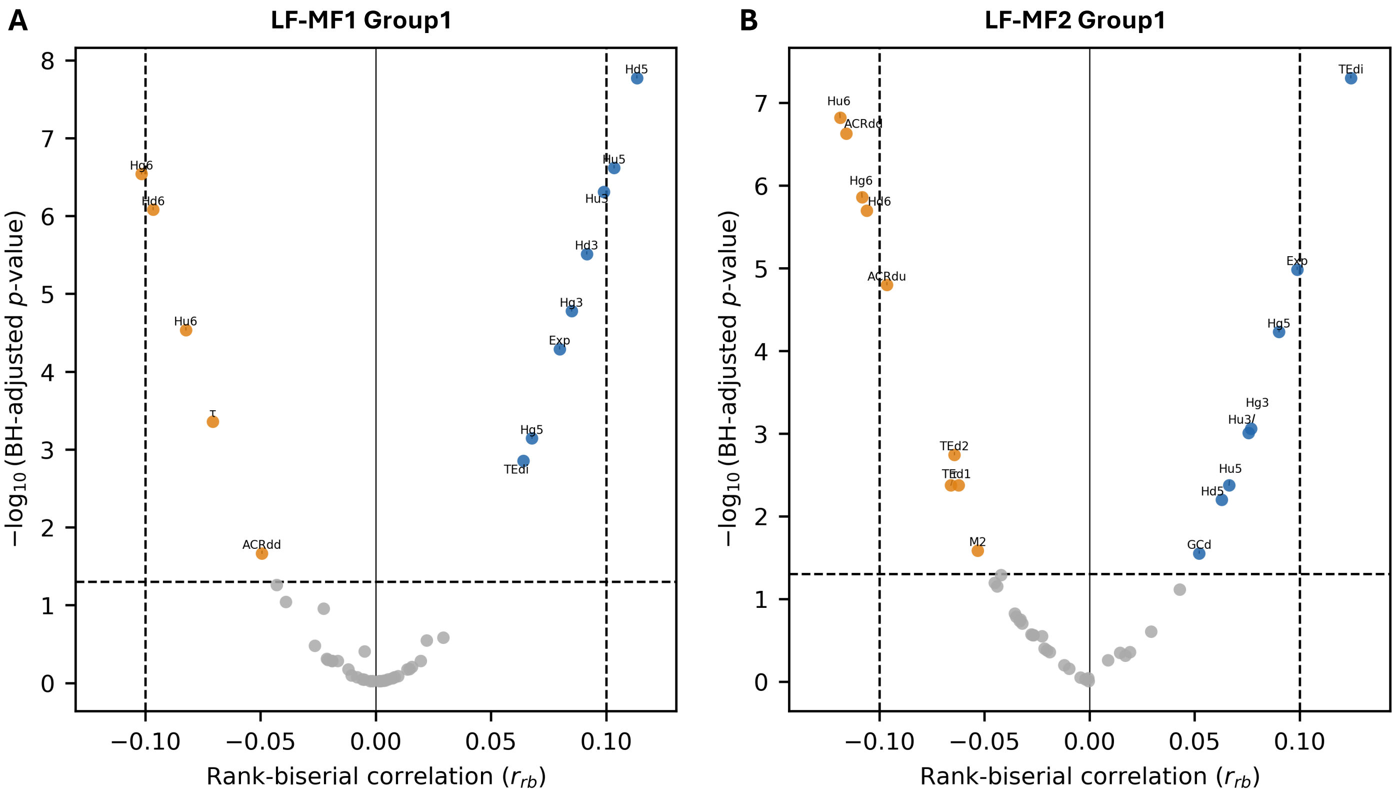


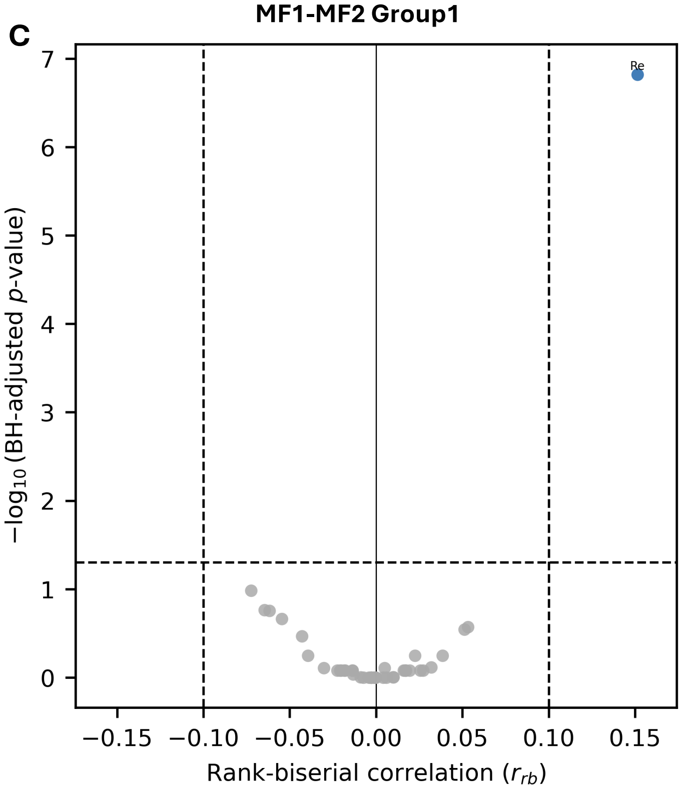


**Fig. S37. *B. rapa* group1 gene pairs and volcano plots of feature distributions.** Volcano plots showing the results of two-sided Mann-Whitney U tests comparing genomic feature distributions between subgenome pairs in *Brassica rapa* for LF-MF1 Group1 (**A**), LF-MF2 Group1 (**B**), and MF1-MF2 Group1 (**C**). The x-axis shows the rank-biserial correlation (*r*~rb~), a measure of effect size ranging from −1 to +1. The y-axis shows the −log~10~ Benjamini-Hochberg (BH) FDR-adjusted *p*-value. In each comparison, blue points indicate features with significantly higher values in the more dominant subgenome (LF in LF-MF1 and LF-MF2 comparisons; MF1 in the MF1-MF2 comparison); orange points indicate features with significantly higher values in the less dominant subgenome. Grey points are not significant. The horizontal dashed line marks the FDR significance threshold (BH-adjusted *P* = 0.05). Vertical dashed lines mark an effect size threshold of |*r*~rb~| = 0.1.

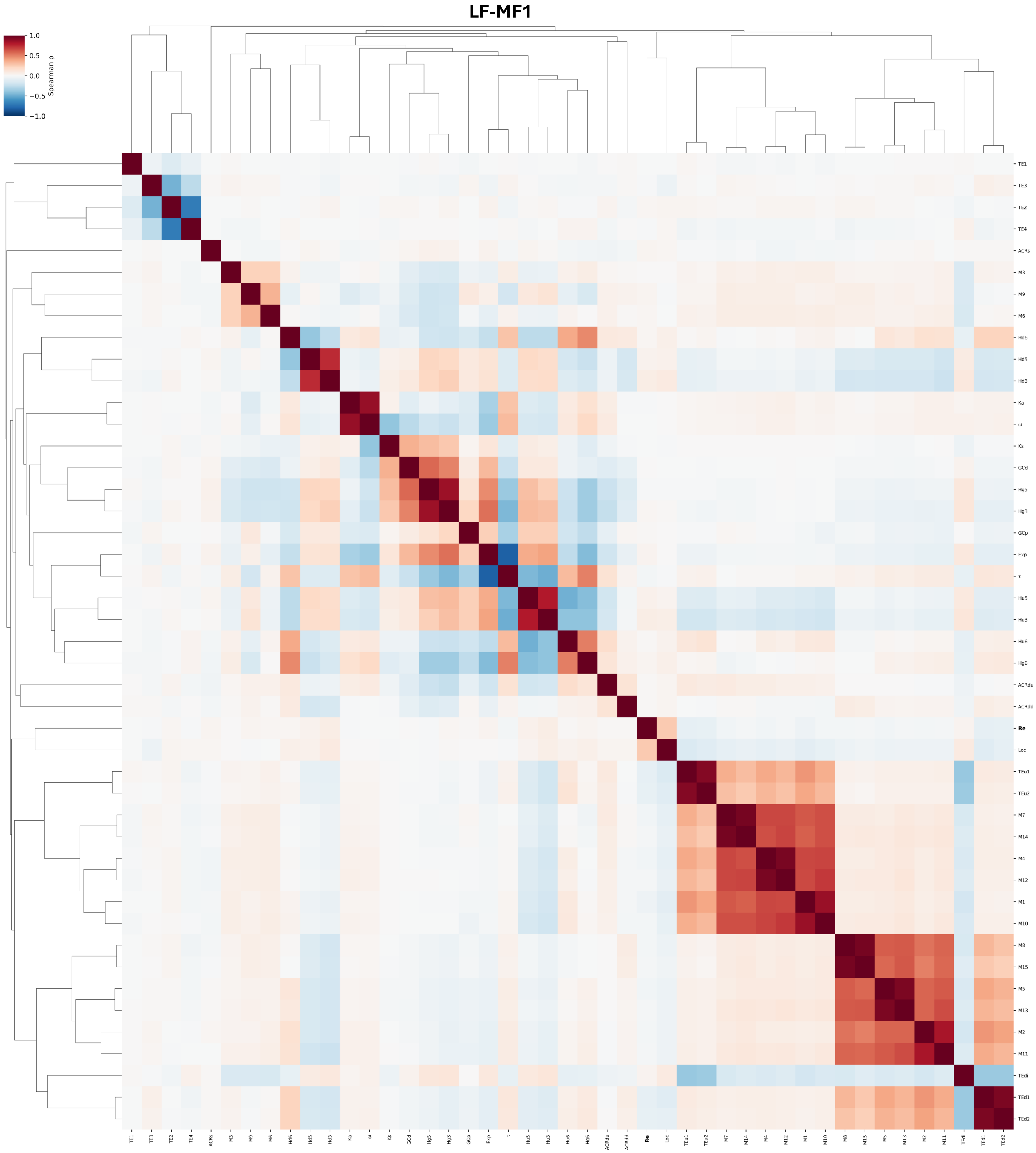


**Fig. S38. Spearman correlation heatmap of *B. rapa* LF-MF1 genomic and epigenomic features.** Pairwise Spearman rank correlation coefficients were computed across all features using complete pairwise deletion for missing values. Features are clustered on both axes using average linkage on a distance matrix defined as 1 − |ρ|, where ρ is the Spearman correlation coefficient; this metric groups co-varying feature blocks symmetrically regardless of the direction of correlation. The color scale represents the Spearman ρ value (red = positive correlation, blue = negative correlation). Recombination rate (Re) is shown in bold.

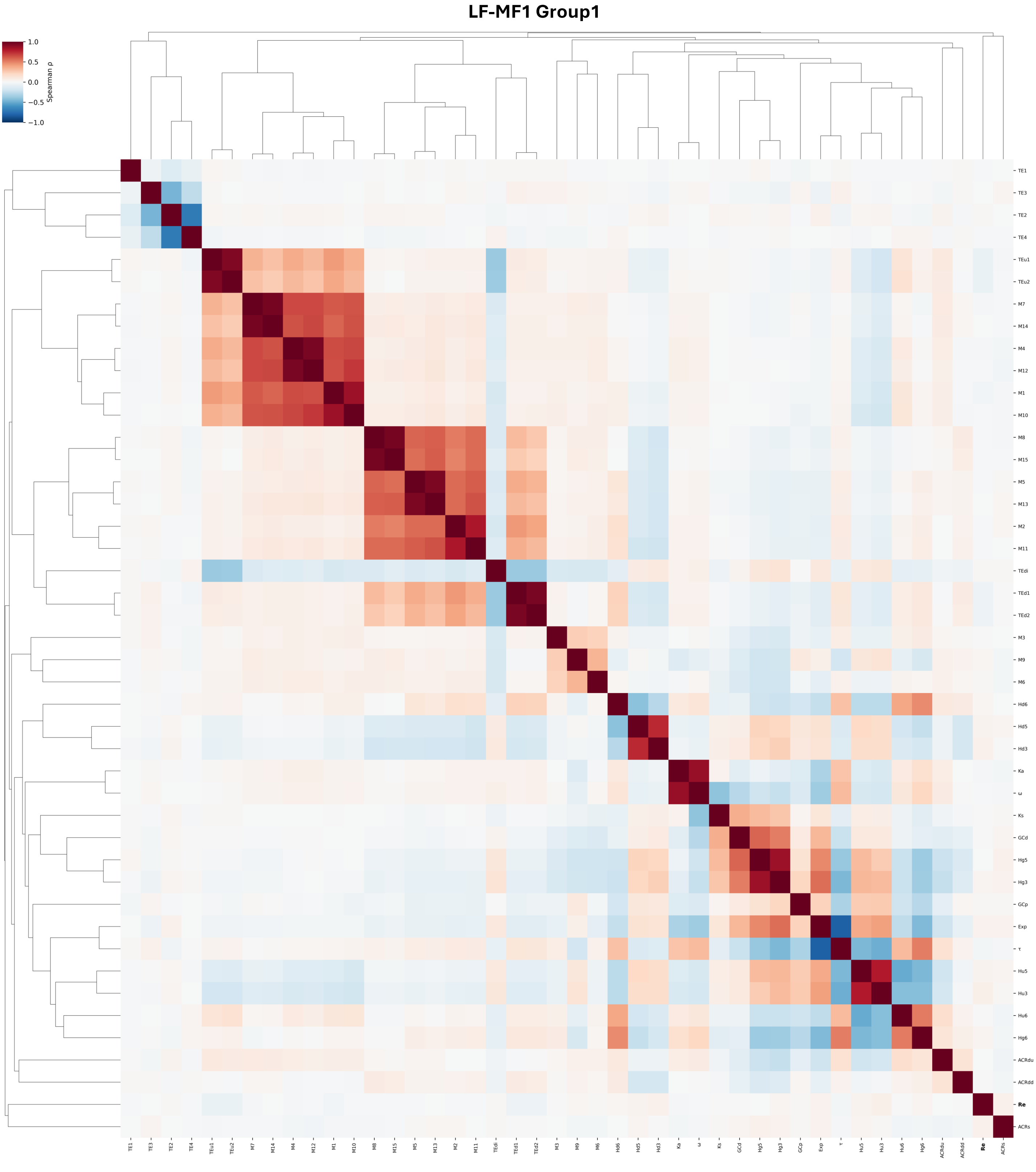


**Fig. S39. Spearman correlation heatmap of *B. rapa* LF-MF1 group1 genomic and epigenomic features.** Pairwise Spearman rank correlation coefficients were computed across all features using complete pairwise deletion for missing values. Features are clustered on both axes using average linkage on a distance matrix defined as 1 − |ρ|, where ρ is the Spearman correlation coefficient; this metric groups co-varying feature blocks symmetrically regardless of the direction of correlation. The color scale represents the Spearman ρ value (red = positive correlation, blue = negative correlation). Recombination rate (Re) is shown in bold.

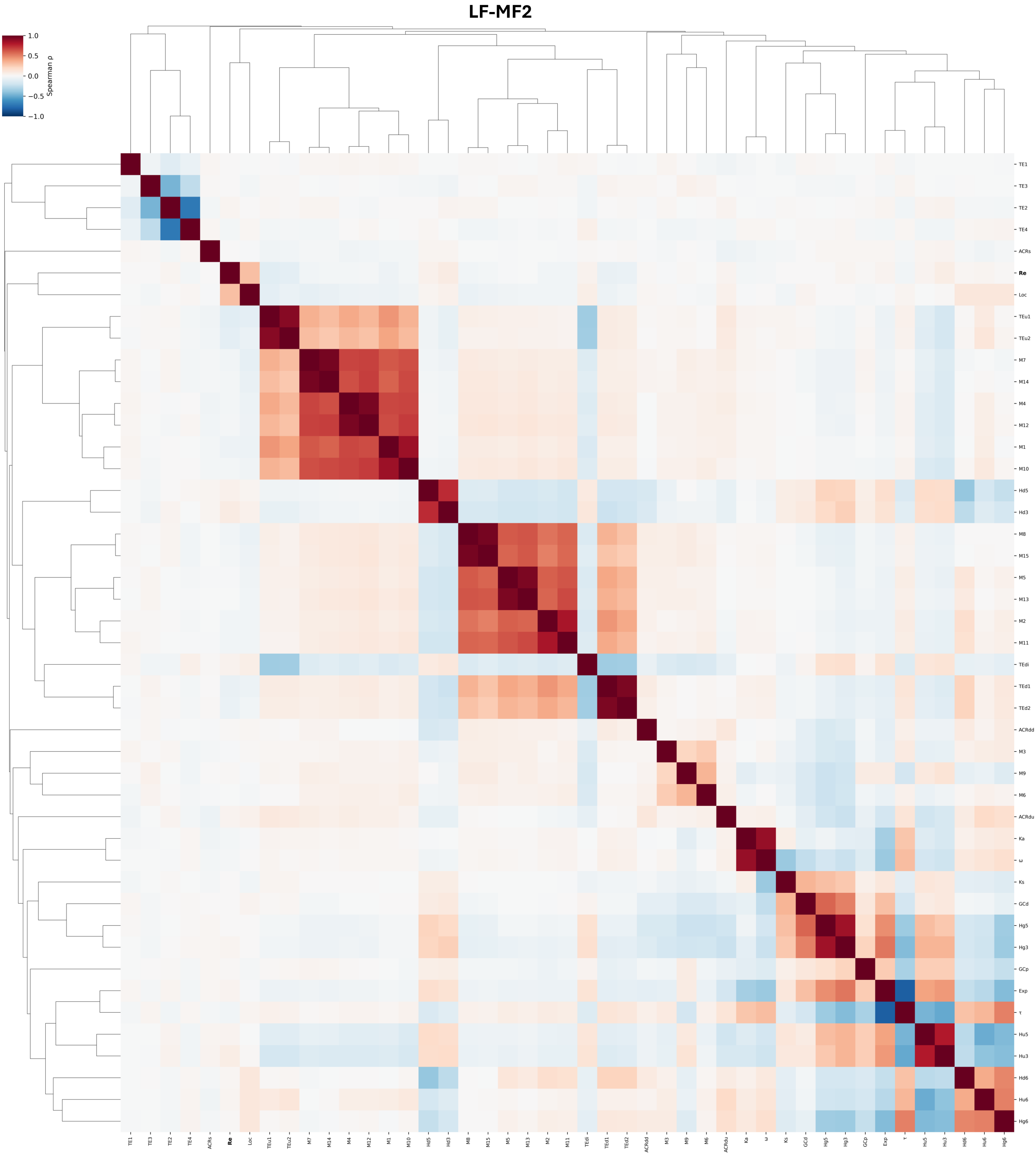


**Fig. S40. Spearman correlation heatmap of *B. rapa* LF-MF2 genomic and epigenomic features.** Pairwise Spearman rank correlation coefficients were computed across all features using complete pairwise deletion for missing values. Features are clustered on both axes using average linkage on a distance matrix defined as 1 − |ρ|, where ρ is the Spearman correlation coefficient; this metric groups co-varying feature blocks symmetrically regardless of the direction of correlation. The color scale represents the Spearman ρ value (red = positive correlation, blue = negative correlation). Recombination rate (Re) is shown in bold.

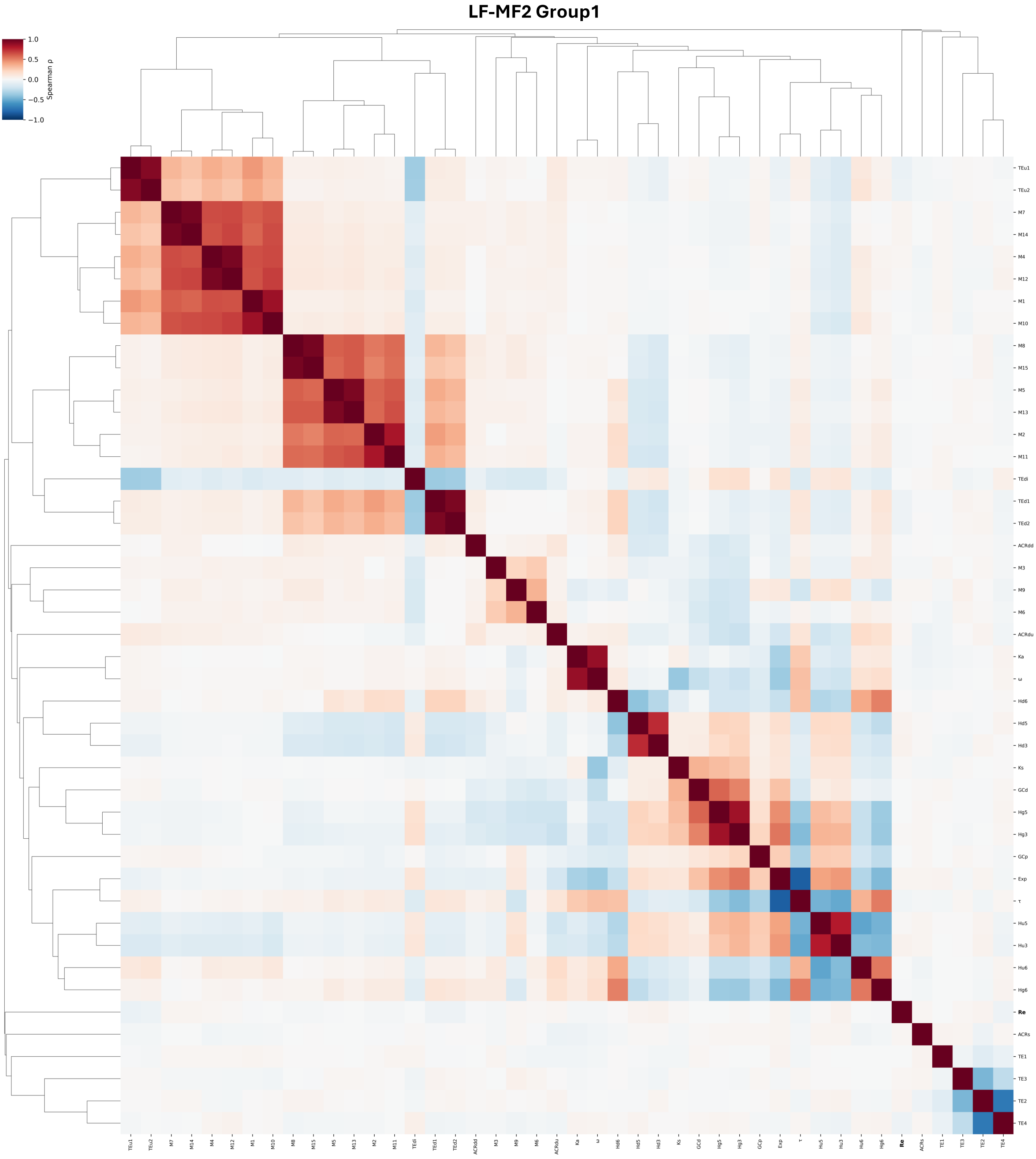


**Fig. S41. Spearman correlation heatmap of *B. rapa* LF-MF2 group1 genomic and epigenomic features.** Pairwise Spearman rank correlation coefficients were computed across all features using complete pairwise deletion for missing values. Features are clustered on both axes using average linkage on a distance matrix defined as 1 − |ρ|, where ρ is the Spearman correlation coefficient; this metric groups co-varying feature blocks symmetrically regardless of the direction of correlation. The color scale represents the Spearman ρ value (red = positive correlation, blue = negative correlation). Recombination rate (Re) is shown in bold.

## MF1-MF2

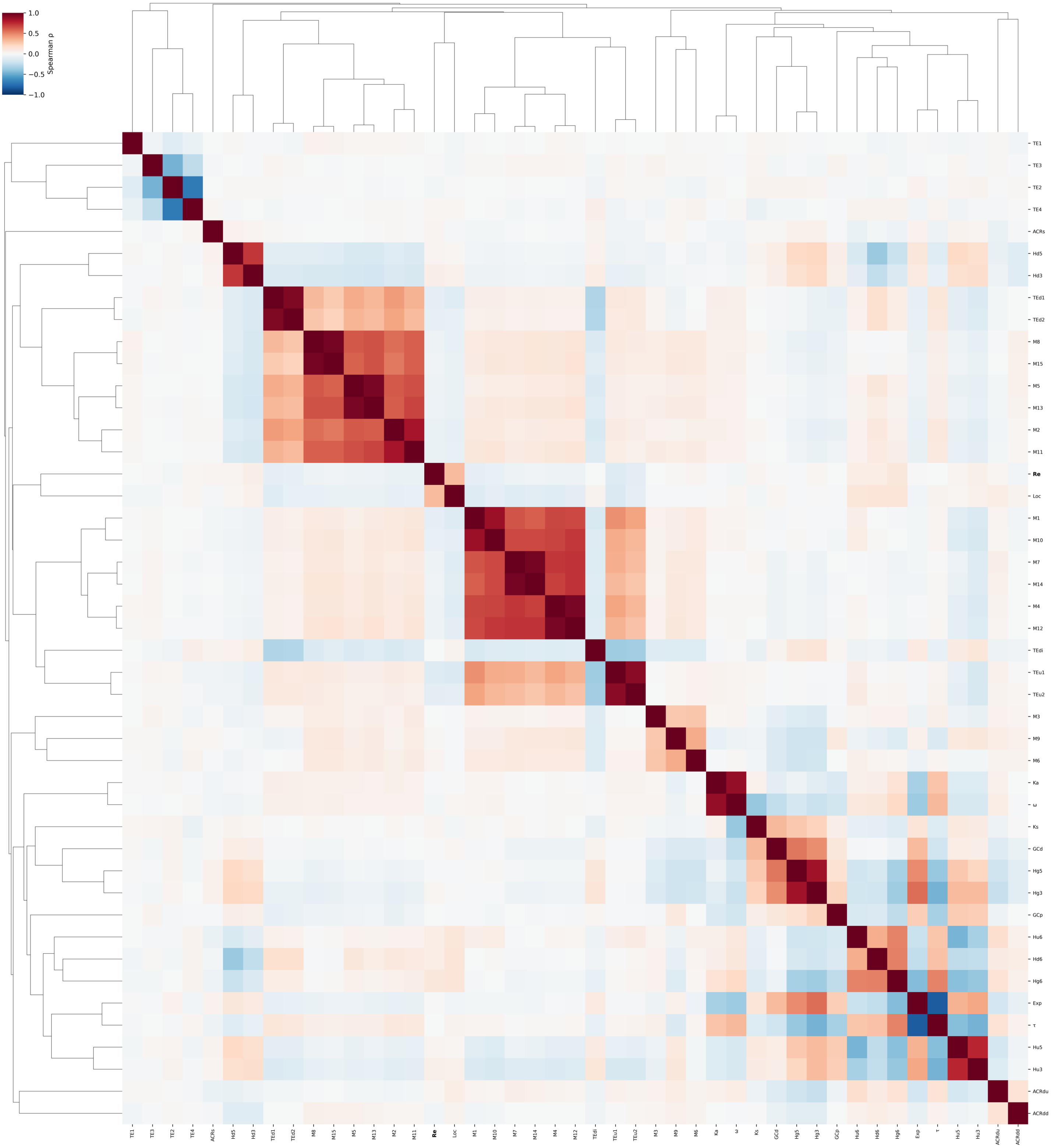


**Fig. S42. Spearman correlation heatmap of *B. rapa* MF1-MF2 genomic and epigenomic features.** Pairwise Spearman rank correlation coefficients were computed across all features using complete pairwise deletion for missing values. Features are clustered on both axes using average linkage on a distance matrix defined as 1 − |ρ|, where ρ is the Spearman correlation coefficient; this metric groups co-varying feature blocks symmetrically regardless of the direction of correlation. The color scale represents the Spearman ρ value (red = positive correlation, blue = negative correlation). Recombination rate (Re) is shown in bold.

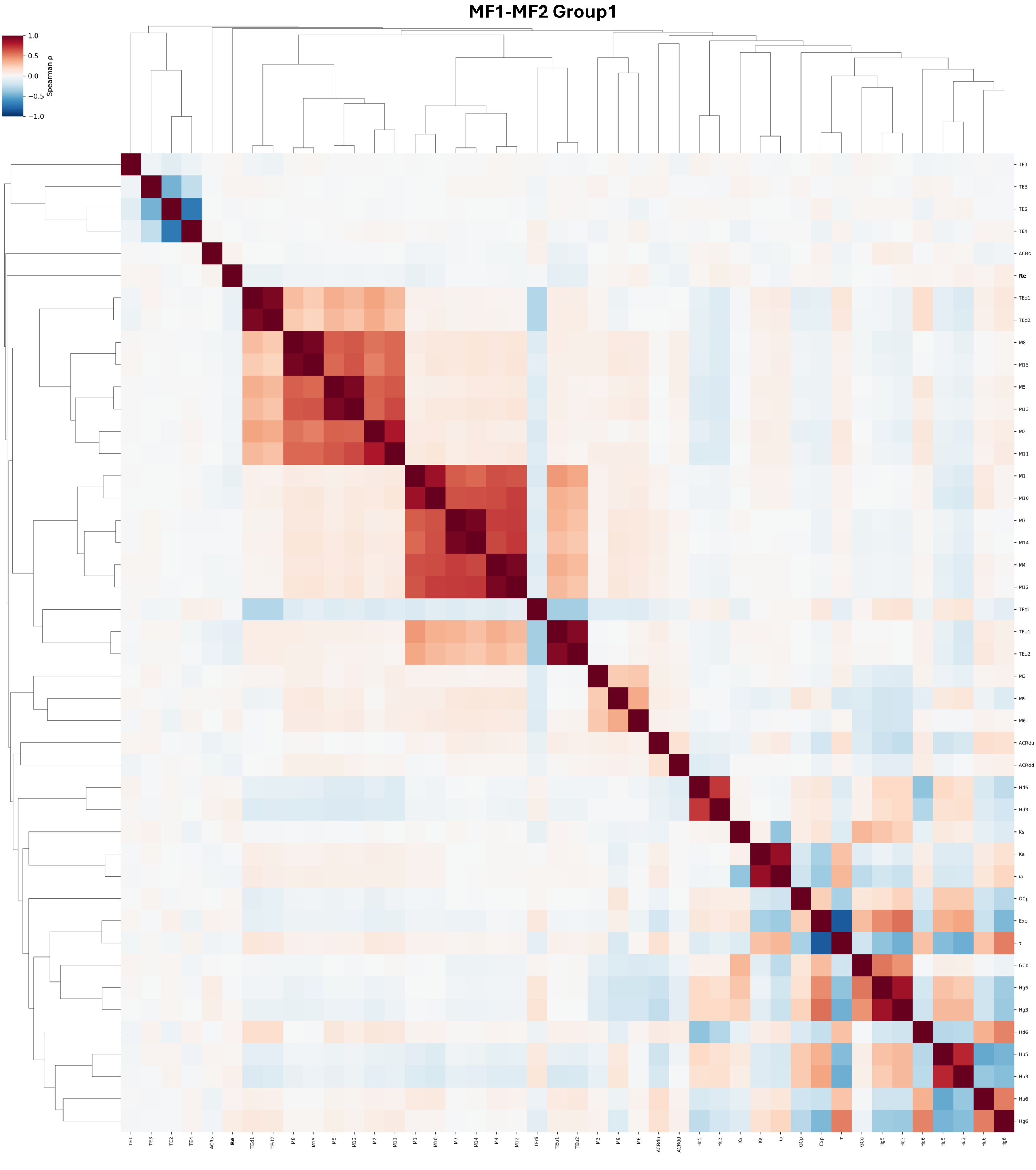


**Fig. S43. Spearman correlation heatmap of *B. rapa* MF1-MF2 group1 genomic and epigenomic features.** Pairwise Spearman rank correlation coefficients were computed across all features using complete pairwise deletion for missing values. Features are clustered on both axes using average linkage on a distance matrix defined as 1 − |ρ|, where ρ is the Spearman correlation coefficient; this metric groups co-varying feature blocks symmetrically regardless of the direction of correlation. The color scale represents the Spearman ρ value (red = positive correlation, blue = negative correlation). Recombination rate (Re) is shown in bold.

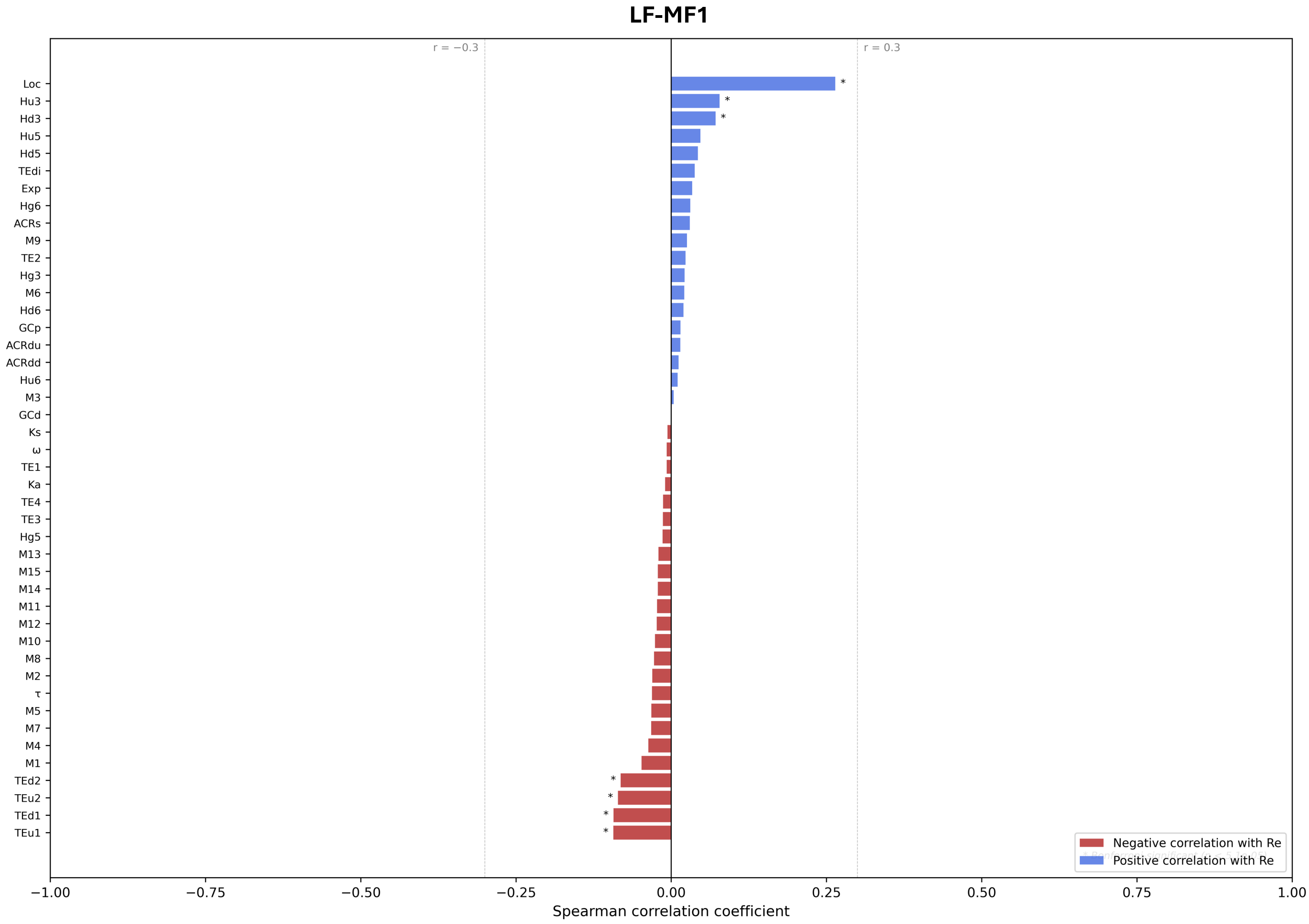


**Fig. S44. Spearman correlation of recombination rate (Re) with all *B. rapa* LF-MF1 features.** Features are ranked by Spearman ρ with Re (most negative at bottom). Bar color indicates direction of correlation: negative (red) or positive (blue). Asterisks indicate feature pairs surviving Bonferroni correction across all n(n−1)/2 pairwise tests within the group (α = 0.05). Dashed reference lines are drawn at ρ = ±0.3.

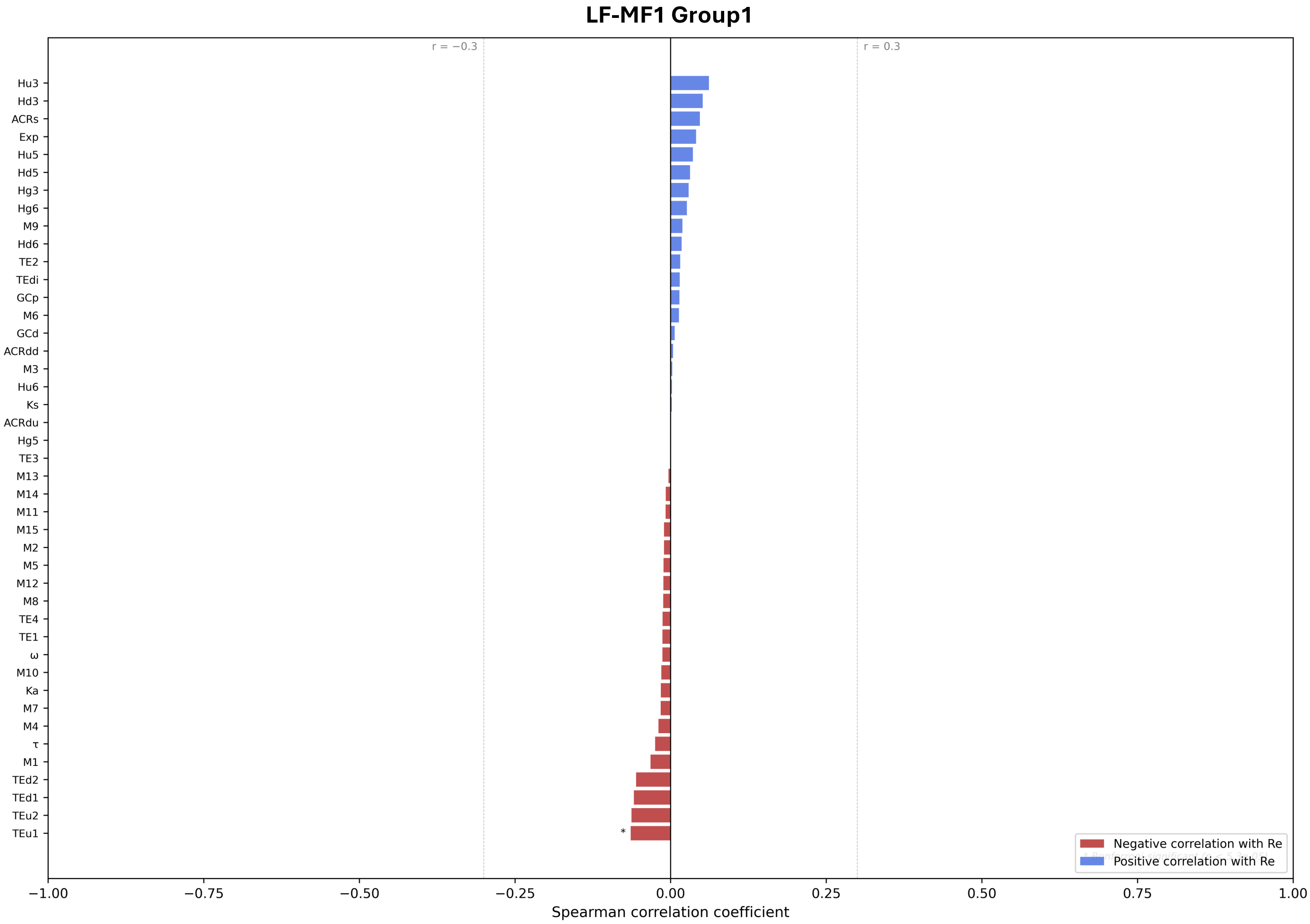


**Fig. S45. Spearman correlation of recombination rate (Re) with all *B. rapa* LF-MF1 group1 features.** Features are ranked by Spearman ρ with Re (most negative at bottom). Bar color indicates direction of correlation: negative (red) or positive (blue). Asterisks indicate feature pairs surviving Bonferroni correction across all n(n−1)/2 pairwise tests within the group (α = 0.05). Dashed reference lines are drawn at ρ = ±0.3.

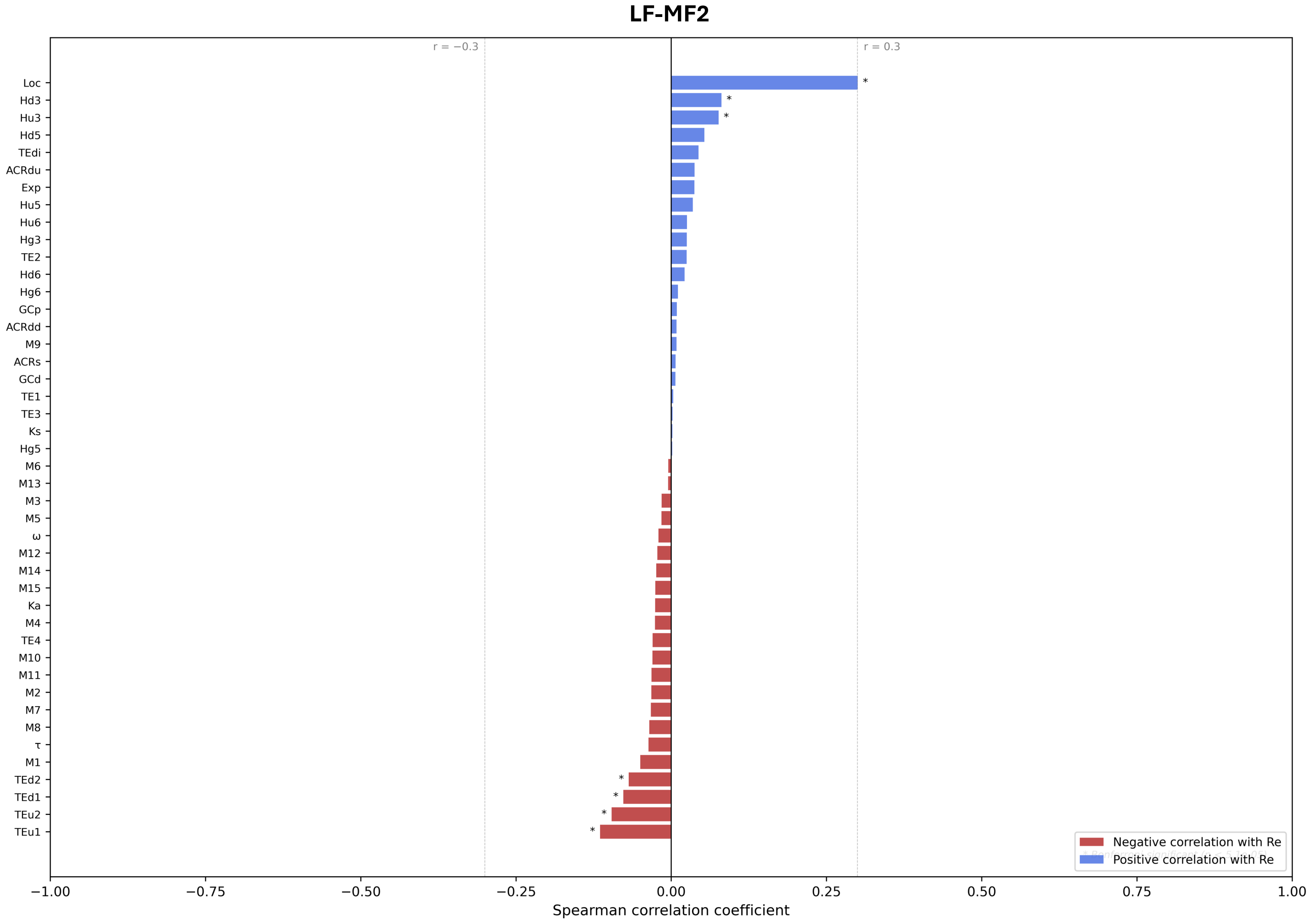


**Fig. S46. Spearman correlation of recombination rate (Re) with all *B. rapa* LF-MF2 features.** Features are ranked by Spearman ρ with Re (most negative at bottom). Bar color indicates direction of correlation: negative (red) or positive (blue). Asterisks indicate feature pairs surviving Bonferroni correction across all n(n−1)/2 pairwise tests within the group (α = 0.05). Dashed reference lines are drawn at ρ = ±0.3.

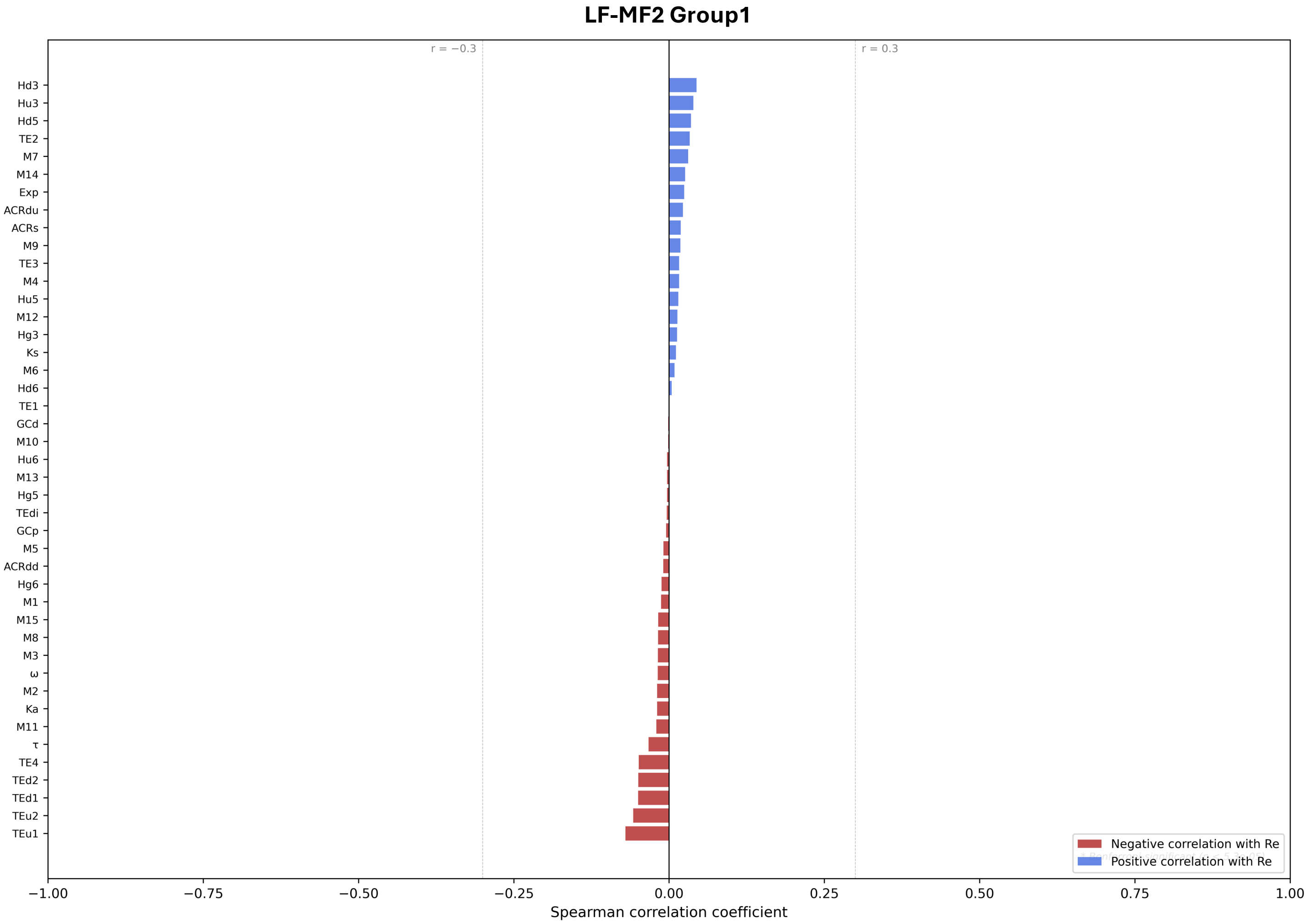


**Fig. S47. Spearman correlation of recombination rate (Re) with all *B. rapa* LF-MF2 group1 features.** Features are ranked by Spearman ρ with Re (most negative at bottom). Bar color indicates direction of correlation: negative (red) or positive (blue). Asterisks indicate feature pairs surviving Bonferroni correction across all n(n−1)/2 pairwise tests within the group (α = 0.05). Dashed reference lines are drawn at ρ = ±0.3.

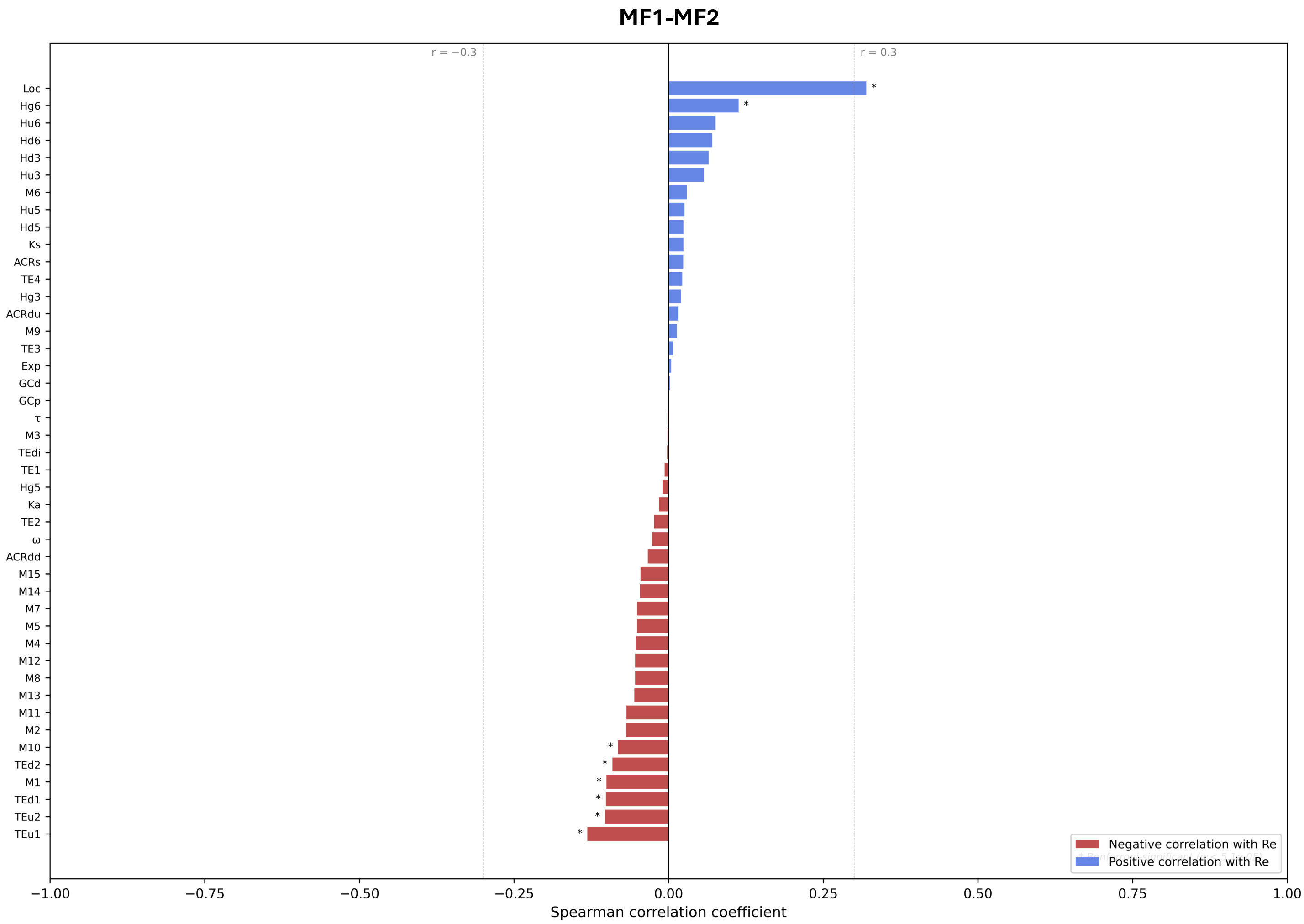


**Fig. S48. Spearman correlation of recombination rate (Re) with all *B. rapa* MF1-MF2 features.** Features are ranked by Spearman ρ with Re (most negative at bottom). Bar color indicates direction of correlation: negative (red) or positive (blue). Asterisks indicate feature pairs surviving Bonferroni correction across all n(n−1)/2 pairwise tests within the group (α = 0.05). Dashed reference lines are drawn at ρ = ±0.3.

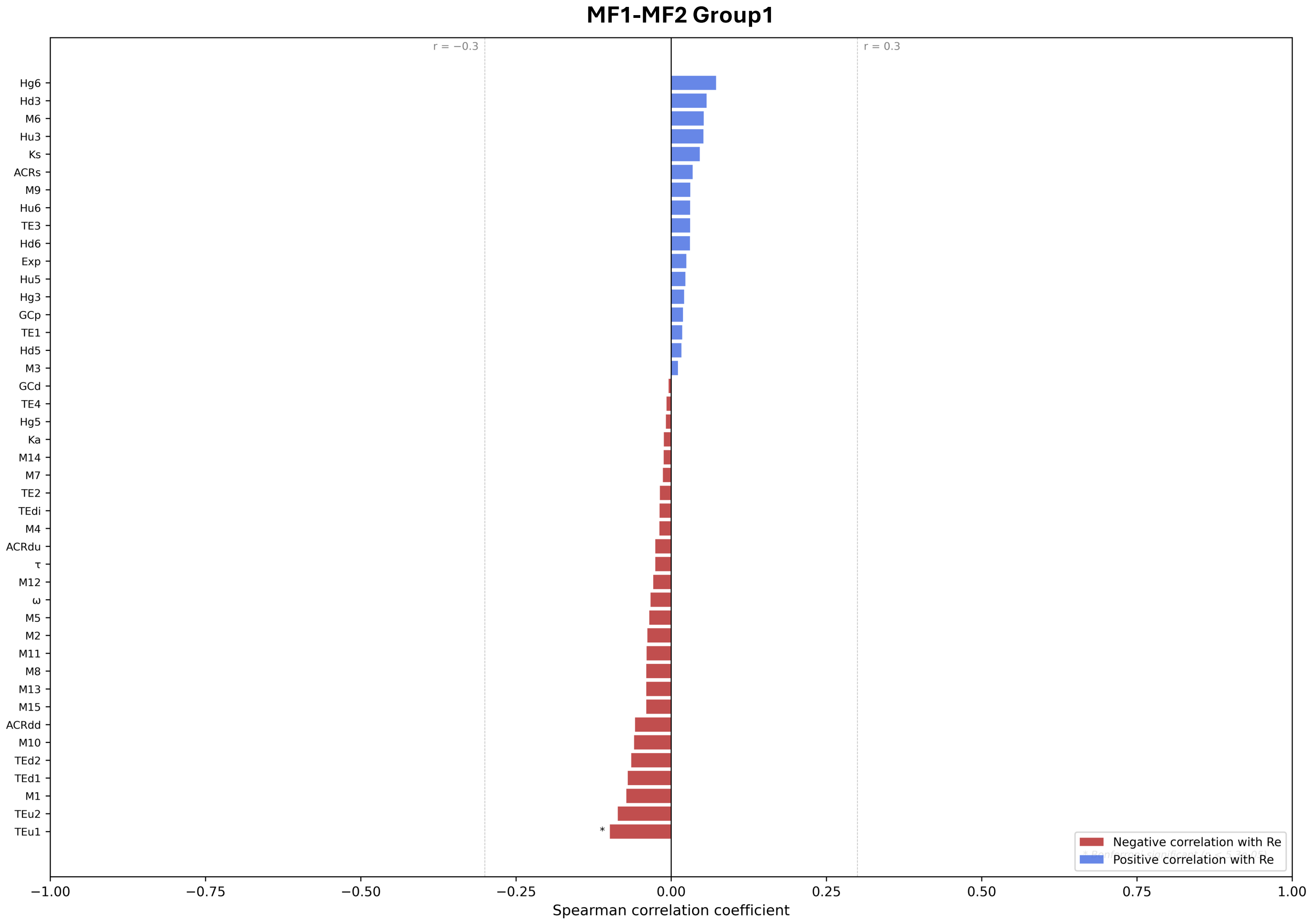


**Fig. S49. Spearman correlation of recombination rate (Re) with all *B. rapa* MF1-MF2 group1 features.** Features are ranked by Spearman ρ with Re (most negative at bottom). Bar color indicates direction of correlation: negative (red) or positive (blue). Asterisks indicate feature pairs surviving Bonferroni correction across all n(n−1)/2 pairwise tests within the group (α = 0.05). Dashed reference lines are drawn at ρ = ±0.3.

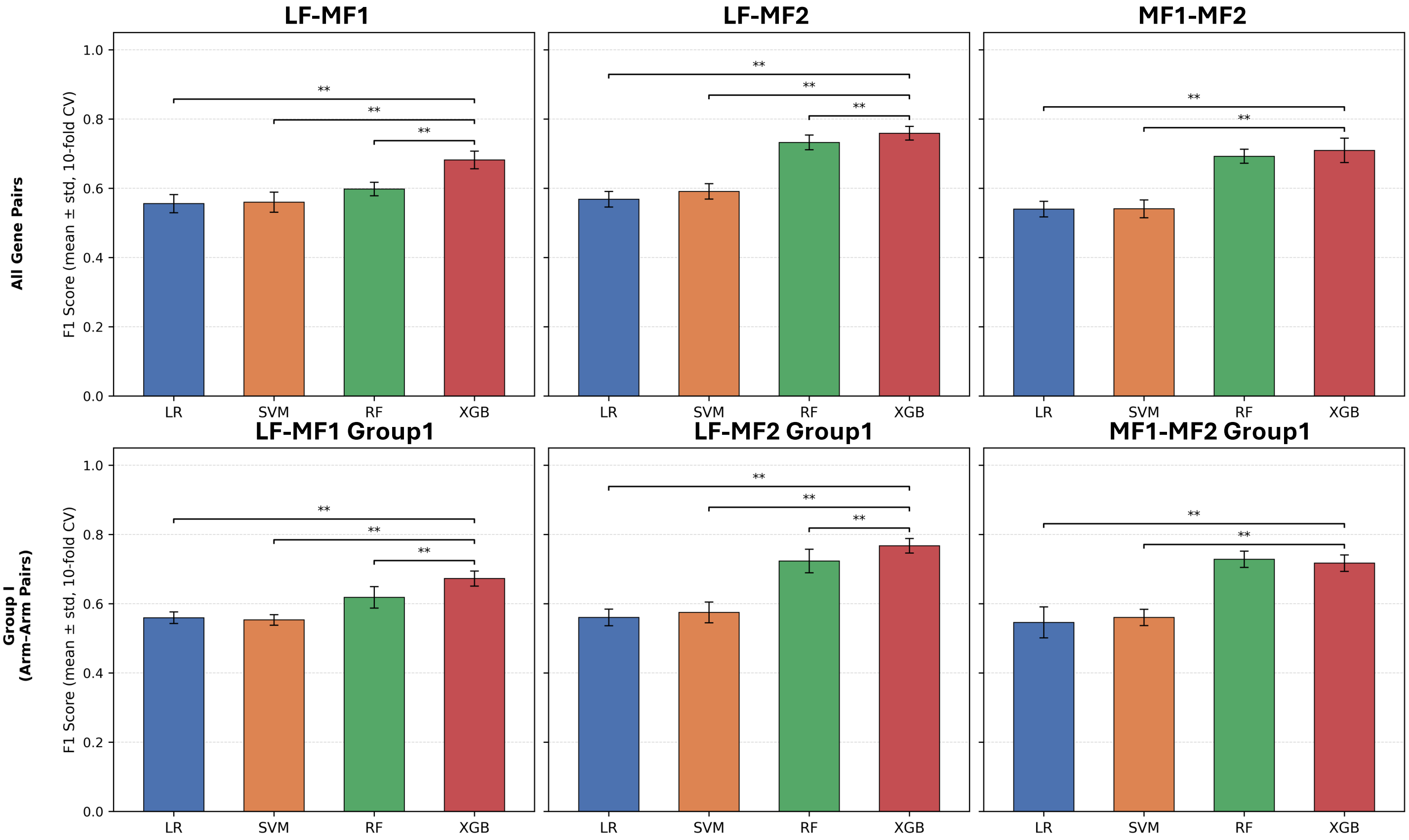


**Fig. S50. Comparison of classification performance across model architectures for each *B. rapa* subgenome pair.** Mean F1 score (± standard deviation across 10 stratified cross-validation folds) is shown for each model architecture: Logistic Regression (LR), Support Vector Machine (SVM), Random Forest (RF), and XGBoost (XGB). Pairwise significance was assessed between each baseline architecture and XGB using two-sided Wilcoxon signed-rank tests on paired fold-level F1 scores, with Benjamini-Hochberg false discovery rate correction applied within each comparison group independently. Only comparisons reaching adjusted $P < 0.05$ are annotated (* $P < 0.05$, ** $P < 0.01$, *** $P < 0.001$). Comparisons across subgenome pairs or between all-pairs and Group I panels are not made. All models used default hyperparameters; feature scaling was applied per fold for LR and SVM only.

### LF-MF1 All Genes

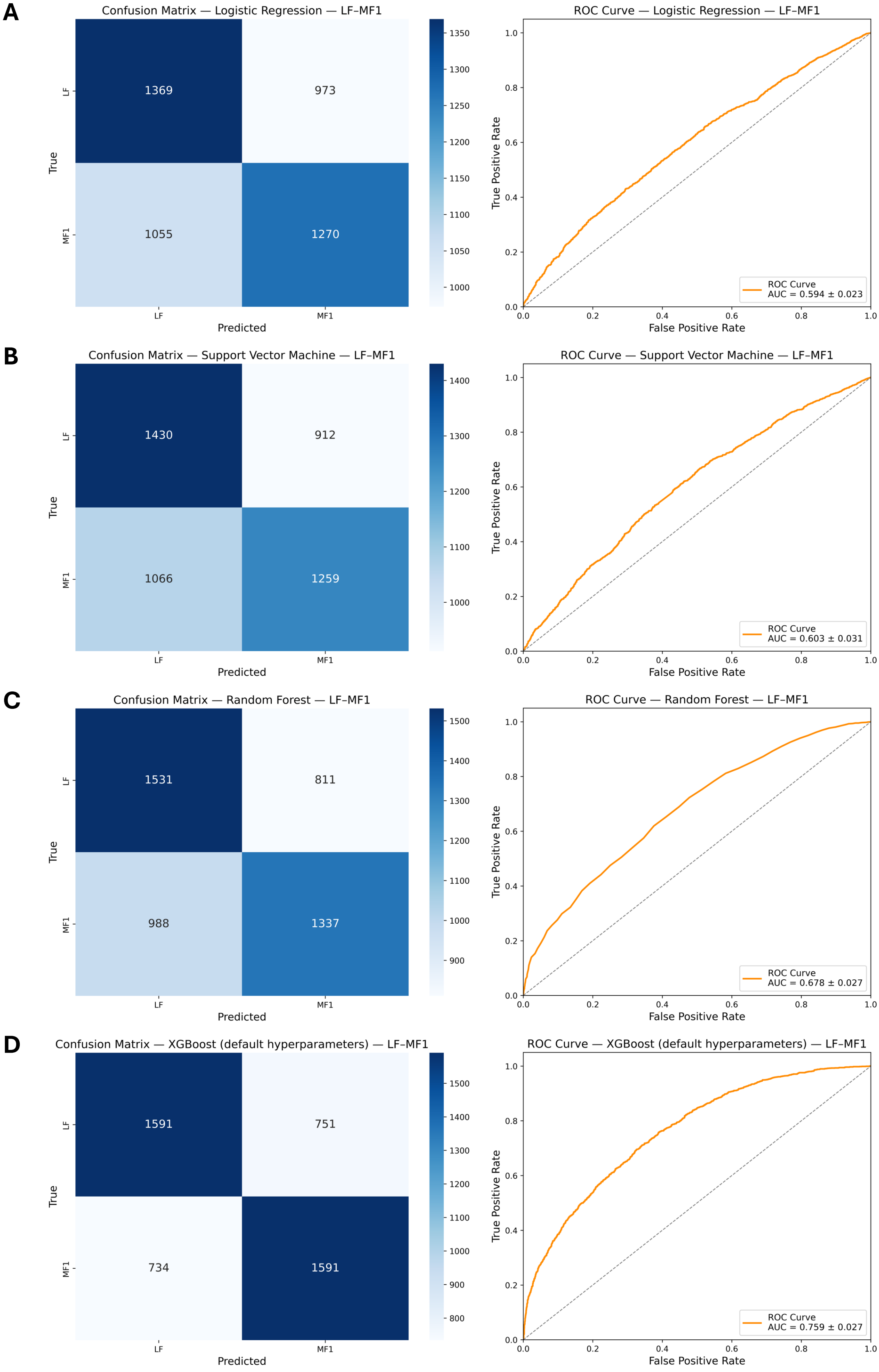


**Fig. S51. Benchmarking unoptimized model performance on *B. rapa* LF-MF1 dataset.** Confusion matrices and ROC/AUC from logistic regression (**A**), support vector machine (**B**), random forest (**C**), and XGBoost (**D**) model architectures.

LF-MF2 All Genes

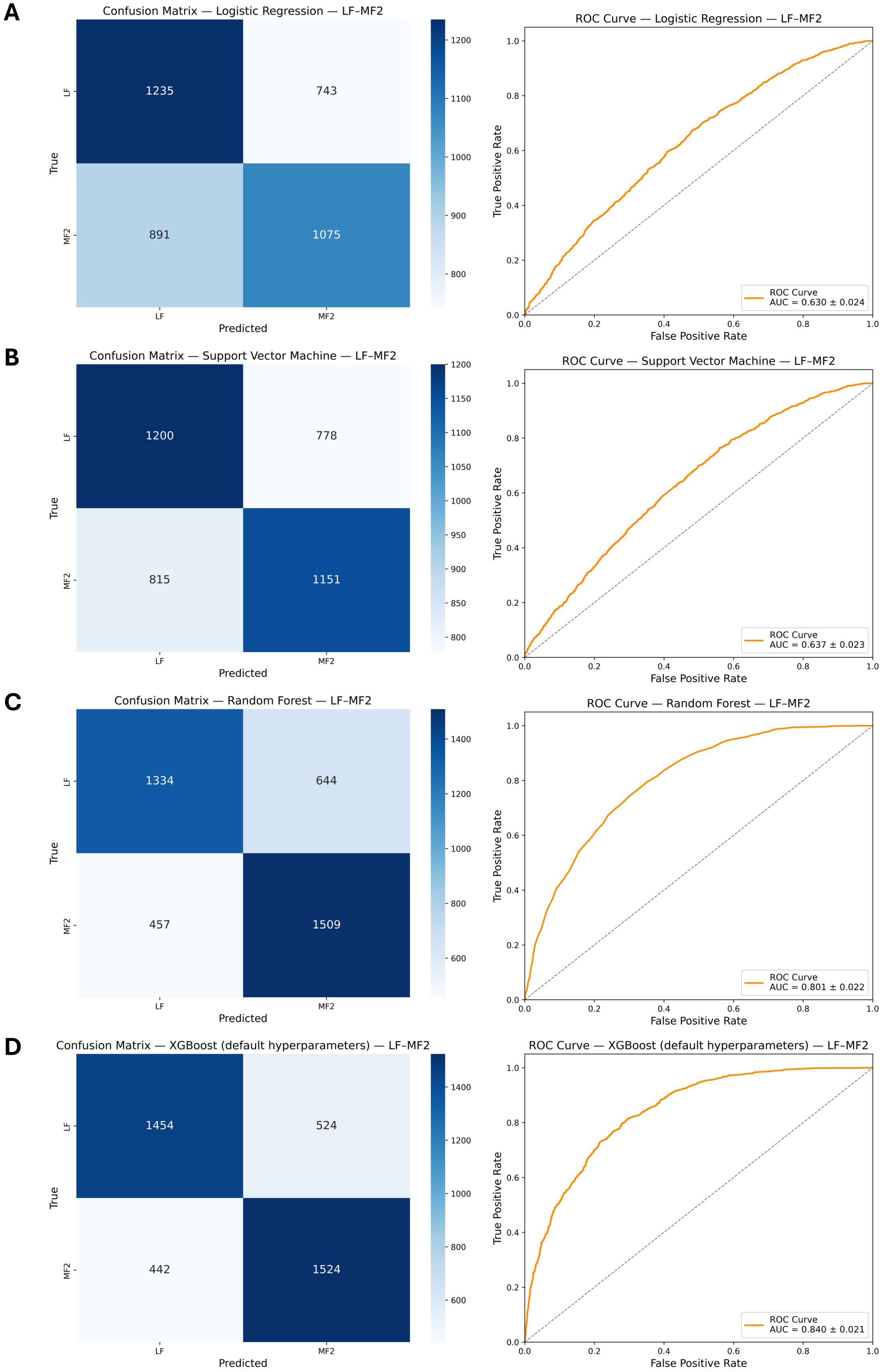


**Fig. S52. Benchmarking unoptimized model performance on *B. rapa* LF-MF2 dataset.** Confusion matrices and ROC/AUC from logistic regression (**A**), support vector machine (**B**), random forest (**C**), and XGBoost (**D**) model architectures.

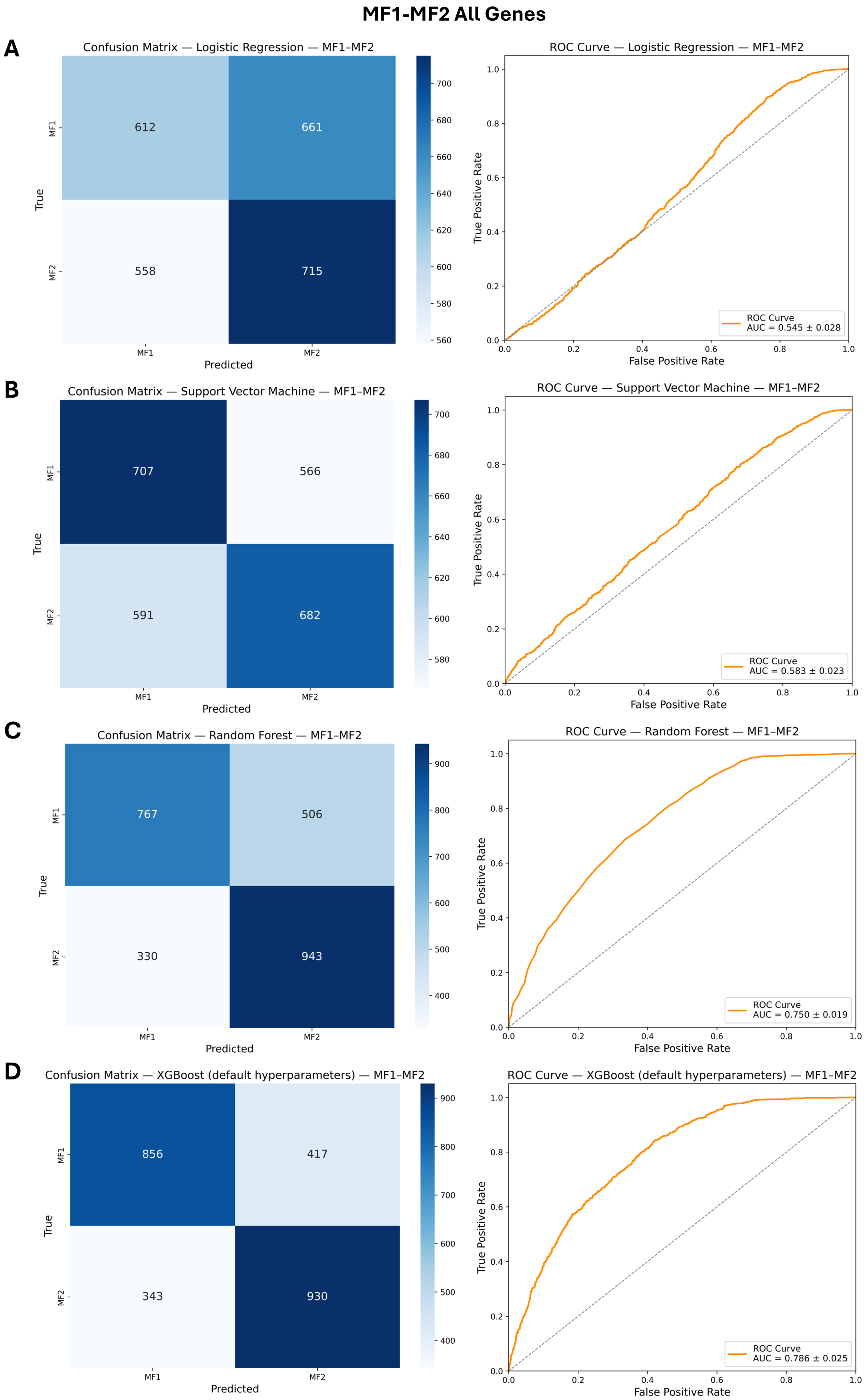


**Fig. S53. Benchmarking unoptimized model performance on *B. rapa* MF1-MF2 dataset.** Confusion matrices and ROC/AUC from logistic regression (**A**), support vector machine (**B**), random forest (**C**), and XGBoost (**D**) model architectures.

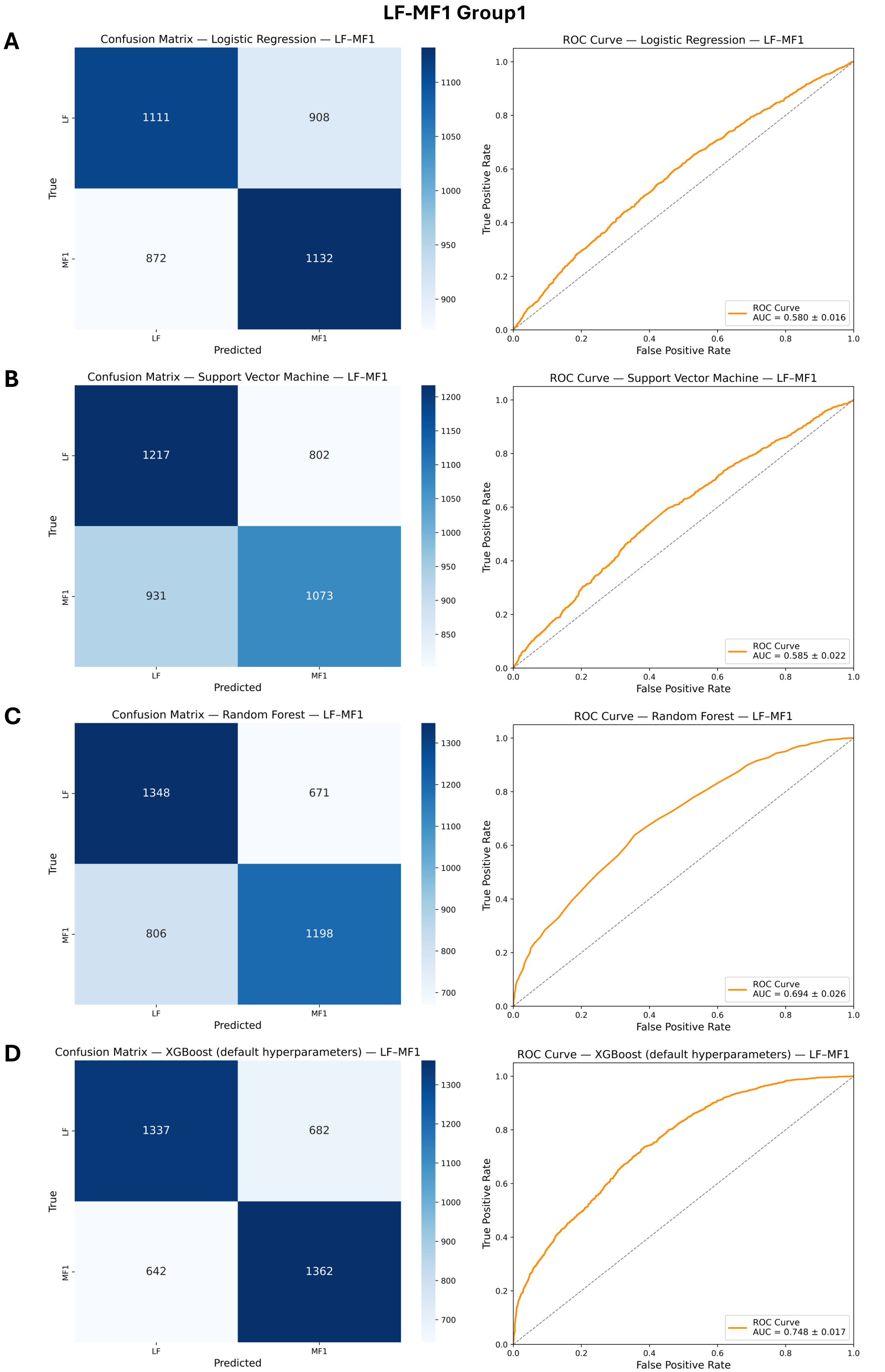


**Fig. S54. Benchmarking unoptimized model performance on *B. rapa* LF-MF1 group1 dataset.** Confusion matrices and ROC/AUC from logistic regression (**A**), support vector machine (**B**), random forest (**C**), and XGBoost (**D**) model architectures.

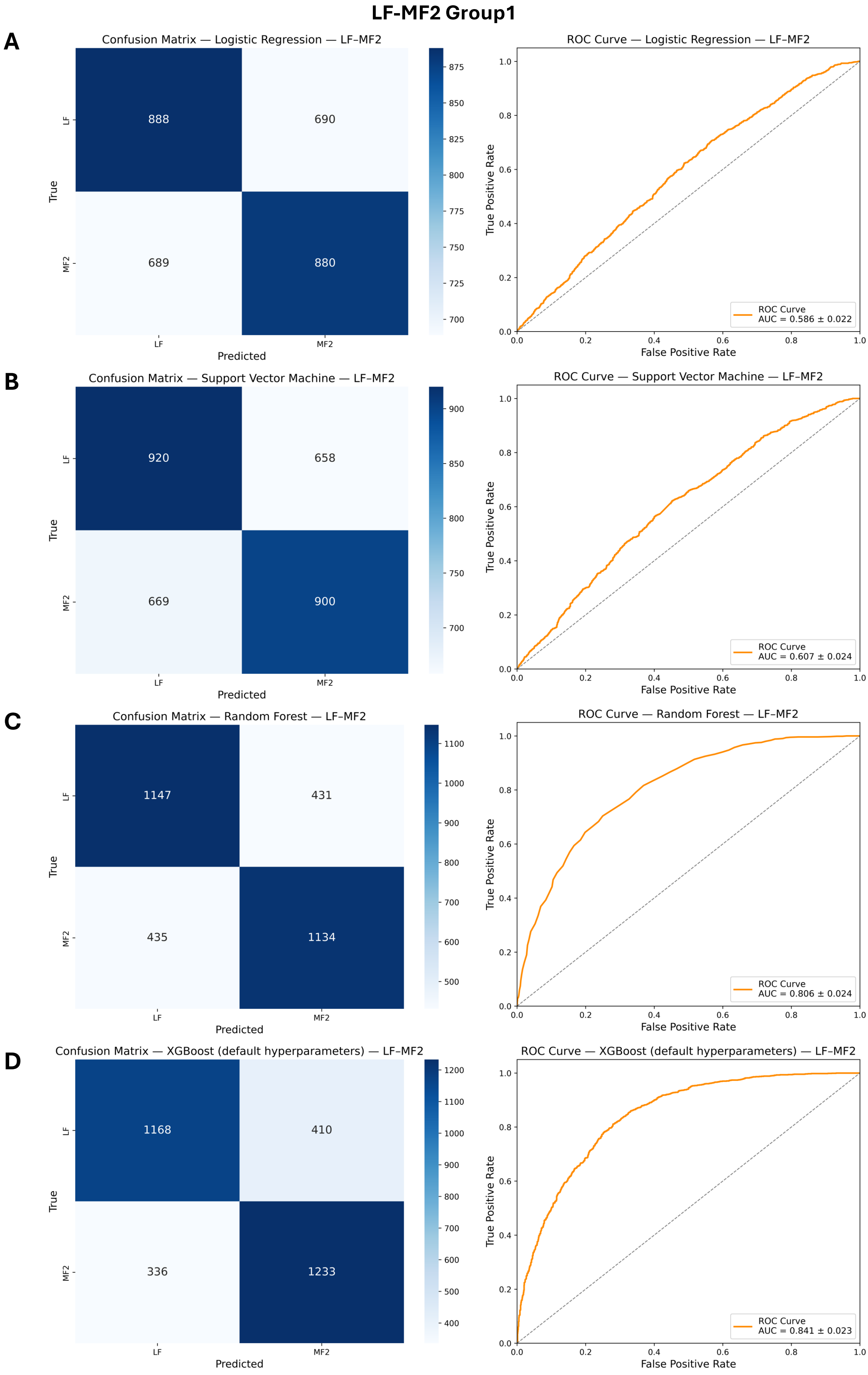


**Fig. S55. Benchmarking unoptimized model performance on *B. rapa* LF-MF2 group1 dataset.** Confusion matrices and ROC/AUC from logistic regression (**A**), support vector machine (**B**), random forest (**C**), and XGBoost (**D**) model architectures.

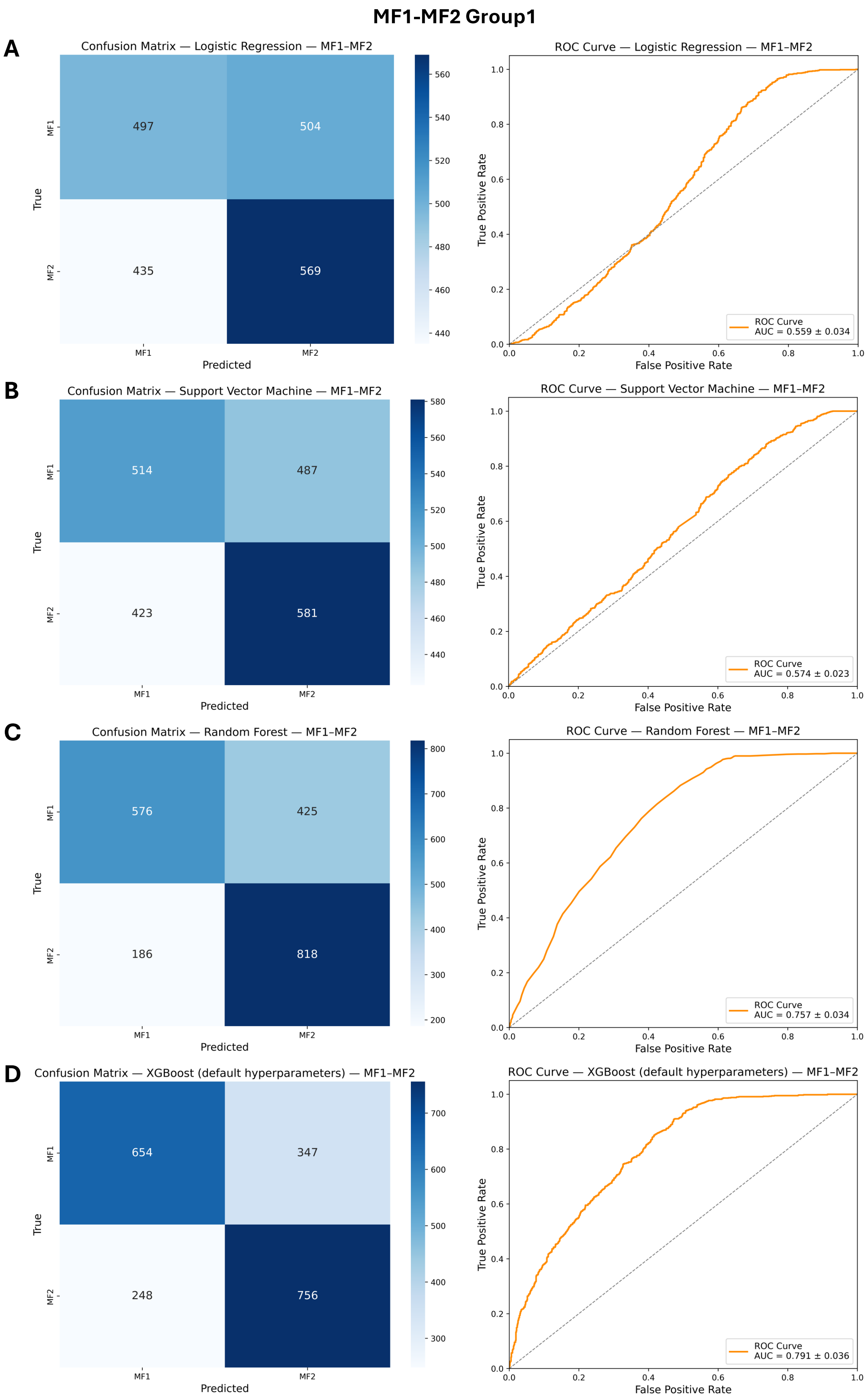


**Fig. S56. Benchmarking unoptimized model performance on *B. rapa* MF1-MF2 group1 dataset.** Confusion matrices and ROC/AUC from logistic regression (**A**), support vector machine (**B**), random forest (**C**), and XGBoost (**D**) model architectures.

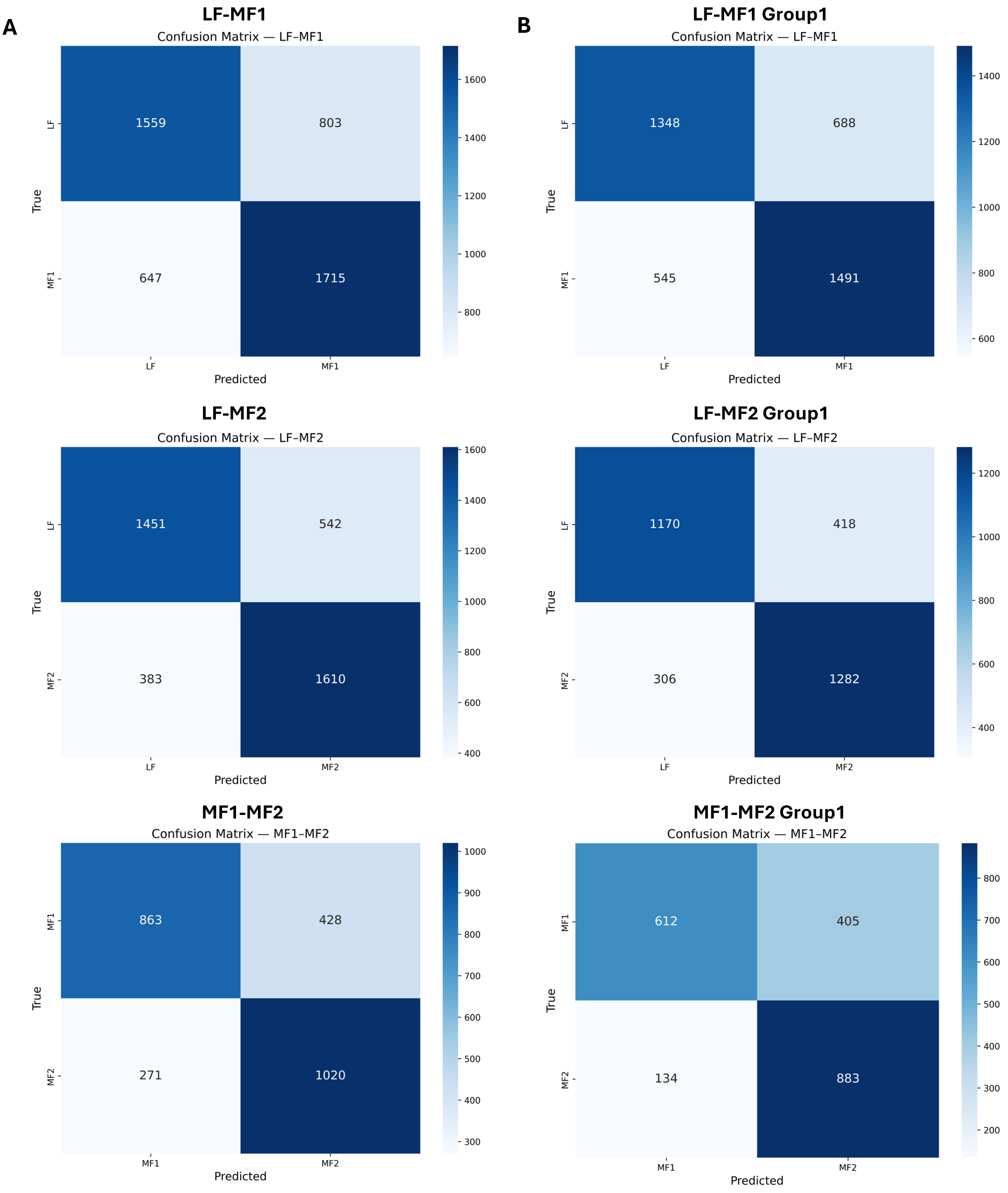


**Fig. S57. Confusion matrices of six final XGBoost *B. rapa* models.** All genes (**A**) and group1 genes (**B**).

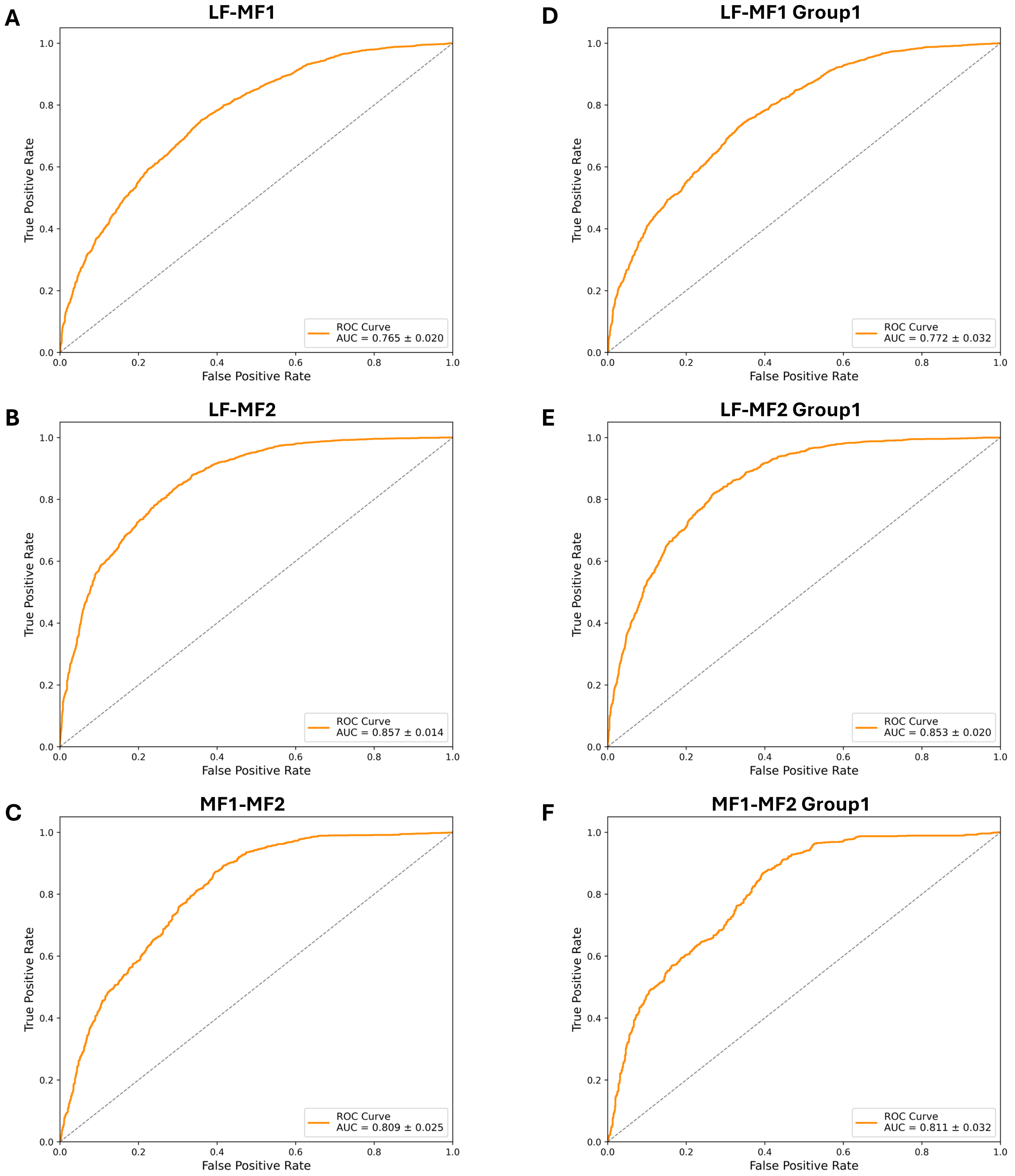


**Fig. S58. Performance metrics for final *B. rapa* models.** ROC/AUC for all-genes models LF-MF1 (**A**), LF-MF2 (**B**), MF1-MF2 (**C**), LF-MF1 group1 (**D**), LF-MF2 group1 (**E**), MF1-MF2 group1 (**F**) showing mean and standard deviation across 10 folds; F1 scores for six final *B. rapa* models, mean and standard deviation across 10 folds (**G**).

**G**

| Subgenome Pair | Mean F1 ± std |
|---|---|
| LF-MF1 | 0.703±0.019 |
| LF-MF1 group1 | 0.707±0.031 |
| LF-MF2 | 0.777±0.012 |
| LF-MF2 group1 | 0.780±0.021 |
| MF1-MF2 | 0.745±0.031 |
| MF1-MF2 group1 | 0.766±0.025 |

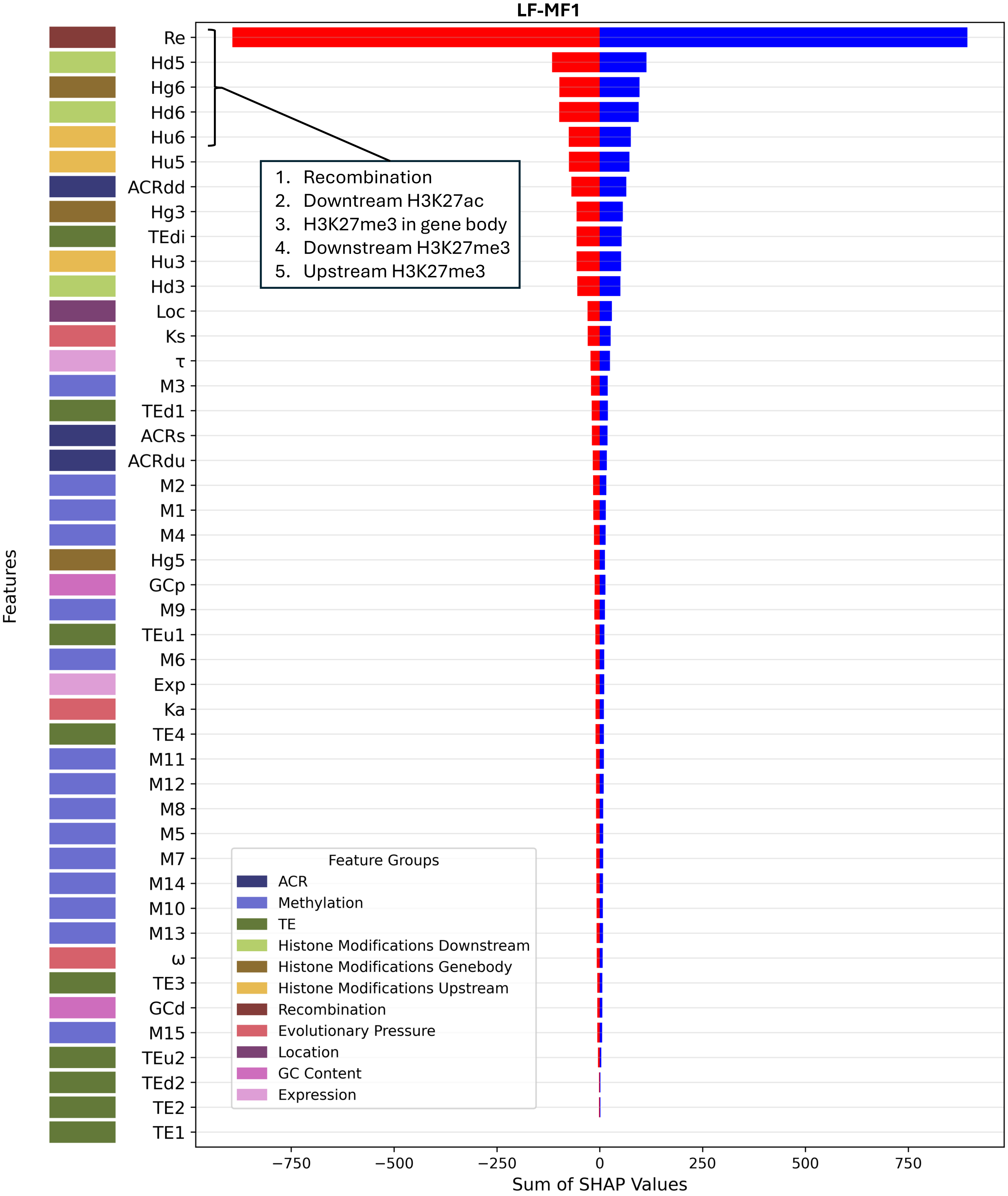


**Fig. S59. Directional SHAP feature importance profile for *B. rapa* LF-MF1.** Features are ranked by the sum of their absolute SHAP values across all test-fold samples (ascending order, highest-importance features at top). For each feature, SHAP values are summed separately across positive contributions (blue; favoring MF1 classification) and negative contributions (red; favoring LF classification), reflecting total directional influence across the test set rather than per-sample magnitude. Colored tiles in the left margin indicate feature group membership. The five features contributing most to classification, as determined by summed absolute SHAP value, are listed in the inset.

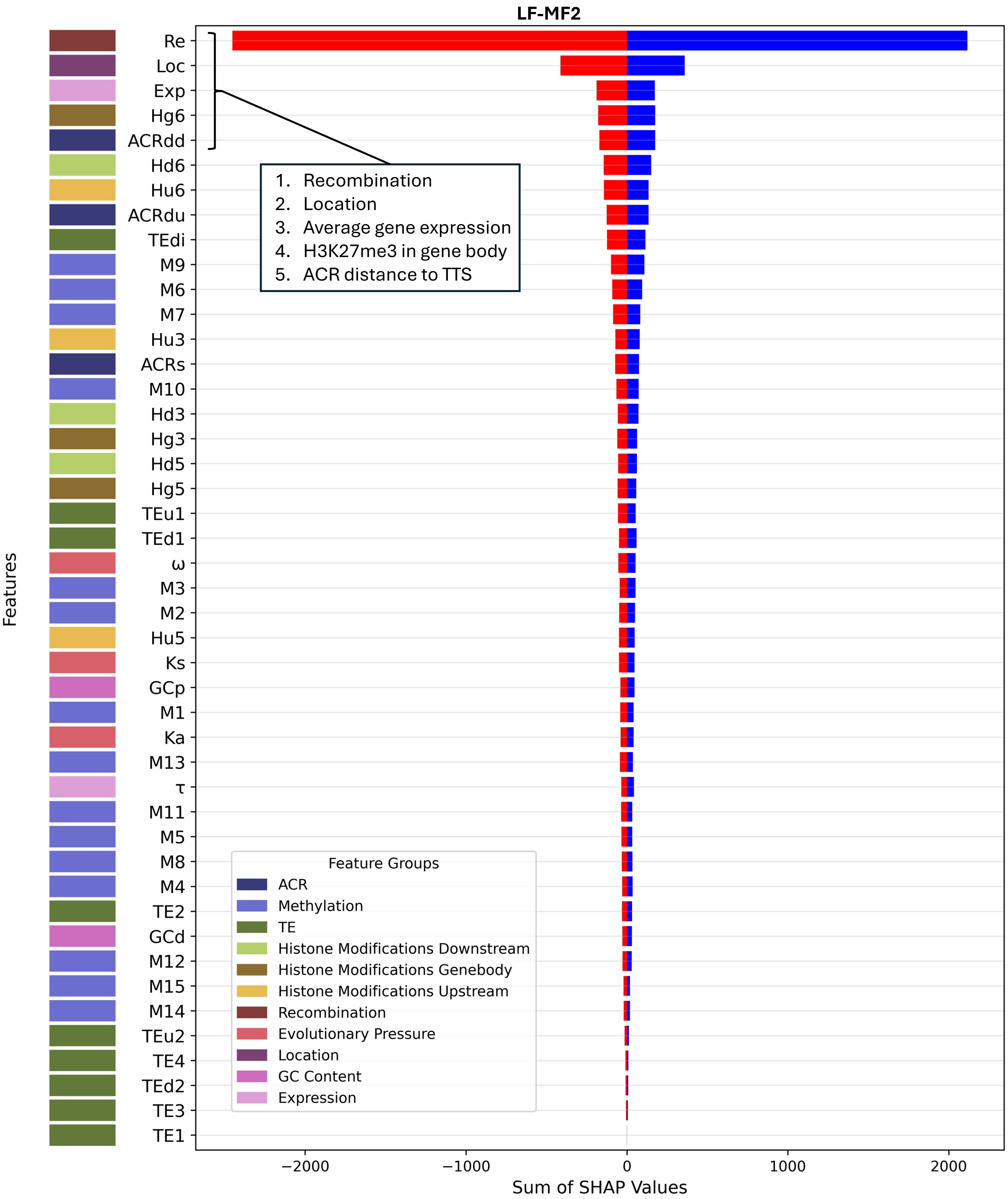


**Fig. S60. Directional SHAP feature importance profile for *B. rapa* LF-MF2.** Features are ranked by the sum of their absolute SHAP values across all test-fold samples (ascending order, highest-importance features at top). For each feature, SHAP values are summed separately across positive contributions (blue; favoring MF2 classification) and negative contributions (red; favoring LF classification), reflecting total directional influence across the test set rather than per-sample magnitude. Colored tiles in the left margin indicate feature group membership. The five features contributing most to classification, as determined by summed absolute SHAP value, are listed in the inset.

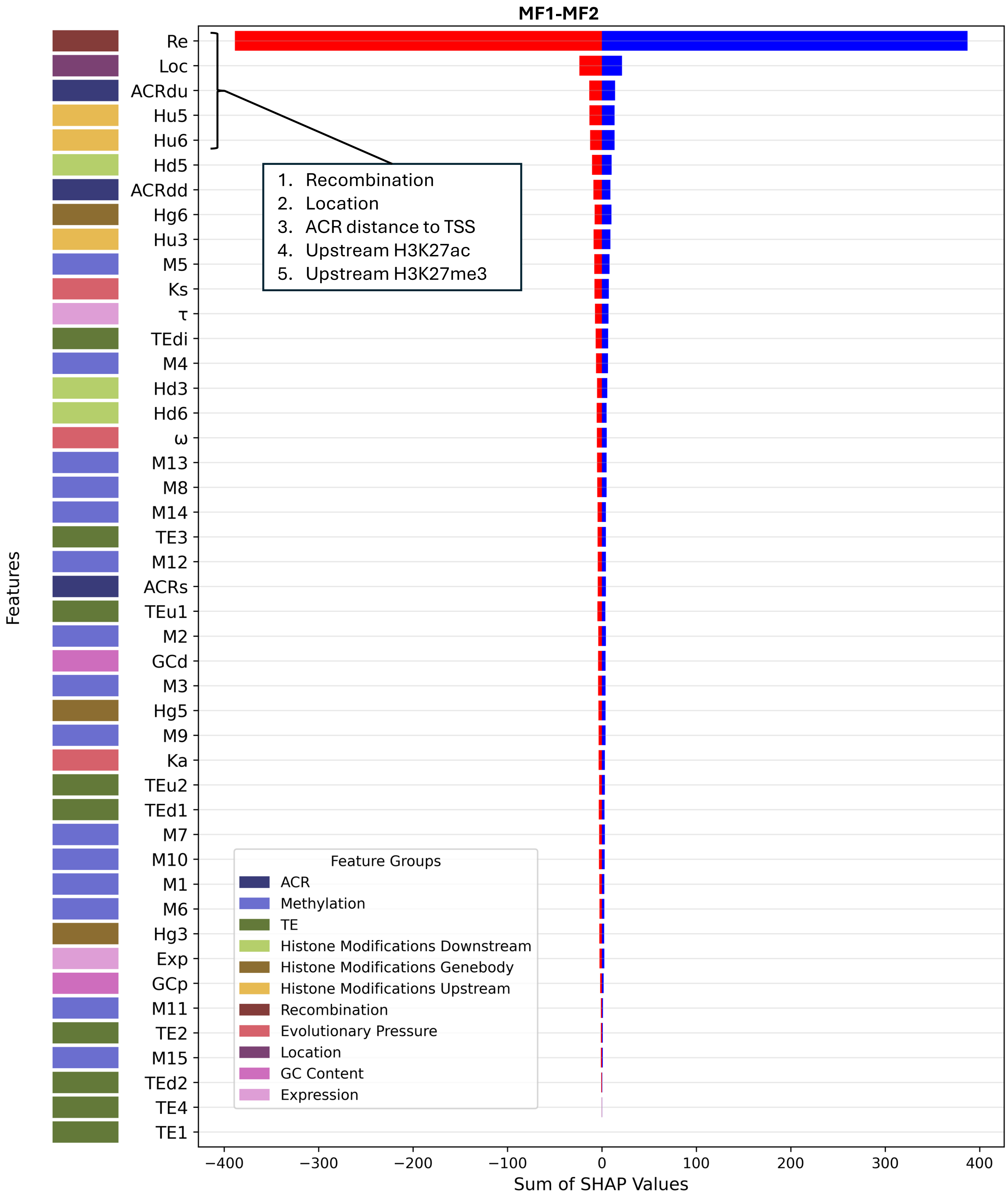


**Fig. S61. Directional SHAP feature importance profile for *B. rapa* MF1-MF2.** Features are ranked by the sum of their absolute SHAP values across all test-fold samples (ascending order, highest-importance features at top). For each feature, SHAP values are summed separately across positive contributions (blue; favoring MF2 classification) and negative contributions (red; favoring MF1 classification), reflecting total directional influence across the test set rather than per-sample magnitude. Colored tiles in the left margin indicate feature group membership. The five features contributing most to classification, as determined by summed absolute SHAP value, are listed in the inset.

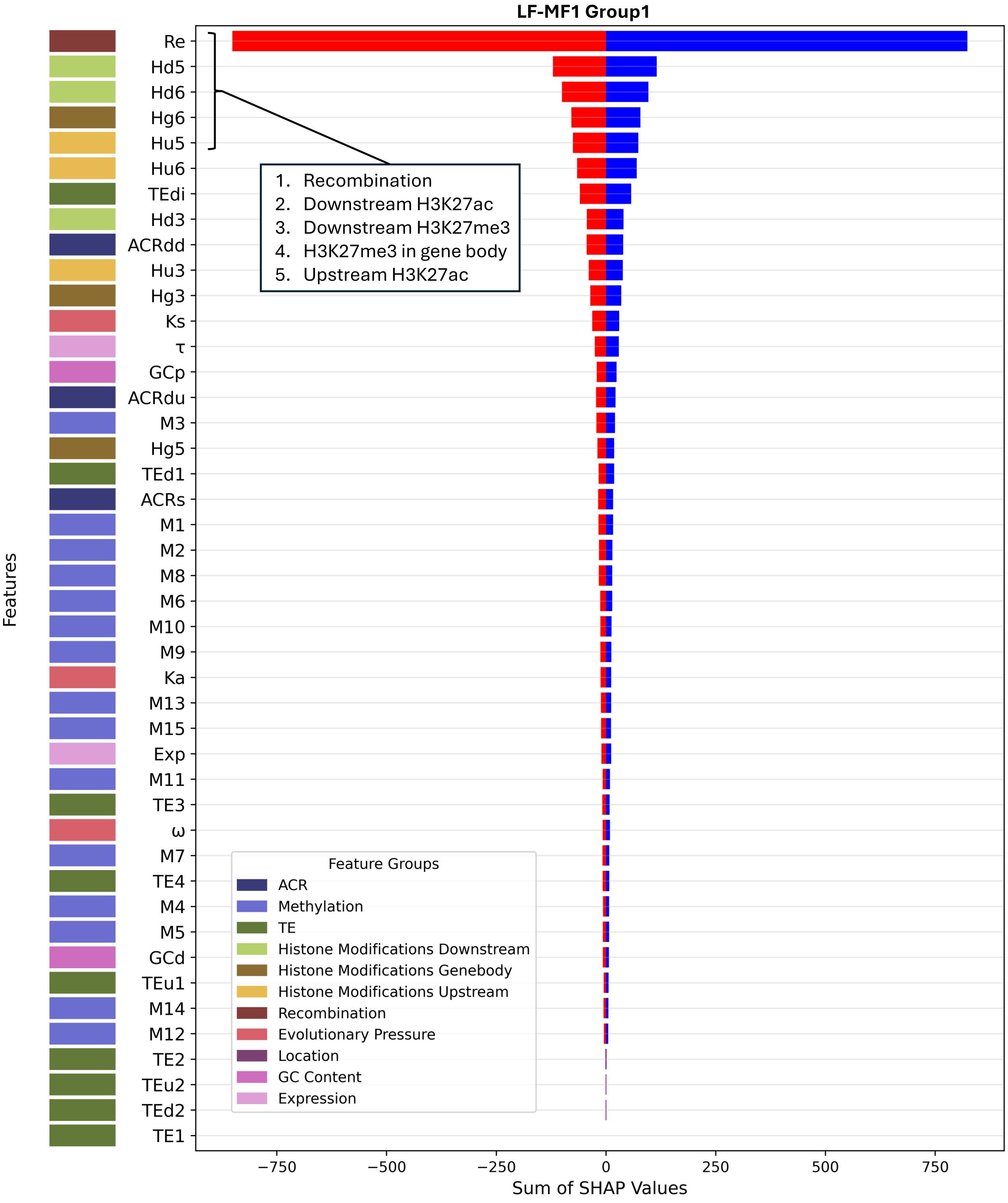


**Fig. S62. Directional SHAP feature importance profile for *B. rapa* LF-MF1 group1.** Features are ranked by the sum of their absolute SHAP values across all test-fold samples (ascending order, highest-importance features at top). For each feature, SHAP values are summed separately across positive contributions (blue; favoring MF1 classification) and negative contributions (red; favoring LF classification), reflecting total directional influence across the test set rather than per-sample magnitude. Colored tiles in the left margin indicate feature group membership. The five features contributing most to classification, as determined by summed absolute SHAP value, are listed in the inset.

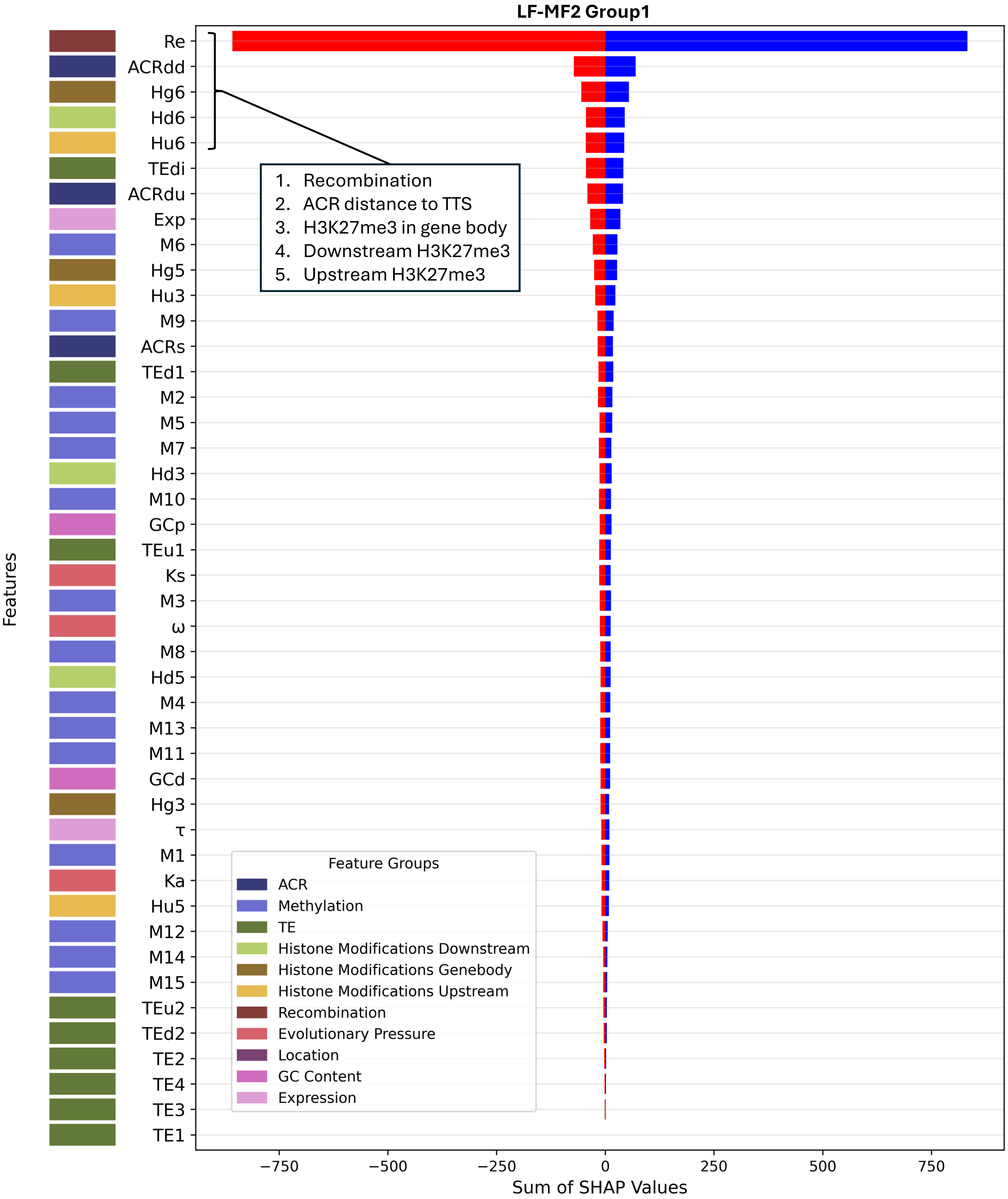


**Fig. S63. Directional SHAP feature importance profile for *B. rapa* LF-MF2 group1.** Features are ranked by the sum of their absolute SHAP values across all test-fold samples (ascending order, highest-importance features at top). For each feature, SHAP values are summed separately across positive contributions (blue; favoring MF2 classification) and negative contributions (red; favoring LF classification), reflecting total directional influence across the test set rather than per-sample magnitude. Colored tiles in the left margin indicate feature group membership. The five features contributing most to classification, as determined by summed absolute SHAP value, are listed in the inset.

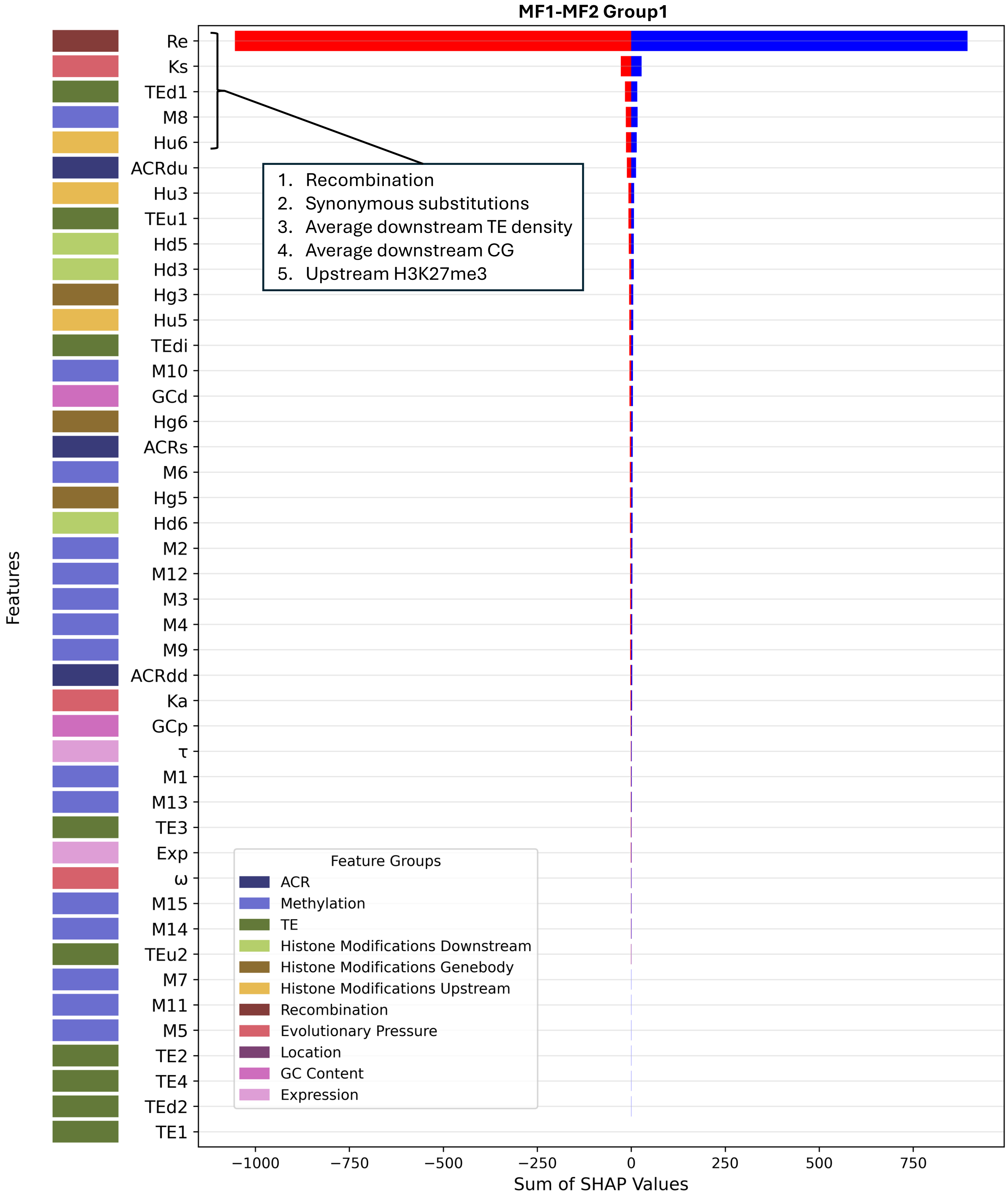


**Fig. S64. Directional SHAP feature importance profile for *B. rapa* MF1-MF2 group1.** Features are ranked by the sum of their absolute SHAP values across all test-fold samples (ascending order, highest-importance features at top). For each feature, SHAP values are summed separately across positive contributions (blue; favoring MF2 classification) and negative contributions (red; favoring MF1 classification), reflecting total directional influence across the test set rather than per-sample magnitude. Colored tiles in the left margin indicate feature group membership. The five features contributing most to classification, as determined by summed absolute SHAP value, are listed in the inset.

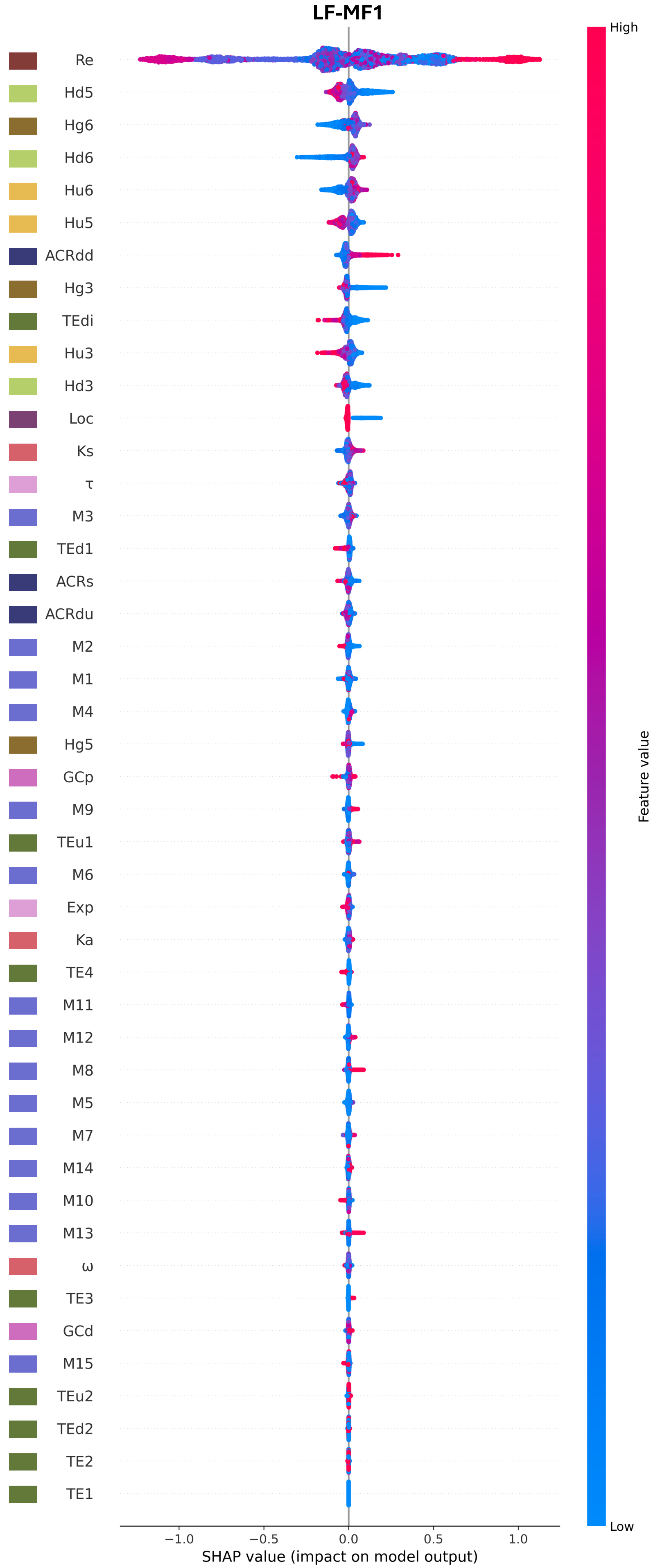


| Re | Recombination |
|---|---|

| | Expression Metrics |
|---|---|
| Exp | Avg expression |
| τ | Tau (tissue specificity index) |

| | GC Content |
|---|---|
| GCd | GC content cDNA |
| GCp | GC content promoter |

| | Methylation |
|---|---|
| M1 | CHH upstream avg |
| M2 | CHH downstream avg |
| M3 | CHH gene body avg |
| M4 | CHG upstream avg |
| M5 | CHG downstream avg |
| M6 | CHG gene body avg |
| M7 | CG upstream avg |
| M8 | CG downstream avg |
| M9 | CG gene body avg |
| M10 | CHH upstream max |
| M11 | CHH downstream max |
| M12 | CHG upstream max |
| M13 | CHG downstream max |
| M14 | CG upstream max |
| M15 | CG downstream max |

| | ACRs |
|---|---|
| ACRdu | ACR distance to TSS |
| ACRdd | ACR distance to TTS |
| ACRs | Nearest summit |

| | Histone Mod. Upstream |
|---|---|
| Hu3 | H3K4me3 up |
| Hu5 | H3K27ac up |
| Hu6 | H3K27me3 up |

| | Histone Mod. Gene Body |
|---|---|
| Hg3 | H3K4me3 gene body |
| Hg5 | H3K27ac gene body |
| Hg6 | H3K27me3 gene body |

| | Histone Mod. Downstream |
|---|---|
| Hd3 | H3K4me3 down |
| Hd5 | H3K27ac down |
| Hd6 | H3K27me3 down |

| | Transposable Elements (TEs) |
|---|---|
| TEu1 | Avg upstream density |
| TEd1 | Avg downstream density |
| TEu2 | Max upstream density |
| TEd2 | Max downstream density |
| TE1 | Helitrons |
| TE2 | Terminal inverted repeat elements |
| TE3 | LTR retrotransposons |
| TE4 | LINE and SINE elements |
| TEdi | TE distance |

| | Evolutionary Pressure |
|---|---|
| Ka | Nonsynonymous substitutions |
| Ks | Synonymous substitutions |
| ω | Omega, Ka/Ks ratio |

**Fig. S65. SHAP summary beeswarm plot for *B. rapa* LF-MF1.** Features are ranked by mean absolute SHAP value (highest at top). Each point represents one test-fold sample, with horizontal position indicating the SHAP value (impact on model output); points are jittered vertically for visibility where they overlap. Point color encodes the raw feature value for that sample, ranging from low (blue) to high (red). Positive SHAP values indicate a contribution toward MF1 classification; negative SHAP values indicate a contribution toward LF classification. SHAP values were computed on held-out test-fold samples across 10 stratified cross-validation folds.

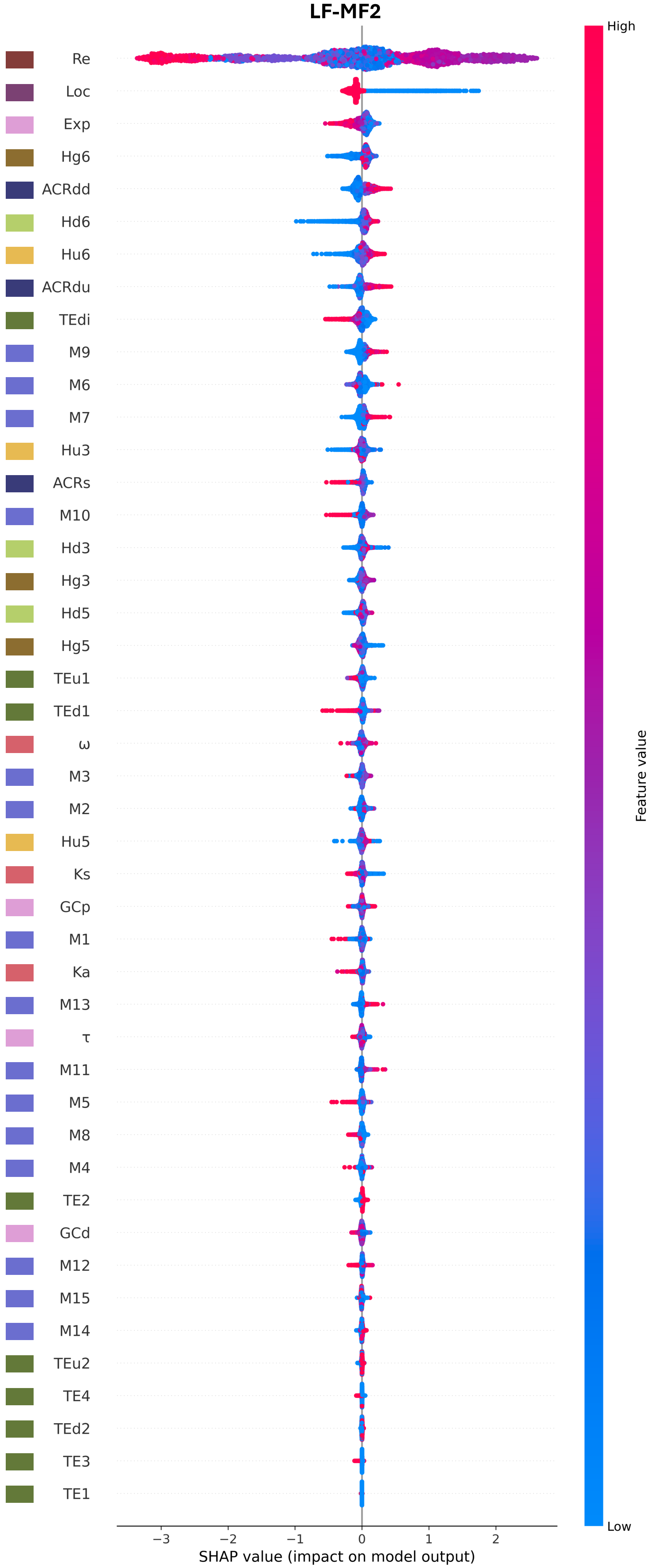


| Re | Recombination |
|---|---|

| | Expression Metrics |
|---|---|
| Exp | Avg expression |
| τ | Tau (tissue specificity index) |

| | GC Content |
|---|---|
| GCd | GC content cDNA |
| GCp | GC content promoter |

| | Methylation |
|---|---|
| M1 | CHH upstream avg |
| M2 | CHH downstream avg |
| M3 | CHH gene body avg |
| M4 | CHG upstream avg |
| M5 | CHG downstream avg |
| M6 | CHG gene body avg |
| M7 | CG upstream avg |
| M8 | CG downstream avg |
| M9 | CG gene body avg |
| M10 | CHH upstream max |
| M11 | CHH downstream max |
| M12 | CHG upstream max |
| M13 | CHG downstream max |
| M14 | CG upstream max |
| M15 | CG downstream max |

| | ACRs |
|---|---|
| ACRdu | ACR distance to TSS |
| ACRdd | ACR distance to TTS |
| ACRs | Nearest summit |

| | Histone Mod. Upstream |
|---|---|
| Hu3 | H3K4me3 up |
| Hu5 | H3K27ac up |
| Hu6 | H3K27me3 up |

| | Histone Mod. Gene Body |
|---|---|
| Hg3 | H3K4me3 gene body |
| Hg5 | H3K27ac gene body |
| Hg6 | H3K27me3 gene body |

| | Histone Mod. Downstream |
|---|---|
| Hd3 | H3K4me3 down |
| Hd5 | H3K27ac down |
| Hd6 | H3K27me3 down |

| | Transposable Elements (TEs) |
|---|---|
| TEu1 | Avg upstream density |
| TEd1 | Avg downstream density |
| TEu2 | Max upstream density |
| TEd2 | Max downstream density |
| TE1 | Helitrons |
| TE2 | Terminal inverted repeat elements |
| TE3 | LTR retrotransposons |
| TE4 | LINE and SINE elements |
| TEdi | TE distance |

| | Evolutionary Pressure |
|---|---|
| Ka | Nonsynonymous substitutions |
| Ks | Synonymous substitutions |
| ω | Omega, Ka/Ks ratio |

**Fig. S66. SHAP summary beeswarm plot for *B. rapa* LF-MF2.** Features are ranked by mean absolute SHAP value (highest at top). Each point represents one test-fold sample, with horizontal position indicating the SHAP value (impact on model output); points are jittered vertically for visibility where they overlap. Point color encodes the raw feature value for that sample, ranging from low (blue) to high (red). Positive SHAP values indicate a contribution toward MF2 classification; negative SHAP values indicate a contribution toward LF classification. SHAP values were computed on held-out test-fold samples across 10 stratified cross-validation folds.

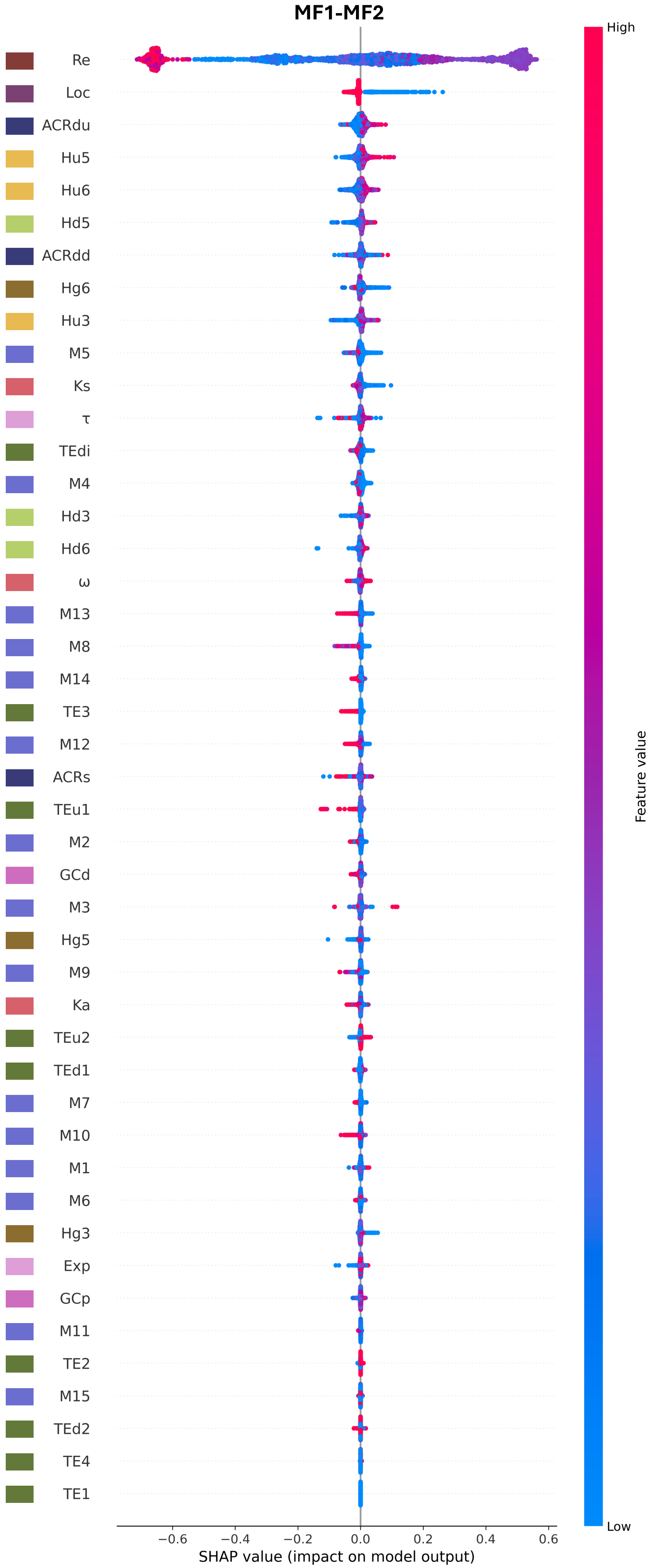


| Re | Recombination |
|---|---|

| | Expression Metrics |
|---|---|
| Exp | Avg expression |
| τ | Tau (tissue specificity index) |

| | GC Content |
|---|---|
| GCd | GC content cDNA |
| GCp | GC content promoter |

| | Methylation |
|---|---|
| M1 | CHH upstream avg |
| M2 | CHH downstream avg |
| M3 | CHH gene body avg |
| M4 | CHG upstream avg |
| M5 | CHG downstream avg |
| M6 | CHG gene body avg |
| M7 | CG upstream avg |
| M8 | CG downstream avg |
| M9 | CG gene body avg |
| M10 | CHH upstream max |
| M11 | CHH downstream max |
| M12 | CHG upstream max |
| M13 | CHG downstream max |
| M14 | CG upstream max |
| M15 | CG downstream max |

| | ACRs |
|---|---|
| ACRdu | ACR distance to TSS |
| ACRdd | ACR distance to TTS |
| ACRs | Nearest summit |

| | Histone Mod. Upstream |
|---|---|
| Hu3 | H3K4me3 up |
| Hu5 | H3K27ac up |
| Hu6 | H3K27me3 up |

| | Histone Mod. Gene Body |
|---|---|
| Hg3 | H3K4me3 gene body |
| Hg5 | H3K27ac gene body |
| Hg6 | H3K27me3 gene body |

| | Histone Mod. Downstream |
|---|---|
| Hd3 | H3K4me3 down |
| Hd5 | H3K27ac down |
| Hd6 | H3K27me3 down |

| | Transposable Elements (TEs) |
|---|---|
| TEu1 | Avg upstream density |
| TEd1 | Avg downstream density |
| TEu2 | Max upstream density |
| TEd2 | Max downstream density |
| TE1 | Helitrons |
| TE2 | Terminal inverted repeat elements |
| TE3 | LTR retrotransposons |
| TE4 | LINE and SINE elements |
| TEdi | TE distance |

| | Evolutionary Pressure |
|---|---|
| Ka | Nonsynonymous substitutions |
| Ks | Synonymous substitutions |
| ω | Omega, Ka/Ks ratio |

**Fig. S67. SHAP summary beeswarm plot for *B. rapa* MF1-MF2.** Features are ranked by mean absolute SHAP value (highest at top). Each point represents one test-fold sample, with horizontal position indicating the SHAP value (impact on model output); points are jittered vertically for visibility where they overlap. Point color encodes the raw feature value for that sample, ranging from low (blue) to high (red). Positive SHAP values indicate a contribution toward MF2 classification; negative SHAP values indicate a contribution toward MF1 classification. SHAP values were computed on held-out test-fold samples across 10 stratified cross-validation folds.

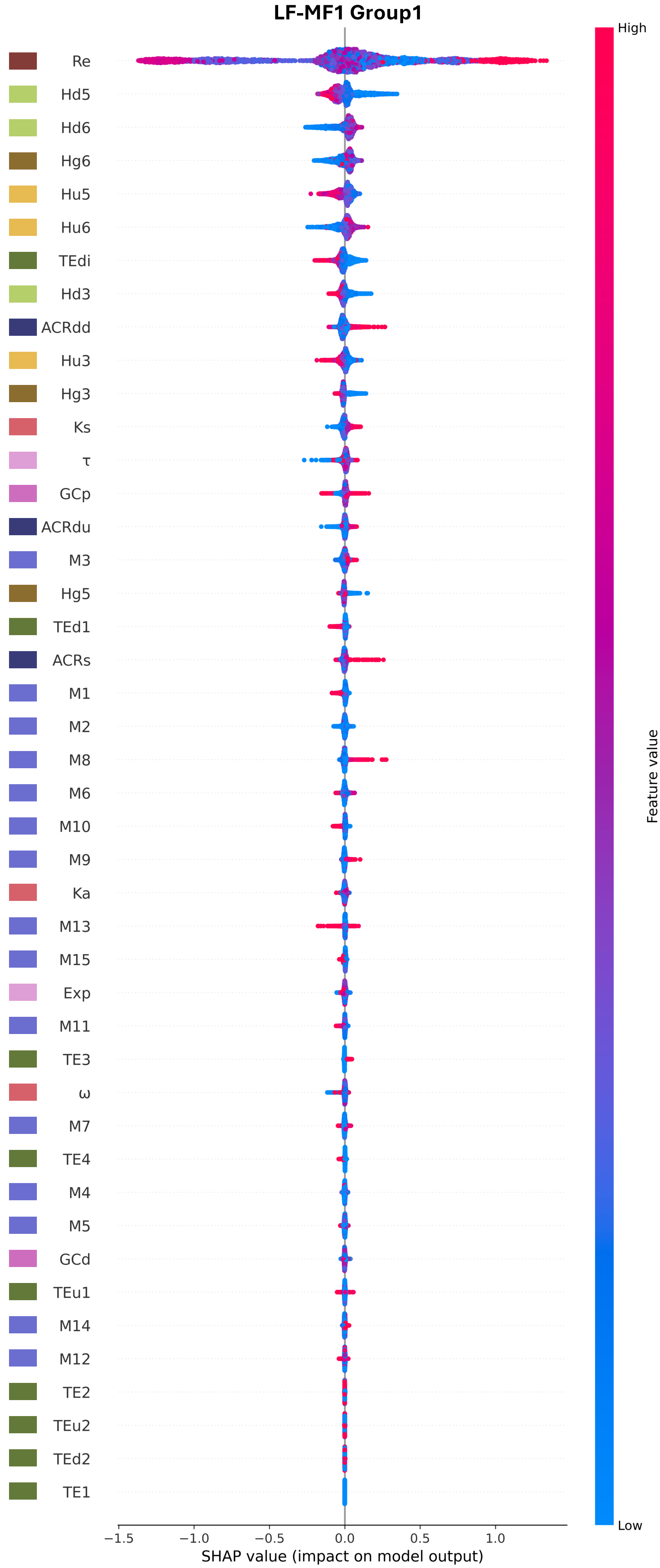


| Re | Recombination |
|---|---|

| | Expression Metrics |
|---|---|
| Exp | Avg expression |
| τ | Tau (tissue specificity index) |

| | GC Content |
|---|---|
| GCd | GC content cDNA |
| GCp | GC content promoter |

| | Methylation |
|---|---|
| M1 | CHH upstream avg |
| M2 | CHH downstream avg |
| M3 | CHH gene body avg |
| M4 | CHG upstream avg |
| M5 | CHG downstream avg |
| M6 | CHG gene body avg |
| M7 | CG upstream avg |
| M8 | CG downstream avg |
| M9 | CG gene body avg |
| M10 | CHH upstream max |
| M11 | CHH downstream max |
| M12 | CHG upstream max |
| M13 | CHG downstream max |
| M14 | CG upstream max |
| M15 | CG downstream max |

| | ACRs |
|---|---|
| ACRdu | ACR distance to TSS |
| ACRdd | ACR distance to TTS |
| ACRs | Nearest summit |

| | Histone Mod. Upstream |
|---|---|
| Hu3 | H3K4me3 up |
| Hu5 | H3K27ac up |
| Hu6 | H3K27me3 up |

| | Histone Mod. Gene Body |
|---|---|
| Hg3 | H3K4me3 gene body |
| Hg5 | H3K27ac gene body |
| Hg6 | H3K27me3 gene body |

| | Histone Mod. Downstream |
|---|---|
| Hd3 | H3K4me3 down |
| Hd5 | H3K27ac down |
| Hd6 | H3K27me3 down |

| | Transposable Elements (TEs) |
|---|---|
| TEu1 | Avg upstream density |
| TEd1 | Avg downstream density |
| TEu2 | Max upstream density |
| TEd2 | Max downstream density |
| TE1 | Helitrons |
| TE2 | Terminal inverted repeat elements |
| TE3 | LTR retrotransposons |
| TE4 | LINE and SINE elements |
| TEdi | TE distance |

| | Evolutionary Pressure |
|---|---|
| Ka | Nonsynonymous substitutions |
| Ks | Synonymous substitutions |
| ω | Omega, Ka/Ks ratio |

**Fig. S68. SHAP summary beeswarm plot for *B. rapa* LF-MF1 group1.** Features are ranked by mean absolute SHAP value (highest at top). Each point represents one test-fold sample, with horizontal position indicating the SHAP value (impact on model output); points are jittered vertically for visibility where they overlap. Point color encodes the raw feature value for that sample, ranging from low (blue) to high (red). Positive SHAP values indicate a contribution toward MF1 classification; negative SHAP values indicate a contribution toward LF classification. SHAP values were computed on held-out test-fold samples across 10 stratified cross-validation folds.

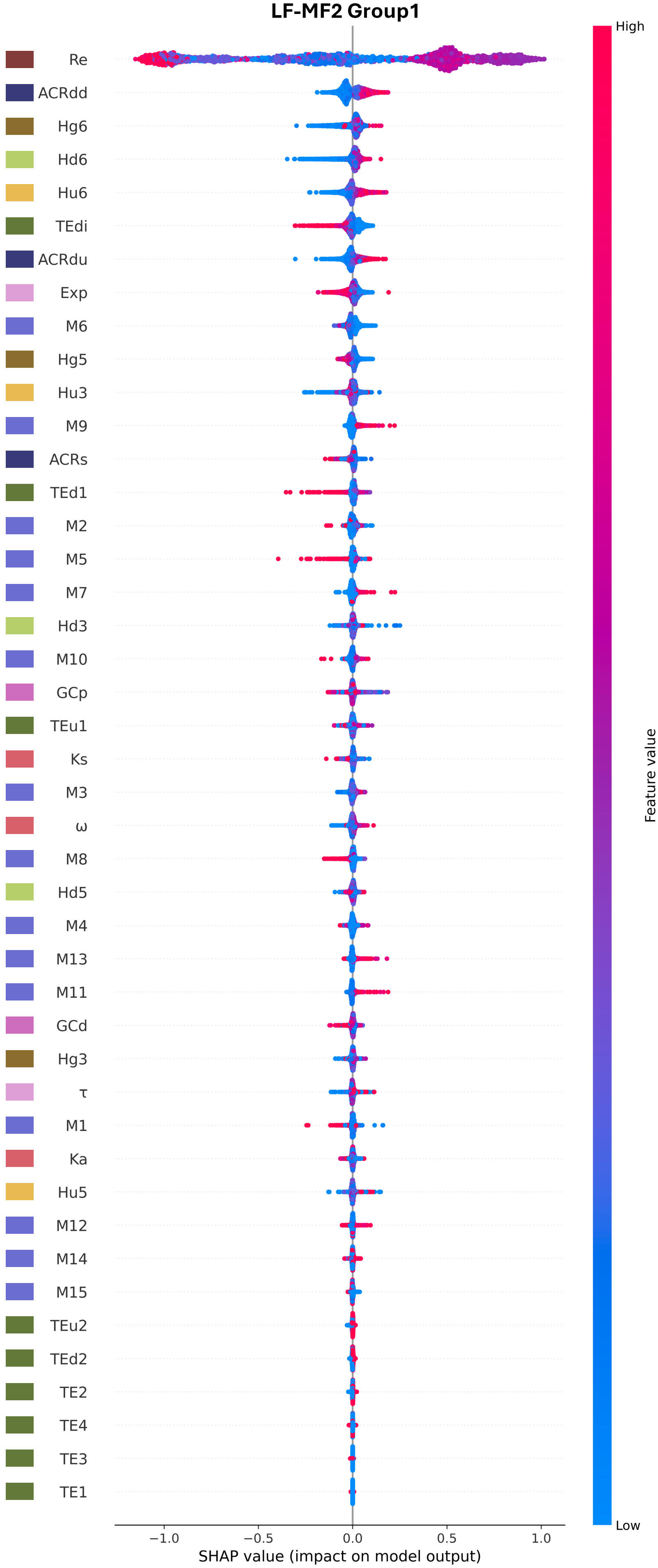


| Re | Recombination |
|---|---|

| | Expression Metrics |
|---|---|
| Exp | Avg expression |
| τ | Tau (tissue specificity index) |

| | GC Content |
|---|---|
| GCd | GC content cDNA |
| GCp | GC content promoter |

| | Methylation |
|---|---|
| M1 | CHH upstream avg |
| M2 | CHH downstream avg |
| M3 | CHH gene body avg |
| M4 | CHG upstream avg |
| M5 | CHG downstream avg |
| M6 | CHG gene body avg |
| M7 | CG upstream avg |
| M8 | CG downstream avg |
| M9 | CG gene body avg |
| M10 | CHH upstream max |
| M11 | CHH downstream max |
| M12 | CHG upstream max |
| M13 | CHG downstream max |
| M14 | CG upstream max |
| M15 | CG downstream max |

| | ACRs |
|---|---|
| ACRdu | ACR distance to TSS |
| ACRdd | ACR distance to TTS |
| ACRs | Nearest summit |

| | Histone Mod. Upstream |
|---|---|
| Hu3 | H3K4me3 up |
| Hu5 | H3K27ac up |
| Hu6 | H3K27me3 up |

| | Histone Mod. Gene Body |
|---|---|
| Hg3 | H3K4me3 gene body |
| Hg5 | H3K27ac gene body |
| Hg6 | H3K27me3 gene body |

| | Histone Mod. Downstream |
|---|---|
| Hd3 | H3K4me3 down |
| Hd5 | H3K27ac down |
| Hd6 | H3K27me3 down |

| | Transposable Elements (TEs) |
|---|---|
| TEu1 | Avg upstream density |
| TEd1 | Avg downstream density |
| TEu2 | Max upstream density |
| TEd2 | Max downstream density |
| TE1 | Helitrons |
| TE2 | Terminal inverted repeat elements |
| TE3 | LTR retrotransposons |
| TE4 | LINE and SINE elements |
| TEdi | TE distance |

| | Evolutionary Pressure |
|---|---|
| Ka | Nonsynonymous substitutions |
| Ks | Synonymous substitutions |
| ω | Omega, Ka/Ks ratio |

**Fig. S69. SHAP summary beeswarm plot for *B. rapa* LF-MF2 group1.** Features are ranked by mean absolute SHAP value (highest at top). Each point represents one test-fold sample, with horizontal position indicating the SHAP value (impact on model output); points are jittered vertically for visibility where they overlap. Point color encodes the raw feature value for that sample, ranging from low (blue) to high (red). Positive SHAP values indicate a contribution toward MF2 classification; negative SHAP values indicate a contribution toward LF classification. SHAP values were computed on held-out test-fold samples across 10 stratified cross-validation folds.

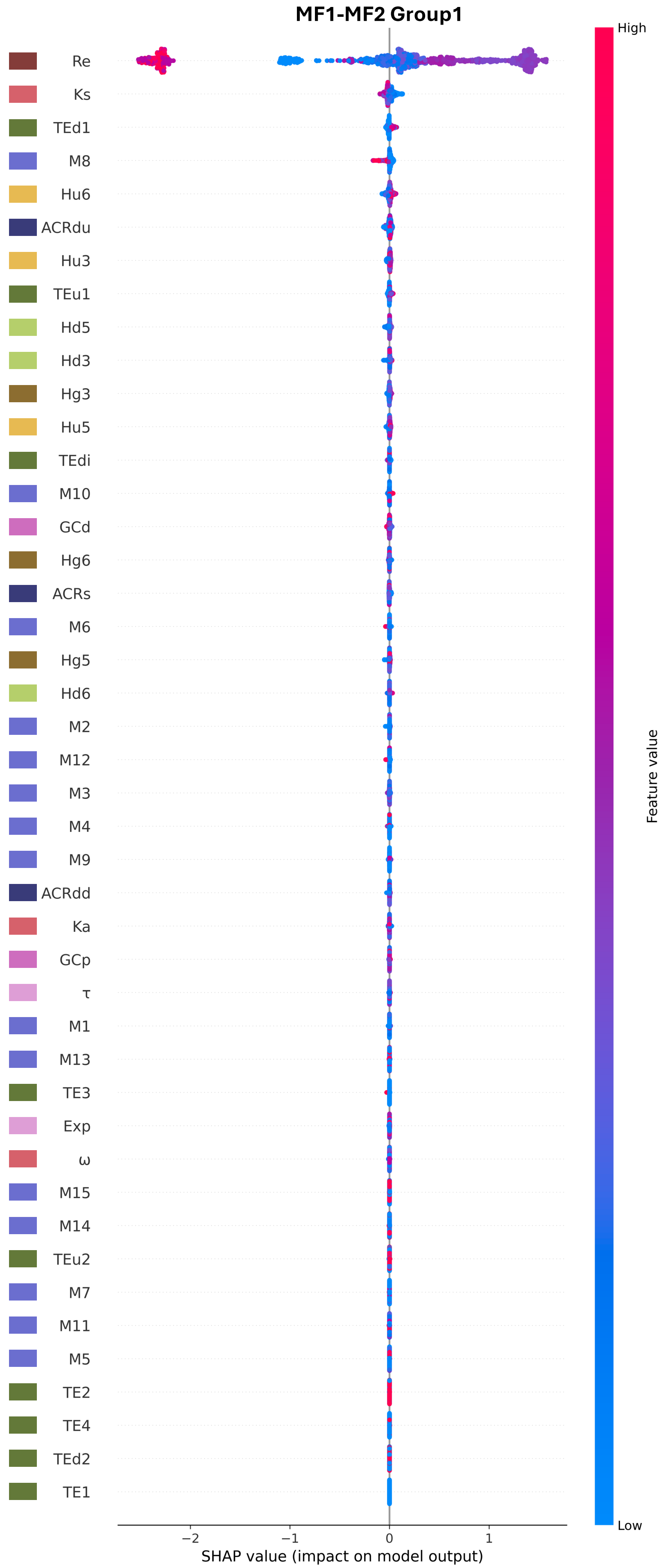


| Re | Recombination |
|---|---|

| | Expression Metrics |
|---|---|
| Exp | Avg expression |
| τ | Tau (tissue specificity index) |

| | GC Content |
|---|---|
| GCd | GC content cDNA |
| GCp | GC content promoter |

| | Methylation |
|---|---|
| M1 | CHH upstream avg |
| M2 | CHH downstream avg |
| M3 | CHH gene body avg |
| M4 | CHG upstream avg |
| M5 | CHG downstream avg |
| M6 | CHG gene body avg |
| M7 | CG upstream avg |
| M8 | CG downstream avg |
| M9 | CG gene body avg |
| M10 | CHH upstream max |
| M11 | CHH downstream max |
| M12 | CHG upstream max |
| M13 | CHG downstream max |
| M14 | CG upstream max |
| M15 | CG downstream max |

| | ACRs |
|---|---|
| ACRdu | ACR distance to TSS |
| ACRdd | ACR distance to TTS |
| ACRs | Nearest summit |

| | Histone Mod. Upstream |
|---|---|
| Hu3 | H3K4me3 up |
| Hu5 | H3K27ac up |
| Hu6 | H3K27me3 up |

| | Histone Mod. Gene Body |
|---|---|
| Hg3 | H3K4me3 gene body |
| Hg5 | H3K27ac gene body |
| Hg6 | H3K27me3 gene body |

| | Histone Mod. Downstream |
|---|---|
| Hd3 | H3K4me3 down |
| Hd5 | H3K27ac down |
| Hd6 | H3K27me3 down |

| | Transposable Elements (TEs) |
|---|---|
| TEu1 | Avg upstream density |
| TEd1 | Avg downstream density |
| TEu2 | Max upstream density |
| TEd2 | Max downstream density |
| TE1 | Helitrons |
| TE2 | Terminal inverted repeat elements |
| TE3 | LTR retrotransposons |
| TE4 | LINE and SINE elements |
| TEdi | TE distance |

| | Evolutionary Pressure |
|---|---|
| Ka | Nonsynonymous substitutions |
| Ks | Synonymous substitutions |
| ω | Omega, Ka/Ks ratio |

**Fig. S70. SHAP summary beeswarm plot for *B. rapa* MF1-MF2 group1.** Features are ranked by mean absolute SHAP value (highest at top). Each point represents one test-fold sample, with horizontal position indicating the SHAP value (impact on model output); points are jittered vertically for visibility where they overlap. Point color encodes the raw feature value for that sample, ranging from low (blue) to high (red). Positive SHAP values indicate a contribution toward MF2 classification; negative SHAP values indicate a contribution toward MF1 classification. SHAP values were computed on held-out test-fold samples across 10 stratified cross-validation folds.

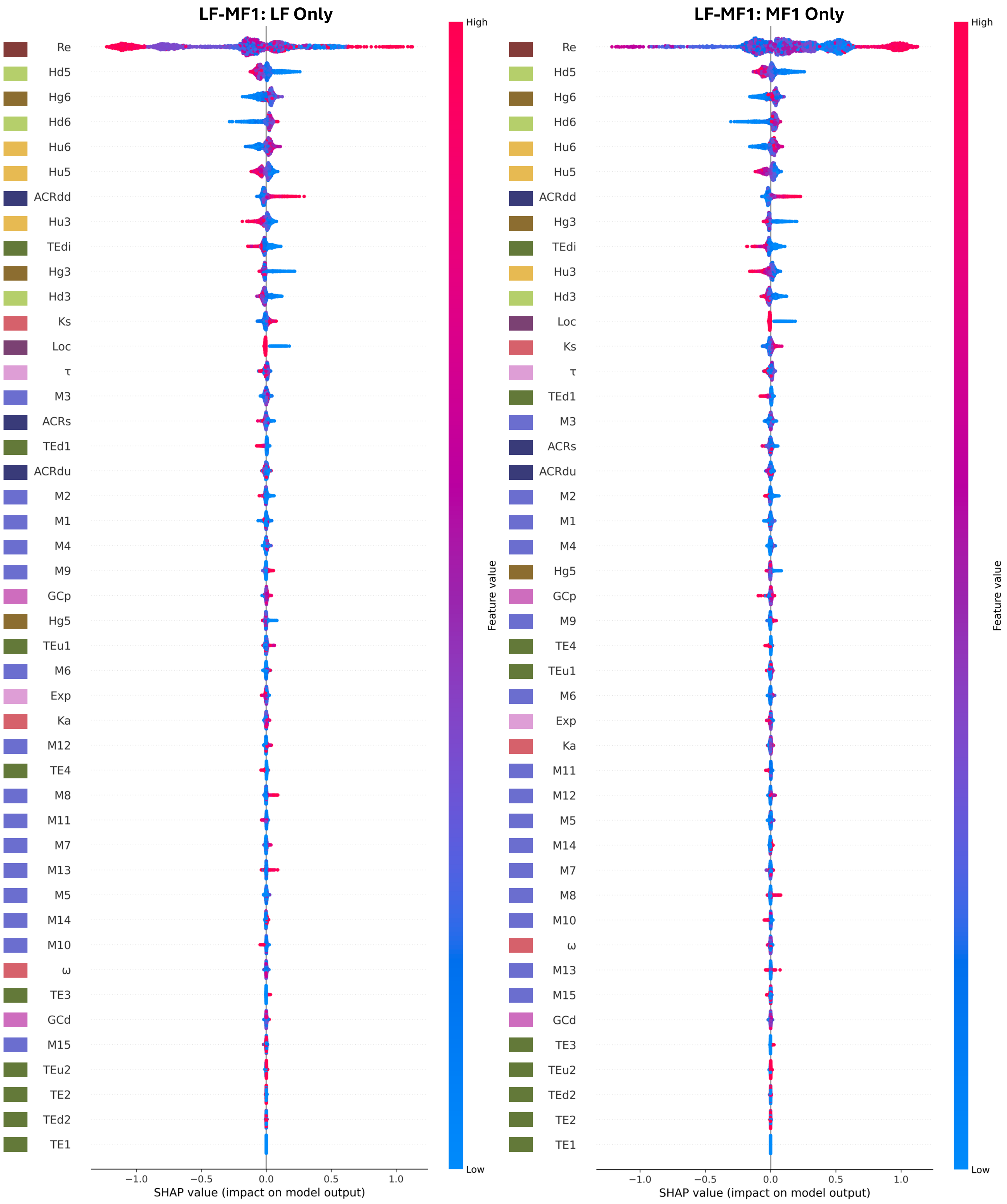


**Fig. S71. Per-label SHAP summary beeswarm plots for LF-MF1.**
(See pg 78)

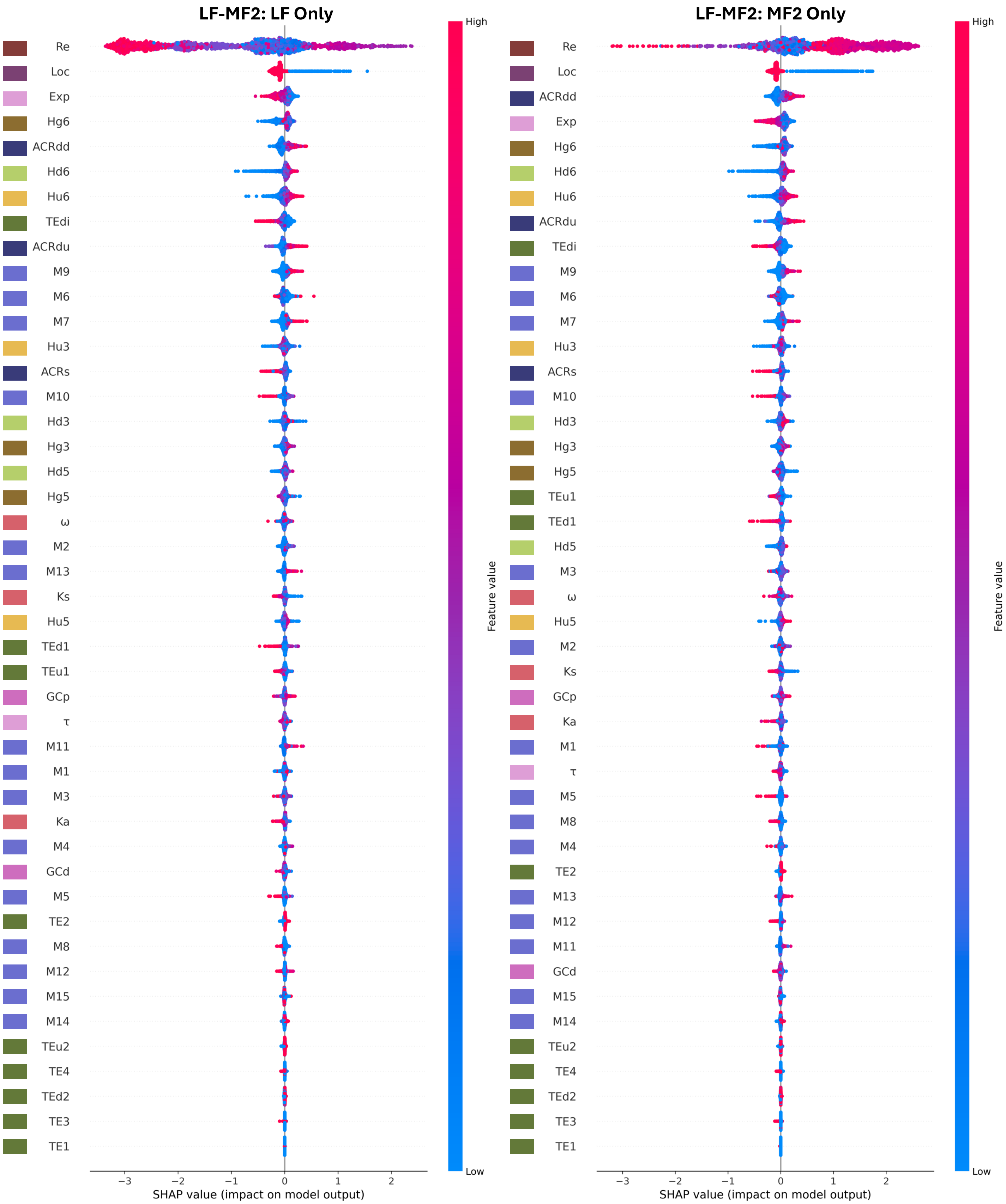


**Fig. S72. Per-label SHAP summary beeswarm plots for LF-MF2.**
(See pg 78)

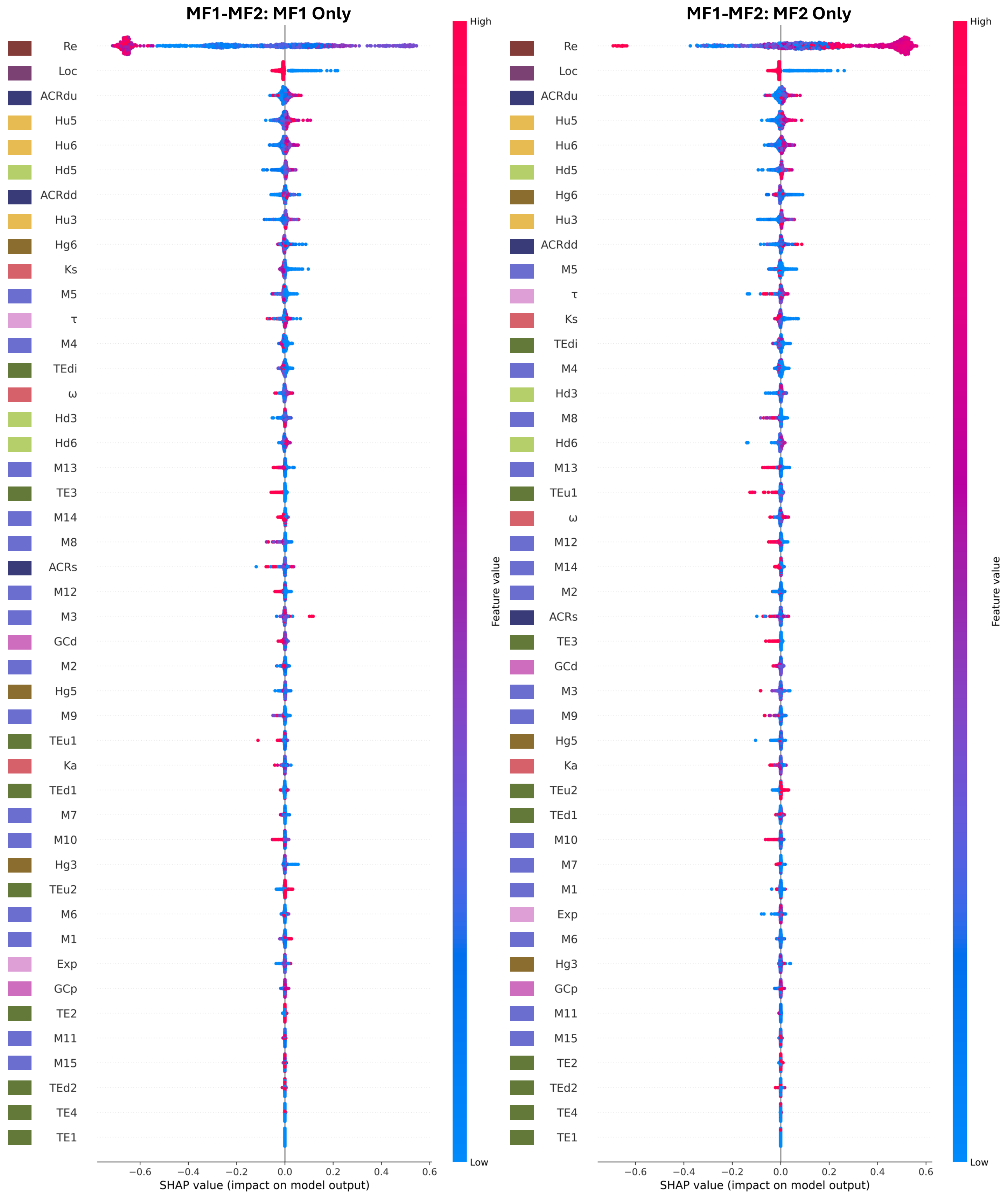


**Fig. S73. Per-label SHAP summary beeswarm plots for MF1-MF2.**
(See pg 78)

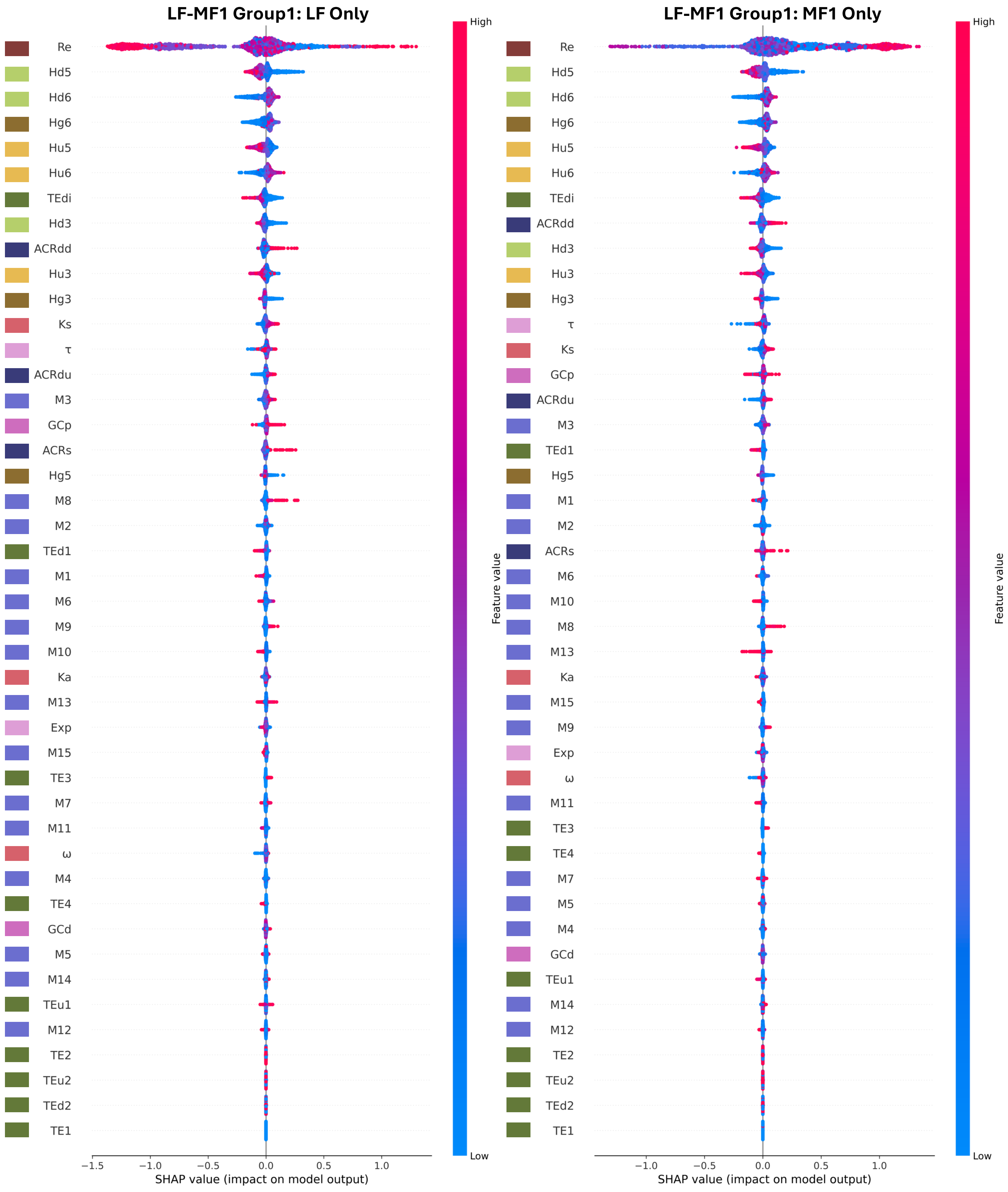


**Fig. S74. Per-label SHAP summary beeswarm plots for LF-MF1 group1.**
(See pg 78)

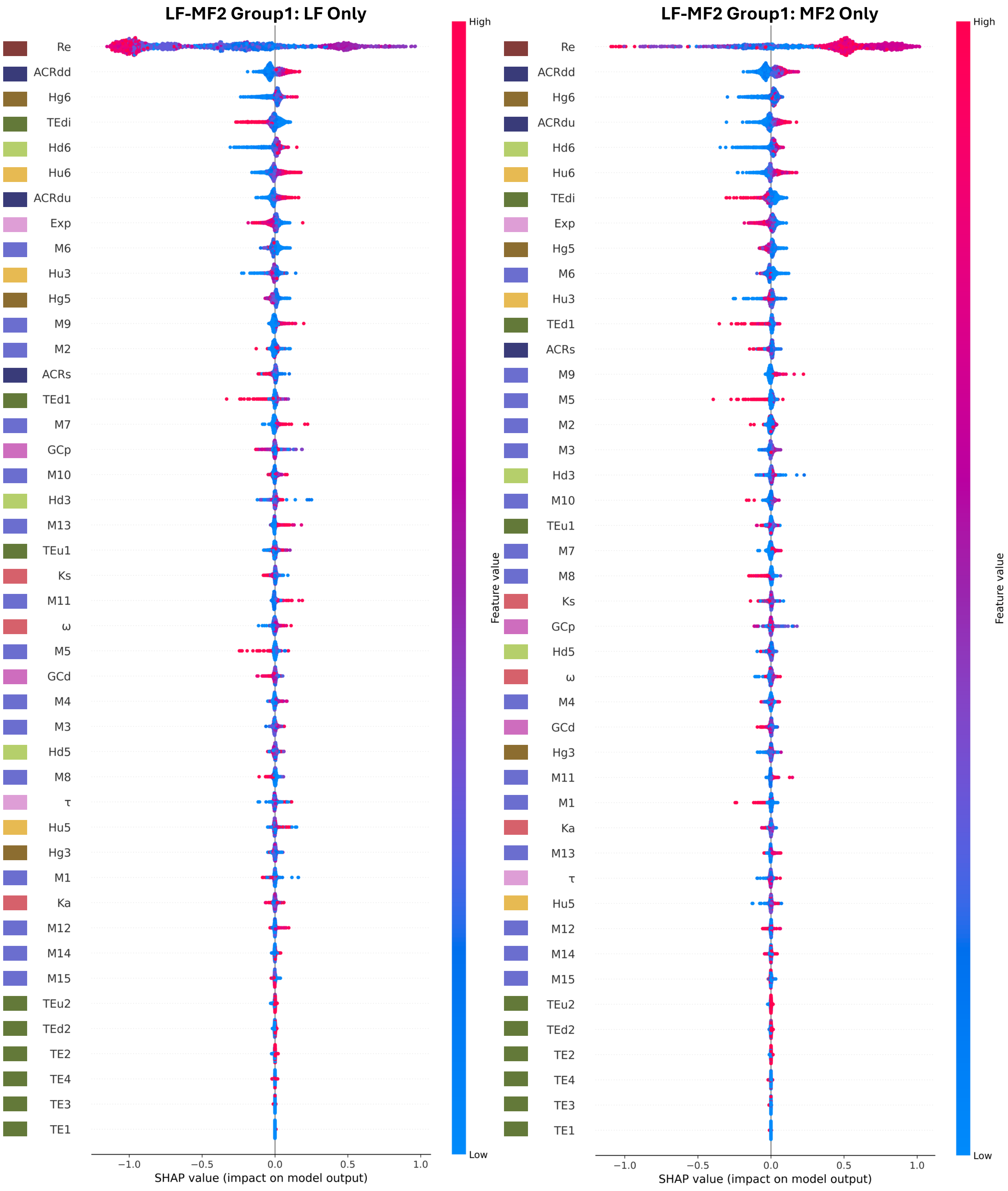


**Fig. S75. Per-label SHAP summary beeswarm plots for LF-MF2 group1.**
(See pg 78)

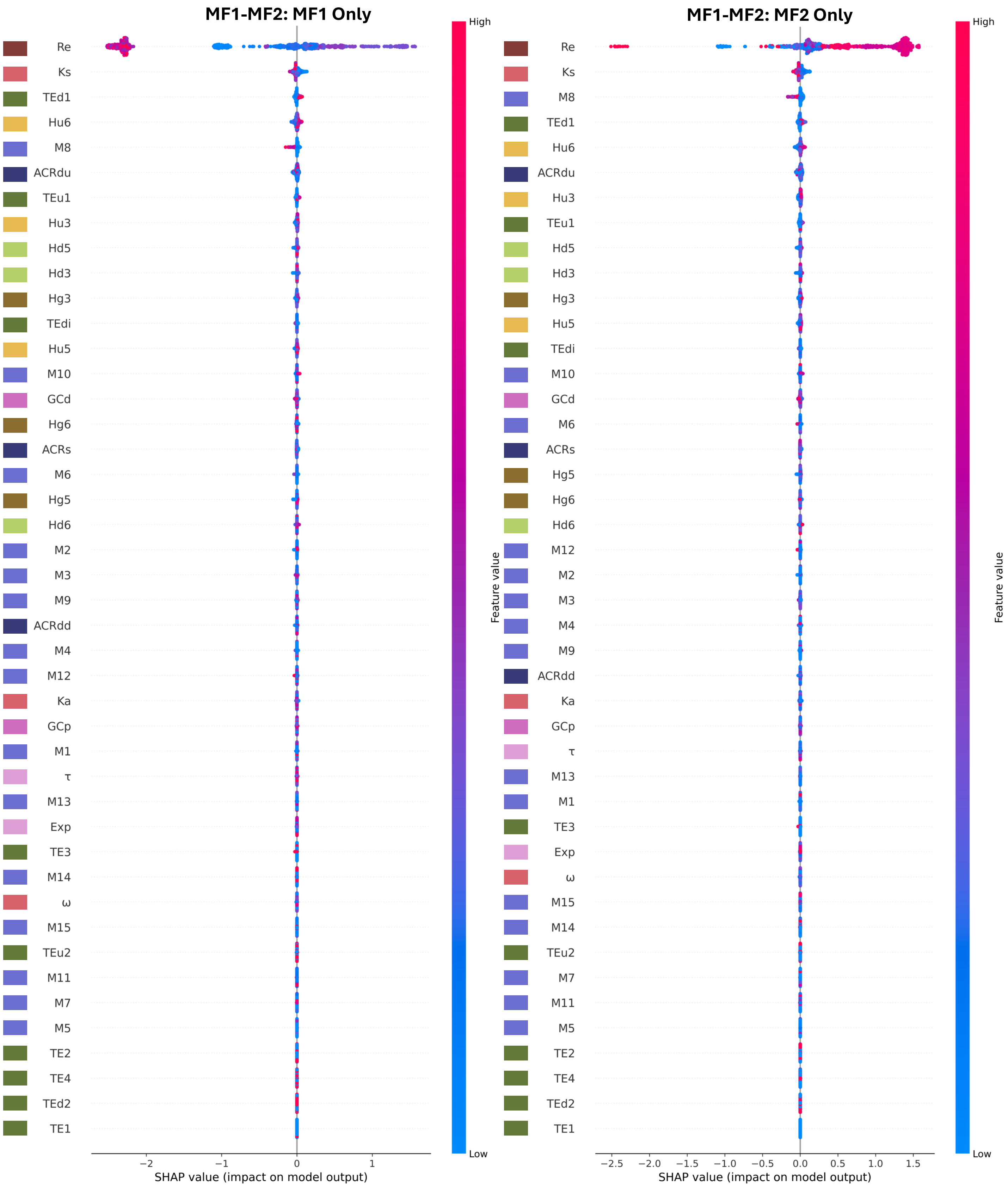


**Fig. S76. Per-label SHAP summary beeswarm plots for MF1-MF2 group1.**
(See pg 78)

**Per-label SHAP summary beeswarm plots for *B. rapa*.** SHAP values are shown separately for the more dominant subgenome (LF in LF-MF1, LF-MF2; left) and the less dominant subgenome (MF1 in LF-MF1, MF2 in LF-MF2, MF1-MF2; right) samples. Within each panel, features are ranked independently by mean absolute SHAP value for that label's samples (highest at top); rank order therefore differs between panels and reflects asymmetry in the features driving dominant versus submissive subgenome classification. Each point represents one test-fold sample, with horizontal position indicating the SHAP value (impact on model output) and vertical jitter applied where points overlap. Point color encodes the raw feature value for that sample, ranging from low (blue) to high (red); gray points indicate missing (NaN) feature values. Positive SHAP values indicate a contribution toward the more non-dominant subgenome classification; negative SHAP values indicate a contribution toward the more dominant classification, and this directionality is consistent across both panels. SHAP values were computed on held-out test-fold samples across 10 stratified cross-validation folds. For SHAP values computed across all samples regardless of label, see Figure S65-S70. For summed directional SHAP contributions per feature, see Figure S59-S64.

**Fig. S71. Per-label SHAP summary beeswarm plots for LF-MF1.**
**Fig. S72. Per-label SHAP summary beeswarm plots for LF-MF2.**
**Fig. S73. Per-label SHAP summary beeswarm plots for MF1-MF2.**
**Fig. S74. Per-label SHAP summary beeswarm plots for LF-MF1 group1.**
**Fig. S75. Per-label SHAP summary beeswarm plots for LF-MF2 group1.**
**Fig. S76. Per-label SHAP summary beeswarm plots for MF1-MF2 group1.**

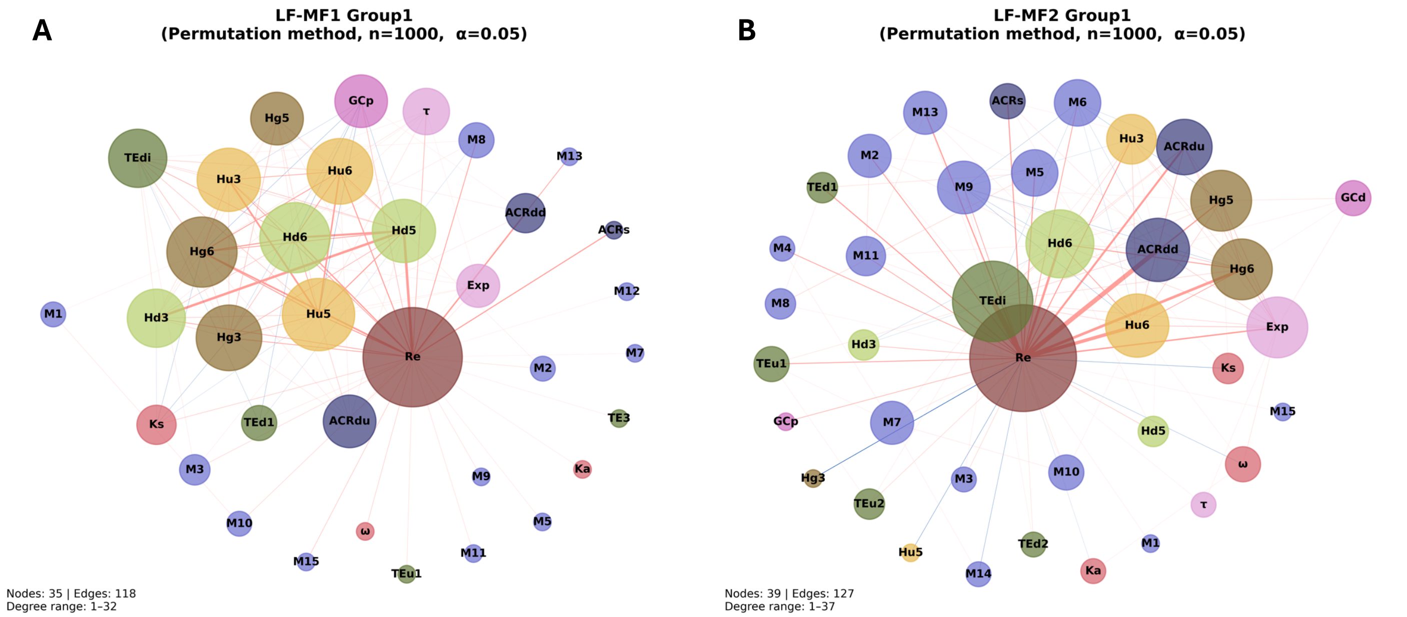


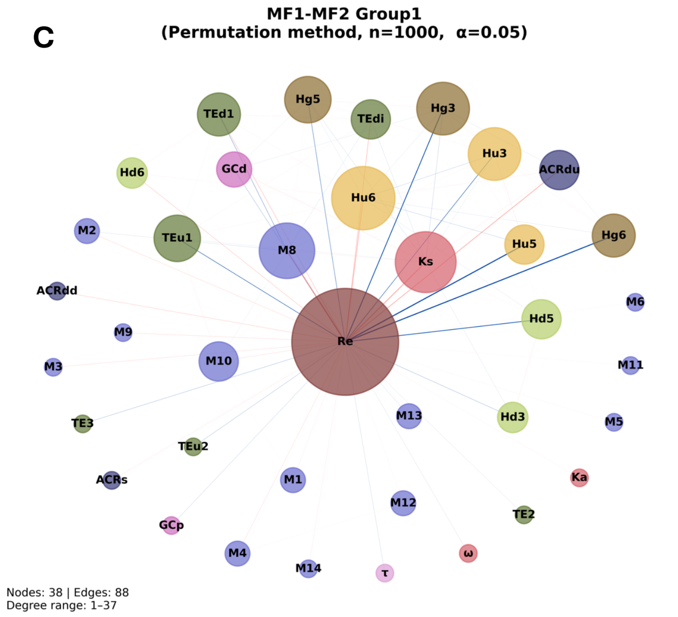


**D**

| | LF-MF1 Group1 | LF-MF2 Group1 | MF1-MF2 Group1 |
|---|---|---|---|
| 1 | Re | Re | Re |
| 2 | Hu5 | TEdi | Hu6 |
| 3 | Hd6 | Hd6 | Ks |
| 4 | Hg6 | ACRdd | M8 |
| 5 | Hg3 | Hu6 | Hg3 |

**Fig. S77. SHAP co-variation networks for *B. rapa* group1 subgenome classification.** Networks for LF-MF1 group1 (**A**), LF-MF2 group1 (**B**), and MF1-MF2 group1 (**C**). Nodes represent features, colored by feature group, and sized by degree (number of significant co-varying partners). Edges connect feature pairs with significant pairwise SHAP co-variation (permutation test, n = 1,000, α = 0.05), with color indicating direction: positive co-variation (features vary together in model importance; red) or negative co-variation (features vary in opposition; blue), and width scaled to reflect relative co-variation magnitude within each panel independently. Each panel reports the total node and edge count, degree range (minimum to maximum node degree in that network), and edge weight range (minimum to maximum co-variation score in that network). The top five features by node degree are listed for each network (**D**).

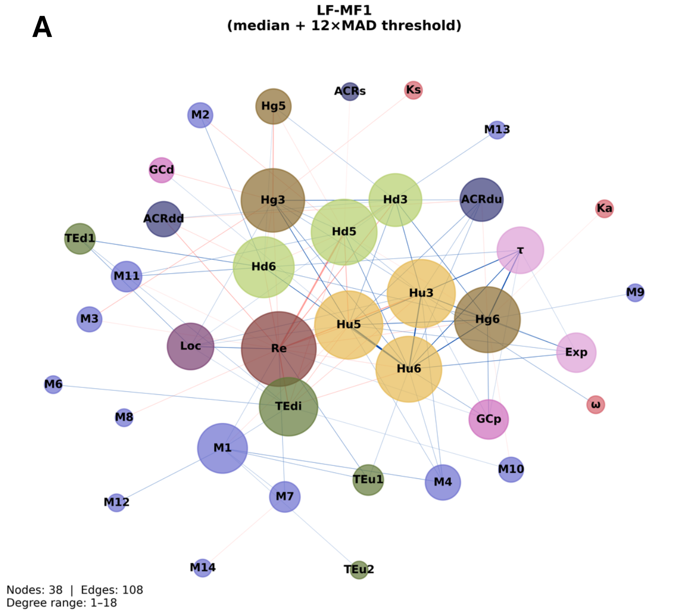


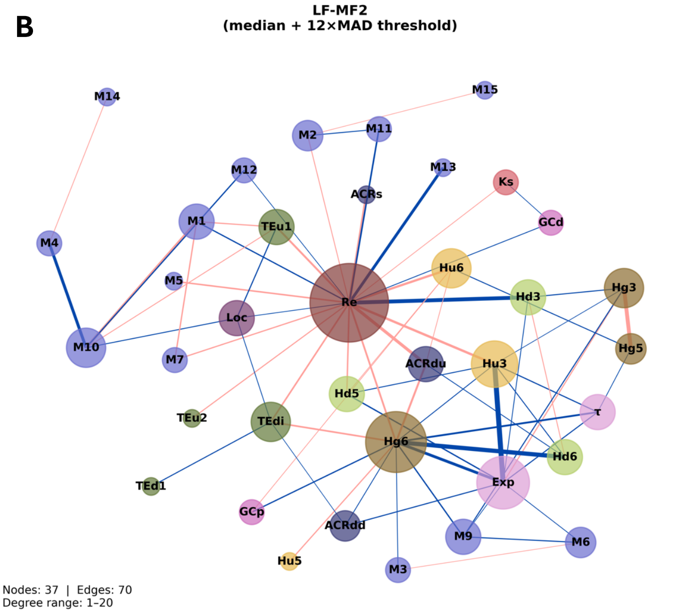


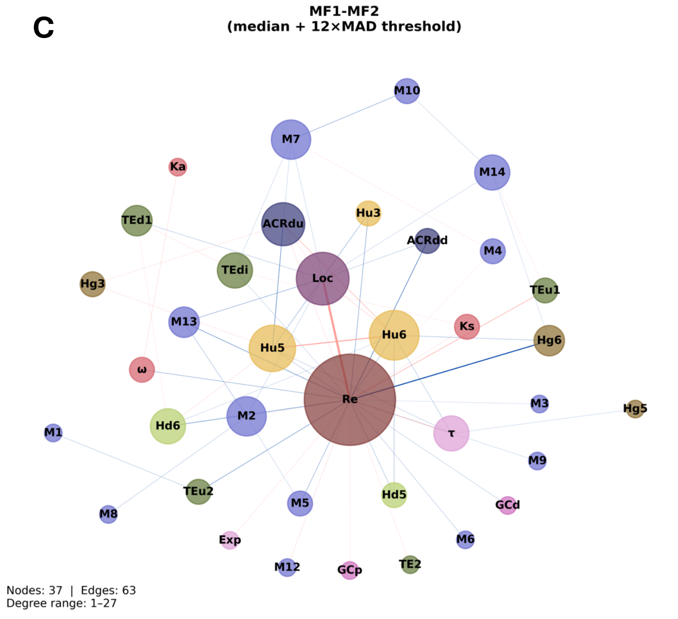


**Fig. S78. SHAP feature interaction networks for *B. rapa*.** Interaction networks shown for the LF-MF1 (**A**), LF-MF2 (**B**), MF1-MF2 (**C**) all-gene datasets. SHAP interaction values quantify the degree to which pairs of features jointly influence model predictions beyond what is expected from their individual contributions alone. For each feature pair, mean Shapley interaction values were computed across all test-set samples. Edges connect pairs whose absolute mean interaction value exceeded a robust effect size threshold (median + 12 × median absolute deviation, computed from non-zero pairs within each group) and are colored by interaction direction: positive (pink) edges indicate synergistic interactions, in which the combined contribution of two features exceeds the sum of their individual effects, and negative (blue) edges indicate antagonistic interactions, in which the combined contribution is less than the sum. Node size is proportional to degree (number of retained edges) and node color indicates feature category, as in Fig. 6.

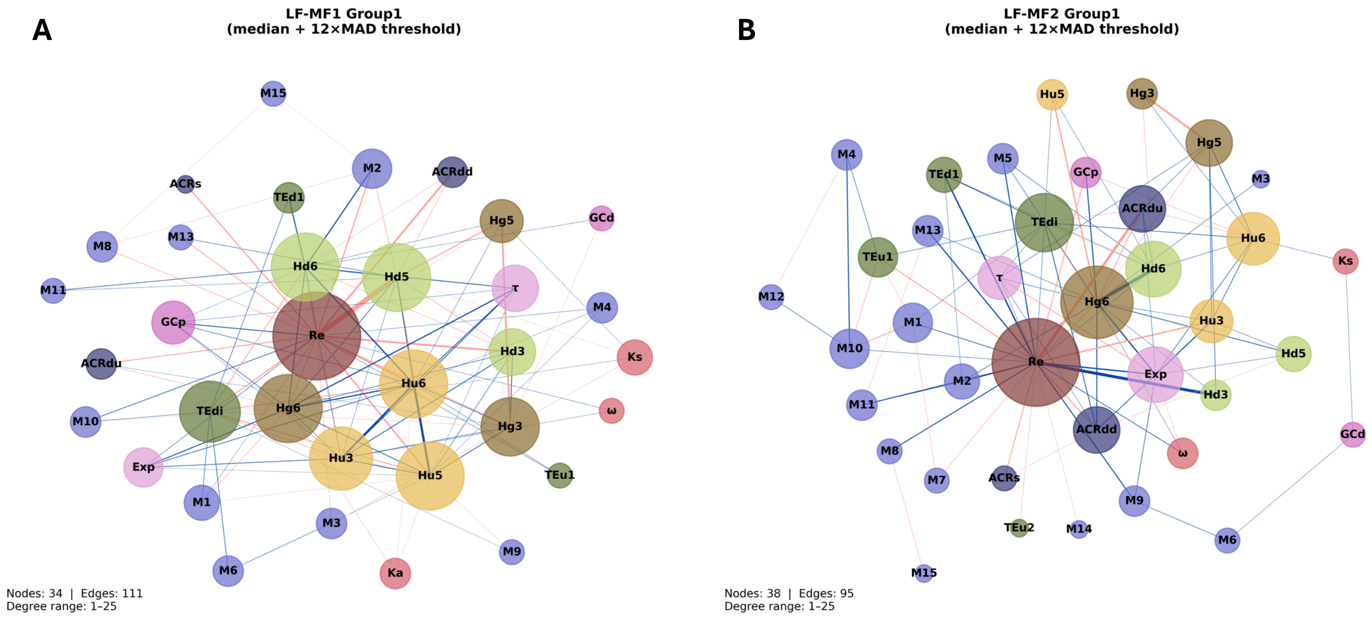


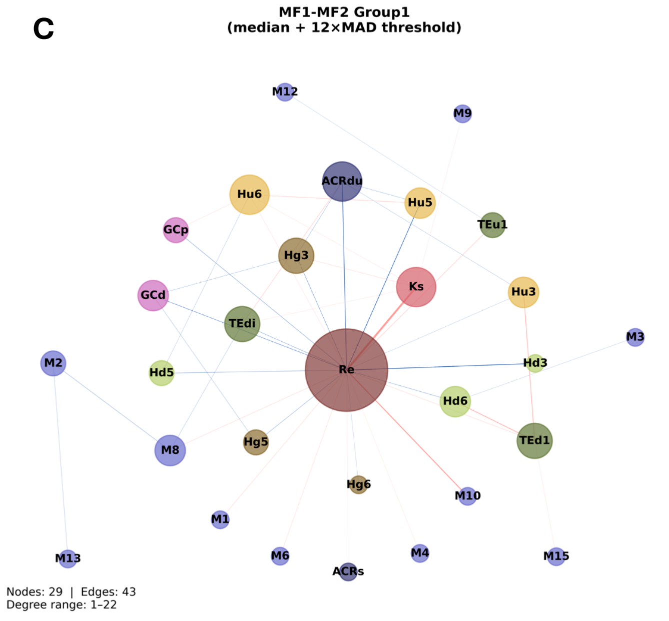


**Fig. S79. SHAP feature interaction networks for *B. rapa* group1.** Interaction networks shown for the LF-MF1 (**A**), LF-MF2 (**B**), MF1-MF2 (**C**) group1 datasets. SHAP interaction values quantify the degree to which pairs of features jointly influence model predictions beyond what is expected from their individual contributions alone. For each feature pair, mean Shapley interaction values were computed across all test-set samples. Edges connect pairs whose absolute mean interaction value exceeded a robust effect size threshold (median + 12 × median absolute deviation, computed from non-zero pairs within each group) and are colored by interaction direction: positive (pink) edges indicate synergistic interactions, in which the combined contribution of two features exceeds the sum of their individual effects, and negative (blue) edges indicate antagonistic interactions, in which the combined contribution is less than the sum. Node size is proportional to degree (number of retained edges) and node color indicates feature category, as in Fig. 6.

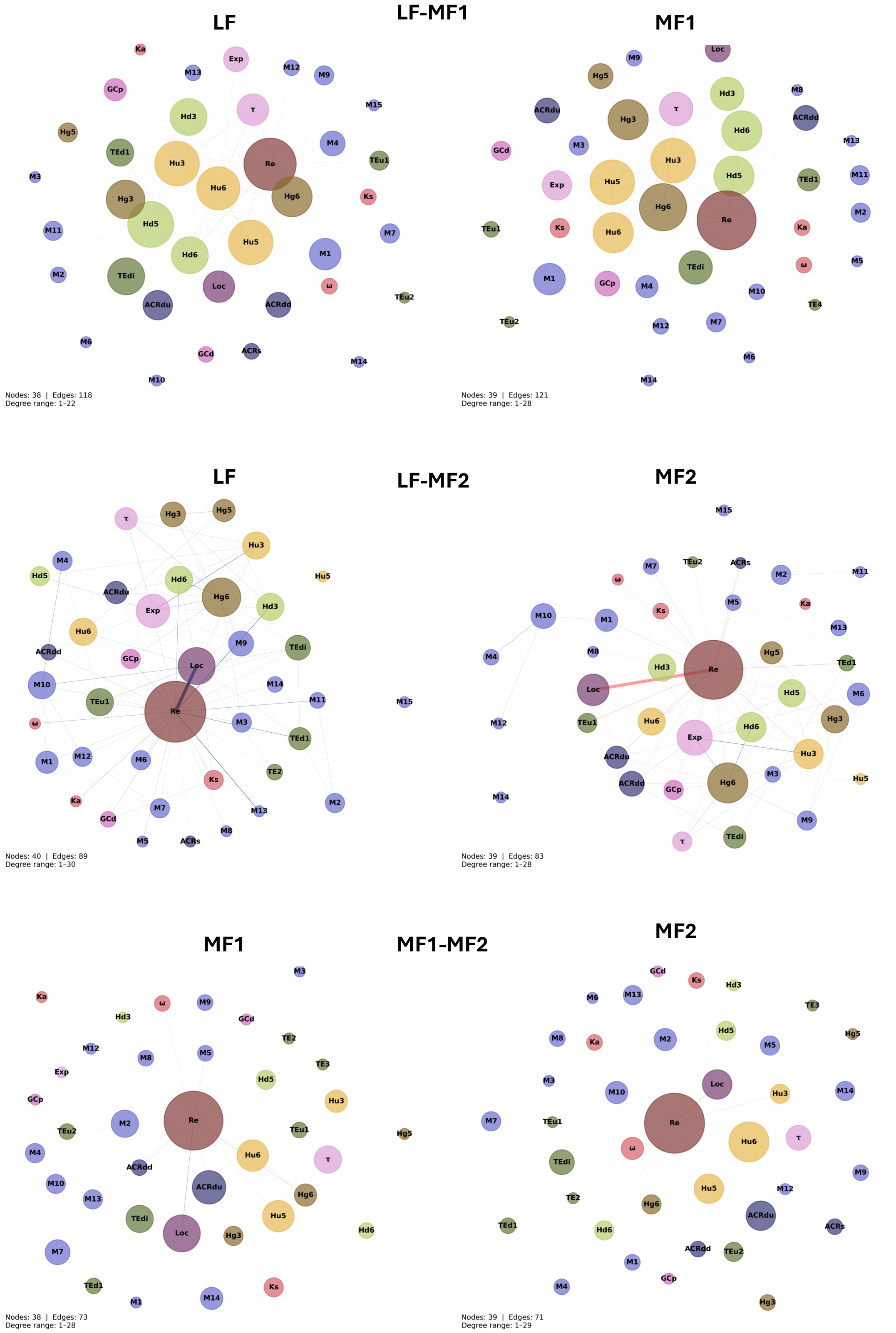


**Fig. S80. SHAP feature interaction networks for *B. rapa* per label.** Networks show pairwise Shapley interaction values computed separately for genes of the more dominant subgenome (left column) and the more non-dominant subgenome (right column). Edges connect feature pairs whose absolute mean Shapley interaction value exceeded the group-specific robust effect size threshold (median + 12 × median absolute deviation of non-zero pairs), computed from all-samples interactions and applied uniformly to both per-label subsets within each group to ensure directly comparable filtering across columns. Positive (pink) edges indicate synergistic interactions and negative (blue) edges indicate antagonistic interactions. Node size is proportional to degree and node color indicates feature category, as in Fig. 6.

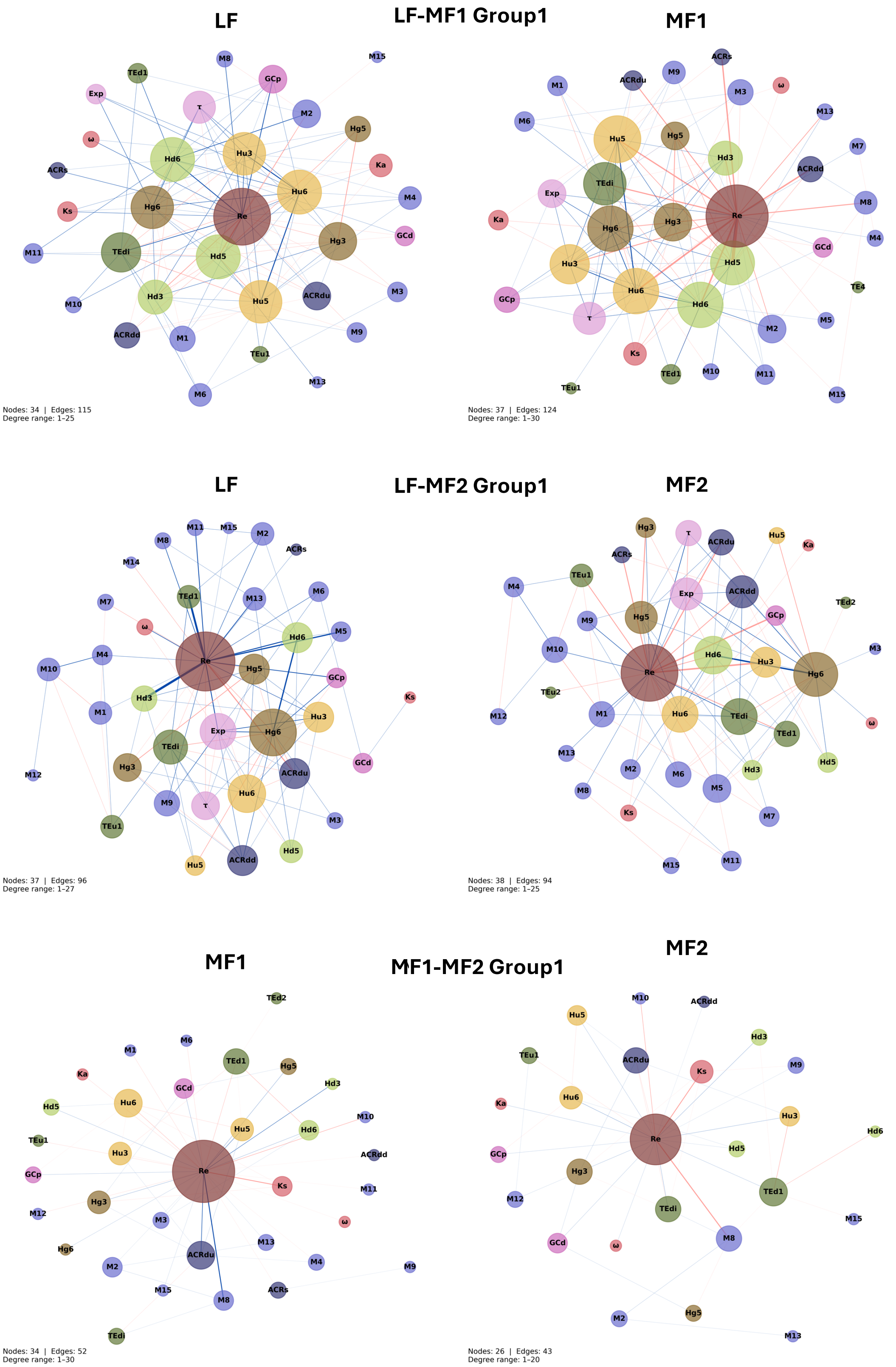


**Fig. S81. SHAP feature interaction networks for *B. rapa* group1 per label.** Networks show pairwise Shapley interaction values computed separately for genes of the more dominant subgenome (left column) and the more non-dominant subgenome (right column). Edges connect feature pairs whose absolute mean Shapley interaction value exceeded the group-specific robust effect size threshold (median + 12 × median absolute deviation of non-zero pairs), computed from all-samples interactions and applied uniformly to both per-label subsets within each group to ensure directly comparable filtering across columns. Positive (pink) edges indicate synergistic interactions and negative (blue) edges indicate antagonistic interactions. Node size is proportional to degree and node color indicates feature category, as in Fig. 6.

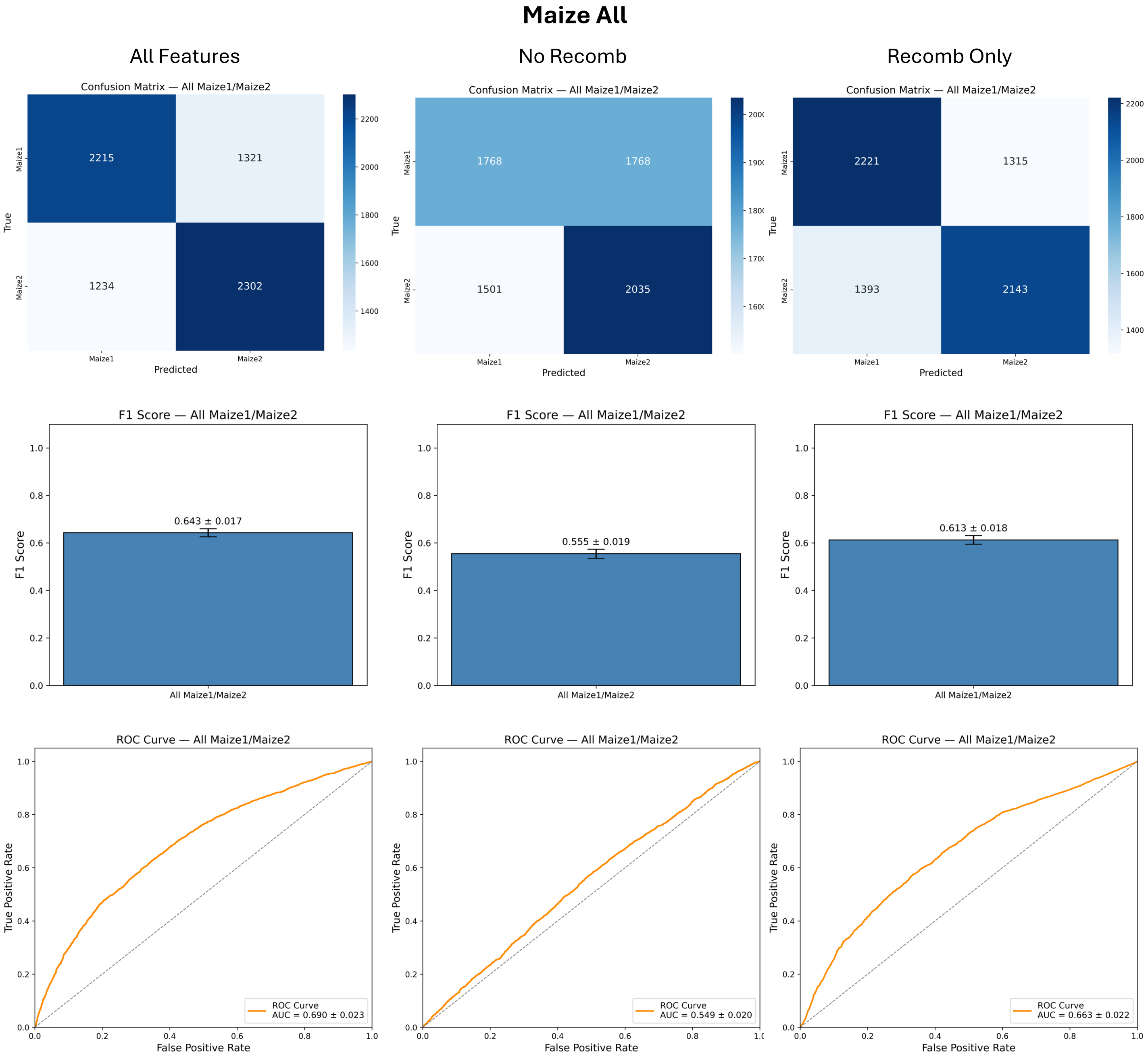


**Fig. S82. Recombination rate ablation analysis for maize1-maize2.** Classification performance of the final Optuna-tuned XGBoost model is shown under three feature conditions: all features (left), recombination rate excluded (center), and recombination rate as the sole feature (right). Each panel shows the pooled confusion matrix (top), mean F1 score ± standard deviation across 10 stratified cross-validation folds (middle), and the mean ROC curve ± standard deviation with AUC across folds (bottom). The positive class is maize2. All three conditions used identical stratified 10-fold splits and the same tuned hyperparameters; the model was not retrained under the reduced feature conditions. Confusion matrix color scales are independent across panels.

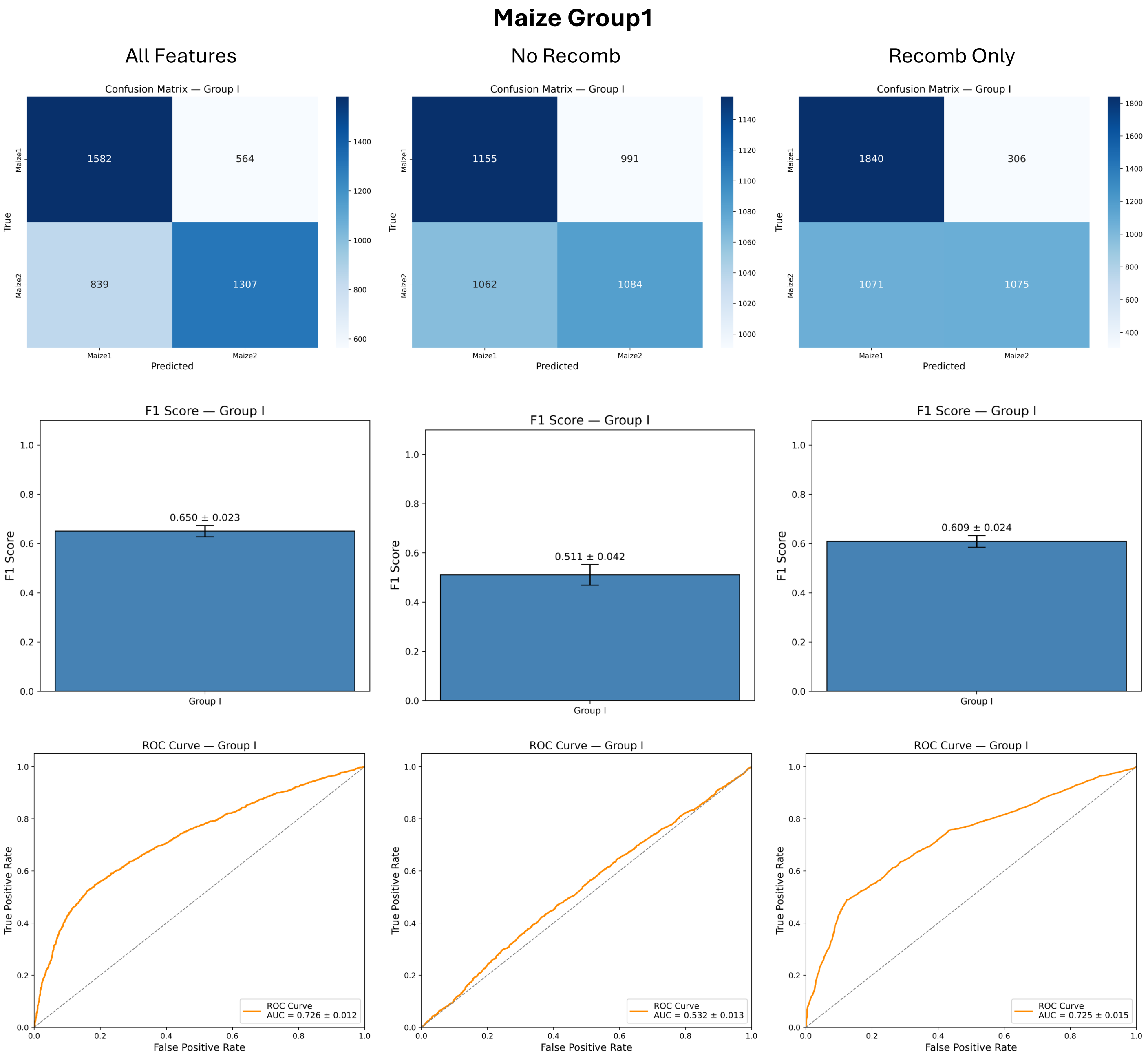


**Fig. S83. Recombination rate ablation analysis for maize group1.** Classification performance of the final Optuna-tuned XGBoost model is shown under three feature conditions: all features (left), recombination rate excluded (center), and recombination rate as the sole feature (right). Each panel shows the pooled confusion matrix (top), mean F1 score ± standard deviation across 10 stratified cross-validation folds (middle), and the mean ROC curve ± standard deviation with AUC across folds (bottom). The positive class is maize2. All three conditions used identical stratified 10-fold splits and the same tuned hyperparameters; the model was not retrained under the reduced feature conditions. Confusion matrix color scales are independent across panels.

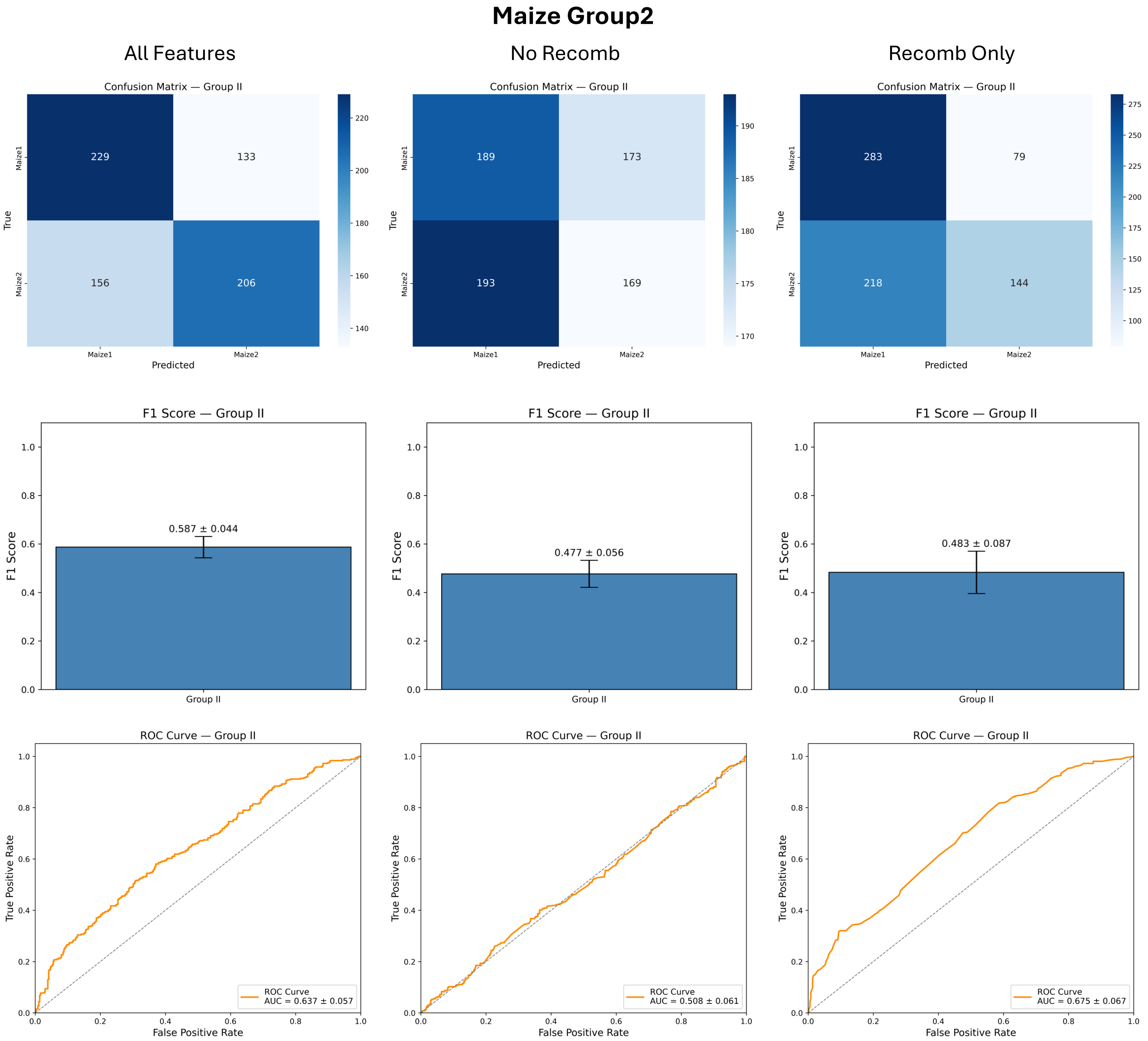


**Fig. S84. Recombination rate ablation analysis for maize group2.** Classification performance of the final Optuna-tuned XGBoost model is shown under three feature conditions: all features (left), recombination rate excluded (center), and recombination rate as the sole feature (right). Each panel shows the pooled confusion matrix (top), mean F1 score ± standard deviation across 10 stratified cross-validation folds (middle), and the mean ROC curve ± standard deviation with AUC across folds (bottom). The positive class is maize2. All three conditions used identical stratified 10-fold splits and the same tuned hyperparameters; the model was not retrained under the reduced feature conditions. Confusion matrix color scales are independent across panels.

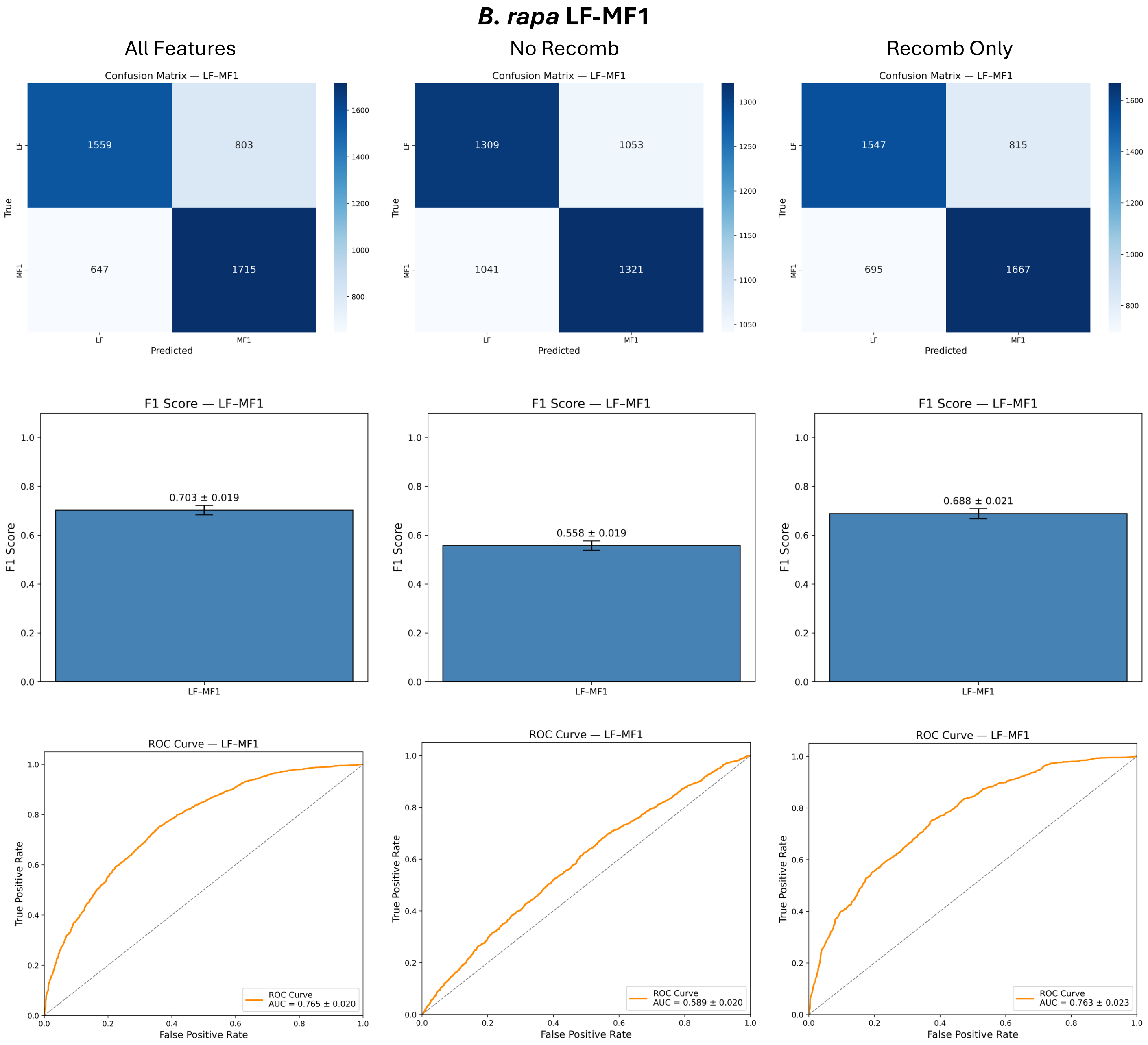


**Fig. S85. Recombination rate ablation analysis for *B. rapa* LF-MF1.** Classification performance of the final Optuna-tuned XGBoost model is shown under three feature conditions: all features (left), recombination rate excluded (center), and recombination rate as the sole feature (right). Each panel shows the pooled confusion matrix (top), mean F1 score ± standard deviation across 10 stratified cross-validation folds (middle), and the mean ROC curve ± standard deviation with AUC across folds (bottom). The positive class is MF1. All three conditions used identical stratified 10-fold splits and the same tuned hyperparameters; the model was not retrained under the reduced feature conditions. Confusion matrix color scales are independent across panels.

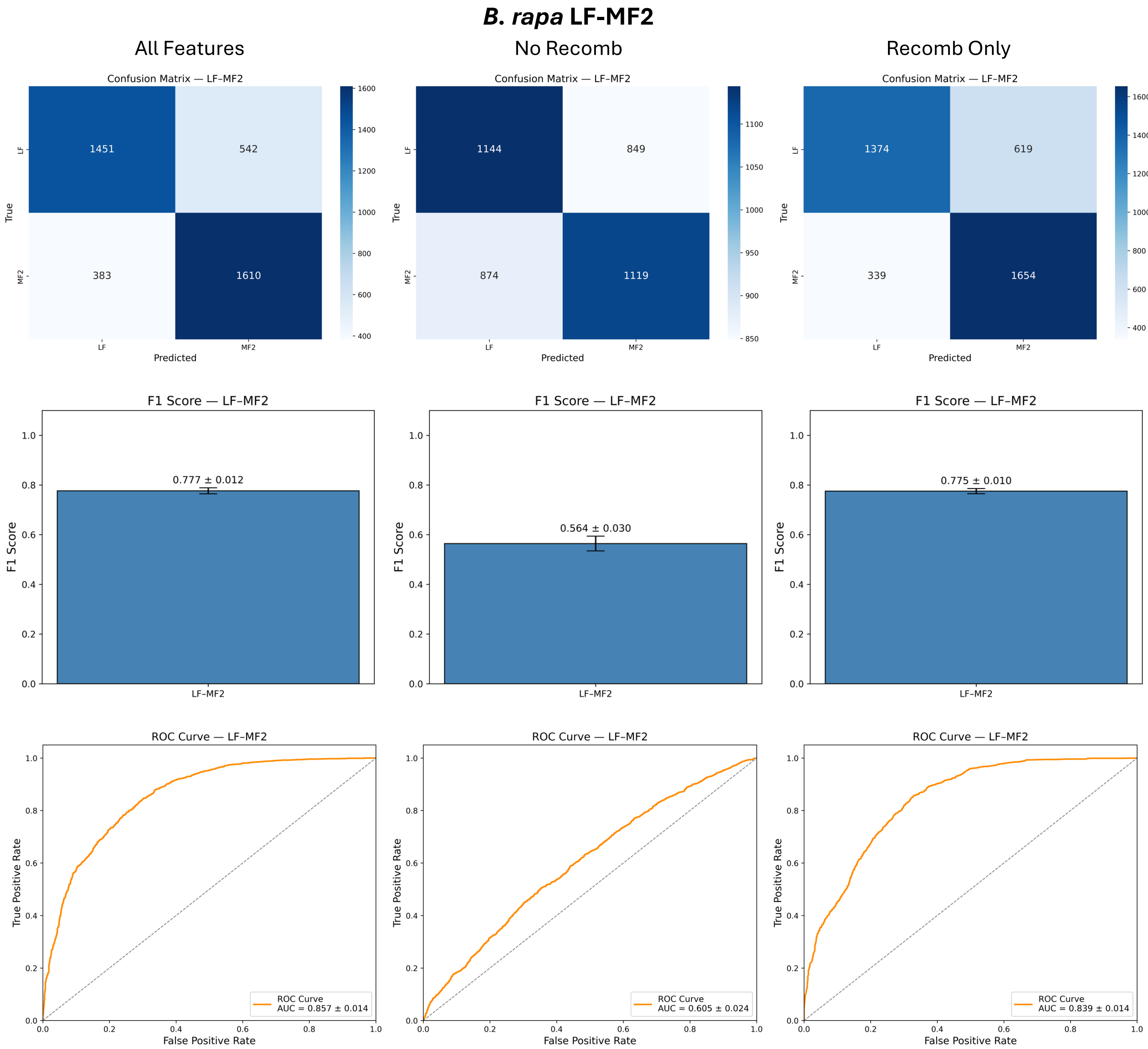


**Fig. S86. Recombination rate ablation analysis for *B. rapa* LF-MF2.** Classification performance of the final Optuna-tuned XGBoost model is shown under three feature conditions: all features (left), recombination rate excluded (center), and recombination rate as the sole feature (right). Each panel shows the pooled confusion matrix (top), mean F1 score ± standard deviation across 10 stratified cross-validation folds (middle), and the mean ROC curve ± standard deviation with AUC across folds (bottom). The positive class is MF2. All three conditions used identical stratified 10-fold splits and the same tuned hyperparameters; the model was not retrained under the reduced feature conditions. Confusion matrix color scales are independent across panels.

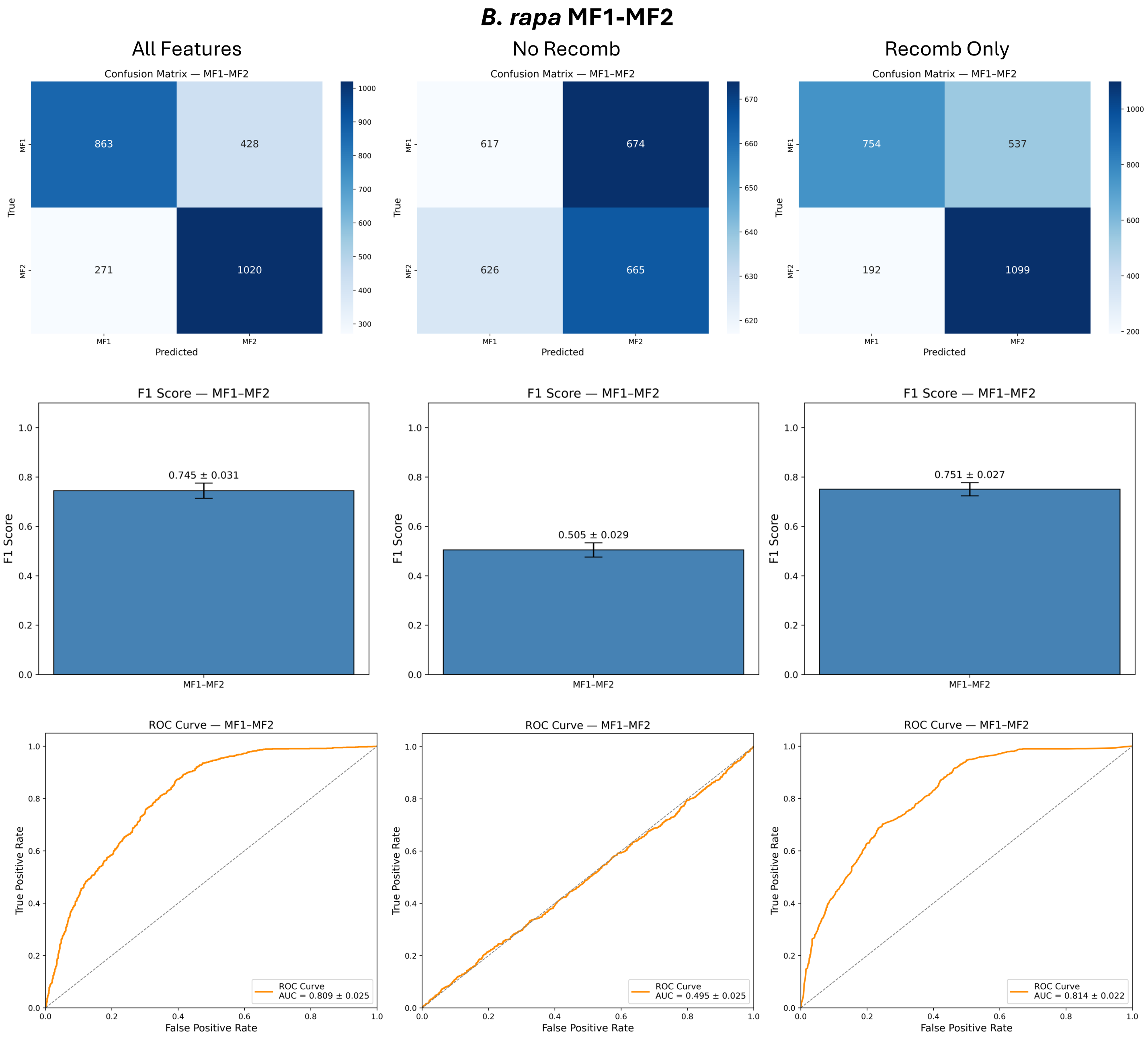


**Fig. S87. Recombination rate ablation analysis for *B. rapa* MF1-MF2.** Classification performance of the final Optuna-tuned XGBoost model is shown under three feature conditions: all features (left), recombination rate excluded (center), and recombination rate as the sole feature (right). Each panel shows the pooled confusion matrix (top), mean F1 score ± standard deviation across 10 stratified cross-validation folds (middle), and the mean ROC curve ± standard deviation with AUC across folds (bottom). The positive class is MF2. All three conditions used identical stratified 10-fold splits and the same tuned hyperparameters; the model was not retrained under the reduced feature conditions. Confusion matrix color scales are independent across panels.

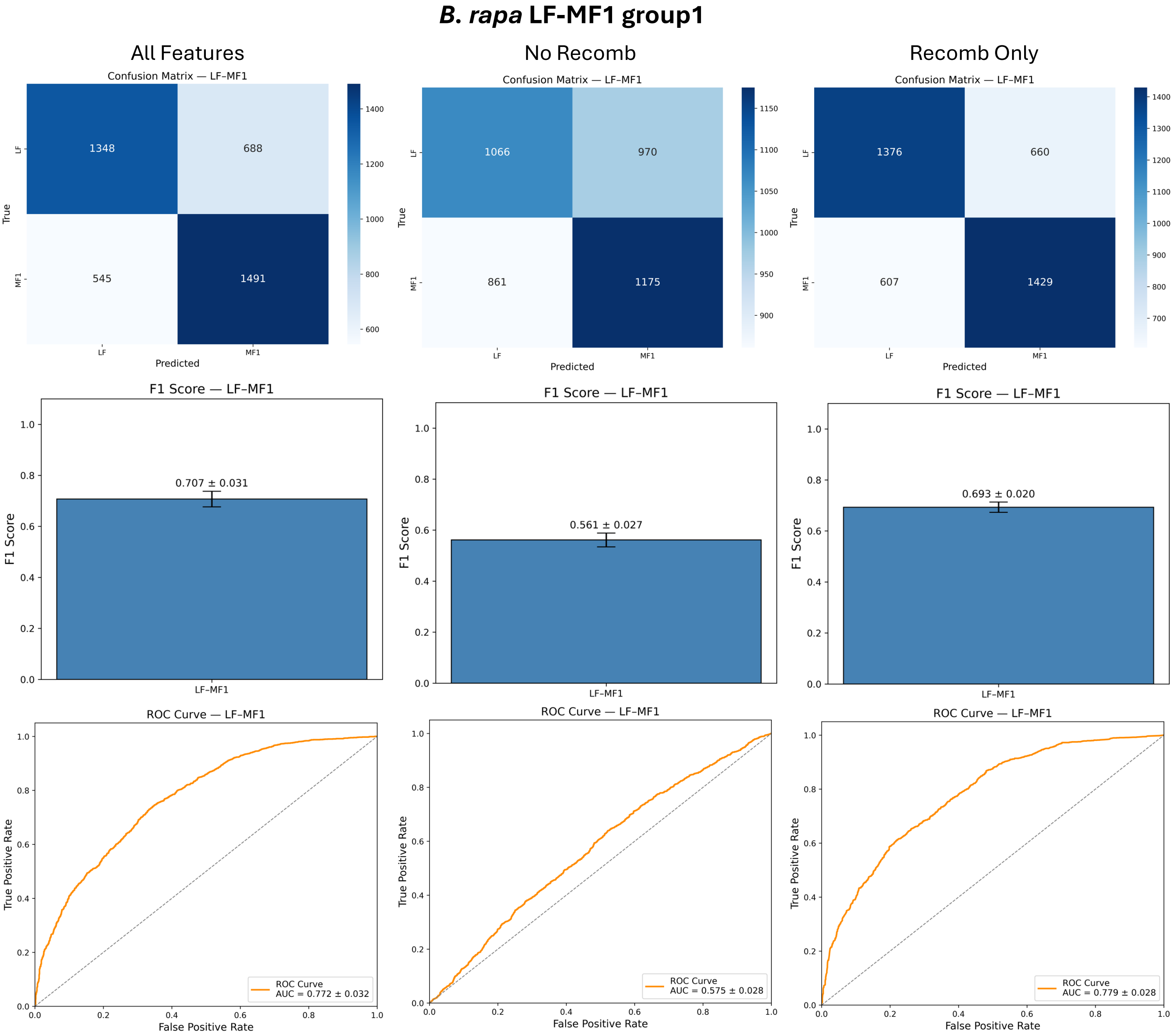


**Fig. S88. Recombination rate ablation analysis for *B. rapa* LF-MF1 group1.** Classification performance of the final Optuna-tuned XGBoost model is shown under three feature conditions: all features (left), recombination rate excluded (center), and recombination rate as the sole feature (right). Each panel shows the pooled confusion matrix (top), mean F1 score ± standard deviation across 10 stratified cross-validation folds (middle), and the mean ROC curve ± standard deviation with AUC across folds (bottom). The positive class is MF1. All three conditions used identical stratified 10-fold splits and the same tuned hyperparameters; the model was not retrained under the reduced feature conditions. Confusion matrix color scales are independent across panels.

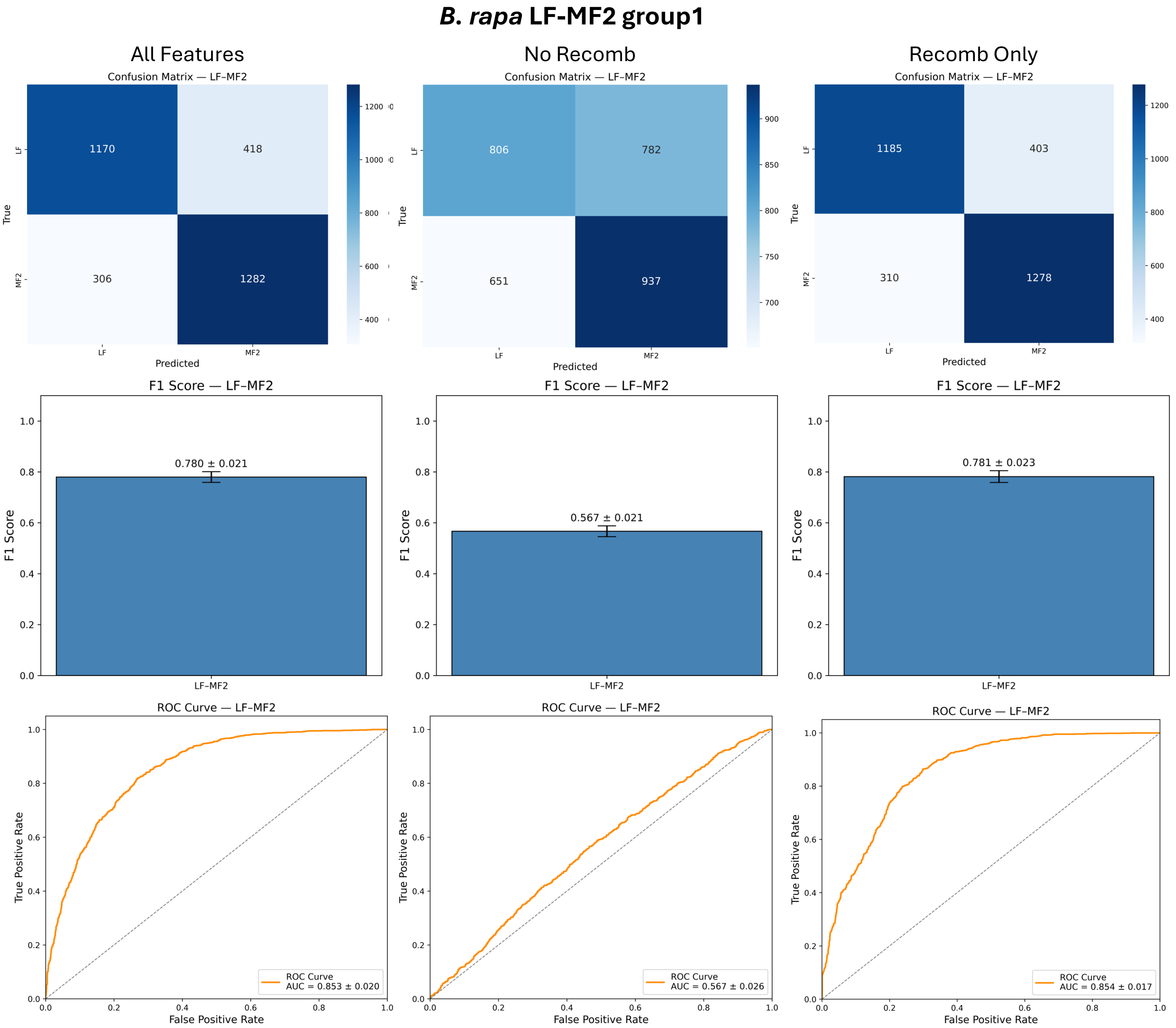


**Fig. S89. Recombination rate ablation analysis for *B. rapa* LF-MF2 group1.** Classification performance of the final Optuna-tuned XGBoost model is shown under three feature conditions: all features (left), recombination rate excluded (center), and recombination rate as the sole feature (right). Each panel shows the pooled confusion matrix (top), mean F1 score ± standard deviation across 10 stratified cross-validation folds (middle), and the mean ROC curve ± standard deviation with AUC across folds (bottom). The positive class is MF2. All three conditions used identical stratified 10-fold splits and the same tuned hyperparameters; the model was not retrained under the reduced feature conditions. Confusion matrix color scales are independent across panels.

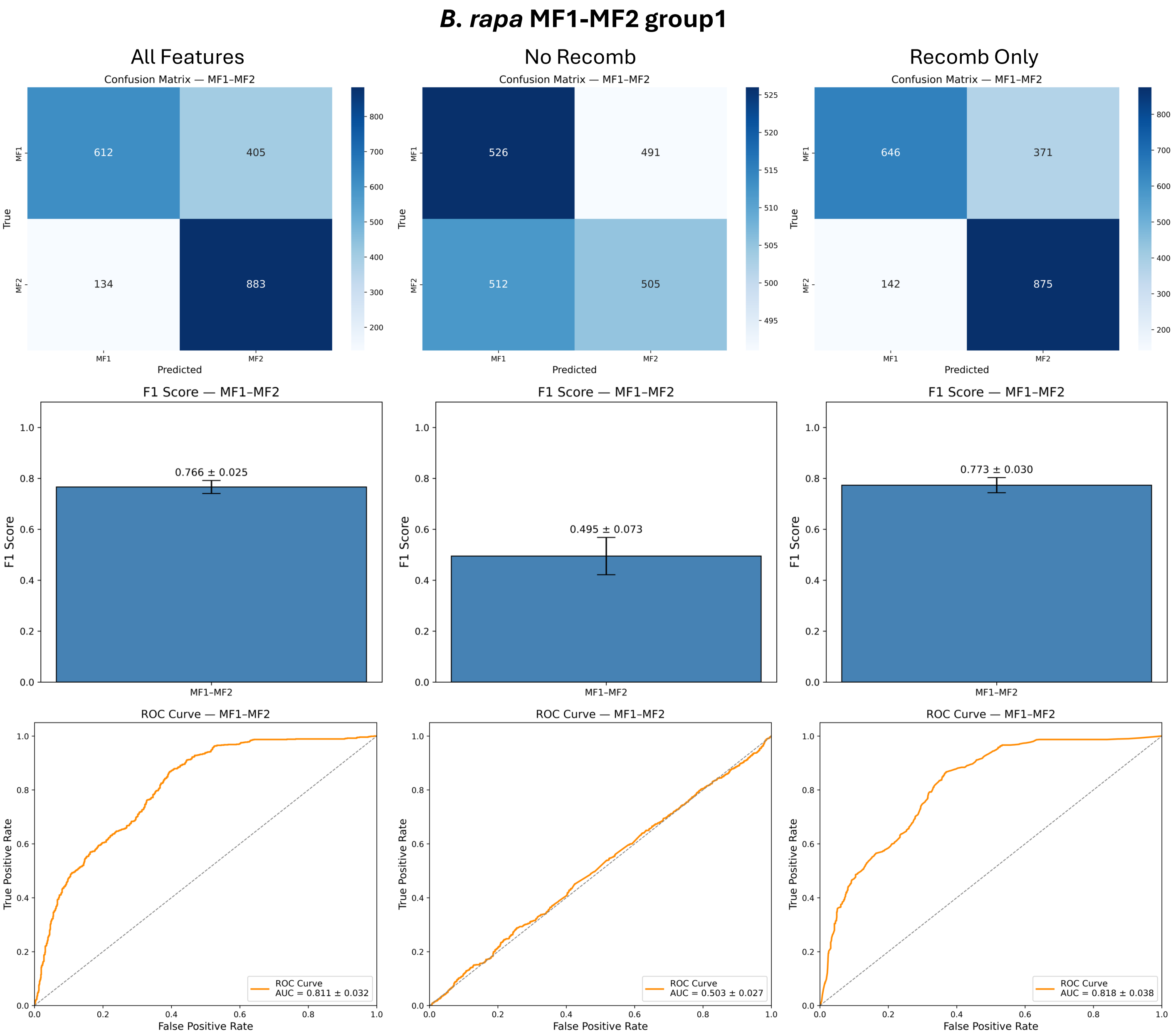


**Fig. S90. Recombination rate ablation analysis for *B. rapa* MF1-MF2 group1.** Classification performance of the final Optuna-tuned XGBoost model is shown under three feature conditions: all features (left), recombination rate excluded (center), and recombination rate as the sole feature (right). Each panel shows the pooled confusion matrix (top), mean F1 score ± standard deviation across 10 stratified cross-validation folds (middle), and the mean ROC curve ± standard deviation with AUC across folds (bottom). The positive class is MF2. All three conditions used identical stratified 10-fold splits and the same tuned hyperparameters; the model was not retrained under the reduced feature conditions. Confusion matrix color scales are independent across panels.

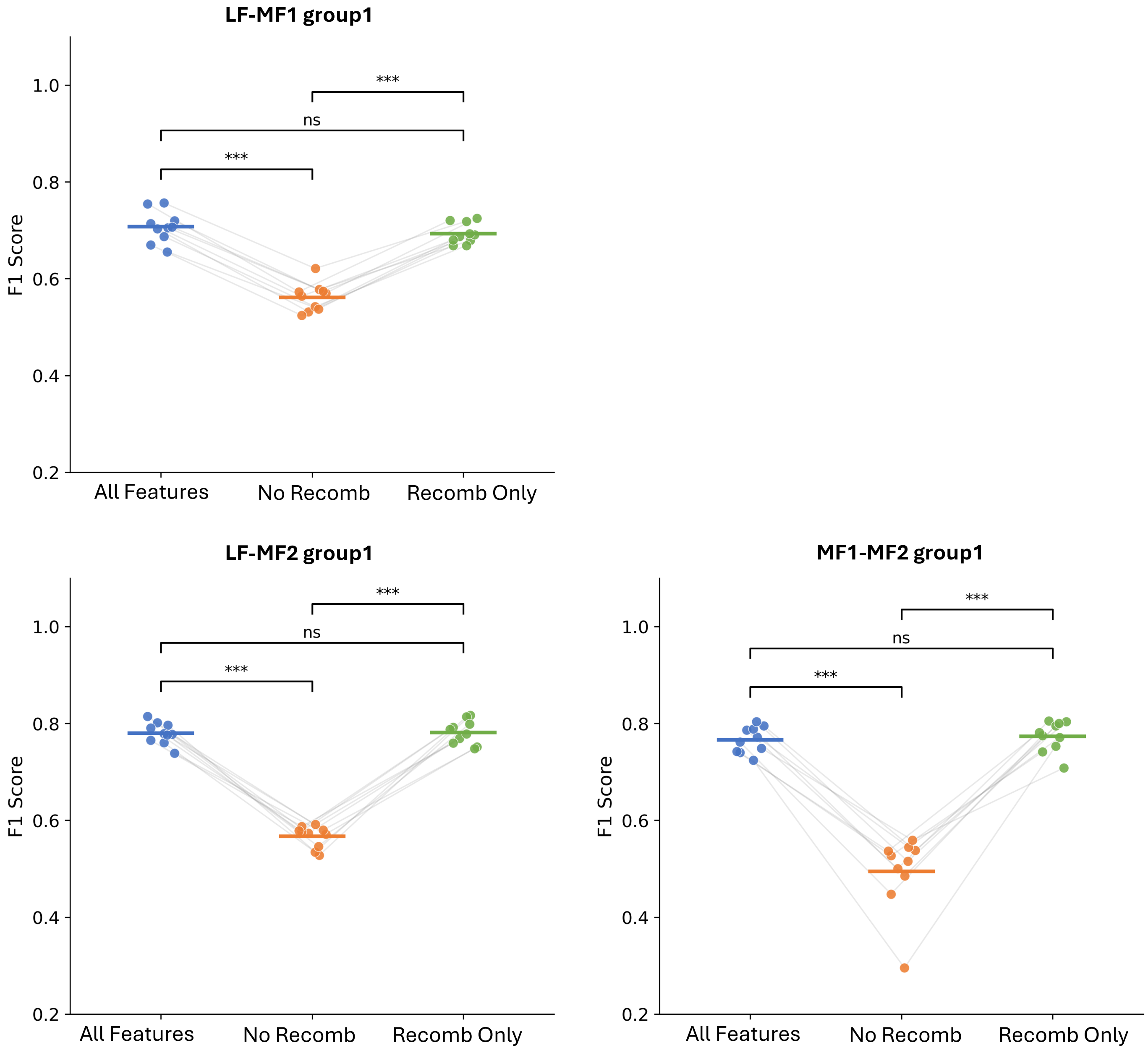


**Fig. S91. Per-fold F1 score comparison across ablation conditions for *B. rapa* group1.** F1 scores from each of the 10 stratified CV folds are shown for three feature conditions: all features (blue), recombination rate excluded (orange), and recombination rate as the sole feature (green). Each point represents one fold; gray lines connect matching folds across conditions to make the paired structure explicit. The horizontal bar within each condition indicates the mean F1 across folds. The same Optuna-tuned XGBoost hyperparameters and identical fold splits were used across all three conditions; models was not retrained under reduced feature conditions. Pairwise differences in F1 between conditions were assessed using two-sided paired t-tests on the 10 per-fold scores. Benjamini-Hochberg FDR correction was applied within each group independently (three tests: All Features vs. No Recomb, All Features vs. Recomb Only, No Recomb vs. Recomb Only; * $P < 0.05$, ** $P < 0.01$, *** $P < 0.001$).

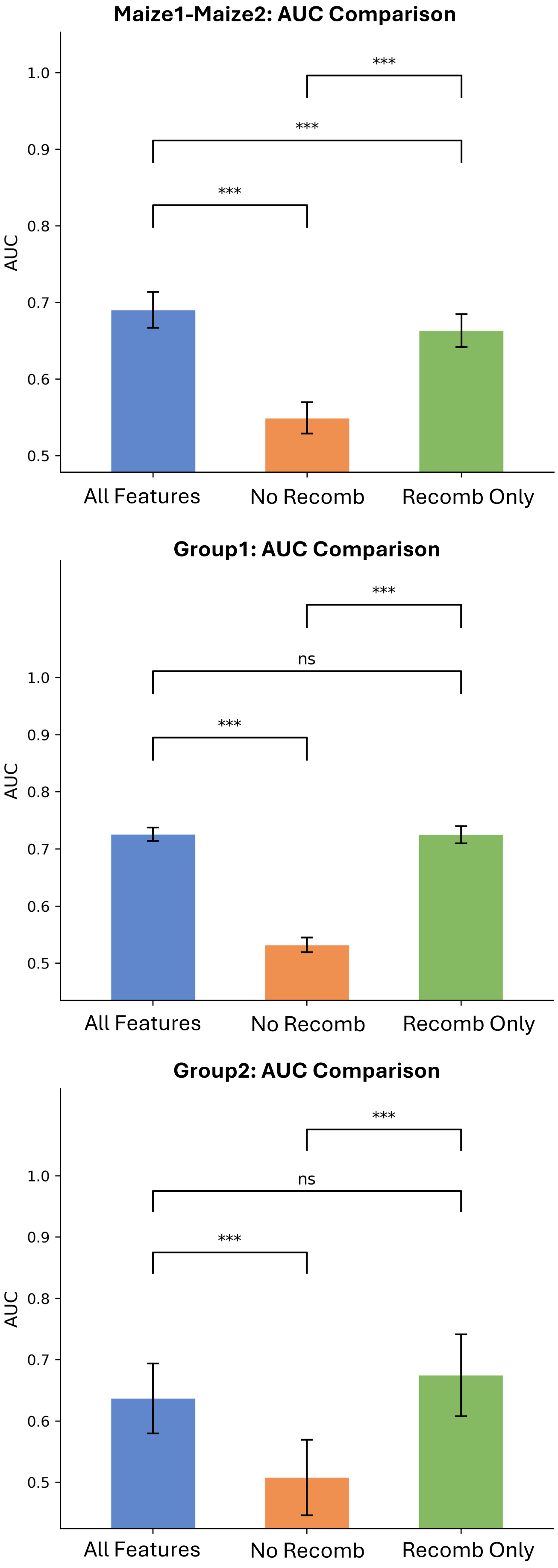


**Fig. S92. AUC comparison across ablation conditions for maize.** Mean AUC ± standard deviation across 10 stratified cross-validation folds is shown for three feature conditions: all features (blue), recombination rate excluded (orange), and recombination rate as the sole feature (green). The same Optuna-tuned XGBoost hyperparameters and identical fold splits were used across all three conditions; the model was not retrained under reduced feature conditions. Pairwise differences in AUC between conditions were assessed using the paired DeLong test applied to predicted probabilities pooled across all test-fold samples. Benjamini-Hochberg false discovery rate correction was applied within each group independently (three tests: All Features vs. No Recomb, All Features vs. Recomb Only, No Recomb vs. Recomb Only; * $P < 0.05$, ** $P < 0.01$, *** $P < 0.001$).

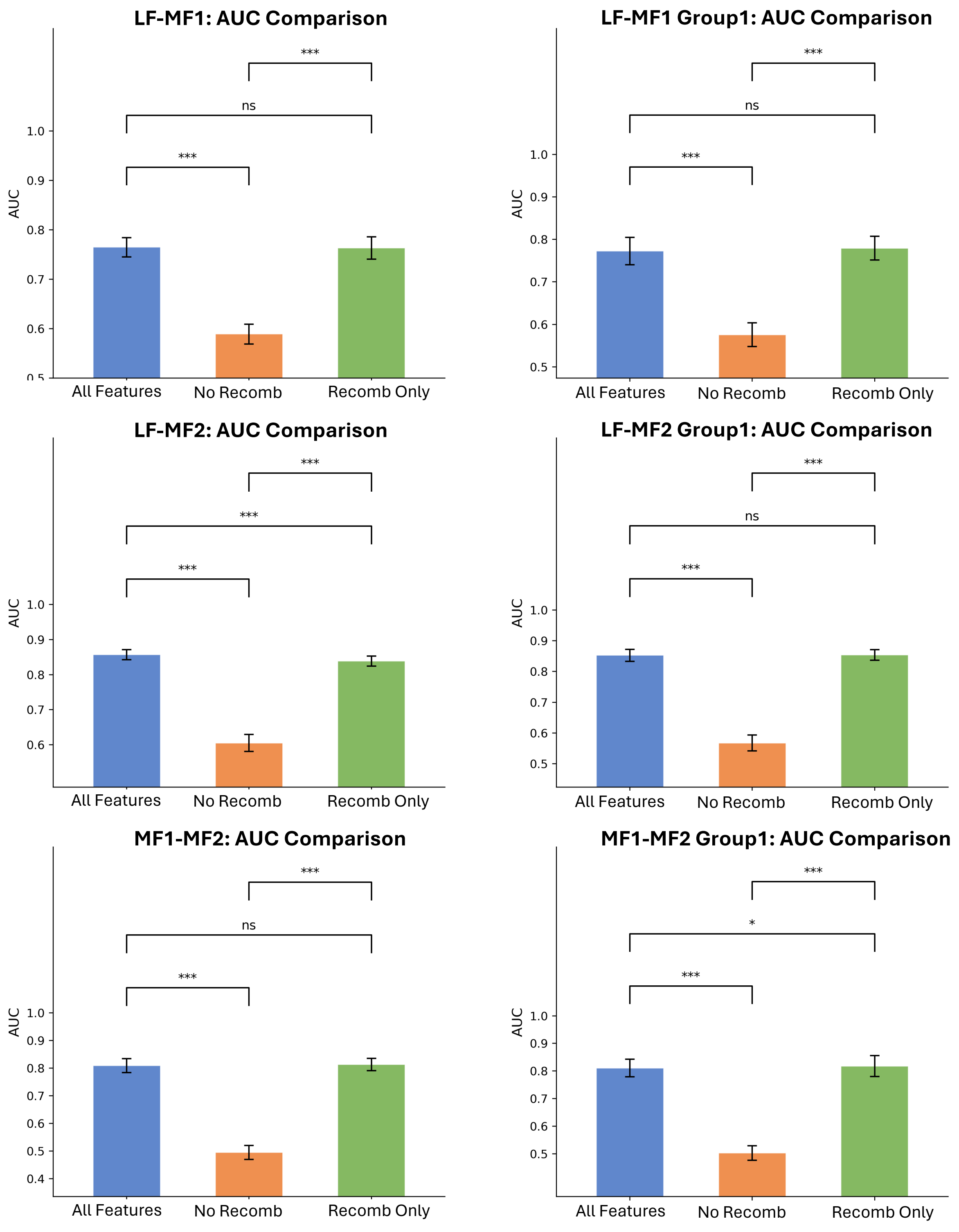


**Fig. S93. AUC comparison across ablation conditions for *B. rapa*.** Mean AUC ± standard deviation across 10 stratified cross-validation folds is shown for three feature conditions: all features (blue), recombination rate excluded (orange), and recombination rate as the sole feature (green) for full subgenome sets (left) and group1 (right). The same Optuna-tuned XGBoost hyperparameters and identical fold splits were used across all three conditions; the model was not retrained under reduced feature conditions. Pairwise differences in AUC between conditions were assessed using the paired DeLong test applied to predicted probabilities pooled across all test-fold samples. Benjamini-Hochberg false discovery rate correction was applied within each group independently (3 tests: All Features vs. No Recomb, All Features vs. Recomb Only, No Recomb vs. Recomb Only; * $P < 0.05$, ** $P < 0.01$, *** $P < 0.001$).

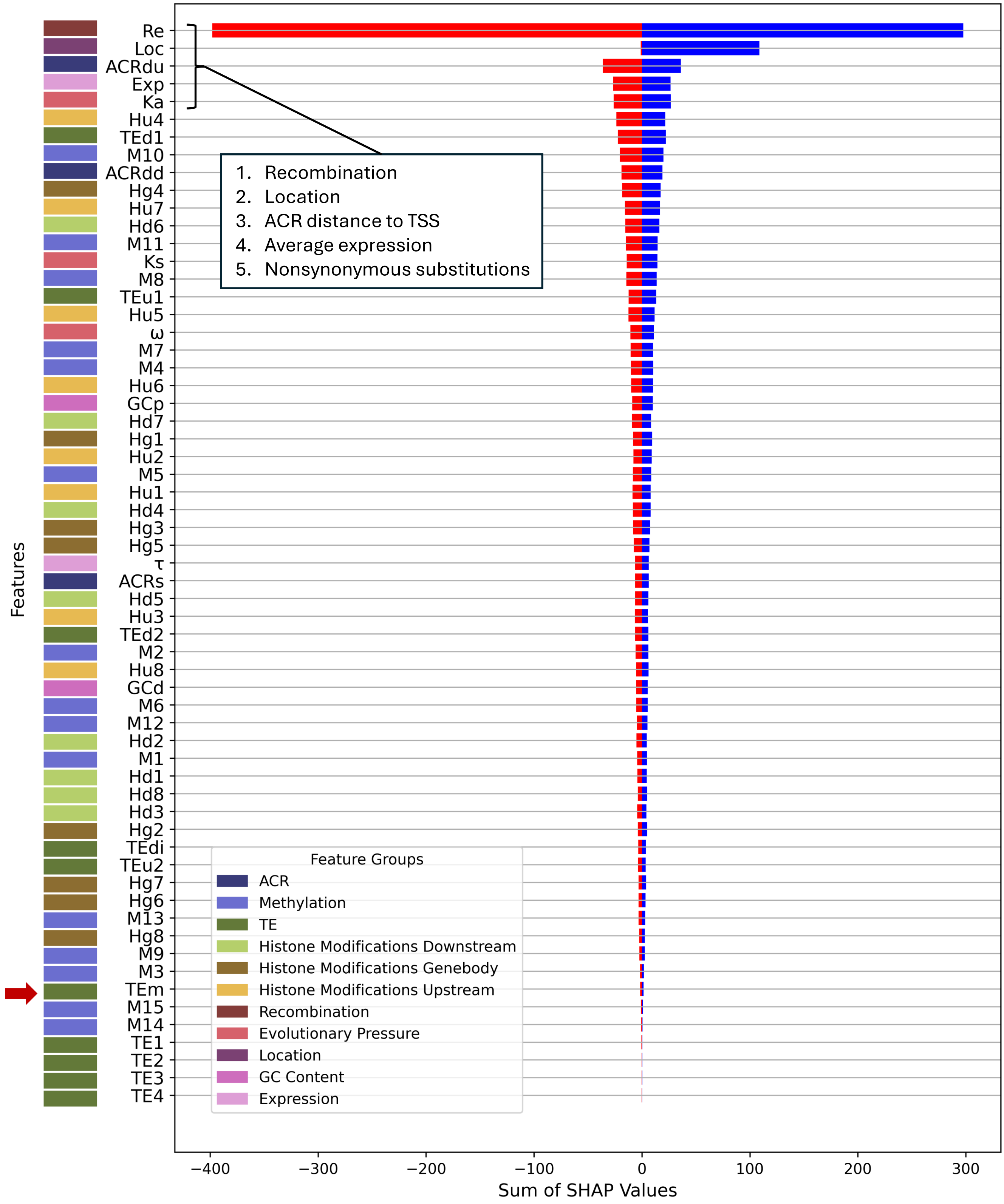


**Fig. S94. Directional SHAP feature importance profile for maize1-maize2 including MITEs.** Features including MITEs (TEm with red arrow indication) are ranked by the sum of their absolute SHAP values across all test-fold samples (ascending order, highest-importance features at top). For each feature, SHAP values are summed separately across positive contributions (blue; favoring maize2 classification) and negative contributions (red; favoring maize1 classification), reflecting total directional influence across the test set rather than per-sample magnitude. Colored tiles in the left margin indicate feature group membership. The five features contributing most to classification, as determined by summed absolute SHAP value, are listed in the inset.

**Table S1. Benchmark of unoptimized model performances across five maize subgenome datasets.**

| Model Comparison | Model* | F1 Mean ± Std | F1 Pooled | Precision Pooled | Recall Pooled | Accuracy Pooled | AUC Pooled |
|---|---|---|---|---|---|---|---|
| All | LR | 0.5482 ± 0.0249 | 0.5485 | 0.5438 | 0.5533 | 0.5452 | 0.5704 |
| | SVM | 0.5251 ± 0.0189 | 0.5253 | 0.5498 | 0.5029 | 0.5462 | 0.5649 |
| | RF | 0.5895 ± 0.0161 | 0.5894 | 0.6113 | 0.5691 | 0.6041 | 0.639 |
| | XGB | 0.6060 ± 0.0152 | 0.606 | 0.6146 | 0.5977 | 0.612 | 0.6569 |
| Group1 | LR | 0.5384 ± 0.0198 | 0.5386 | 0.5473 | 0.5302 | 0.5464 | 0.5763 |
| | SVM | 0.5375 ± 0.0243 | 0.5375 | 0.5695 | 0.509 | 0.5627 | 0.59 |
| | RF | 0.6311 ± 0.0147 | 0.6311 | 0.691 | 0.5807 | 0.661 | 0.6944 |
| | XGB | 0.6433 ± 0.0205 | 0.6435 | 0.6689 | 0.6199 | 0.657 | 0.7064 |
| Group2 | LR | 0.5301 ± 0.0435 | 0.531 | 0.5144 | 0.5487 | 0.5146 | 0.5022 |
| | SVM | 0.3353 ± 0.1713 | 0.3706 | 0.4625 | 0.3092 | 0.4742 | 0.4671 |
| | RF | 0.5201 ± 0.0919 | 0.5263 | 0.5538 | 0.5014 | 0.5481 | 0.5788 |
| | XGB | 0.5948 ± 0.0667 | 0.5958 | 0.5983 | 0.5933 | 0.5969 | 0.6418 |
| Group3 | LR | 0.8332 ± 0.0303 | 0.8329 | 0.8114 | 0.8556 | 0.8291 | 0.8977 |
| | SVM | 0.7360 ± 0.0474 | 0.7375 | 0.7237 | 0.7518 | 0.7336 | 0.8162 |
| | RF | 0.8393 ± 0.0256 | 0.8393 | 0.8371 | 0.8415 | 0.8396 | 0.8935 |
| | XGB | 0.8121 ± 0.0243 | 0.8126 | 0.8282 | 0.7975 | 0.8168 | 0.9035 |
| Group4 | LR | 0.8053 ± 0.0395 | 0.8061 | 0.8504 | 0.7663 | 0.8157 | 0.9083 |
| | SVM | 0.6637 ± 0.0503 | 0.6642 | 0.7337 | 0.6067 | 0.6933 | 0.7681 |
| | RF | 0.8396 ± 0.0252 | 0.8398 | 0.8756 | 0.8067 | 0.8461 | 0.9063 |
| | XGB | 0.8144 ± 0.0259 | 0.8158 | 0.8421 | 0.791 | 0.8213 | 0.9096 |

Note: Model architectures compared include logistic regression (LR), support vector machine (SVM), random forest (RF), and XGBoost (XGB). All models were evaluated on the same feature matrix using stratified 10-fold cross-validation, preserving class proportions across folds. For each fold, a fresh model instance was trained on the held-out training partition; models requiring feature scaling (LR, SVM) used a StandardScaler fit on the training fold only and applied to the test fold, preventing data leakage. F1 Mean ± Std reports the mean and standard deviation of F1 scores computed independently at each of the 10 test folds (classification threshold = 0.5). Pooled metrics (F1, Precision, Recall, Accuracy, AUC) are computed by concatenating predicted values across all 10 test folds before metric calculation, treating the full dataset as a single held-out set, providing a single aggregate performance estimate across the full sample. AUC for pooled metrics was computed from the ROC curve using continuous predicted probabilities. Per-fold F1 mean ± standard deviation, per-fold AUC mean ± standard deviation, confusion matrices, and ROC curves for each model are shown in Figure S8-S11.

**Table. S2. Optuna hyperparameter search ranges utilized for five maize and six *B. rapa* XGBoost classifier models.**

| Hyperparameter | Search Range (Maize) | Search Range (*B. rapa*) | Hyperparameter Description |
|---|---|---|---|
| learning_rate | [0.001, 0.3]* | [0.001, 0.3]* | Sampled on a log scale. Controls the contribution of each tree to the overall model. Lower values produce more robust models requiring more trees; higher values are more computationally efficient but risk overfitting. |
| n_estimators | [100, 1000] | [100, 1000] | Specifies the maximum number of gradient boosted trees. Acts as a ceiling; early stopping (patience = 30 rounds, monitored on a held-out inner validation fold) halts training earlier if validation log-loss plateaus. |
| max_depth | [3, 12][a] or [3, 8][b] | [3, 10][a] or [3, 8][b] | Sets the maximum depth of each tree. Higher values allow more complex relationships to be modelled but increase overfitting risk. Upper bound is tier-dependent (see footnotes). |
| subsample | [0.5, 1.0] | [0.5, 1.0] | Controls the fraction of training samples randomly selected for each tree. Lower values increase diversity across the ensemble but risk underfitting. |
| colsample_bytree | [0.5, 1.0] | [0.5, 1.0] | Sets the fraction of features randomly sampled for each tree during training. Encourages the model to rely on different feature subsets across trees to reduce overfitting. |
| min_child_weight | [1, 10] | [1, 15] | Controls the minimum sum of instance weights (Hessian) required to make a split. Higher values make the model more conservative; lower values allow more complex trees. Upper bound is higher for B. rapa to provide additional regularisation headroom for smaller training sets. |
| gamma | [0, 5] | [0, 5] | Minimum loss reduction required to make a split. Controls tree complexity by pruning splits that do not sufficiently reduce loss. |
| reg_alpha | [0, 5] | [0, 5] | L1 regularisation term on leaf weights. Encourages sparsity by driving some feature weights toward zero. |
| reg_lambda | [0.1, 15] | [0.1, 15] | L2 regularisation term on leaf weights. Penalises the sum of squared weights to reduce model complexity. |

* learning_rate was sampled on a log scale.

[a] Maize: max_depth ceiling of 12 for All and group1 models (large training sets, ~6,350 and ~3,860 samples per fold); ceiling of 8 for groups 2, 3, and 4 (small training sets, ~500-940 samples per fold) to reduce overfitting risk.

*B. rapa*: max_depth ceiling of 10 for LF-MF1 All, LF-MF1 group1, and LF-MF2 All models (large tier, ~4,250-4,650 samples per fold); ceiling of 8 for LF-MF2 group1, MF1-MF2 All, and MF1-MF2 group1 models (small tier, ~2,300-2,860 samples per fold).

[b] Small-tier ceiling of 8 is shared between maize and *B. rapa*.

**Table. S3. Best model hyperparameter values for five maize classifier models.**

| Model | All | Group1 | Group2 | Group3 | Group4 |
|---|---|---|---|---|---|
| learning_rate | 0.001139 | 0.1996 | 0.117 | 0.002412 | 0.2366 |
| n_estimators | 424 | 137 | 553 | 627 | 804 |
| max_depth | 9 | 12 | 8 | 6 | 7 |
| subsample | 0.8326 | 0.9844 | 0.9708 | 0.6229 | 0.8879 |
| colsample_bytree | 0.5349 | 0.5892 | 0.7040 | 0.6055 | 0.6764 |
| min_child_weight | 4 | 5 | 7 | 1 | 1 |
| gamma | 1.342 | 3.908 | 3.344 | 1.942 | 3.464 |
| reg_alpha | 3.628 | 2.568 | 2.325 | 2.741 | 4.124 |
| reg_lambda | 9.852 | 7.342 | 7.964 | 13.71 | 13.36 |
| Best F1 (CV) | 0.6461 | 0.6568 | 0.6206 | 0.8521 | 0.8525 |

Best F1: mean binary F1 (pos_label = 1, Maize2/WGD=1) across 10 stratified cross-validation folds, evaluated at the best Optuna trial.
All models: 300 trials (All, group1) or 500 trials (groups2–4).
Early stopping: 30 rounds on held-out inner validation fold.

**Table S4. Benchmark of unoptimized model performances across across six *B. rapa* subgenome datasets.**

| Model Comparison | Model* | F1 Mean ± Std | F1 Pooled | Precision Pooled | Recall Pooled | Accuracy Pooled | AUC Pooled |
|---|---|---|---|---|---|---|---|
| **LF-MF1** | LR | 0.5557 ± 0.0263 | 0.556 | 0.5662 | 0.5462 | 0.5655 | 0.5935 |
| | SVM | 0.5597 ± 0.0291 | 0.5601 | 0.5799 | 0.5415 | 0.5762 | 0.6017 |
| | RF | 0.5977 ± 0.0195 | 0.5978 | 0.6224 | 0.5751 | 0.6145 | 0.6782 |
| | XGB | 0.6818 ± 0.0256 | 0.6818 | 0.6793 | 0.6843 | 0.6818 | 0.7597 |
| **LF-MF1 Group I** | LR | 0.5595 ± 0.0166 | 0.5598 | 0.5549 | 0.5649 | 0.5575 | 0.5803 |
| | SVM | 0.5531 ± 0.0153 | 0.5532 | 0.5723 | 0.5354 | 0.5692 | 0.5848 |
| | RF | 0.6183 ± 0.0309 | 0.6186 | 0.641 | 0.5978 | 0.6329 | 0.6941 |
| | XGB | 0.6726 ± 0.0215 | 0.6729 | 0.6663 | 0.6796 | 0.6709 | 0.7477 |
| **LF-MF2** | LR | 0.5681 ± 0.0225 | 0.5682 | 0.5913 | 0.5468 | 0.5857 | 0.6296 |
| | SVM | 0.5910 ± 0.0224 | 0.591 | 0.5967 | 0.5855 | 0.5961 | 0.6363 |
| | RF | 0.7326 ± 0.0212 | 0.7327 | 0.7009 | 0.7675 | 0.7208 | 0.7996 |
| | XGB | 0.7591 ± 0.0198 | 0.7593 | 0.7441 | 0.7752 | 0.7551 | 0.8395 |
| **LF-MF2 Group I** | LR | 0.5604 ± 0.0238 | 0.5607 | 0.5605 | 0.5609 | 0.5618 | 0.5858 |
| | SVM | 0.5748 ± 0.0298 | 0.5756 | 0.5777 | 0.5736 | 0.5783 | 0.6059 |
| | RF | 0.7234 ± 0.0339 | 0.7237 | 0.7246 | 0.7228 | 0.7248 | 0.8055 |
| | XGB | 0.7675 ± 0.0211 | 0.7677 | 0.7505 | 0.7859 | 0.7629 | 0.8406 |
| **MF1-MF2** | LR | 0.5398 ± 0.0224 | 0.5398 | 0.5196 | 0.5617 | 0.5212 | 0.5439 |
| | SVM | 0.5407 ± 0.0258 | 0.5411 | 0.5465 | 0.5357 | 0.5456 | 0.5793 |
| | RF | 0.6925 ± 0.0201 | 0.6929 | 0.6508 | 0.7408 | 0.6716 | 0.7498 |
| | XGB | 0.7094 ± 0.0352 | 0.7099 | 0.6904 | 0.7306 | 0.7015 | 0.7864 |
| **MF1-MF2 Group I** | LR | 0.5460 ± 0.0449 | 0.5479 | 0.5303 | 0.5667 | 0.5317 | 0.5588 |
| | SVM | 0.5604 ± 0.0235 | 0.5608 | 0.544 | 0.5787 | 0.5461 | 0.5733 |
| | RF | 0.7282 ± 0.0235 | 0.7281 | 0.6581 | 0.8147 | 0.6953 | 0.756 |
| | XGB | 0.7171 ± 0.0240 | 0.7176 | 0.6854 | 0.753 | 0.7032 | 0.791 |

Note: Model architectures compared include logistic regression (LR), support vector machine (SVM), random forest (RF), and XGBoost (XGB). All models were evaluated on the same feature matrix using stratified 10-fold cross-validation, preserving class proportions across folds. For each fold, a fresh model instance was trained on the held-out training partition; models requiring feature scaling (LR, SVM) used a StandardScaler fit on the training fold only and applied to the test fold, preventing data leakage. F1 Mean ± Std reports the mean and standard deviation of F1 scores computed independently at each of the 10 test folds (classification threshold = 0.5). Pooled metrics (F1, Precision, Recall, Accuracy, AUC) are computed by concatenating predicted values across all 10 test folds before metric calculation, treating the full dataset as a single held-out set, providing a single aggregate performance estimate across the full sample. AUC for pooled metrics was computed from the ROC curve using continuous predicted probabilities. Per-fold F1 mean ± standard deviation, per-fold AUC mean ± standard deviation, confusion matrices, and ROC curves for each model are shown in Figure S50-S56.

**Table. S5. Best model hyperparameter values for size *B. rapa* classifier models.**

| Model | LF–MF1 All | LF–MF1 Group1 | LF–MF2 All | LF–MF2 Group1 | MF1–MF2 All | MF1–MF2 Group1 |
|---|---|---|---|---|---|---|
| learning_rate | 0.003405 | 0.002097 | 0.007705 | 0.001172 | 0.001323 | 0.005584 |
| n_estimators | 509 | 945 | 672 | 940 | 423 | 649 |
| max_depth | 9 | 10 | 10 | 8 | 8 | 4 |
| subsample | 0.6735 | 0.6841 | 0.9188 | 0.8451 | 0.9164 | 0.7956 |
| colsample_bytree | 0.6771 | 0.7799 | 0.6456 | 0.6498 | 0.7986 | 0.9688 |
| min_child_weight | 5 | 3 | 1 | 1 | 1 | 15 |
| gamma | 3.001 | 0.4984 | 3.007 | 0.07678 | 4.979 | 4.993 |
| reg_alpha | 3.123 | 3.281 | 0.4706 | 0.006539 | 0.9142 | 3.501 |
| reg_lambda | 6.668 | 14.95 | 0.3775 | 0.1138 | 1.997 | 11.58 |
| Best F1 (CV) | 0.7064 | 0.711 | 0.7843 | 0.7857 | 0.7527 | 0.7699 |

Best F1: mean binary F1 (pos_label = 1, non-dominant subgenome/WGD=1) across 10 stratified cross-validation folds, evaluated at the best Optuna trial.
LF–MF1 All, LF–MF1 group1, LF–MF2 All: 400 trials (large tier).
LF–MF2 group1, MF1–MF2 All, MF1–MF2 group1: 600 trials (small tier).
Early stopping: 30 rounds on held-out inner validation fold.
